# High-throughput discovery of regulatory effector domains in human RNA-binding proteins

**DOI:** 10.1101/2024.07.19.604317

**Authors:** Abby R. Thurm, Yaara Finkel, Cecelia Andrews, Xiangmeng S. Cai, Colette Benko, Lacramioara Bintu

## Abstract

RNA regulation plays an integral role in tuning gene expression and is controlled by thousands of RNA-binding proteins (RBPs). We develop and use a high-throughput recruitment assay (HT-RNA-Recruit) to identify regulatory domains within human RBPs by recruiting over 30,000 protein tiles from 367 RBPs to a reporter mRNA. We discover over 100 unique RNA-regulatory effectors in 86 distinct RBPs, presenting evidence that RBPs contain functionally separable domains that dictate their post-transcriptional control of gene expression, and identify some with unique activity at 5’ or 3’UTRs. We identify some domains that downregulate gene expression both when recruited to DNA and RNA, and dissect their mechanisms of regulation. Finally, we build a synthetic RNA regulator that can stably maintain gene expression at desired levels that are predictable by a mathematical model. This work serves as a resource for human RNA-regulatory effectors and expands the synthetic repertoire of RNA-based genetic control tools.

**Highlights:**

- HT-RNA-Recruit identifies hundreds of RNA-regulatory effectors in human proteins.
- Recruitment to 5’ and 3’ UTRs identifies regulatory domains unique to each position.
- Some protein domains have both transcriptional and post-transcriptional regulatory activity.
- We develop a synthetic RNA regulator and a mathematical model to describe its behavior.

## Introduction

RNA-binding proteins (RBPs) play integral roles in the coordination of mRNA fate and the post-transcriptional regulation of gene expression^1^. Previous work has annotated RNA-binding domains and measured transcriptome-wide binding profiles for many of the over 1,000 known human RBPs^2^. Subsequent large-scale studies have identified mRNA regulatory activity for hundreds of human RBPs, underscoring the importance of integrating RBP-mediated mRNA regulation in quantitative measurements and predictions of human gene expression^3,4^. Most previous studies have focused on characterizing full-length RBPs, as it remains to be determined whether human RBPs are broadly composed of distinct domains that confer their mRNA-regulatory roles.

The identification of separable regulatory or ‘effector’ domains in DNA-binding transcription factors (TFs) across species has led not only to generalizable models of TF organization, but has also enabled high-throughput screening of tens of thousands of TF effector domains for gene regulatory potential using tethering or recruitment assays^5–8^. These information-rich datasets have aided the discovery of new effector domains and for high-throughput annotation of TF regulatory activity at large scale. Recent work in yeast has demonstrated that RBPs can similarly be evaluated for mRNA regulatory capacity using large-scale tethering screens of proteomic fragments^9^. These results significantly improved the annotation of RBPs with previously unknown function and confirmed that many RBPs are functionally modular with specific regions within them that encode regulatory activity through interactions with cellular processing factors^10^. Specifically, several mRNA regulatory domains were found to overlap intrinsically disordered regions (IDRs), indicating that regions of RBP regulatory function correlate with their structural features^11^. These results demonstrate that widespread evaluation and annotation of human RBP regulatory domains is now both possible and necessary for accurate measurement of mRNA regulation.

Here, we develop a high-throughput, pooled recruitment assay (HT-RNA-Recruit) in human cells to measure RNA regulatory activity for tens of thousands of protein tiles. We use this approach to identify new RNA-regulatory effectors through unbiased tiling of over 300 human RBPs as well as testing all Pfam-annotated domains in known RBPs for regulatory activity. Furthermore, we take advantage of our synthetic system to systematically vary recruitment positioning and stoichiometry on the reporter RNA and show the strength of RNA-regulatory effectors is dependent both on which UTR they are recruited to and on the number of available binding sites on the mRNA. In addition, we use a previously developed high-throughput DNA recruitment system to compare the transcriptional and post-transcriptional regulatory capacity of a subset of RBP and TF tiles. While most effector domains regulate gene expression when recruited to either DNA or RNA, we find several that can act at both levels of control. In particular, the KRAB domain from the transcription factor ZNF10 exerts transcriptional silencing effects even when recruited via tethering to the reporter mRNA. Finally, we use one of the strongest RNA-regulatory effectors to build an inducible RNA-regulatory synthetic protein, and develop a mathematical model describing its dose-dependence and kinetic behavior over time.

## Results

### High-throughput recruitment identifies protein tiles with RNA-regulatory activity within RNA binding proteins

To develop a high-throughput recruitment assay to RNA, which we call HT-RNA-Recruit, we took advantage of the well-established MS2 hairpin sequence and its cognate protein binder, MS2 phage capsid protein (MCP). We installed 24 copies of the MS2 hairpin in the 3’UTR of the RNA of a reporter gene constitutively expressing a synthetic surface marker and fluorescent marker Citrine, which we integrated into K562 cells at the AAVS1-Safe Harbor locus on chromosome 19 (**Fig. 1A**). Libraries of 80-amino acid protein were cloned as fusions to MCP (**Fig. S1A**) and delivered via pooled lentivirus to cells expressing the MS2 reporter. Recruitment of tiles that downregulate reporter RNA levels or inhibit translation lead to lower expression of both Citrine and the surface marker (**Fig. 1A**), enabling magnetic separation of large numbers of ON (reporter-expressing) and OFF (non-expressing) cells (**Fig. S1B**).

**Figure 1.**
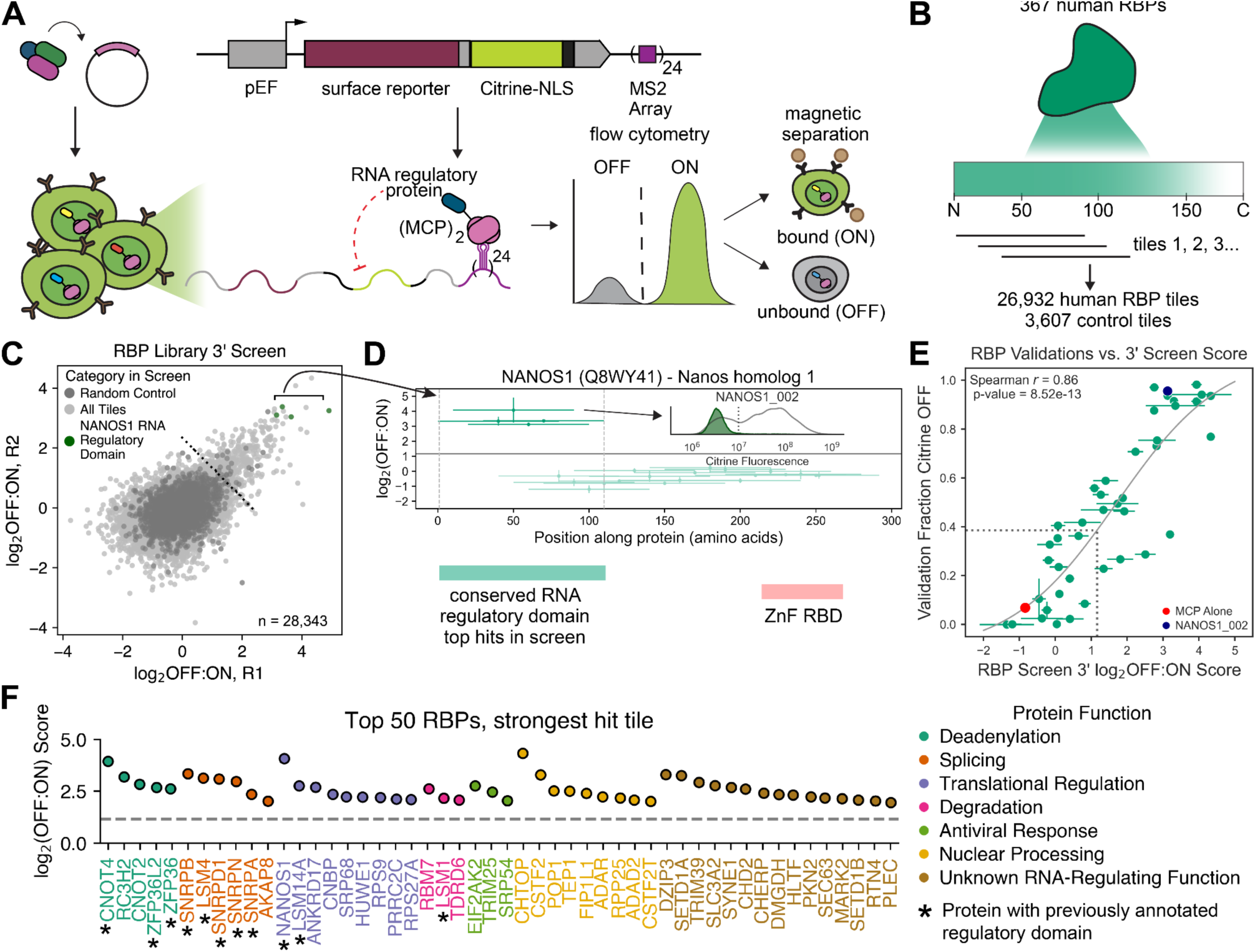
High-throughput recruitment to RNA discovers hundreds of protein tiles with RNA-regulating capabilities. (A) Overview of high-throughput RNA recruitment assay. A library of protein tiles is cloned as a pool fused to the MS2 capsid protein (MCP) dimer and delivered to cells expressing a reporter gene encoding a surface marker that enables magnetic separation of cells, a Citrine reporter gene, and 24 copies of the MS2 stem loop for 3’UTR recruitment of the protein library. After 10 days of recruitment, cells are separated into ON and OFF populations and domains are sequenced. (B) Schematic of RBP library design, which includes all possible 80 amino acid tiles for 367 human RBPs. (C) Log_2_(OFF:ON) (positive = RNA downregulated) enrichment scores plotted per replicate of the human RBP screen in K562 cells. Light gray, all members; dark gray, random controls; green, tiles from NANOS1 known RNA regulatory domain. (D) Example tiling plot of NANOS1, a known translational modulator. X-axis, position of tile along protein; y-axis, recruitment screen score. Line, length of tile; dot, tile score; vertical error bars, standard deviation of 2 biological screen replicates. The solid line represents the screen hit cutoff and dashed vertical lines represent the edges of the determined regulatory domain span. Inset: Flow cytometry plot of individual recruitment of top tile MCP-NANOS1_002 (green fill) versus MCP alone (grey line, no fill). Green box indicates location of a previously-annotated RNA-regulatory domain; red box indicates Pfam-annotated RNA-binding domain. (E) Individual validation measurements for 48 tested tiles. Grey dashed lines represent screen cutoff score (vertical) and corresponding Citrine OFF cutoff (horizontal). Red dot, MCP alone; blue dot, NANOS1_002. Error bars represent an average of two biological replicates. (F) Top 50 RBPs with tile hits in the recruitment screen, binned into categories of RNA-related function (hand-annotated by literature evidence, in **Table S1**) and ranked by top tile screen score. Starred proteins are those whose top tiles overlapped with previously reported regulatory domains. Dashed line, recruitment hit cutoff.

We used an 1100-member RBP census curated from the Pfam database and wide-scale RNA crosslinking and pulldown (eCLIP)^12^ experiments to then formulate a smaller subset of these RBPs to investigate for ‘effector’ domain activity. We first used GO term annotations to exclude RBPs with no known cellular function, those annotated to be involved in DNA-templated transcription, and enzymatic components of the ribosome and spliceosome. We also excluded RBPs involved in rRNA processing and further filtered out proteins involved in other nuclear processes like chromosome maturation. We were left with a list of 367 RBPs plus four negative control proteins (actin, albumin, tubulin, and IgG, **Table S1**). We tiled along the length of each protein in 80aa windows with a 10aa tiling window to create 25,954 unique protein tiles (**Fig. 1B**). Finally, we queried the Pfam database for any known domains in the original 1100 proteins and selected those that were <80 amino acids long (978 domains). For domains shorter than 80aa, we added neighboring sequence from the native protein on both ends to reach 80aa and keep a consistent tile size throughout the library. (**Fig. 1B**). To this, we added 3,597 tiles of random protein sequence and 10 known well-expressing tiles that have transcriptional activating or repressing activity for a total library size of 30,539 members. This library was cloned in a pool as a fusion to 3x-FLAG-tagged MCP in a lentiviral vector and delivered to reporter-expressing K562 cells in two biological replicates (**Methods**).

After 10 days of selection and recruitment, we performed magnetic separation and subsequent sequencing of the protein tiles expressed in the separated ON and OFF populations, then computed the log_2_(OFF:ON) ratio for each tile using read counts in each population (**Fig. S1B**). The screen measurements were reproducible (Spearman’s *r* = 0.36, p < 1×10^-16^), and consisted of 28,343 tiles that passed a sequencing depth threshold (**Fig. 1C, Methods**). We defined a hit threshold as three standard deviations above the mean of the random control population (**Fig. 1C**), resulting in 438 hit tiles. Four of the top hit tiles in the screen overlap a known RNA regulatory domain in the protein NANOS1^13^, showing that our high-throughput method can reliably identify protein domains that act as RNA downregulators (**Fig. 1D**). When cloned and recruited individually to the reporter RNA, the second tile of NANOS1 (aa 11-90) led to a reduction of Citrine levels in all cells (**Fig. 1D inset**). To further confirm the validity of our calculated screen enrichment scores, we tested 48 MCP-tile fusions in individual cell lines and measured the fraction of cells that had no Citrine expression (**Fig. S1C-F**). The individual Citrine measurements correlated well with the corresponding high-throughput enrichment scores, or screen scores (Spearman’s *r* = 0.85) (**Fig. 1E**, **Fig. S1C-F, Table S3**). To measure reporter RNA levels directly, we performed HCR-Flow-RNA-FISH (**Methods**) on individual cell lines made with eight top hit tiles after MCP-mediated recruitment and found that all tested tiles induced decreased RNA levels correlated with the observed decrease in Citrine fluorescence, indicating that at least this subset of tiles from the library is directly increasing RNA degradation rather than inhibiting translation (**Fig. S1G**).

We overlapped the hits in our screen with results from a previous large-scale tethering study^3^ performed using full-length RBPs. Despite the differences in study design (this work uses protein tiles, the previous work tethered full-length proteins), we observed reasonable concordance. Of the 195 proteins tested in both studies, 12 had hits in both studies, 115 were not hits in either, 36 were hits only in this study (at the tile level), and 32 were hits only in the previous work (at the full-protein level) (**Fig. S1H**). The top tile hits from the 12 overlapping proteins comprised some of the strongest hits in our screen (**Fig. S1H**). Through literature review, we manually compiled a list of the biological processes that RBPs with top hit tiles were known to participate in (**Table S1**). The top 50 RBPs with strong hit tiles in our screen spanned a wide range of known RNA-related functions, from well-known members of the CCR4-NOT deadenylation complex (CNOT4, CNOT2^14^) to those with known RNA-binding potential but with unknown function in controlling RNA fate (DZIP3, SETD1A) (**Fig. 1F**). We noted that some of these top hit tiles were located within larger, previously-annotated protein regions known to regulate mRNA stability (**Fig. 1F, starred proteins**), and expanded our analysis from looking solely at isolated tiles to searching for broader evidence of both previously-annotated or newly-discovered regulatory domains.

### Annotation of RBP regulatory domains contextualizes protein structure and functions

We defined an RNA-regulatory effector as two or more overlapping hit tiles or a single most N- or C-terminal hit tile (**Fig. 2A**). We used these rules to define 101 effectors in 86 unique proteins from a subset of the 438 tiles that were hits in the screen. Using both existing annotations and manual literature curation, we found evidence of previously-described regulatory activity for 17 of our newly defined domains. 54 domains also overlapped annotations in the Pfam/Interpro database, of which 6 were known RNA-binding domains; however, the rest of the 48 known annotations did not designate RNA regulatory activity, such as chromodomains or kinase domains in enzymatic RBPs (**Fig. 2B, Table S2, Document S1**). This leaves 78 newly annotated RNA-regulatory effectors for which we found no previous description of regulating mRNA binding, translation, or degradation (**Fig. 2B**).

**Figure 2.**
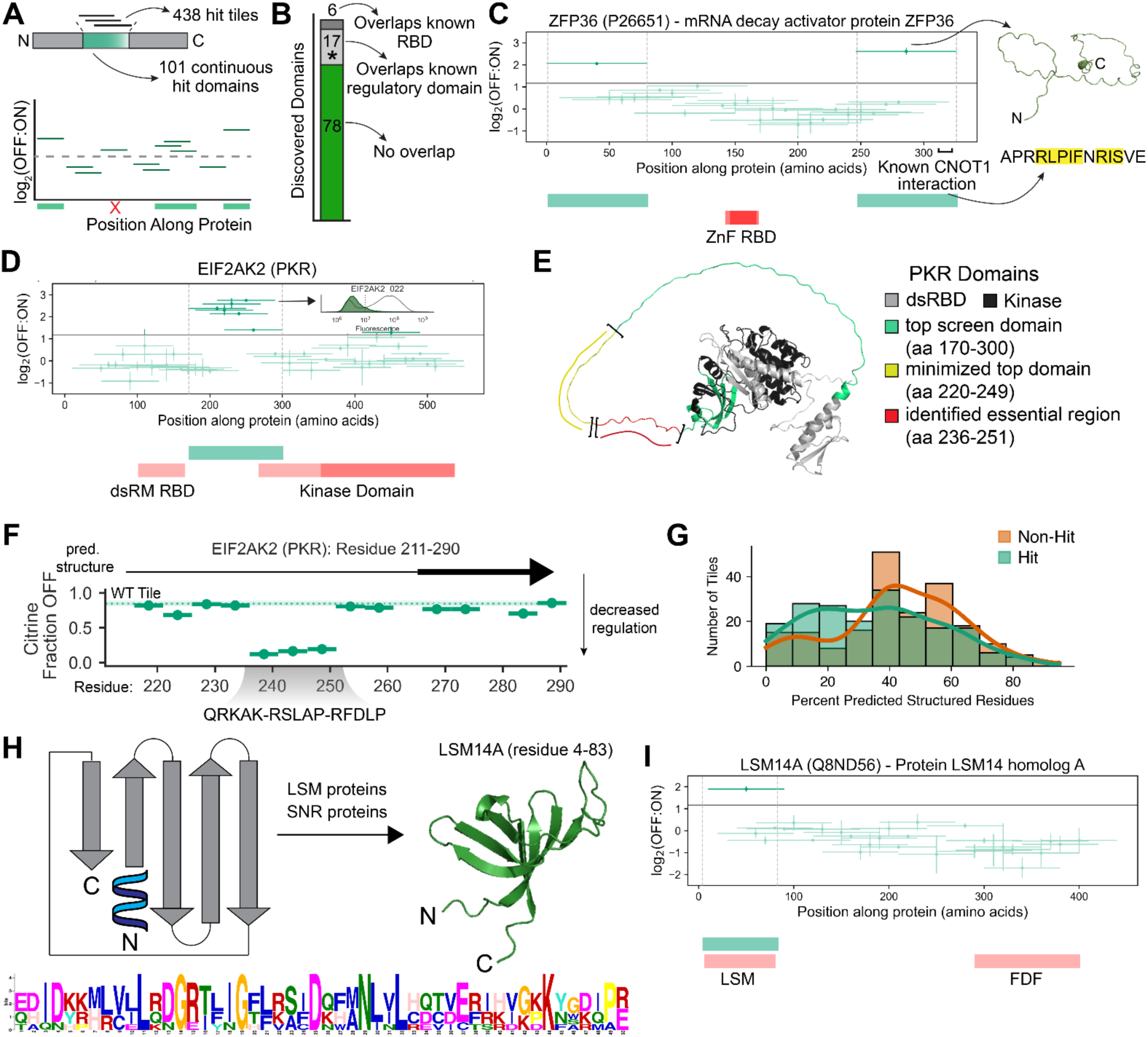
Annotation of regulatory domains in human RNA binding proteins. (A) Schematic of domain identification criteria. Single hit tiles at the N- or C-terminus or overlapping hit tiles were considered regulatory domains, but not single tiles with no overlapping neighboring hits. (B) Bar chart of 101 identified regulatory domains. 78 did not overlap either an RNA-binding domain (RBD) or previously annotated regulatory domain, 6 overlapped RBDs, and 17 overlapped known domains. (C) Tiling plot of TTP (ZFP36), known mRNA degradation activator. Vertical dashed lines indicate boundaries of newly-identified RNA-regulatory effector domain. Green boxes, newly identified RNA-regulatory effector domains; red boxes, Pfam-annotated domains. Identified C-terminal regulatory domain overlaps a known CNOT1 interaction motif (inset, right, cross-species conserved residues in yellow) and is disordered (structure, inset). (D) Tiling plot of PKR (EIF2AK2). Inset: Flow cytometry plot of individual recruitment of top tile EIF2AK2_0222 (green fill) versus MCP alone (grey line, no fill). (E) Alphafold predicted structure of PKR. Grey, annotated dsRM RBD; black, annotated kinase domain; green, our identified regulatory domain; yellow, computationally minimized sequence of regulatory domain (**Methods**); red, essential region as identified by deletion scanning mutagenesis in **F**. (F) Deletion scanning mutagenesis of EIF2AK2 tile 22 (residue 211-290). Each line is a 5aa deletion, with x-axis showing the deleted residues; dot is the fraction OFF score for that deletion as measured by flow cytometry. Shading represents the fraction OFF for the wild-type (non-deleted) tile. Vertical error bars are the average of two biological replicates. Gradient shows the essential residues whose deletion caused decreased RNA downregulation. (G) Predicted disordered character of 195 tested hit tiles (green) vs. 195 non-hit tiles (orange), as predicted by Jpred4. (H) 2-D schematic (left) and example Alphafold structure (right) of the LSm domains that were enriched in non-disordered hit tiles. Below, the MEME suite rendering of the LSm motif sequence. (I) Tiling plot of LSM14A (LSm domain structure shown in **H**).

Due to the high amount of overlap between neighboring tiles, we could more precisely define regions of activity within the set of previously-identified RNA-regulatory domains (**Fig. 1F, starred proteins**). For example, the protein CNOT4 was known to interact C-terminally with CNOT1, the catalytic member of the CCR4-NOT deadenylation complex^15^, but an exact interaction motif or region has not been reported. Our screen identifies a C-terminal regulatory effector in CNOT4 at amino acids 331-480, giving a more specific range where an interaction motif may be located (**Fig. S2A**).

Another protein with previously-annotated RNA regulatory regions is tristetraprolin (TTP, encoded by the gene ZFP36), which is also known to interact both N- and C-terminally with elements of the CCR4-NOT complex^16,17^. We identified both N- and C-terminal effectors in TTP that are each one tile long, indicating that the specific interaction motifs are likely within the first and last 10-20 amino acids of the protein and would not be present in subsequent or preceding tiles (**Fig. 2C**). Focusing on the C-terminal effector, we noticed that this most C-terminal tile indeed overlaps a conserved CNOT1 interaction motif (RLPIXRIS, aa 315-323^16^). This region is also predicted to have extremely disordered character by AlphaFold^18^, consistent with previous structural evidence showing that disordered protein regions can tightly interact with structured regions of CNOT1^10,11,16^ (**Fig. 2C**).

A newly identified RNA-regulatory effector in the protein Protein Kinase R (PKR, encoded by the gene EIF2AK2) had high screen scores for each tile and is located between two other annotated domains: a dsRM double-stranded RNA binding domain (dsRBD) and the catalytic kinase domain (**Fig. 2D**). PKR binds via its dsRBD to non-native double-stranded RNAs over 30 nucleotides in length, normally those of RNA viral genomes^19,20^. This induces dimerization of PKR kinase domains, autophosphorylation, and subsequent phosphorylation of the protein eIF2a to inhibit host translation and induce apoptosis of infected cells^21^.

However, PKR was not previously annotated to have a role in the direct degradation or regulation of RNAs that it binds. We mapped our discovered effector onto the AlphaFold structure of PKR and found that it corresponds with a disordered region directly between the dsRBD and kinase domain, suggesting that this domain is both functionally and structurally distinct and may play a role in PKR regulation of viral infection (**Fig. 2E**). We additionally used AlphaFold to predict the structure of the effector alone and confirmed that its disorder is maintained when folded outside the context of the full protein (**Fig. S2B**). To identify smaller, specific regions within this effector that are essential for its regulatory activity, we performed both computational domain minimization (the intersection of overlapping tiles) and deletion scanning mutagenesis experiments. By taking only the region of PKR that appeared in all overlapping hit tiles, we computed the ‘minimized’ regulatory domain to span from aa 220-249 in the larger disordered structural region (**Methods, Fig. 2E**). We then selected the top-scoring screen tile (aa 211-290) and created a set of mutants harboring sequential 5 aa deletions along the length of the tile. After performing recruitment experiments with each mutant tile individually, we identified 3 sequential deletion mutants at amino acids 236-251 that abrogated regulatory activity **(Fig. 2E**, **Fig. 2F)**. Mapping this region back onto the PKR structure shows that this essential region is disordered and is almost fully contained within our computationally identified minimized domain.

To determine whether most regulatory tiles and domains were similarly disordered, we used the Jpred4 secondary structure prediction server^22^ to query the expected secondary structure of 195 top hit tiles (those with screen score >=1.6) and 200 non-hit tiles to determine whether most regulatory tiles and domains were similarly disordered (**Fig. 2G**). We found a significant (p = 2.06 × 10^-5^, Kolmogorov-Smirnov statistic 0.24) disenrichment of structured amino acids in hit tiles compared to non-hits, concluding that the RNA-regulatory effectors we identified are more likely to be disordered than highly structured. Based on the rough bimodal appearance of the distribution of the percent structured amino acids for tested hit tiles, we then binned the tiles into structured (>35% structured amino acids, 94 tiles/195 tested) and unstructured (<35% structured aa, 101 tiles/195 tested) categories and submitted each to the MEME server^23^ for motif enrichment analysis. No significant motifs or significant differences in amino acid enrichment were identified in the unstructured tiles, signifying that they are unlikely to share a single interaction partner or other unifying traits (**Fig. S2C-E**).

The only significant motif identified in the structured tiles was a conserved region in the LSM protein domain, found in LSm proteins and the protein components of snRNPs involved in splicing and RNA quality control in the nucleus^24^. LSM domains have a distinct and conserved structure consisting of a short N-terminal alpha-helix followed by a five-stranded B-sheet, and appear in the library in tiles from 9 LSm proteins and 8 SNRPs^25^ (**Fig. 2H**). Of those, the LSM domains of 5 LSm proteins, including LSM14A, and 4 SNRP proteins were hits in our screen (**Fig. 2I, Fig. S2F**). Although the LSm proteins are normally found tightly associated with each other in heptameric rings that regulate splicing and RNA decapping, several individual proteins are known to interact with RNA nuclear and cytosolic degradation machinery on their own^26–29^. Our results suggest that recruitment of a single LSm component is sufficient for recruitment of either the rest of the LSm complex or of general decay factors and the subsequent induction of potent RNA downregulation.

### Regulatory domain identification and strength is dependent on recruitment position

In order to identify RNA-regulatory effectors that potentially interact with cellular mRNA processing machinery specific to one end of the mRNA, such as the 5’ decapping machinery or the 3’ polyadenylation complex^30,31^, we next changed the positioning of the MS2 stem-loops on our reporter RNA. We constructed a version of the reporter identical to the one used in our first recruitment screen, but with the 24 MS2 stem-loop cassette installed ∼100 nucleotides upstream of the translational start site of the surface marker coding region (in the 5’UTR) rather than downstream of the Citrine coding region (**Fig. 3A**). We hereafter refer to the two different reporter constructs as the 5’ and 3’ reporters. We repeated our high-throughput recruitment assay with the same ∼30,000-member RBP tiling library in cells harboring the 5’-recruitment reporter and again found our measurements to be reproducible (**Fig. 3B**, Spearman’s *r* = 0.82, **Fig. S3A**). Unlike the initial 3’ screen, we observed a large, highly populated range of tile scores (**Fig. 3B**), resulting in a clearly bimodal distribution of enrichment ratios even for random tiles designed as negative controls. The high proportion of random tiles that had activity in this screen compared to the 3’ screen shows that there is likely non-specific regulatory activity associated with recruiting proteins in high copy number to the 5’UTR of an mRNA. To confirm that the wider range of screen scores and higher proportion of random tiles with regulatory activity was not due to technical noise, we again tested 48 individual MCP-tile fusions and found the fraction of cells not expressing reporter protein to correlate very well with their corresponding 5’ screen score (**Fig. 3C**, Spearman’s *r* = 0.95, **Fig. S3B**). RT-qPCR against the Citrine reporter of selected individual validation lines also showed a direct correlation between mRNA and protein levels (**Fig. S3C**).

**Figure 3.**
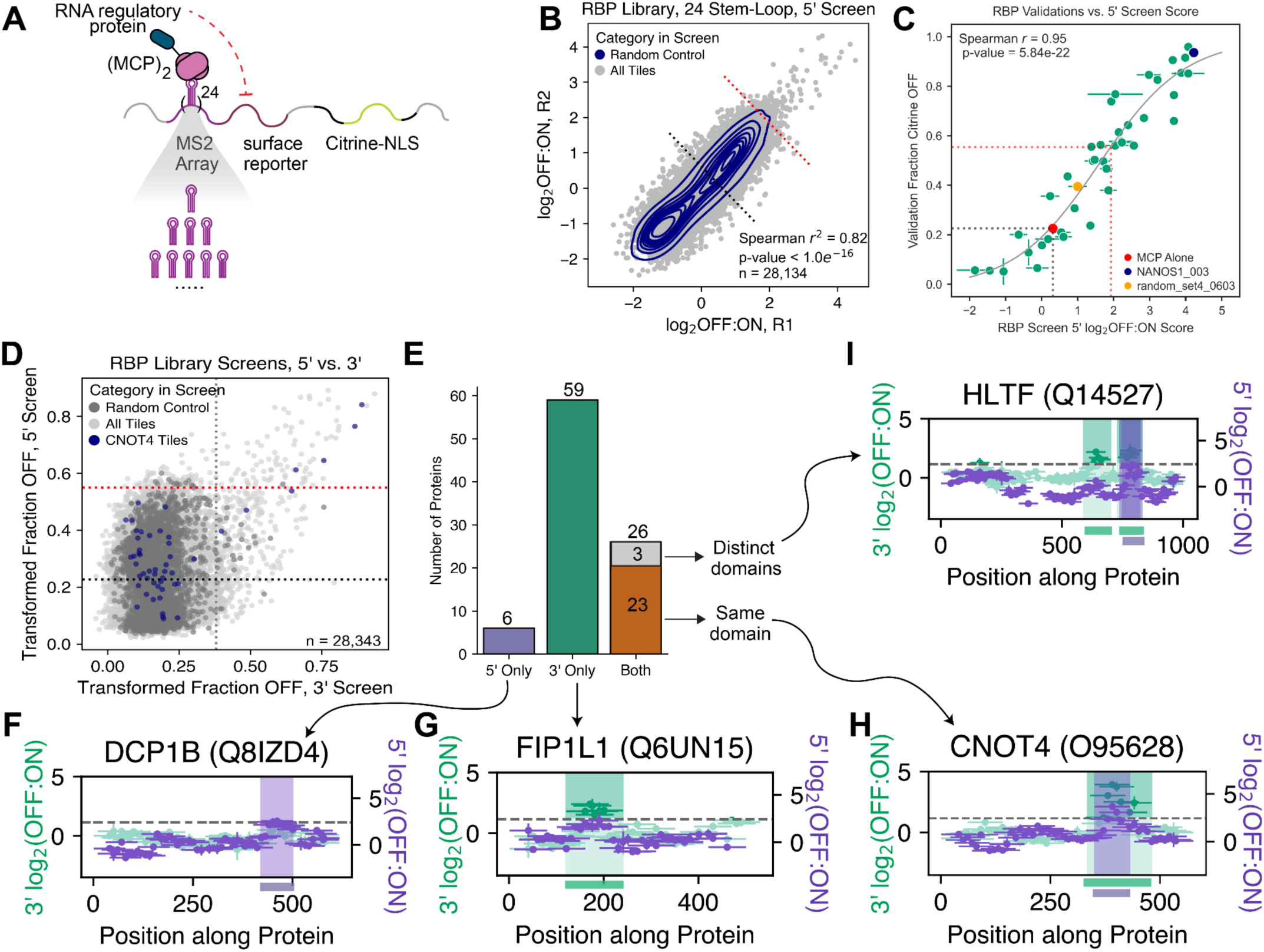
Regulatory domain identification and strength is dependent on recruitment positioning and stoichiometry. (A) Schematic of the 5’UTR recruitment RNA with site for changing numbers of MS2 stem-loops. (B) Log_2_(OFF:ON) enrichment scores plotted per replicate of the RBP library screen at 24 stem-loops in the 5’UTR. Grey dots, all tiles; purple contour lines, random control distribution. ‘High’ threshold (mean+1.5 standard deviations of the random population) in red dashed line, ‘low’ threshold in grey dashed line. (C) Individual validation measurements for 48 selected tiles in 5’UTR reporter cells. Red dashed lines represent the high screen cutoff and corresponding fraction OFF cutoff, grey dashed lines represent the same for low threshold. MCP alone is shown as a red dot, NANOS1_003 in blue, and selected random control random_set2_0603 in orange. (D) Fraction OFF scores (calculated using the fitted transformation from each set of individual validation experiments) for the full RBP library at 24 stem-loops in either the 3’UTR (x-axis) or 5’UTR (y-axis). Light grey, all tiles; dark grey, random controls; purple, tiles from CNOT4. Vertical grey dashed line, transformed cutoff for 3’ screen; horizontal red dashed line, transformed high cutoff for 5’ screen; horizontal black dashed line, transformed low cutoff for 5’ screen. (E) Summary of 91 total RBPs that were annotated with regulatory domains in the 5’UTR screen (purple), 3’UTR screen (green), or both (orange/grey). (F) Tiling plot for DCP1B showing both 24 stem-loop 3’UTR scores (green) and 24 stem-loop 5’UTR scores (purple). 5’UTR domains are shaded in purple. (G) Tiling plot for FIP1L1 with its 3’UTR domain (green shading). (H) Tiling plot for CNOT4, with its 3’UTR domain (green shading) containing its 5’UTR domain (purple). (I) Tiling plot for HLTF.

Given the high percentage of random control hits in our 5’ screen, we wondered if our high recruitment stoichiometry (24 MS2 stem-loops) in the 5’UTR of the reporter mRNA was causing non-specific regulatory effects by sterically inhibiting translation, and if this problem would be alleviated by reducing the number of MS2 stem-loops. Reducing the number of stem-loops (from 24, to 7, 3, 2, and 1) reduces the number of cells OFF for both individually chosen strong RNA-regulatory tiles and a random tile (**Fig. S3D**). However, re-running the high-throughput measurements for a smaller library (called the RBP Hit Library, 3,149 members, **Methods**) consisting of top hits from both the 3’UTR and 5’UTR screens, moderate hits from the 5’UTR screen, and some negative controls, with a 5’UTR, 7 stem-loop reporter mRNA still returns many of the random controls with scores above MCP alone (**Fig. S3E-S3J**). These results suggest that reducing the number of 5’UTR stem loops from 24 to 7 does not eliminate widespread downregulation of Citrine expression (**Fig. S3F**, population under red dotted line), leaving the mechanism of 5’UTR reporter perturbation to be determined.

We returned to the results of the 5’UTR 24 stem-loop screen to identify tiles that were clearly RNA regulators above any perturbation of the reporter that could be attributed to non-specific 5’UTR disruption. After fitting a logistic regression curve to correlate our low-throughput validations with their respective screen scores (**Fig. 3C**), we used these results to calculate two thresholds for this screen: the first is a “low” threshold at screen score = 0.30, corresponding to recruitment of the negative control (MCP alone) (**Fig. 3B, 3C**, black dotted line). The second “high” threshold was calculated by using the mean + 1.5 standard deviations of the random tiles’ scores (**Fig. 3B, 3C**, red dotted line). Although this “high” threshold excludes many tiles that have moderately high screen scores and clearly reduce Citrine levels in validation measurements (169 tiles above the ‘high’ threshold vs. 9,984 tiles between the two thresholds), using a more stringent cutoff for domain discovery ensures any annotated domains have strong regulatory activity and are not identified solely due to non-specific modulation of reporter levels.

To query whether our RNA-regulatory effectors exhibited any position-dependence and therefore putative reliance on end-specific RNA-processing machinery, we directly compared effectors identified from recruitment to the 3’ and 5’ UTRs. We used our individual validation curves for both the 3’ and 5’ 24 stem-loop reporter screens (**Fig. 1E, 3C** respectively) to calculate the fraction of cells with Citrine OFF for all tile scores. These transformed absolute fraction OFF scores can then be used to directly compare tile effects between screens, as they are not affected by differences in magnetic separation purity or sequencing depth that affect calculated enrichment ratios between high-throughput measurements taken at different times (**Methods**).

When comparing the 3’ vs. 5’ transformed OFF scores for each tile sequenced in both 24 stem-loop screens using the full RBP tile library, we noticed a strong 5’ bias with many more tiles having higher transformed OFF fractions in the 5’ screen (**Fig. 3D**), as expected from the tendency of 5’ recruitment to non-specifically reduce Citrine at moderate levels (see discussion above). Once we applied the higher 5’ hit threshold and searched for domains, we found 32 proteins with strong 5’ RNA-regulatory effectors (**Fig. 3E, Table S2**). 6 of these proteins had effectors unique to 5’ recruitment (**Fig. 3E**), such as DCP1B, a component of the mRNA decapping complex (**Fig. 3F**). 59 proteins were annotated with RNA-regulatory effectors active only in the 3’ screen, though the majority of these domains had 5’ screen scores that were above the low detection threshold but under the high 5’ threshold (**Fig. 3E, 3G**). One example of such a protein is FIP1L1, which is a known subunit of the CPSF (cleave and polyadenylation specificity factor) that regulates 3’ processing and mRNA stability^32^ (**Fig. 3G**). 23 proteins had one or two strong effectors active when recruited at either the 5’ or 3’ UTR (**Fig. 3E, 3H**). One protein (ADAD2, **Fig. S3K**) had one effector active only in the 5’ recruitment screen and one that was unique to the 3’ screen, and the remaining 2 proteins had multiple 3’ effectors and a single effector active in both screens: HLTF (**Fig. 3I**) and SETD1B^33^ (**Fig. S3L**). In summary, this resulted in a total of 91 unique proteins annotated with 108 unique RNA-regulatory effectors overall (**Table S2**).

### Discovery of domains that can downregulate gene expression when recruited to either RNA or DNA

Given that some members of our RBP tile library come from proteins known to affect RNA processing in the nucleus, we asked if any tiles in our library act as direct transcriptional regulators. Since the MS2 mRNA reporter is transcribed from the genomic integration site, some of the effects we see with MS2-MCP-mediated recruitment could be a result of co-transcriptional interactions of the MCP-tile fusion with chromatin regulators or transcriptional machinery, thus affecting transcription rather than post-transcriptional regulation of RNA export, degradation or translation. In order to test tiles for direct effects on transcription, we recruited tiles directly to DNA upstream of the Citrine reporter, by fusing the RBP Hit Library (3,149 members, consisting of all top hits from the 3’, 24 stem-loop screen; some hits from the 5’, 24 stem-loop screen; and a group of negative controls, as in **Fig. S3F**) to rTetR, a doxycycline-inducible DNA-binding domain used in previous transcriptional effector screens^5^. We delivered this new fusion library to K562 cells expressing a reporter that encodes the same surface reporter and Citrine, but lacks the MS2 stem loops and instead contains a 9xTetO binding site array for rTetR upstream of the strong pEF promoter that drives reporter expression (**Fig. 4A**).

**Figure 4.**
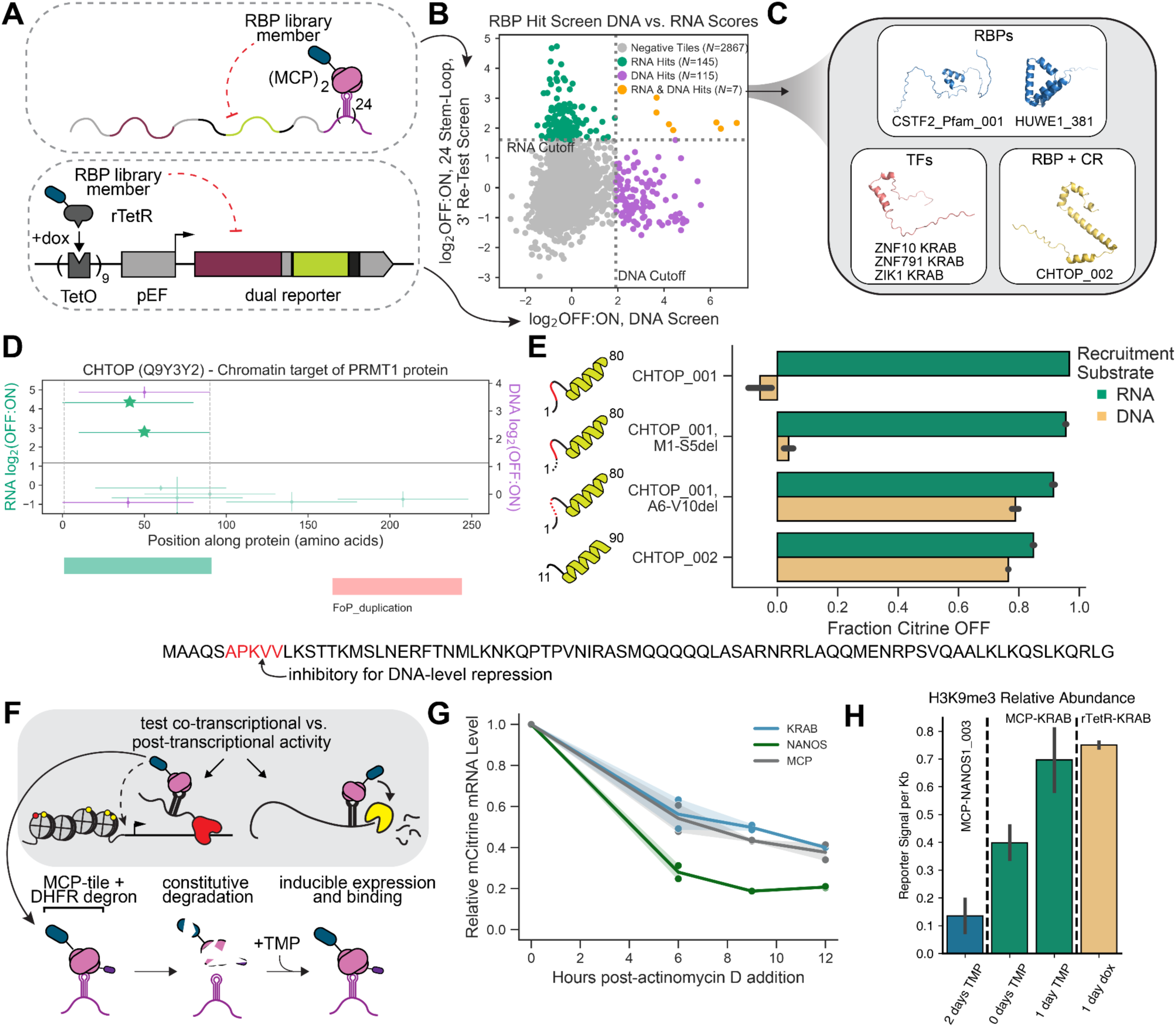
DNA-level recruitment investigates dual DNA- and RNA-mediated control by RBP regulatory domains. (A) Top, schematic of MS2 reporter construct. Bottom, schematic of TetO reporter construct. RBP tiles are cloned to the dox-inducible DNA-binding domain rTetR and recruited to a reporter expressing the same surface reporter and Citrine in the MS2 reporters. (B) Average log_2_(OFF:ON) enrichment scores (two replicates) for the RBP Hit library in the TetO DNA screen (x-axis) and the batch-retest 3’UTR 24 stem-loop RNA screen (y-axis). Purple, DNA hits; green, RNA hits; yellow, dual hits; grey, rest of tiles; vertical dashed line, DNA screen cutoff; horizontal dashed line, RNA screen cutoff. (C) Alphafold predicted structures of the categories of non-random-control dual hits. Blue, tiles from known RNA regulators; yellow, known RBP + chromatin regulator CHTOP; pink, KRAB domain. (D) Tiling plot for CHTOP for the original RBP Library 3’UTR screen (green) and the DNA screen (purple). Stars represent tiles re-tested in the smaller batch screen at the 3’UTR, 24 stem-loop reporter. (E) Summary of individual flow cytometry measurements of CHTOP deletions, with the identified putative DNA-inhibitory region shown in red on the schematics at left and each deletion shown as a dashed line. Green bars, fraction Citrine OFF when measured on the MCP-MS2 RNA recruitment system at 24 stem-loops in the 3’UTR; yellow bars, fraction Citrine OFF when measured on the rTetR-TetO DNA system. Bottom, sequence of CHTOP regulatory region with inhibitory 5 amino acids in red. Error bars are standard deviations of two biological replicates. (F) Top, schematic of possible mechanisms of transcriptional vs. post-transcriptional regulation during RNA recruitment. Bottom, schematic of DHFR-MCP degradation and TMP-induced stabilization. (G) Citrine mRNA levels over time as measured by flow cytometry of HCR-Flow-RNA-FISH, normalized to timepoint 0 for each experiment for cells expressing the 5’UTR, 7 stem-loop reporter. Timecourses are taken after the concurrent addition of actinomycin D and TMP (at time=0) for DHFR-MCP alone (grey), DHFR-MCP-ZNF10_KRAB (blue), and DHFR-MCP-NANOS1_003 (green). Shading is standard deviation of two biological replicates. (H) Relative H3K9me3 levels as measured by CUT&RUN, integrated over the 5kb locus of the 5’UTR, 7 stem-loop reporter, for TMP and dox recruitment of DHFR-MCP-NANOS1_003 (blue), DHFR-MCP-ZNF10_KRAB (green), and rTetR-ZNF10_KRAB (yellow). Error bars are standard deviations of two biological replicates.

After selection, we added doxycycline at 1,000 ng/mL to induce binding of rTetR and recruitment of the fused RBP Hit Library tiles to the reporter locus. We performed magnetic separation identically to the MCP-fusion screens after 7 days of doxycycline treatment (**Fig. S4A**). At the same time, we also re-screened this smaller library fused to MCP and recruited at the 3’ UTR of the 24xMS2 stem loop reporter as an additional re-test of results from our large initial screen (**Fig. S4B**).

After performing both screens, which showed good reproducibility (**Fig. S4C-S4F**), we compared each tile’s average screen score when recruited to DNA vs. RNA and found the hits to be almost completely mutually exclusive (**Fig. 4B**): 145 tiles were hits only in the RNA-recruitment screen and 115 only in the DNA-recruitment screen. Of the 103 proteins with tiles in the library, only 14 proteins had at least one tile called as a hit in both screens (though not necessarily the same tile), while 47 proteins had one or more RNA hit tiles only and 42 proteins had only DNA hit tiles (**Fig. S4G**). However, only 7 individual tiles showed strong regulatory activity both when recruited to 3’ RNA (via MCP) and to DNA (via rTetR): three KRAB domains from zinc-finger transcription factors, one random control tile, and one tile each from ubiquitin E3 ligase HUWE1, the 3’ RNA processing factor CSTF2, and CHTOP, a protein known to bind both chromatin regulator PRMT1 and act as a component of the nuclear export complex TREX^34^ (**Fig. 4C**).

Since CHTOP was previously annotated to interact with both chromatin and RNA regulatory pathways, we further investigated the tile that we found to have dual repressive activity: CHTOP_002, spanning amino acids 11-90 (**Fig. 4D**). Another tile, CHTOP_001, from amino acids 1-80, is a consistently strong RNA regulator when recruited to both the 3’ and 5’ of RNA but lacks any activity when recruited to DNA (**Fig. 4D**). Both of the following two tiles - CHTOP_003 or CHTOP_004 - were inert when recruited to either RNA or DNA (**Fig. S4H**). We hypothesized that either: (1) the last 10 amino acids of CHTOP_002 (81-90) were conferring the transcriptional repressive activity unique to that tile, or (2) the first 10 amino acids of CHTOP_001 (1-10) were inhibiting its ability to repress when recruited to DNA. To this end, we created systematic deletions of these two tiles (**Fig. 4E**). Deleting the last 10 amino acids of CHTOP_002 had no effect on its activity (**Fig. S4H**), allowing us to reject the first hypothesis above. Deleting amino acids 6-10 from CHTOP_001 led to a gain in DNA regulatory activity comparable to wild-type CHTOP_002, while maintaining its strong RNA regulating capability (**Fig. 4E, S4I**), while deleting the amino acids 1-5 had no effect on either its RNA- or DNA-level activity (**Fig. S4H**). This led us to conclude that the second five amino acids of CHTOP (APKVV) specifically inhibited transcriptional repressive activity (**Fig. 4E - schematic under graphs**). It remains to be determined how this sequence regulates CHTOP activity at its endogenous targets.

We next turned to investigate the biggest class of dual RNA/DNA-regulating tiles: the KRAB domains, which belong to the largest class of transcriptional repressor domains most commonly found in zinc finger transcription factors^35^. We selected the KRAB domain from ZNF10, which is commonly used as an epigenetic silencing tool^36^, and asked if its effects in the RNA recruitment assay were due to increasing rates of RNA degradation or through *in trans* interactions with the reporter locus while bound to the reporter mRNA (**Fig. 4F, top**). To do so, we developed a system for inducible recruitment with MCP-MS2 to measure the dynamics of RNA regulation over time. We fused the (trimethoprim) TMP-dependent DHFR degron to the N-terminus of MCP and a HaloTag to its C-terminus, then cloned ZNF10 KRAB fused C-terminally to the HaloTag (**Fig. 4F, bottom**). We confirmed that there was very little protein before TMP addition and that MCP-tile levels robustly stabilized as early as 6 hours (as measured by HaloTag stain) after addition of 10 μM TMP (**Fig. S4J**). We then used this system to test the timescale of RNA degradation separate from putative transcriptional silencing. We added actinomycin D (actD; to inhibit RNA polymerase activity and stop transcription) and TMP (to stabilize DHFR-MCP fusions) and measured the RNA half-life with HCR-Flow-RNA-FISH (**Fig. 4G**). The reporter mRNA half-life when bound only by DHFR-MCP alone was 7.63 +/- 0.002 hours, consistent with measurements of a very similar reporter molecule^33^ (**Fig. 4G**). The half-life decreased to 3.83 hours when bound by DHFR-MCP-NANOS1_003 (a known interactor with RNA degradation machinery) but showed virtually no change when bound by DHFR-MCP-KRAB (8.36 hours) (**Fig. 4G**), suggesting that MCP-KRAB recruitment does not increase RNA degradation.

Finally, to test the chromatin-mediated transcriptional effect of KRAB when recruited to the locus via RNA, we recruited DHFR-MCP-KRAB and measured the abundance of histone H3, lysine 9 trimethylation (H3K9me3), a repressive histone mark, at the reporter locus (**Fig. 4H**). KRAB domains are known to associate tightly with protein KAP1, which nucleates the assembly of a transcriptional repressive complex including the histone H3 methylase SETDB1^37^. We hypothesized that the presence of H3K9me3 at the reporter locus upon RNA-mediated recruitment would indicate recruitment of KAP1 and SETDB1 by KRAB *in trans.* KRAB-containing proteins are known to bind DNA directly and recruit the methylase complex, but have not been shown to act by binding to RNA and recruiting H3K9me3 to the locus co-transcriptionally. MCP-KRAB led to increased H3K9me3 levels after one day of RNA-mediated recruitment, similarly to rTetR-KRAB (DNA-mediated recruitment), and significantly higher than the RNA-mediated recruitment of MCP-NANOS1_003 which is known to act via RNA degradation (**Fig. 4H, Fig. S4K-L**). Together, the increase in repressive histone modifications and unchanged RNA lifetime indicate that the repressive effect of KRAB is through transcriptional regulation at the DNA locus both when recruited to DNA and RNA.

### Synthetic RNA-level control of gene expression expands working models for gene regulation

A potential application of our newly discovered RNA-regulatory effector domains is to tune RNA levels in gene regulatory circuits. To utilize this type of regulation in synthetic biology, we need to develop a predictive mathematical model of RNA-mediated regulation and compare it to the classical synthetic transcriptional regulation, such as the one mediated by KRAB. To test how different levels of the RNA-regulatory effector change mRNA expression dynamics, we adapted our recruitment system by designing a cell line stably expressing DHFR-MCP-HaloTag-NANOS1_003 (as in **Fig. 4G-H**, a strong regulator in both the 3’ and 5’ screens), which we named synNANOS (**Fig. 5A**). We first showed that synNANOS can efficiently downregulate reporter expression in the presence of saturating TMP (**Methods**, **Fig. S5A**). By increasing the TMP concentration, we can gradually increase the levels of synNANOS as measured by HaloTag staining (**Fig. 5B**). These data allow us to mathematically fit the dependence of synNANOS as a function of TMP concentration using a Michaelis-Menten equation (**Fig. 5B**, blue line). The TMP-dependent increase of stabilized synNANOS leads to a gradual decrease in the Citrine reporter mean fluorescence intensity (**Fig. 5C**): log-fold changes in TMP concentrations lead to linear decreases in measured Citrine expression for a wide range of TMP.

**Figure 5.**
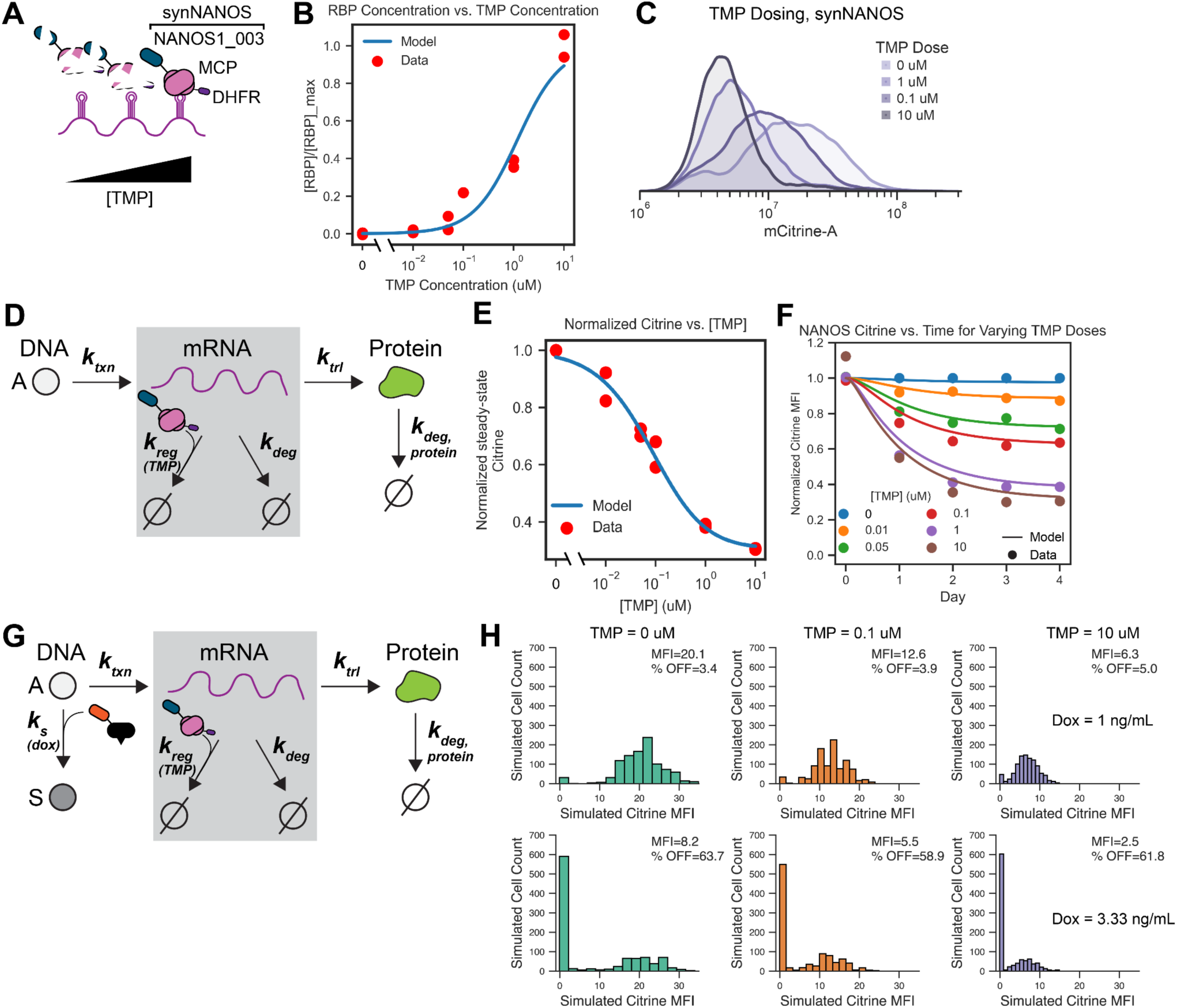
A tunable RBP creates synthetic RNA-level gene regulation and expands gene expression models. (A) Schematic of dose-tunable DHFR-MCP construct, which is expressed more stably with increasing TMP. (B) Model fit for relative synNANOS levels, as measured using HaloTag staining, at different TMP doses to determine ***K_D,TMP_***, or relative TMP dependence on RBP expression levels. Model RMSE=0.08, Data SD=0.35. (C) Flow cytometry distributions of synNANOS recruited to the 7 stem-loop 5’UTR reporter at varying concentrations of TMP. (D) Schematic for a mathematical model of gene regulation at the RNA level, where an active gene (A) produces mRNA at a constant rate ***k_txn_***. mRNA is either degraded by a TMP-dependent RBP at a rate ***k_reg_***, or by constant cellular mRNA degradation at a rate ***k_deg_***. Protein translation and protein degradation occur at constant rates ***k_trl_*** and ***k_deg, protein_***, respectively. (E) Model fit to Citrine levels after recruitment of synNANOS at varying TMP doses after 4 days of recruitment. Model RMSE=0.03, Data SD=0.28. (F) Model fit to Citrine levels after recruitment of synNANOS at varying TMP doses over time. Model RMSE=0.06, Data SD=0.25. (G) Overview of Gillespie simulation assumptions: the gene can either be in the active (A) or silenced (S) state, the transition between which is controlled by a transcriptional silencer at a dox-dependent rate ***k_s_***. Cells can produce mRNA in either the A or S states at a constant rate ***k_txn_***; mRNA is then degraded by the TMP-controlled RBP at a rate ***k_reg_*** or the constitutive rate ***k_deg_.*** Protein is made and degraded at the constant rates ***k_trl_*** and ***k_deg,protein_***, respectively. (H) Gillespie simulation results for cells hypothetically expressing a dox-inducible transcriptional silencer and a TMP-inducible RNA regulator, for two dox doses (top row, 1 ng/mL; bottom row, 3.33 ng/mL) and three TMP doses (columns, L-R: 0, 0.1, and 10 μM). Y-axis, number of simulated cells; X-axis, simulated Citrine fluorescence intensity.

In contrast, when we perform transcriptional control by recruiting the ZNF10 KRAB domain fused to rTetR at different concentrations of doxycycline at the 9xTetO response element upstream of the promoter (**Fig. S5B**), the response to the inducer is abrupt, as reported before^38^: rather than gradual shifts in MFI, we observe a sharp change in total MFI between 2-10ng/ml of dox (**Fig. S5C-D**). This response likely comes from the previously observed and modeled single-cell stochasticity associated with chromatin-mediated gene silencing^39,40^, and emphasizes the need for development of a new mathematical model incorporating RNA regulation that describes gradual MFI decreases mediated by RBPs (**Fig. 5D**).

To model RNA control, we considered a gene in the actively transcribing state where cells produce mRNA at the rate of transcription ***k_trx_***. This mRNA can be basally degraded by the cell at rate ***k_deg_***, or it can be degraded faster when bound by the RNA regulatory domain at a rate ***k_reg_*** (**Fig. 5D**), which is characteristic of a specific domain. We assume any protein translated from existing mRNA is translated at a rate ***k_trl_*** and is subsequently degraded at a rate ***k_deg, protein_***. From this model, we can derive an expression for the steady state reporter levels as a function of TMP concentration and further reduce the number of free parameters by experimentally measuring the natural mRNA degradation rate and how synNANOS concentration scales with TMP concentration. Using HCR-Flow-RNA-FISH to measure RNA degradation kinetics after inhibiting transcription with actinomycin D (**Fig. 4G**), we approximate the reporter RNA degradation rate in the absence of synNANOS to be ***k_deg_*** = 0.1/hr. The presence of synNANOS decreases the steady state reporter mRNA levels (mRNA_ss_) compared to maximum levels (in its absence) in a manner that depends on its associated degradation rate (***k_reg_***, eq. 1, **Methods** for derivation).

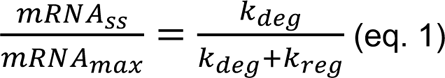

The synNANOS-associated rate of degradation (***k_reg_***) depends on the concentration of synNANOS, which changes with TMP and is described by eq. 2 (see **Methods** for derivation):

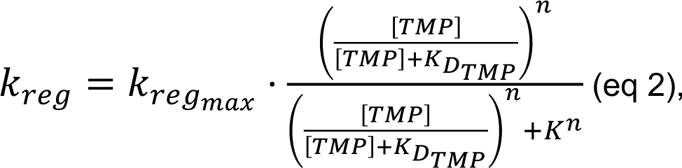

where ***K_D(TMP)_*** is the Michaelis-Menten constant describing how synNANOS concentration depends on TMP concentration, ***k_reg, max_*** is the maximum synNANOS-associated degradation rate occurring at saturating synNANOS levels, *K* is the equilibrium binding of synNANOS to RNA, and *n* is the Hill coefficient.

We first extract ***K_D(TMP)_*** = 1.19 by directly fitting the synNANOS protein levels as a function of TMP (**Fig. 5B**). Then, using ***k_deg_*** and ***K_D(TMP)_*** as input to equations 1&2, we can fit the steady-state Citrine levels (normalized to maximum) as a function of TMP concentration (**Fig. 5E**, blue line), with free parameters *K* and *n* describing the binding of synNANOS to RNA. ***k_reg,max_*** describes the rate of synNANOS-associated mRNA degradation at saturating synNANOS levels, with higher values of ***k_reg, max_*** corresponding to higher sensitivity to TMP and more mRNA degradation at lower concentrations of RBP (**Fig. S5E**), and can be tuned by changing the regulatory domain in a synthetic RBP to create a more or less potent RNA degrader.

Finally, we can use the parameters fit to steady-state mRNA levels at varying TMP concentrations to predict reporter mRNA degradation kinetics over time (**Fig. 5F**). We find that the model predicts steady-state Citrine levels to be reached by 2-3 days of TMP treatment, and that varying TMP doses allows steady-state relative Citrine levels from 1 to as low as 0.4. We compared our RNA degradation model to a model of KRAB-mediated transcriptional silencing, where varying dox concentration directly modulates ***k_s_*** (the probability of a cell to be in a completely silent transcriptional state) as previously described^38^ (**Fig. S5F**) . By fitting this model to our rTetR-KRAB recruitment data, we could describe KRAB silencing over time at varying dox doses (**Fig. S5G**). Unlike in the RNA regulation model, transcriptionally-regulated Citrine levels are not predicted to reach intermediate steady state by 5 days of recruitment, and higher doses of dox trend sharply toward a normalized fluorescence of 0 as recruitment continues (**Fig. S5G**). The transcriptional silencing and RNA degradation models accurately capture the different dynamics and magnitudes of KRAB and synNANOS recruitment effects on Citrine expression and allow us to extract parameters that are most relevant to the distinct mechanisms of gene silencing employed by transcription factors vs. RBPs.

## Discussion

Over 1,000 RBPs exist in human cells and coordinate each step of the mRNA lifecycle, making them the largest class of regulators of post-transcriptional gene expression^41^. Despite previous high-throughput work annotating protein regions with mRNA regulatory potential across the entire yeast proteome and large-scale studies of full-length human RBPs, we lack a comprehensive “parts list” of the RBP domains that control mRNA regulation in human biology. By engineering a high-throughput RNA recruitment assay in human cells, we tested over 30,000 protein tiles from 367 RBPs for mRNA regulatory capacity. We identify over 100 regulatory domains from 86 distinct RBPs, creating the first resource of compact RNA-regulatory effectors from human proteins. We find RNA-regulatory effectors in RBPs with a wide range of RNA-dependent processes, such as deadenylation or quality-control RNA degradation, as well as in RBPs with no previously known regulatory function. Although there are some Pfam-annotated domains that partially overlap our newly annotated domains, they do not denote regulatory activity and are largely distinct from our RNA-regulatory effectors. We also performed Alphafold structural prediction of each of our discovered regulatory domains and found that almost all of them are predicted to have the same structure as their corresponding region in the full-length RBP (as determined by Alphafold or PDB structures, **Table S2**). These analyses combined indicate that in many RBPs, RNA-regulatory effectors are both functionally and structurally distinct from other functional domains such as RNA-binding domains. This is the first time RNA-regulatory effectors have been reported at scale for human RBPs, and these findings are concordant with those from previous studies performed with individual human proteins^13^ and high-throughput yeast studies^9,11^.

Since our recruitment assays utilize relatively short 80 amino acid tiles, the majority of our RNA-regulatory effectors are likely mediating protein-protein interactions with full-length RNA-regulatory complexes that contain larger proteins with enzymatic function, including mRNA degradation or translation. Our analyses find that RNA-regulatory effectors tend to be disordered, which has also been shown in yeast and in isolated examples of human proteins^42^, and supports a hypothesis that RBPs are likely to recruit their cognate cellular complexes through general hydrophobic interactions driven by their disordered character and variable sequence motifs^43^. However, we found no significant enrichment of amino acid composition or widespread presence of conserved short linear motifs (SLiMs) in the sequences of our annotated RNA-regulatory effectors, despite the fact that many full-length RBPs have been shown to work through convergent pathways and interact with the same cellular machinery to exert their effects^10,44,45^. This is possibly a limitation coming from the number of effectors tested in this study rather than a lack of consensus motifs that bind RNA regulatory machinery: if there are tens of divergent motifs, our 101 RNA-regulatory effectors would not be a sufficient sample size to pinpoint the motif rules. For example, the few SLiMs that have been identified in binding partners of the major deadenylation complex CCR4-NOT are not conserved across RBPs; different RBPs known to recruit the same CCR4-NOT components do so with different protein sequences or at different regions of the interacting protein. TTP and NANOS1 both interact with CNOT1 through divergent sequences that share almost no sequence characteristics - a C-terminal motif in TTP (that we find in its C-terminal regulatory domain), and a so-called NOT1-interacting motif (NIM) in the regulatory domain we identify in NANOS1^46,47^. In the future, we can use our high-throughput assay to dissect these sequence rules by performing deep mutational scanning to identify more specific smaller essential regions and putative motifs within our domains.

The vast majority of mRNA tethering assays insert the RNA stem-loops, or site of recruitment, into the 3’UTR of the mRNA reporter substrate^48^. We demonstrate here that 3’UTR recruitment indeed reports more specifically on the regulatory strength of a tethered protein than 5’UTR recruitment, which is sensitive to perturbation even when bound by some inert controls. We hypothesize that stem-loop placement upstream of the ribosome-binding site and start codon interferes with ribosome scanning and translation initiation through steric hindrance by very stable RNA structures^49–51^. Decreasing the number of stem-loops from 24 to 7 does not resolve these non-specific effects, likely because it still results in a much more highly structured 5’UTR than would be expected in endogenous RNAs. This position-dependence of recruitment could be studied more systematically beyond 3’and 5’ UTRs. For example, our assay could be extended by inserting stem-loops in distal introns or non-coding exons, where RBPs have been shown to have higher natural residence than in UTRs most commonly used in synthetic assays^2,52^.

While here we focused on RNA-binding proteins, recent studies have proposed that human transcription factors could also possess RNA binding and regulatory activity^53,54^. In addition, a significant number of proteins have been designated as both DNA- and RNA-binding, and most have been posited to have different functions depending on which substrate they are bound to^55^. At the level of effector domains, we find that the vast majority of RNA-regulatory effectors do not have an effect on transcription when recruited directly to DNA upstream of the promoter, which aligns with previous work suggesting that most gene regulatory proteins have effects specific to the RNA or DNA sequence they most natively bind^56,57^. However, we do find a handful of domains that work in both contexts. We show that multiple KRAB domains, including the one from ZNF10, act as transcriptional repressors whether bound to DNA or tethered to RNA and recruit the same repressive histone machinery from both substrates. It remains to be determined if this is the case for the entire KRAB family or other strong transcriptional repressors, but is another example of regulatory proteins retaining their native function regardless of the substrate they bind. It is possible that some proteins retain bona fide dual-DNA/RNA-regulatory function, such as CHTOP, known to affect both TREX-mediated RNA export and PRMT-mediated histone arginine methylation^58,59^ - but most strong RNA-regulatory effectors appear to have only one main function.

Recent advances in programmable RNA targeting and editing^60,61^ have drastically increased the possibilities for synthetic RNA manipulation and engineering. We describe the discovery of hundreds of compact RNA-regulating protein tiles that can be fused to RNA-targeting proteins for selective mRNA downregulation at varying levels, as we report tiles with wide ranges of regulatory strengths. By testing thousands of tiles at different UTR positioning and in lower stoichiometry, we also produce a catalog of tiles that are more likely to be effective when endogenously recruited at low copy number. Addition of a DHFR degron to our tested domains makes them drug-inducible, an added advantage for building synthetic circuits at the RNA level or testing RNA-targeting protein therapeutics. Finally, by observing that incorporation of RNA-mediated control into existing models for gene expression creates accurate predictions of the dynamics of regulatory domain-mediated RNA degradation: we show that protein output can be finely tuned at the RNA level, thus facilitating incorporation of RNA regulation into more extensive gene regulatory networks for multi-input synthetic systems. We modeled such a multi-input system by writing a Gillespie simulation for cells that express both rTetR-KRAB (for transcriptional control) and synNANOS (for RNA control), and simulated gene expression profiles for cells treated with different doses of their respective inducers (dox and TMP). In this system, dox tunes the fraction of cells stably silenced while TMP tunes the average expression level of the non-silenced cells, allowing the system to reach different bistable populations that would not be achievable with transcriptional or RNA-mediated control alone (**Fig. 5G-H, S5H**).

Overall, our work to build a high-throughput RNA recruitment assay and test over 30,000 protein fragments for RNA regulatory activity expands knowledge of both RBP organization into regulatory domains and of existing protein sequences with specific RNA control capabilities. Our results and the tools we developed lay the foundation for future work to investigate more deeply the sequence, positioning and stoichiometry rules that govern RBP regulation in the large scale that is required for such assays.

## Supporting information

Table S3. Individual validation flow cytometry, qPCR, and Flow-FISH data.

Table S1. RBP Library sequences, enrichment scores, literature review

## Author Contributions

A.R.T. and L.B. conceptualized and designed the study. A.R.T. designed and generated all MS2 reporters and reporter cell lines. A.R.T. designed the libraries. A.R.T. and X.S.C. cloned and screened the libraries. A.R.T. analyzed screen data. A.R.T. generated plasmids and individual cell lines and performed individual recruitment assay experiments with assistance from C.B. A.R.T. performed qPCR of individual cell lines. Y.F., X.S.C., and A.R.T. performed HCR-RNA-Flow-FISH experiments. Y.F. and A.R.T. performed transcriptional inhibition experiments. A.R.T. and Y.F. performed HaloTag staining experiments. A.R.T. performed and analyzed CUT&RUN experiments. C.A. wrote the steady-state and kinetic RNA degradation models, the KRAB silencing model, and the Gillespie simulation, with input from A.R.T. and L.B. A.R.T. and L.B. wrote the manuscript with contributions from all authors. L.B. supervised the study.

## Acknowledgements

We thank Sage Allen for experimental assistance and extraordinary lab managerial skills; Ernst Pulido for assistance with preliminary recruitment experiments; Taihei Fujimori for data processing and visualization assistance; Joydeb Sinha and Nicole DelRosso for experimental advice; and Prof. Lauren Hagler, Prof. Dan Herschlag, Prof. Mike Bassik, Julia Schaepe, Benjamin Doughty, and all members of the Bintu lab for helpful conversations. This work was supported by NIH-NCI F30CA287739-01 (A.R.T.), a Stanford Bio-X Bowes Fellowship (A.R.T.), a Stanford Sarafan Chem-H Chemistry-Biology Interface Fellowship (A.R.T.), NIH T32GM145402 (A.R.T.), NIH-NHGRI R01HG011866 (L.B.), NIH-NIGMS MIRA R35GM12894701 (L.B.), and NIH R01 GM132899 (D.H.).

## Supplemental Information

**Table S1.** RBP Library sequences and enrichment scores from the 3’ 24 stem-loop screen and 5’ 24 stem-loop screen (tab 1); literature review and annotations of top hit proteins from 3’ 24 stem-loop screen (tab 2); RBP HIt Library sequences and enrichment scores from the 3’ 24 stem-loop batch re-test screen, 5’ 7 stem-loop screen, and DNA screen (tab 3).

**Table S2.** All discovered regulatory domains from the RBP Library 3’ (tab 1) and 5’ (tab 2) screens, including overlapping with Pfam database annotations and Alphafold structural predictions.

**Table S3.** Individual validation flow cytometry, qPCR, and Flow-FISH data.

**Document S1.** Tiling plots for 3’ 24 stem-loop RBP Library screen, related to Figs. 1 and 2. Tiling plots for 131 hit RBPs. The x-axis shows the position of tile along the protein with each horizontal line representing an 80 aa tile, while the y-axis shows the screen enrichment score with vertical error bars showing the range from two biological replicate screens. The solid line represents the screen hit cutoff and dashed vertical lines represent the edges of the determined regulatory domain span. Green boxes represent discovered regulatory domains, while red boxes represent known domains from the Pfam/Interpro database.

## Methods

### Resource Availability

#### Lead Contact

Further information and requests for resources and reagents should be directed to and will be fulfilled by the Lead Contact, Lacramioara Bintu.

#### Materials Availability

Information for previously published plasmids is available in the Methods section.

#### Cell culture

Cell culture was performed as described in^6^. Briefly, all experiments were carried out in K562 cells (ATCC, CCL-243, female), which were cultured in a controlled humidified incubator at 37°C and 5% CO2 in RPMI 1640 (Gibco, 11-875-119) media supplemented with 10% FBS (Omega Scientific, 20014T) and 1% Penicillin-Streptomycin-Glutamine (Gibco, 10378016). All MS2 reporter cell lines were generated as in^5^. Reporter DNA was integrated by TALEN-mediated homology-directed repair to integrate donor constructs into the *AAVS1* locus by electroporation of 1 × 10^6^ cells with 1 μg of reporter donor plasmid and 0.5 μg of each TALEN-L (Addgene no. 35431) and TALEN-R (Addgene no. 35432) plasmid using program T-016 on the Nucleofector 2b (Lonza, AAB-1001). After 48 hours of recovery, cells were treated with 500 ng/mL puromycin antibiotic (Invivogen #ant-pr-1) for 7 days to select for a stably integrated population. Fluorescent reporter integration and expression was measured by flow cytometry. HEK293T-LentiX (Takara Bio, 632180, female) cells were used to produce lentivirus (as described below) and were grown in DMEM (Gibco, 10569069) media supplemented with 10% FBS (Omega Scientific, 20014T) and 1% Penicillin-Streptomycin-Glutamine (Gibco, 10378016). These cell lines were not authenticated. All cell lines tested negative for mycoplasma.

#### Lentiviral production and transduction

Small-scale lentiviral production was performed as described in^62^. Briefly, HEK293T Lenti-X cells were seeded at 5 × 10^5^ cells per well in 2 mL of DMEM in 6-well plates. After 24 hours, cells were transfected with 750 ng of an equimolar mixture of three third-generation production plasmids (pMD2.G: Addgene #12259; pRSV-Rev: Addgene #12253; pMDLg/pRRE: Addgene #12251; all gifts from D. Trono) and 750 ng of plasmid encoding the gene of interest. The 4 plasmids were incubated for 15 minutes with 5 μL of polyethylenimine (PEI, Polysciences #23966) before transfection. After 72 hours of incubation, lentivirus was harvested and collected and supernatant was filtered through 0.45 μM PES filters (CELLTREAT #229749). Undiluted, filtered virus was added to K562 cells at a final concentration of 1-2 × 10^5^ cells/mL and centrifuged in 1.5 mL Eppendorf tubes at 1,000 × g for 2 hours, after which supernatant was discarded and cells were cultured for two days in fresh media. After 48 hours of culture, antibiotic selection was initiated with blasticidin (10 μg/mL, Gibco #A1113903) and infection and selection efficiency were monitored daily with flow cytometry on a Bio-Rad ZE5 Cell Analyzer (Bio-Rad #12004278).

Lentiviral production for screens was performed by seeding 9 × 10^6^ HEK293T Lenti-X cells into 15-cm dishes in 30 mL of DMEM. The next day, cells were transfected as above using 11.25 μg packaging plasmid mixture, 11.25 μg cloned library plasmids, and 150 μL PEI. 24 hours after transfection, a full media changed was performed; 72 hours after transfection, supernatant was harvested and filtered with a 0.45 μm PES filter unit (Thermo Scientific #1680045).

#### Human RBP tiling library design

The gene symbols of 1100 human RBPs were passed into the Python package Mygene to extract their Uniprot IDs and associated GO annotations. Uniprot was then accessed through its Python API to pair each gene symbol and Uniprot code with their corresponding amino acid sequence. The list of 1100 proteins was filtered to exclude the GO terms “’rRNA’, ’No GO term found’, ’splic’, ’ribosom’, ’transcription’”, and the proteins albumin, actin, tubulin, IgG, BRIX1, DDX31, PUM2, NANOS1, RRP36, and PKR were added as putative negative and positive controls. We also included the 50 top proteins that were destabilizing hits in^3^, if not already included. 80aa-long protein tiles were generated in 10aa increments along all proteins and duplicates were removed using custom Python scripts. Annotated Pfam domains for all of the original 1100 proteins were collected using the package Prody, which accesses Interpro through a Python API, and filtered for domains under 80aa. Domains 80aa long were added directly to the list of tiles; those shorter than 80aa were expanded by adding native protein sequence on each side of the selected domain until 80aa was reached. 3,597 random tiles from the library generated in^62^ and 10 well-expressing tiles known to be transcriptional activators or repressors^62^ were added to the resulting 26,932 tiles. All 30,539 tiles were then reverse-translated and codon-optimized as follows: codon use was matched to human codon frequencies; a GC content of 20-70% within 50bp windows and maximum 65% GC content was enforced; BsmBI sites were excluded; C homopolymers greater than 7 in length were excluded. Finally, we appended BsmBI restriction sites and primer handles for PCR amplification to all oligos, resulting in a uniform 300nt length for every library member. The library was ordered as a pool from Twist Biosciences.

#### RBP Hit Library design

A hit library was built out of tiles that had been screened in the RBP library at both the 3’ and 5’ reporters and included the following: all tiles that were hits on both reporters (screen score >1.16 on 3’, 0.3 on 5’); tiles that were only hits on the 3’ reporter (>1.16 on 3’, <0.3 on 5’); the top tiles on the 5’ reporter that were not hits at 3’ (>1.2 on 5’, <1.16 on 3’), and a selection of non-hit tiles from both screens (scores <-0.75 at 3’, <0 on 5’) for a total of 3,149 members. All tiles were again codon-optimized following the constraints as above and ordered as a pool from Twist Biosciences.

#### Pooled library cloning

All Twist oligo pools were resuspended to 10 ng/μL in water, and libraries were selectively PCR amplified using primers specific to their appended handles flanking the sequence of each library member. All reactions were prepared in a pre-PCR hood to reduce contamination. A test qPCR reaction was performed using 25 μL Q5 Ultra II High-Fidelity Polymerase (NEB #M0544L), 0.5 μL library pool, 2.5 μL of each 10 μM library amplification primer, 0.25 μL 20X EvaGreen dye (Fisher Scientific #NC0521178), and water to 50 μL. qPCR was performed on a Bio-Rad CFX machine and was analyzed to extract the half-maximum cycle number for dye saturation using the following protocol: initial denaturation at 98C for 30s; 35 cycles of 98C for 10s, 58C for 20s, and 72C for 30s; final extension at 72C for 2 minutes. Two to six PCRs were then performed, depending on library size, in identical conditions using 17 to 21 cycles depending on the qPCR results per library. Amplified libraries were purified with 0.9X SPRISelect (Beckman Coulter #B23317) and elution in 20 μL.

The pAT031 MCP recruitment lentiviral vector and the pJT126 rTetR recruitment vector (Addgene #161926) were digested with 10,000 U/mL Esp3I (NEB #R0734L) for 15 minutes at 37°C, using 1 μL enzyme per 5 μg plasmid. After heat inactivation at 65°C for 20 minutes, pre-digested vector was run on a 0.5% TAE gel until a linearized band could be extracted using the QIAquick Gel Extraction Kit (Qiagen #28704). Amplified libraries were then cloned into their respective digested vectors using the NEBridge Golden Gate Assembly Kit (BsmBI-v2) (NEB #E1602L) as follows: 20 μL reactions were prepared using 2 μL 10x T4 DNA Ligase Reaction Buffer (NEB #B0202S), nuclease-free water, 75 ng of pre-digested vector, 5 ng of amplified library, and 2 μL assembly kit. 24 reactions were prepared to clone the RBP library; 8 reactions were used for the smaller RBP hit library. Each 20 μL reaction was placed in a thermocycler for 65 cycles of 42°C for 5 minutes and 16°C for 5 minutes, then a final digest at 42°C for 5 minutes and heat inactivation at 70C for 20 minutes. Reactions for each library were pooled and purified using the Zymo Clean&Concentrate DNA kit (Zymo #D4004) eluted in 6 μL of water.

25 μL aliquots of Endura DUO electrocompetent cells (Lucigen #60242-2) were thawed on ice and mixed with 2 μL of the purified Golden Gate product. Mixtures were transferred to Gene Pulse Electroporation Cuvettes with a 0.1cm band gap (Bio-Rad #1652089) and electroporated on a Gene Pulser Xcell Total System (Bio-Rad #1652660) under the following conditions: 1.8kV, 10 uF, 600 Ω, and 0.1 cm distance. Cells were recovered in 2 mL of 37°C SOC recovery medium (NEB #B9020S) at 37°C for 1 hour, after which they were plated across 4-8 10”x10” luria broth agar plates with 100 μg/mL carbenicillin. Plates were incubated at 30°C for 14-18 hours, after which colonies were harvested by scraping and pelleted at 3,500xg for 20 minutes. Plasmid pools were extracted using the Qiagen Plasmid Maxi Kit (Qiagen #12162) and library quality was assessed using Illumina sequencing after PCR amplification from the plasmid pool.

#### High-throughput recruitment assays

K562 cells expressing either the 3’ 24 stem-loop, 5’ 24 stem-loop, 5’ 7 stem-loop, or 9xTetO reporters were infected with their corresponding lentiviral libraries by centrifugation at 1,000xg for 2 hours. Libraries were infected in two replicates at ∼300x infection coverage (starting with 45 × 10^6^ K562 cells for the RBP library screens and 10 × 10^6^ K562 cells for the RBP hit library screens, each at a resulting MOI of 0.3). Cells were treated with 10 μg/mL blasticidin (Gibco #A1113903) starting 48 hours post-infection and were selected for seven to nine days, until at least 94% of cells were positive for lentiviral integration as assessed by flow cytometry (BFP positivity for MCP lentivirus, mCherry positivity for rTetR lentivirus). For screens using the MCP-MS2 recruitment system (both RBP library screens, the RBP hit library re-test, and the 7 stem-loop RBP hit library screen), cells were allowed to recover for 24 hours in blasticidin-free media before magnetic separation and harvest (below). For the TetO-rTetR screen, 1,000 ng/mL doxycycline was added once selection was complete and cells were maintained in doxycycline media for 7 days prior to magnetic separation.

For all libraries, cells were maintained in log growth conditions with daily media changes to ensure dilution to ∼5 × 10^5^ cells/mL and replenishment of blasticidin during selection. Maintenance library coverage was kept as high as possible and >1,000x for all of selection. Cells infected with the RBP library were cultured in 1L spinner flasks with constant paddle rotation; cells infected with the RBP Hit library were maintained in T225 flasks. The half-life of doxycycline in the TetO-rTetR screen was assumed to be 24 hours, and half the amount of doxycycline was replaced each day for the 7 days of recruitment.

#### Magnetic separation

At the end of each recruitment assay, a number of cells equivalent to 12,000X coverage per replicate was removed from the flasks for each replicate and pelleted at 300xg for 5 minutes, then washed twice with DPBS to remove IgG from growth media. Pellets were resuspended in magnetic separation blocking buffer (2% BSA, 2mM EDTA pH 8.0 in DPBS) to a final concentration of 20 × 10^6^ cells/mL. Dynabeads M-280 Protein G (Thermo #10004D) were prepared by incubation on a magnetic stand, removal of supernatant, washing in 5x volume of blocking buffer, and subsequent buffer removal and resuspension in the cell mixture. 90 μL of beads were used per every 10 million cells pelleted. Cell-bead suspensions were incubated at room temperature for 75-90 minutes on a nutator to allow for binding, and then incubated on a magnetic stand for 5 minutes to allow for separation of bead-bound and unbound cells. The ‘unbound’ cell fraction was removed as supernatant and placed in a new tube, which was subsequently re-incubated on the magnet and removed one more time to ensure high purity. The bead-bound fraction was resuspended in the same volume of blocking buffer and re-incubated on the nutator for 15 minutes, after which it was incubated on the magnet, the supernatant discarded, and the beads resuspended as the final ‘bound’ fraction. Bound, unbound, and pre-separated cells were analyzed by flow cytometry for separation purity, pelleted, and frozen at -20°C until further processing.

#### Library preparation and sequencing

Genomic DNA was extracted from all pelleted magnetic separation fragments using the QiaAmp Blood&Cell Culture DNA Maxi Kit (Qiagen #13362) following manufacturer’s instructions, using no more than 1 × 10^8^ cells per column. During extraction, beads were removed from the bead-bound fractions using a magnetic stand post-lysis to avoid column damage, and fractions were eluted in buffer EB (Qiagen #19086) rather than buffer AE to avoid PCR inhibition. Library members were amplified by PCR with primers containing Illumina adapter overhangs. Cycle numbers for PCR were determined by qPCR, in which one test reaction for each fraction was performed with the addition of 0.25 μL 20X EvaGreen dye (Fisher #NC0521178) and was analyzed to extract the half-maximum cycle number for dye saturation. Next, between 8-48 PCR reactions were set up for each fraction, with the number of reactions dependent on both the amount of extracted genomic DNA and on required coverage per library size. Reactions were set up as follows: 5-10 μg genomic DNA, 0.5 μL each primer, 50 μL Q5 Ultra II High-Fidelity Polymerase (NEB #M0544L), water to 100 μL. The thermocycling protocol used was as follows: initial denaturation at 98°C for 3 minutes; 17-25 cycles of 98°C for 10s, 63°C for 30s, and 72°C for 30s; and final extension at 72°C for 2 minutes. All reactions for each fraction were mixed and purified using a double-sided SPRIselect cleanup with an initial 0.5X left-sided cleanup and final ratio of 0.75X. Samples were quantified using the Qubit dsDNA HS Assay Kit (Thermo #Q33231) on a Qubit 4 Fluorometer (Fisher #Q33238), run on an Agilent TapeStation (Agilent #G2964AA) to assess library purity, pooled with 30% PhiX Control v3 (Illumina #FC-110-3001), and sequenced on an Illumina NextSeq 550 with 2×150 cycles, on an Illumina HiSeq 2000 with 2×150 cycles, or on an Illumina MiSeq with 2×150 cycles.

#### High-throughput recruitment sequencing analysis

All recruitment assay sequencing data was processed and analyzed using the HT-recruit-Analyze pipeline from (Tycko), available on GitHub (https://github.com/bintulab/HT-recruit-Analyze). Briefly, raw sequencing reads were demultiplexed using bcl2fastq (Illumina) and aligned using ‘makeCounts.py’ to a reference created with ‘makeIndices.py’. The aligned reads were used to compute enrichment scores from the unbound (OFF) to bound (ON) populations for each library member using ‘makeRhos.py’. Depending on the sequencing depth of the assay, library members with fewer than 50-500 reads summed between replicates were excluded from further analysis. The hit threshold for each screen was determined either by using 3 standard deviations above the mean score of the random tile control population, or by adjustment for wide distribution of random control scorers as described in **Results**.

#### Regulatory domain annotation

Only the unbiased tiling screens (3’ and 5’ RBP library screens) were used to assess putative regulatory domains from tile scores. The starts of new domains were defined as the first tile in a string of two or more consecutive hit tiles. If a tile had dropped out of the screen due to low sequencing depth but the tiles on each side of it were hits, the missing tile was considered part of the same contiguous regulatory domain. Domain ends were annotated where the next successive tile in an ongoing domain was no longer a hit. The extended domain length was considered to be the first amino acid of the first hit tile to the last amino acid of the last hit tile. Single hit tiles were not considered to be domains unless they were the most N- or C-terminal tile of a given protein. Minimized regulatory domains were computed as the sequences fully contained within tiles that downregulated the RNA reporters. Amino acids not contained by two or more tiles in the same domain were excluded, leaving the last ten amino acids of the first hit tile spanning until the first ten amino acids of the last hit tile as the minimized region.

#### Individual recruitment assays

Individual protein tiles that were selected for low-throughput validation were ordered as gene fragments from IDT and cloned into either the pAT031 (MCP) or pJT126 (rTetR) recruitment backbones using Golden Gate cloning. K562 cells expressing the appropriate reporter lines were transduced with lentivirus (prepared as described above) and selected with blasticidin (10 μg/mL) beginning 48 hours after infection. Selection continued for 5-9 days or until cells were >95% BFP/mCherry positive, after which the MCP lines were analyzed for Citrine fluorescence in biological replicate using a Bio-Rad ZE5 cytometer measuring >10,000 cells per sample. rTetR lines were split into two plates (all in biological replicate); one plate was left untreated and one plate was treated with 1,000 ng/mL doxycycline with half-media and doxycycline changes performed every day. Cells were analyzed using flow cytometry for Citrine fluorescence every day of doxycycline treatment. Data was analyzed using Cytoflow^63^ and additional analyses and visualizations were performed using custom Python scripts. All cells were gated for live cells, singlets, and mCherry/BFP positivity; from there, either an MCP-only or an rTetR-control/no-dox control was used as a negative control to compute the fraction of Citrine ON and OFF cells. These OFF scores were compared to screen enrichment ratios using a logistic expression; because OFF scores are not related to sequencing depth, they are a better metric for comparing between screens performed at different times (enrichment ratios are calculated having normalized for raw reads, but different screens can have different dynamic ranges depending on library number and magnetic separation purity that makes relative values consistent but absolute values difficult to compare).

#### Protein compositional analysis, motif finding, and structural analysis

Amino acid composition and compositional biases were calculated by comparing the frequency of each amino acid in a group of interest (the top 195 hit tiles, unstructured hit tiles, and structured hit tiles) to that same amino acid frequency in a reference dataset (195 non-hit tiles, randomly chosen). Predicted structures were computed using the Jpred4 server^22^, which assigns each amino acid of the submitted 195 hit or 195 non-hit sequences as ‘helical,’ ‘sheet,’ or ‘unstructured.’ Motif finding analyses of each of the above groups was performed using the MEME suite server^23^, taking the top 3 most confident motifs and excluding overlap between tiles as putative motifs.

#### HCR-Flow-RNA-FISH for Citrine reporter sequence

HCR-FlowFISH was performed as described in^64^. All reagents were prepared using the Molecular Instruments HCR RNA-FISH Bundle with custom probes against the Citrine reporter mRNA and Amplifier B3 (Alexa-647 fluorophore). Briefly, 2.5 × 10^6^ cells per condition were pelleted and resuspended in 4% formaldehyde, then fixed for one hour at room temperature with agitation. Cells were washed 4 times with PBST and resuspended in 70% cold ethanol, incubated at 4°C for 10 minutes, and washed twice again with PBST. Pellets were resuspended in Probe Hybridization Buffer and mixed with custom probes to incubate overnight at 37°C. After overnight incubation, cells were washed 4 times in Probe Wash solution, resuspended in 5x SSCT, and incubated at room temperature for 5 minutes, then spun down and resuspended in Amplification Buffer. While cells were incubating in Amplification Buffer for 30 minutes at room temperature, fluorescent hairpins were prepared by snap cooling from 90°C to room temperature in the dark for 15 minutes. Hairpins were added to cells and incubated for 3 hours in the dark to allow the HCR to occur. Finally, cells were washed 6 times with SSCT, resuspended in 500 mL PBS, and analyzed on a BioRad ZE5 cytometer for Citrine and Alexa-647 fluorescence.

#### RT-qPCR against Citrine reporter mRNA

RT-qPCR was performed using iScript reverse transcription mix (Bio-Rad #1708841) and the SsoAdvanced SYBR Green Supermix (Bio-Rad #1725271), following manufacturer’s specifications. Briefly, RNA was extracted from cell samples using the RNEasy+QiaShredder kits (Qiagen #74106); 500 ng of RNA per sample was added to iScript reverse transcription master mix and incubated for 5 minutes at 25°C, 20 minutes at 46°C, and 1 minute at 95°C. The resulting cDNA was diluted 1:2 and 2 microliters were carried forward into the qPCR reaction, performed using a BioRad CFX Connect Real-Time system (Bio-Rad #1855201). Data was analyzed using custom Python scripts. Primer sequences targeting the Citrine reporter sequence are as follows: Fwd, CCACCTTCGGCTACGGCCTGA; Rev, GCCATGATATAGACGTTGTGG.

#### Preparation of degron-expressing cell lines

Cell lines expressing DHFR-MCP-tile fusions were selected for cells that responded completely to TMP inhibition of the degron. To isolate these populations, 10 μM TMP was added for 4 days, resulting in a bimodal population of Citrine fluorescence. The fully silenced cells were sorted and allowed to reactivate the mRNA reporter through TMP washout and degradation, after which the sorted and reactivated cells were able to fully repress Citrine translation upon addition of TMP.

#### Degron inhibition and HaloTag staining

For all cell lines expressing DHFR-MCP-tile fusions, 10 μM TMP was used as a saturating dose for degron inhibition. A no-TMP (0 μM) control was included for all TMP dosing experiments. HaloTag staining was performed using the Janelia Fluor 646 HaloTag Ligand (Promega #GA1121) as follows: ligand was resuspended in 35.5 μL DMSO to prepare a 200 μM stock solution. The stock solution was diluted to 200 nM in warm RPMI. 2 × 10^5^ cells were pelleted per sample and resuspended in 200 μL diluted ligand solution. Samples were incubated for 15 minutes in a 37°C 5% CO2 incubator, after which they were pelleted and resuspended in 200 μL RPMI for flow cytometry analysis.

#### Transcriptional inhibition experiments

Actinomycin D was used to inhibit transcription and measure RNA degradation rates. At each timepoint, TMP at a final concentration of 10 μM and actinomycin D at a final concentration of 1 μg/mL were added to 1 × 10^6^ cells. Cells were harvested at the end of the timecourse and HCR-Flow-RNA-FISH was performed as described above to measure Citrine mRNA levels. Degradation rates and half-lives were calculated using custom Python scripts.

#### CUT&RUN for detection of H3K9me3

CUT&RUN was performed as described in^39^, using the CUTANA CUT&RUN Kit (14-1048, EpiCypher) and Abcam anti-H3K9me3 antibody (Abcam ab176916). An input of 5 × 10^5^ cells per sample were processed according to the manufacturer’s protocol. Digitonin was used at a final concentration of 0.01% for nuclear permeabilization. Sequencing libraries were prepared and dual-indexed using Illumina adapters (in the CUTANA kit). Libraries were quantified with the Qubit dsDNA HS Assay Kit and fragment sizes assessed using an Agilent TapeStation. Libraries were sequenced using a NextSeq 550 system from Illumina. A custom human genome (hg38) with the reporter integration added was constructed using bowtie2-build. Alignment was performed using bowtie2, and Picard was used to remove duplicate reads. Bedgraph files were generated using bedtools and reads were normalized by total counts per sample, and reported as counts per million. Further analysis was performed using custom Python scripts. Processing scripts are available at https://github.com/bintulab/Spreading_Lensch_2022/tree/main/CUT%26RUN%20Analysis.

#### RBP-mediated RNA degradation model

We derived deterministic equations for the Citrine levels of cells where transcriptional RBP-mediated RNA degradation affects the average mRNA, and consequently Citrine, level of the cell population. We first derived a differential equation describing transcription of the reporter gene, where ***k_trx_*** is the rate of mRNA production from the reporter locus, ***k_deg_*** is the RBP-independent rate of mRNA degradation, ***k_reg_*** is the RBP-mediated rate of mRNA degradation, and mRNA is the concentration of mRNA in the cell:

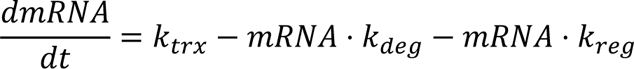

Assuming the maximum mRNA level in the cell occurs when *k_reg_* = 0, the steady-state mRNA levels normalized to the maximum mRNA are thus given by:

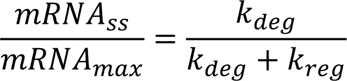

***k_reg_*** is dependent on the average amount of RBP bound to the mRNA, which we model with a Hill binding function. *n* is the Hill coefficient indicating the degree of binding cooperativity and 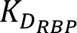 is the binding affinity of RBP to mRNA:

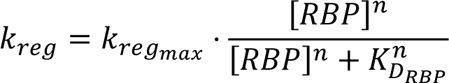

We additionally derived an equation for the concentration of RBP in the cell given TMP concentration with the assumption that the addition of TMP directly inhibits the DHFR degron and controls the amount of stabilized RBP available to bind in the cell:

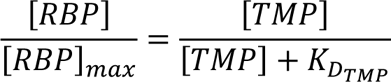

We fit the equation above using scipy.optimize.curve_fit() to extract 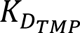, the dependence of RBP stabilization on TMP addition, from HaloTag staining data on DHFR-tagged MCPs at varying concentrations of TMP.

Therefore, we can express ***k_reg_*** as follows:

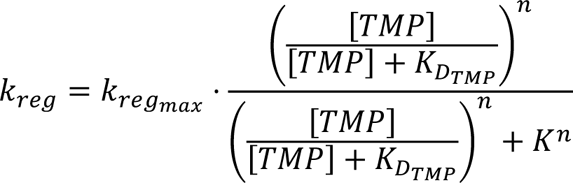

where ***k_reg_max_*** is the maximum rate of RBP-mediated RNA degradation in units days^-1^, 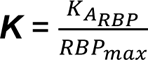 (unitless) is the effective association rate of the RBP to RNA, and ***n*** is the Hill coefficient.

Next, we incorporated translation into the model. We derived an equation for the production of Citrine protein from mRNA with the rates ***k_trl_***, the constant rate of translation, and ***k_deg_protein_***,the constant rate of protein degradation and dilution:

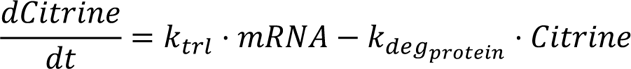

Solving the equation 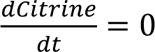, we determine the steady state Citrine level in the cell population:

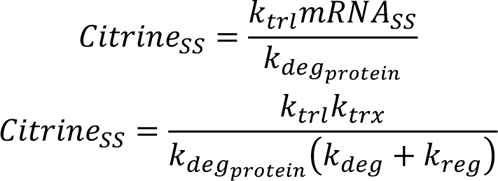

We again assume Citrine levels are maximum when *k_reg_* = 0. Therefore, Citrine_SS_/Citrine_max_ is equivalent to mRNA_SS_/mRNA_max_:

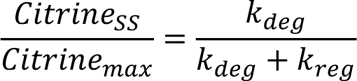

We then solved the system of differential equations with the initial condition Citrine(0) = Citrine_max_ to determine an equation for Citrine(t)/Citrine_max_:

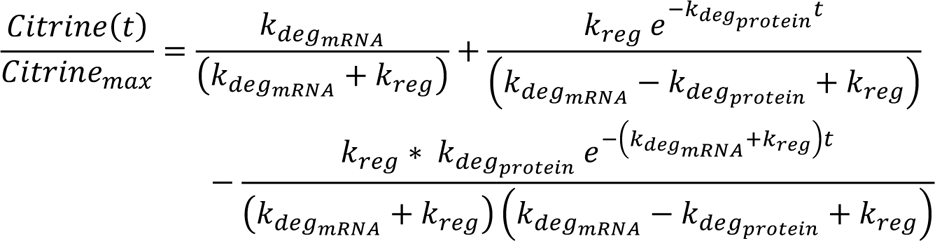

The values of *a*, *b,* and *n* for synNANOS are extracted by fitting the equation for normalized steady-state Citrine levels using scipy.optimize.curve_fit() to normalized Citrine levels after 4 days of TMP addition and subsequent RBP recruitment. We then substituted *a, b,* and *n* into the equation for ***k_reg_*** and used this value to calculate normalized Citrine levels over time for varying TMP doses.

The fit of each model was assessed by calculating the Root Mean Squared Error (RMSE) to quantify the difference between predicted and measured values. The RMSE was compared to the standard deviation (SD) of the measured values.

#### KRAB-dependent transcriptional silencing model

First, the rate of Citrine degradation and dilution was estimated using the following equations fit to 5 days of KRAB recruitment and corresponding measurements of Citrine fluorescence:

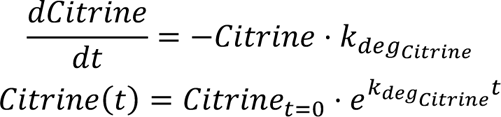

using scipy.optimize.curve_fit().

We derived the change in Citrine fluorescence due to transcriptional silencing from the following system of differential equations. We first derived a differential equation for the rate of silencing on the gene level dependent on **k_s_,** the rate of transcriptional silencing:

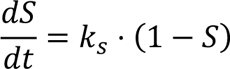

The rate of transcriptional silencing was defined as dependent on dox concentration in an ultrasensitive manner with Hill coefficient ***n***:

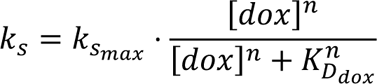

We derived differential equations describing transcription of genes in the active (non-silent) state and translation of the mRNA to Citrine:

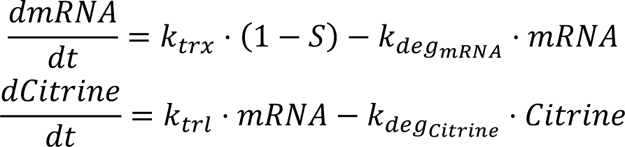

We then solved the system of differential equations with the initial condition Citrine(0) = Citrine_max_ to determine an equation for Citrine(t)/Citrine_max_:

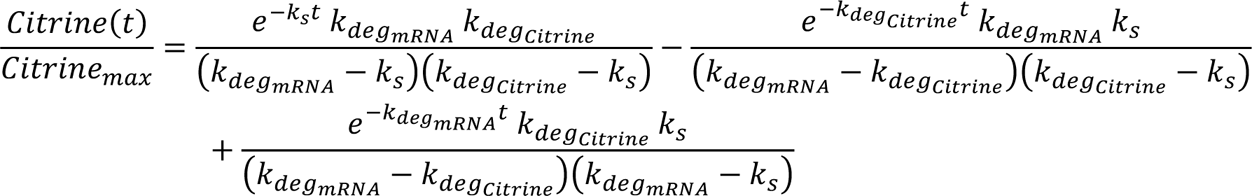

Scipy.optize.curve_fit() was used to fit the resulting function to 5 days of KRAB recruitment, and free parameters ***k_s_, k_s,max_,*** and ***n*** were extracted for each dox dose.

#### Stochastic Simulation of Simultaneous RBP-Mediated Degradation and KRAB Recruitment

Stochastic simulations were performed according to the Gillespie algorithm^65^. Populations of ***n_cells*** were simulated expressing both an RBP RNA degrader and rTetR-KRAB repressor. During the active state, the reporter gene produced mRNA at rate ***k_trx_***. The reporter gene could become silenced at rate ***k_s_*** dependent on dox concentration. In all states, mRNA could be degraded at rate k_reg_ dependent on TMP concentration. The base constants were chosen as follows: {***k_trx_***: 50, ***k_deg_***: 2.39, ***k_reg_max_***: 10.8, b:0.99, ***n_RBP_***: 0.72, ***k_s_max_***:2.1, ***n_KRAB_***: 2.7, ***k_A_TMP_***:1.19, ***k_A_dox_***:7.8}.

#### Data Analysis and Statistics

Statistical analyses were performed in Python using the SciPy package, are two-sided (unless otherwise stated), and are indicated in text or figures/figure legends. “N” for each analysis is indicated in the text, figures, or legends where appropriate. No methods were used to determine whether the data met assumptions of the statistical approach.

#### Data and code availability

Raw HT-recruit and CUT&RUN sequencing files (FASTQs) have been deposited at NCBI SRA at project number PRJNA1112784. All other data reported in this paper will be shared by the lead contact upon request. Any additional information required to reanalyze the data reported in this paper is available from the lead contact upon request.

## Supplemental Figures

**Figure S1.**
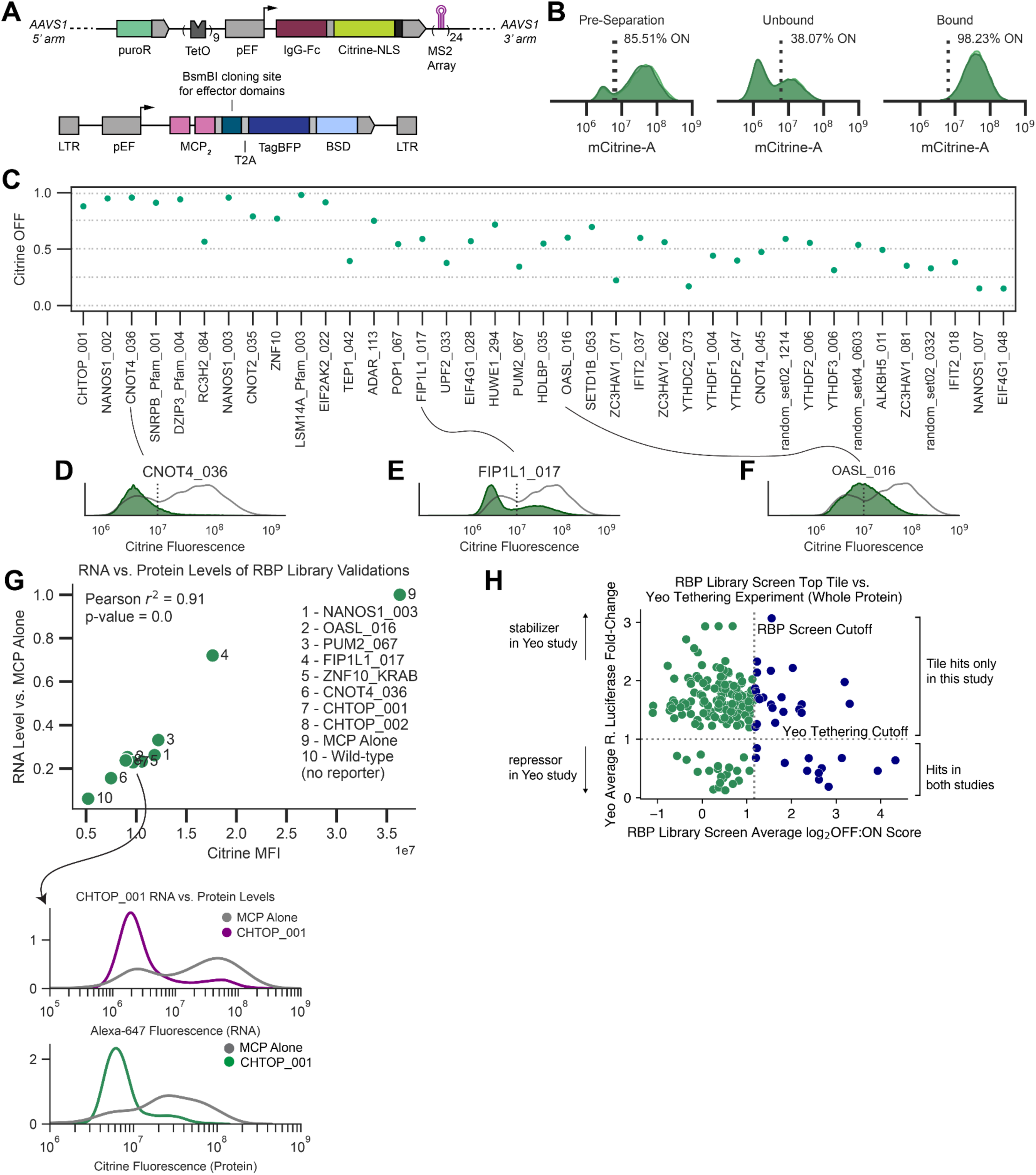
RBP library screen details, related to Figure 1. (A) Full construct schematic of the RNA recruitment reporter integrated into the *AAVS1* safe harbor locus in the first intron of the *PPP1R12C* gene (top) and MCP-fusion vector for pooled domain cloning (bottom). puroR = puromycin resistance, TetO = tetracycline repressor binding sites, IgG-Fc = immunoglobulin G constant region, BSD = blasticidin resistance, LTR = long terminal repeats for lentiviral integration. (B) Flow cytometry distributions on day 10 of recruitment per replicate RBP library screen cells before and after magnetic separation. (C) Summary of RNA downregulation for individually tested screen tiles as measured by flow cytometry, reported in fraction of cells OFF. Tiles are ranked (L-R) by decreasing screen score. Vertical error bars are standard deviation of two biological replicates. (D-F) Example flow cytometry distributions for tested tiles of varying strength: CNOT4_036, FIP1L1_017, OASL_016 (L-R, green fill) versus MCP alone (grey line, no fill). (G) Protein vs. RNA level measurements for 8 selected validations, measured in Citrine fluorescence by flow cytometry (x-axis) and Alexa-647 fluorescence by flow cytometry of HCR-RNA-Flow-FISH (y-axis, normalized to levels of cells expressing MCP alone). Below, example Alexa-647 and mCitrine distributions for RNA and protein level measurements, respectively, of CHTOP_001 and MCP alone. (H) Comparison of 195 proteins tested in both this study and in^3^, with x-axis the recruitment screen score of the top tile tested in this study and y-axis the RNA fold-change of the tethered full protein tested in^3^. Green dots, non-hits in this study; blue dots, hits in this study. Horizontal dashed line is cutoff in previous study (below dashed line = downregulation hit), vertical dashed line is cutoff in our screen.

**Figure S2.**
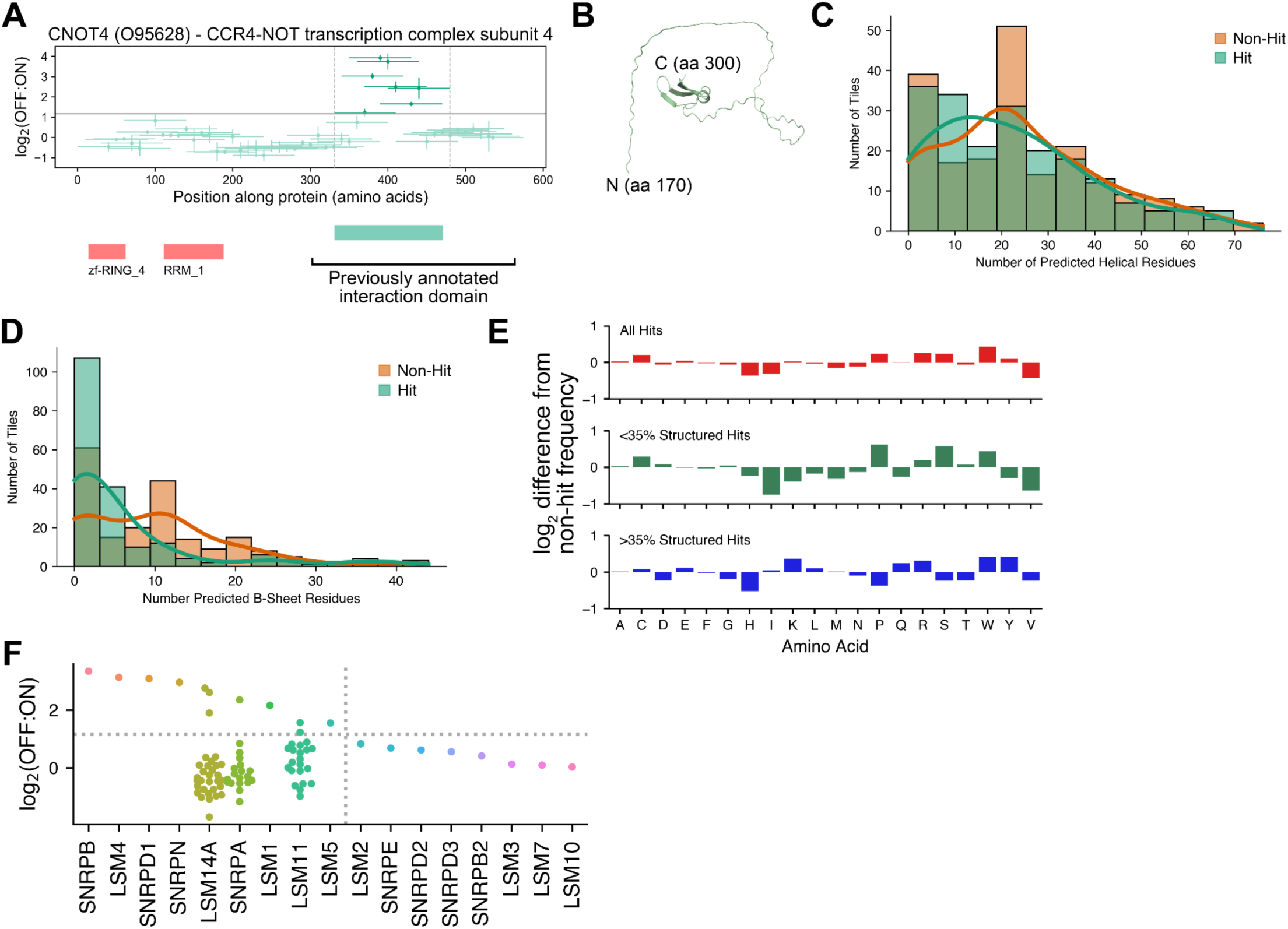
Domain analyses and visualization, related to Figure 2. (A) Tiling plot for CNOT4, showing previously annotated CNOT1 interaction region and more specifically annotated regulatory domain in this study. (B) Alphafold predicted structure of PKR regulatory domain alone. (C) Predicted number of alpha-helical residues for 195 non-hit (orange) vs. hit (green) tiles. (D) Predicted number of B-sheet residues for 195 non-hit (orange) vs. hit (green) tiles. (E) Bar graphs of amino acid frequency enrichment in 195 hit tiles over frequencies in non-hit tiles. Top, all 195 hits; middle, hits with <35% structured residues; bottom, hits with >35% structured residues. (F) Plot of all tiles (one dot = 1 tile) tested from LSm-domain-containing proteins in the RBP library screen. Horizontal dashed line: screen hit cutoff; vertical dashed line: delineating proteins with LSm domain hits (left) from those without (right).

**Figure S3.**
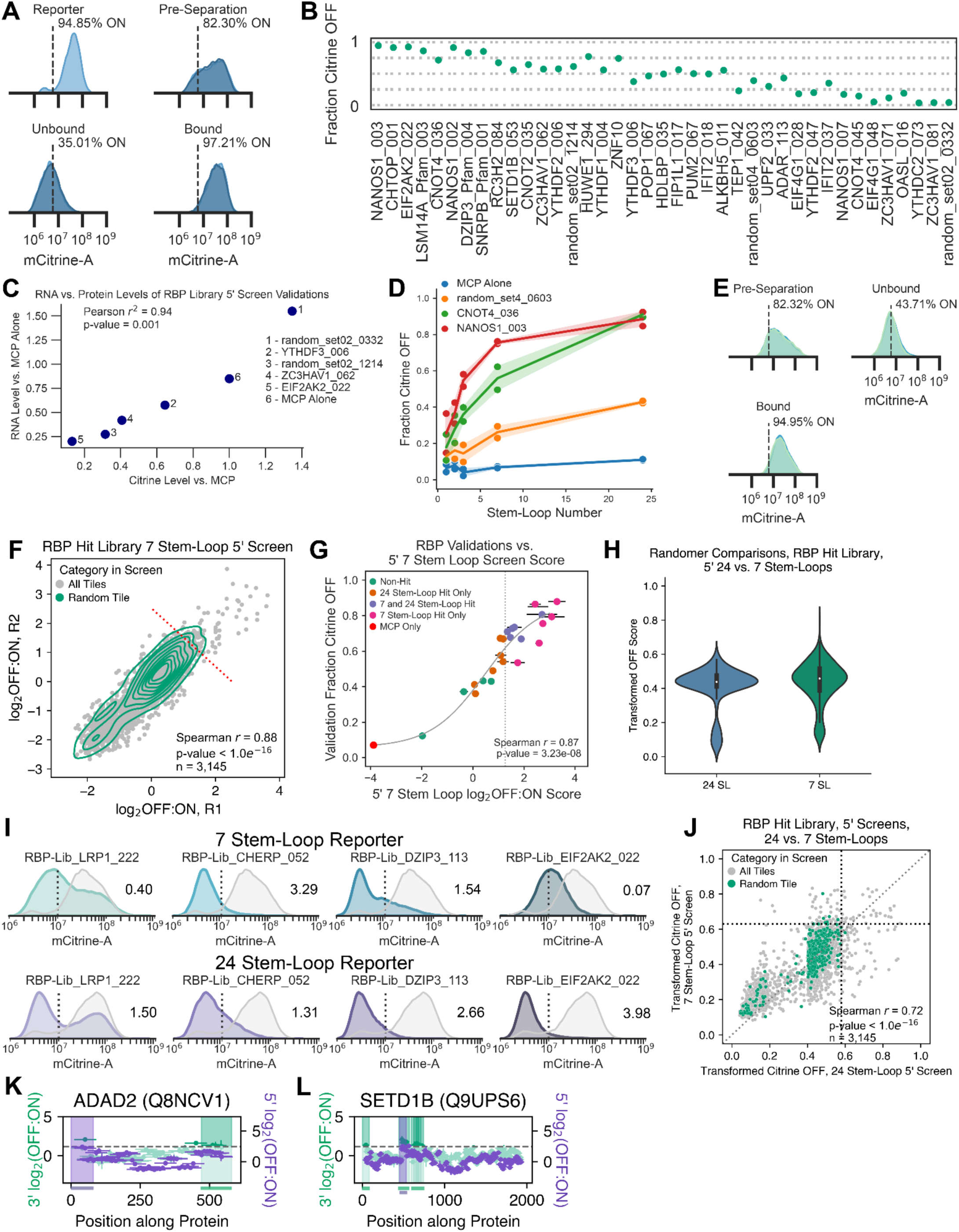
5’UTR screen details, related to Figure 3. (A) Flow cytometry distributions on day 10 of recruitment for replicate 24 stem-loop 5’UTR RBP library screen cells before infection and before and after magnetic separation. (B) Summary of RNA downregulation for individually tested screen tiles as measured by flow cytometry, reported in fraction of cells OFF. Tiles are ranked (L-R) by decreasing screen score. (C) Protein vs. RNA level measurements for 5 selected validations, measured in Citrine fluorescence by flow cytometry (x-axis) and relative values of qPCR against Citrine (y-axis, normalized to Ct values of cells expressing MCP alone). Vertical error is standard deviation of 3 technical replicates, horizontal is standard deviation of two biological replicates. (D) Fraction Citrine OFF (y-axis) of cells harboring 5’UTR reporters with changing numbers of MS2 loops (x-axis), each with MCP alone or 3 different selected tiles. Shading is standard deviation from two biological replicates. (E) Flow cytometry distributions on day 10 of recruitment for replicate 5’UTR 7 stem-loop RBP Hit Library screen cells before and after magnetic separation. (F) Log_2_(OFF:ON) enrichment scores plotted per replicate of the RBP Hit Library (n = 3,145) screen at 7 stem-loops in the 5’UTR. Grey dots, all tiles; green contour, random control; red dashed line, screen threshold (mean + 1.5 standard deviation of random population). (G) Individual validation measurements for 22 selected tiles in 7 stem-loop 5’UTR reporter cells. Tiles are colored by whether or not they were hits in the original 24 stem-loop 5’UTR screen, the 7 stem-loop 5’UTR screen, both, or neither. MCP alone is shown in red. Error bars are standard deviation of two biological replicates. (H) Distributions of transformed OFF screen scores for all random control tiles in the 5’UTR 24 (left) and 7 (right) stem-loop screens. (I) Example flow cytometry distributions for 4 selected tiles (blue/purple fill) at 7 stem-loops (top) or 24 stem-loops. MCP alone for each reporter line is shown in grey fill. Screen scores are reported for each tile on each reporter to the right of its distribution. (J) All RBP Hit Library members (n = 3,145) plotted with their transformed screen score in the 5’UTR at 24 stem-loops (x-axis) versus 7 stem-loops (y-axis). Grey dots, all tiles; green dots, random controls; vertical dashed line, transformed 24 stem-loop high threshold; horizontal dashed line, transformed 7 stem-loop threshold. (K) Tiling plot for ADAD2. (L) Tiling plot for SETD1B.

**Figure S4.**
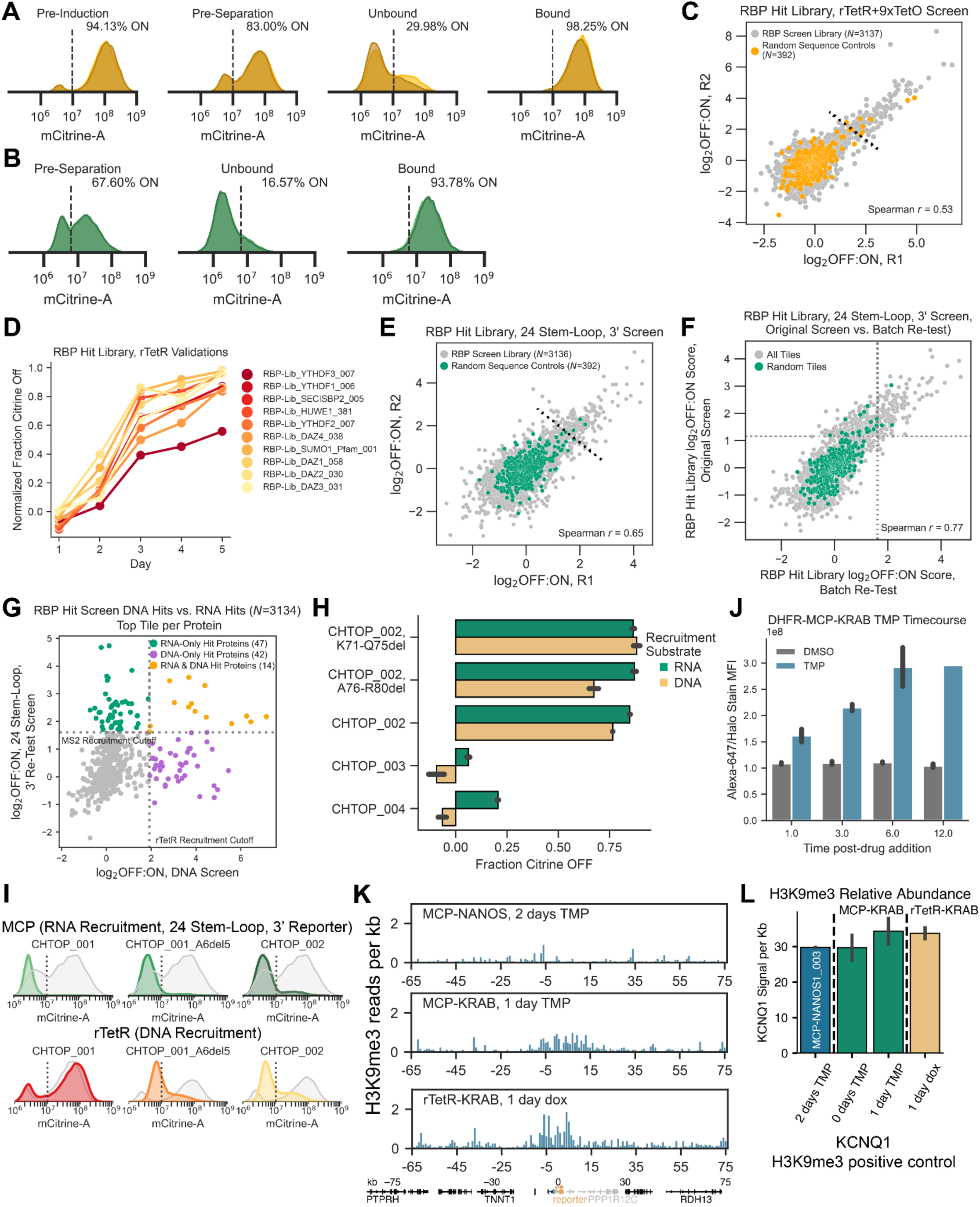
DNA screen and followup details, related to Figure 4. (A) Flow cytometry distributions for no-dox control cells (pre-induction) and on day 7 of dox-mediated recruitment for replicate rTetR-TetO RBP hit library screen cells before and after magnetic separation. (B) Flow cytometry distributions on day 10 of recruitment for replicate 24 stem-loop 5’UTR RBP hit library screen cells before and after magnetic separation. (C) Log_2_(OFF:ON) enrichment scores plotted per replicate of the RBP hit library screen in the rTetR-TetO system. Grey, all tiles; yellow, random controls; black dashed line, screen cutoff (mean + 3 standard deviations of random population). (D) Individual validation measurements of 10 selected hit tiles over 5 days of dox recruitment in rTetR-TetO reporter cells, reported as fraction Citrine OFF for each line normalized to its paired no-dox recruitment control (E) Log_2_(OFF:ON) enrichment scores plotted per replicate of the RBP Hit Library screen on MCP at 24 stem-loops in the 3’UTR. Grey, all tiles; green, random controls; black dashed line, screen cutoff (mean + 3 standard deviations of random population). (F) Average log_2_(OFF:ON) scores for RBP hit library members in the batch retest (x-axis) vs. the original RBP library screen (y-axis). Grey, all tiles; green, random controls; vertical dashed line, batch retest cutoff; horizontal dashed line, original screen cutoff. (G) Average log_2_(OFF:ON) scores for the top scoring tile in the DNA (x-axis) or RNA (y-axis) screens per RBP tested. Purple, proteins with DNA hit tiles; green, proteins with RNA hit tiles; yellow, proteins with hit tiles in both screens. (H) Summary of individual flow cytometry measurements of additional CHTOP tiles and deletions. Green, fraction Citrine OFF when tested fused to MCP in 24 stem-loop, 3’UTR reporter cells; yellow, fraction Citrine OFF when tested fused to rTetR in TetO reporter cells. Error bars are standard deviations of two biological replicates. (I) Flow cytometry distributions for selected CHTOP tiles and deletions on RNA (top, green fill) and DNA (bottom, red/yellow fill) vs. MCP alone (grey fill, top) and no dox (grey fill, bottom) recruitment controls. (J) HaloTag fluorescence of DHFR-MCP-ZNF10_KRAB after various hours of TMP (blue) or DMSO (grey, control) addition, as measured by JaneliaFluor-647 fluorescence after staining of the HaloTag inserted C-terminal to MCP. Error bars are standard deviations of two biological replicates. (K) Genome traces showing normalized CUT&RUN reads against H3K9me3 as a function of distance around the 5’UTR, 7 stem-loop reporter integration site (0 kb on x-axis) after recruitment of MCP-NANOS1_003 (top), MCP-ZNF10_KRAB (middle), or rTetR-KRAB (bottom). Bar plots are average of biological replicates. Bottom, schematic of genomic locus. (L) Quantification of H3K9me3 levels at positive control *KCNQ1* (known high levels of H3K9me3 modification in K562 cells) across the same conditions shown in Fig. 4H.

**Figure S5.**
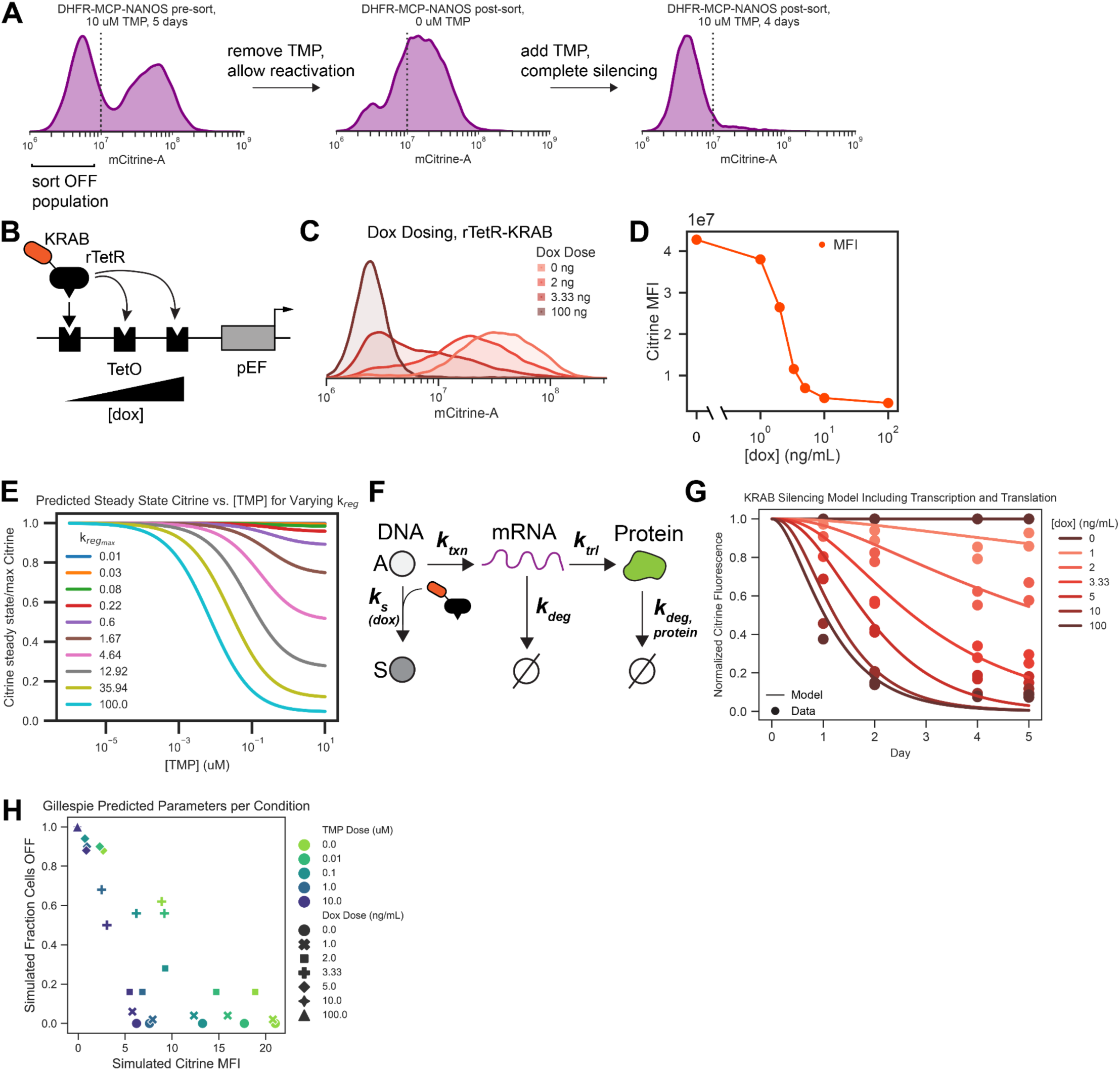
Details of TMP-mediated recruitment and modeling, related to Figure 5. (A) Flow cytometry distributions of 7 stem-loop, 5’UTR reporter cells expressing synNANOS after initial addition of 10 μM TMP, after sorting the silenced cells and allowing reactivation for 5 days, and after re-addition of 10 μM TMP for 4 days (L-R). (B) Schematic of dose-tunable rTetR construct, in which rTetR-KRAB binds at higher occupancy for increasing doses of dox. (C) Flow cytometry distributions of rTetR-KRAB recruited to the same reporter as in Fig. 5B at varying dox doses. (D) Quantification of mean Citrine fluorescence levels for all doses of dox. (E) Predicted Citrine levels for varying levels of model extracted parameter ***k_reg,max_***, proxy for maximum RNA degradation rate upon saturating expression of an RBP. (F) Overview of KRAB-mediated transcriptional silencing model: the gene can either be in the active (A) or silenced (S) state, the transition between which is controlled by a transcriptional silencer at a dox-dependent rate ***k_s_***. Cells can produce mRNA in the A state at a constant rate ***k_txn_***; mRNA is then degraded at the constitutive rate ***k_deg_.*** Protein is made and degraded at the constant rates ***k_trl_*** and ***k_deg,protein_***, respectively. (G) Two-state transcription model fit to Citrine levels after recruitment of rTetR-KRAB at varying dox doses over time. Model RMSE=0.08, Data SD=0.34. (H) Summary of predicted Citrine MFI and fraction cells with Citrine OFF from Gillespie simulation of the dual transcriptional/post-transcriptional regulator system at varying TMP doses (RBP control, colors) and dox doses (transcriptional repressor control, shapes).

