## Supplemental Data 1 for "High-throughput discovery of regulatory effector domains in human RNA-binding proteins"

### EIF2S2 (P20042) - Eukaryotic translation initiation factor 2 subunit 2

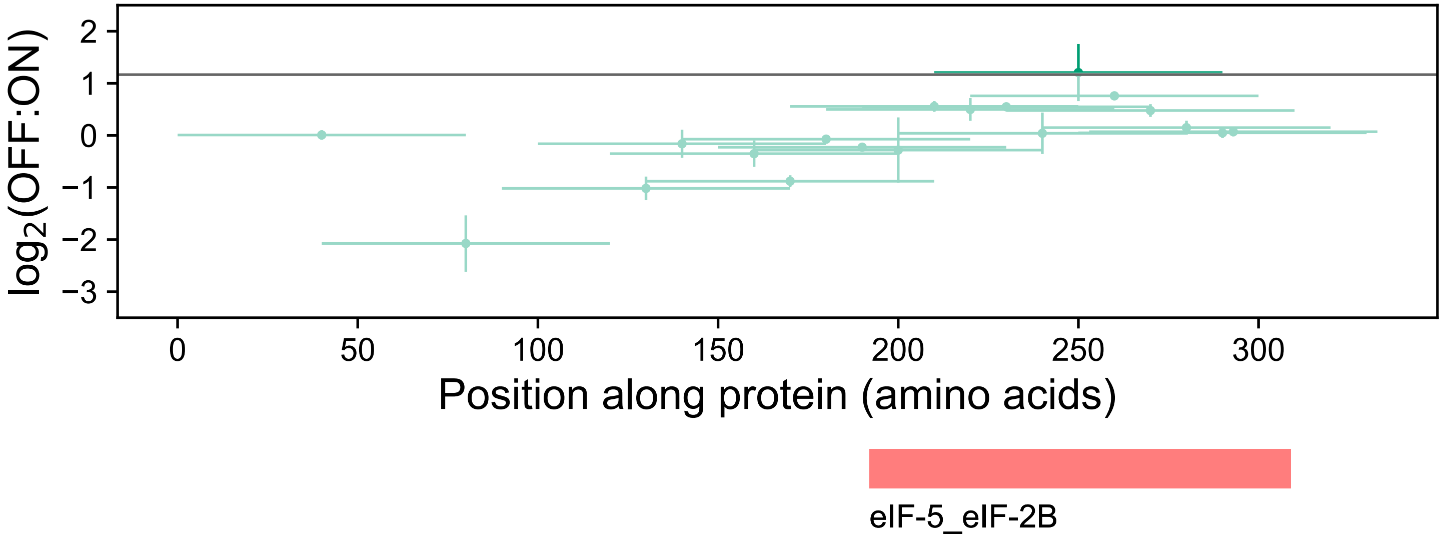

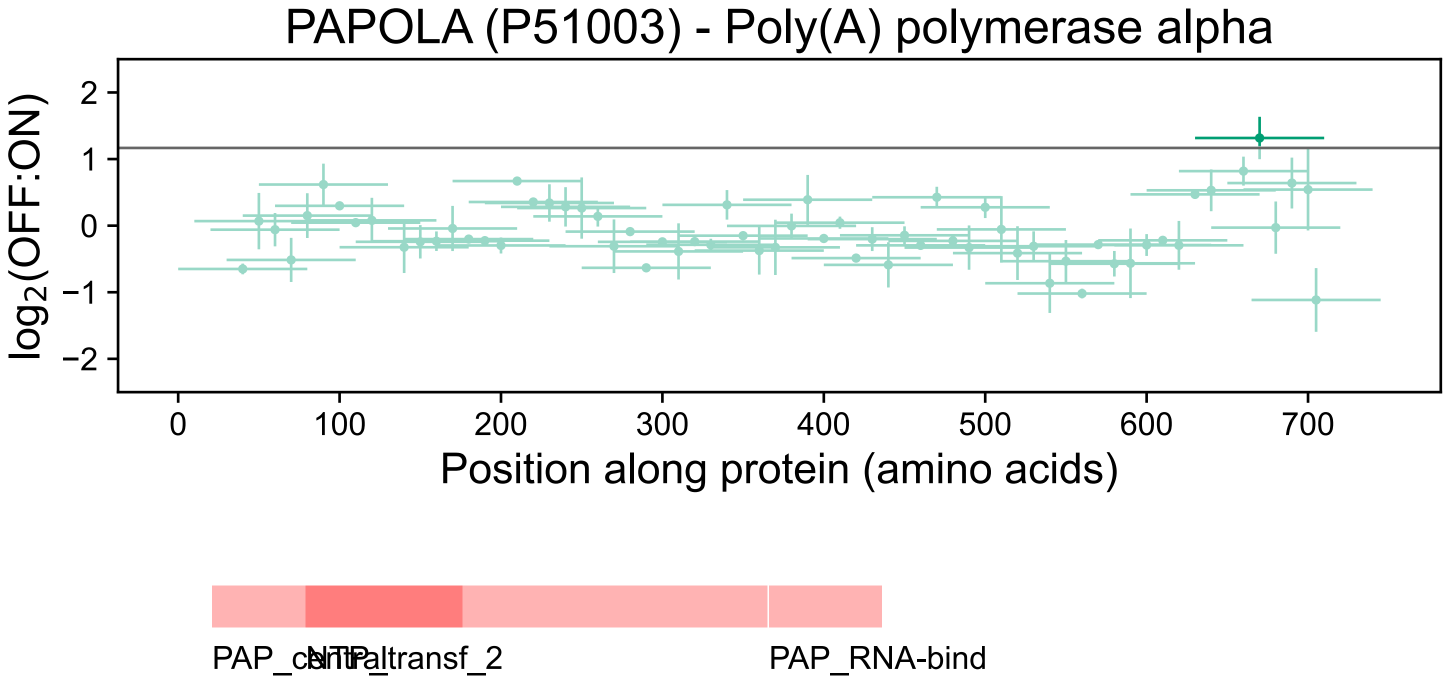

PAPOLA (P51003) - Poly(A) polymerase alpha
log2(OFF:ON)
2
1
0
−1
−2
0
100
200
300
400
500
600
700
Position along protein (amino acids)
PAP_central
NTP_transf_2
PAP_RNA-bind

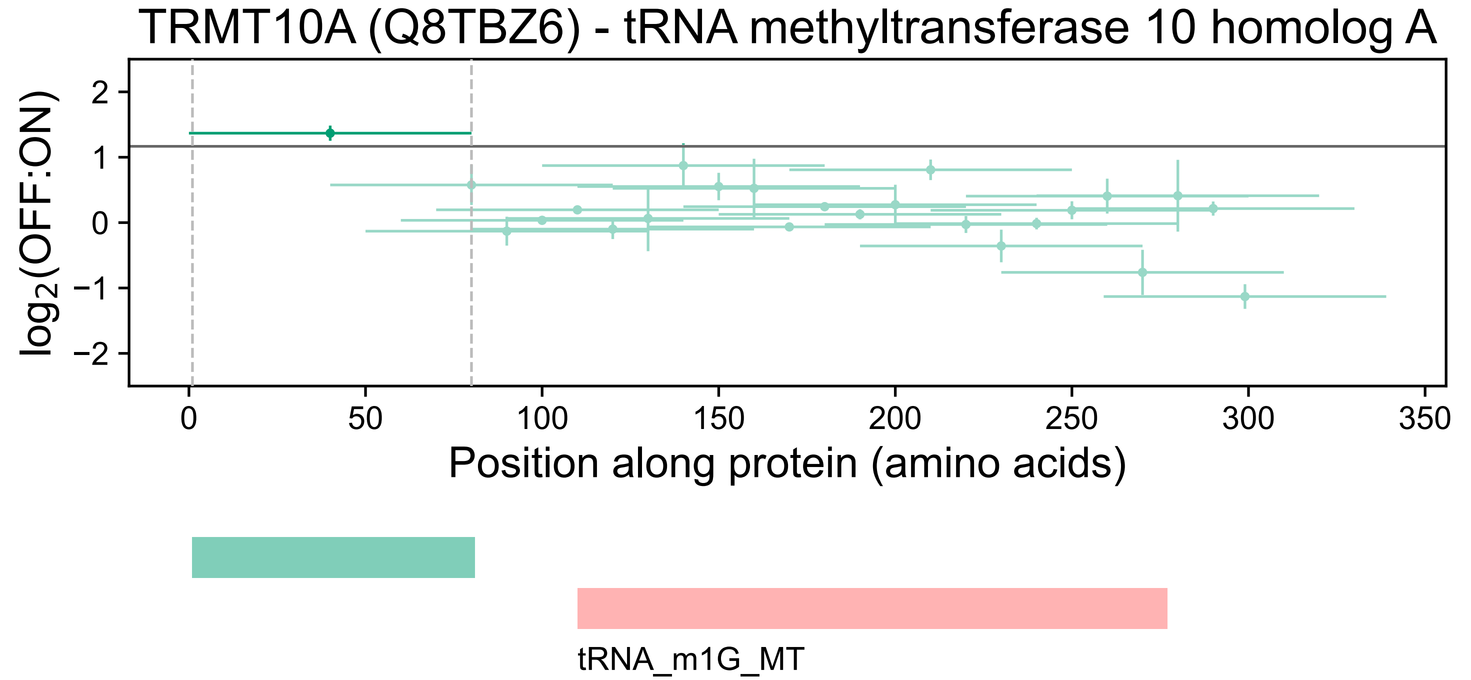
TRRMT10A (Q8TBZ6) - tRNA methyltransferase 10 homolog A
log₂(OFF:ON)
2
1
0
−1
−2
0
50
100
150
200
250
300
350
Position along protein (amino acids)
tRNA_m1G_MT

PTCD3 (Q96EY7) - Pentatricopeptide repeat domain-containing protein 3, mitochondrial

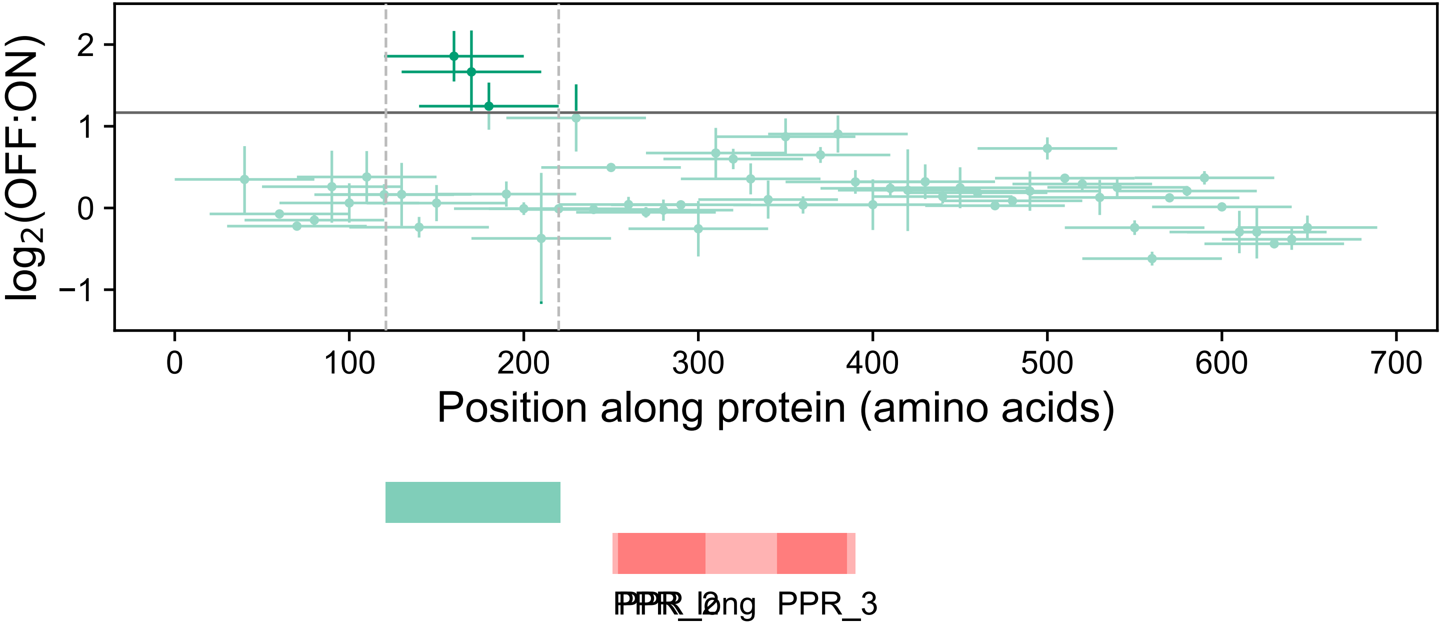

### PATL1 (Q86TB9) - Protein PAT1 homolog 1

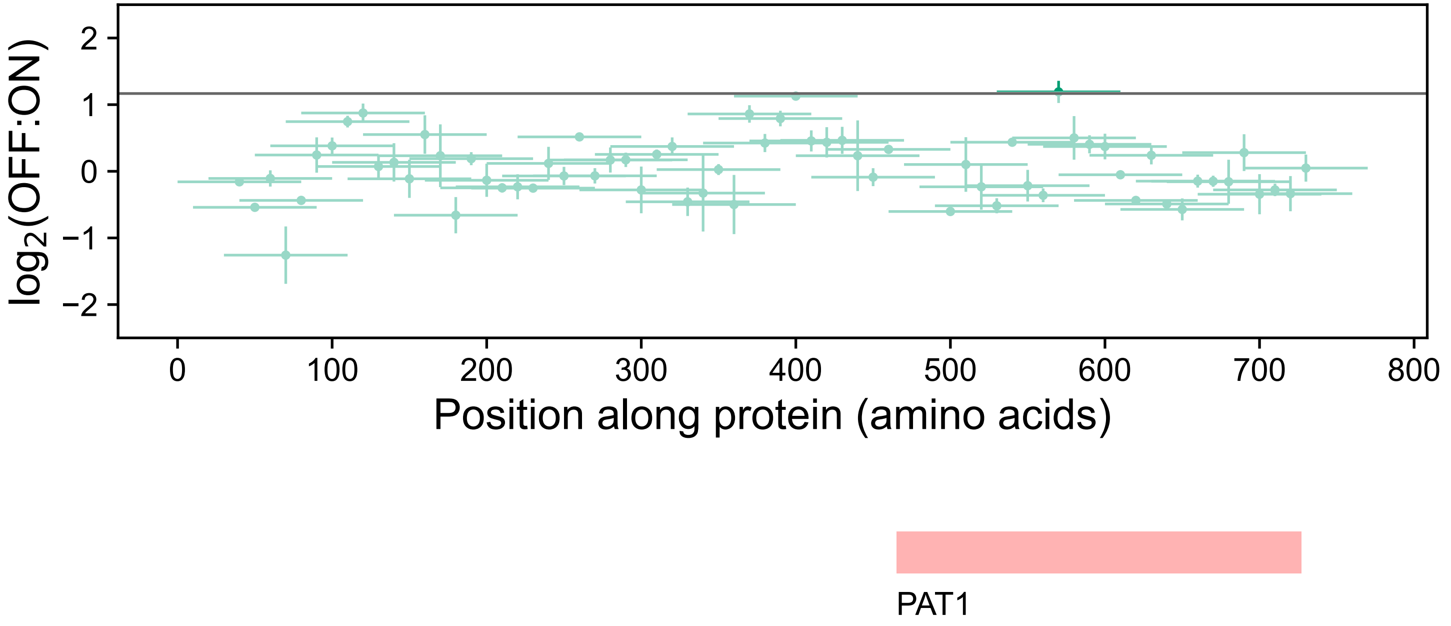

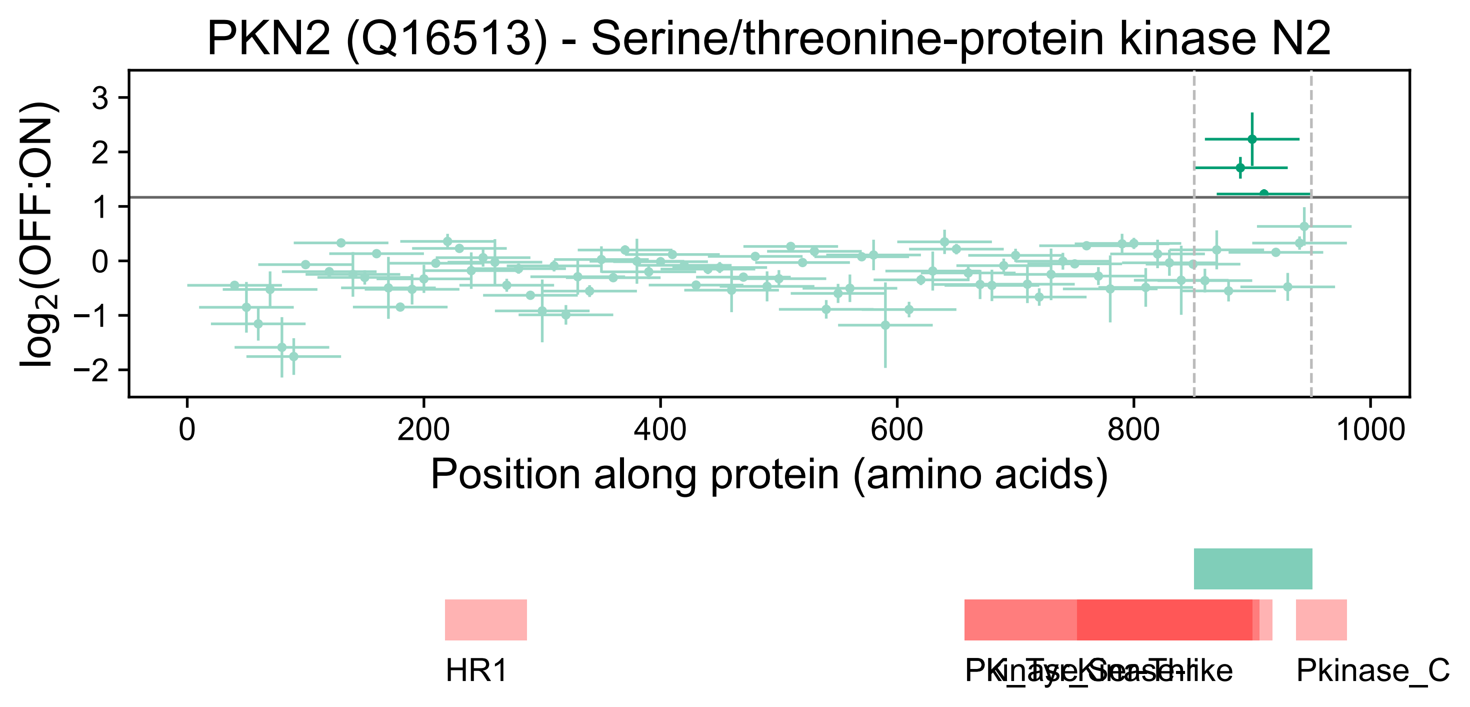
PKN2 (Q16513) - Serine/threonine-protein kinase N2
log2(OFF:ON)
3
2
1
0
−1
−2
0
200
400
600
800
1000
Position along protein (amino acids)
HR1
PK_Tyr_Ser-Thr
Kinase-like
Pkinase_C

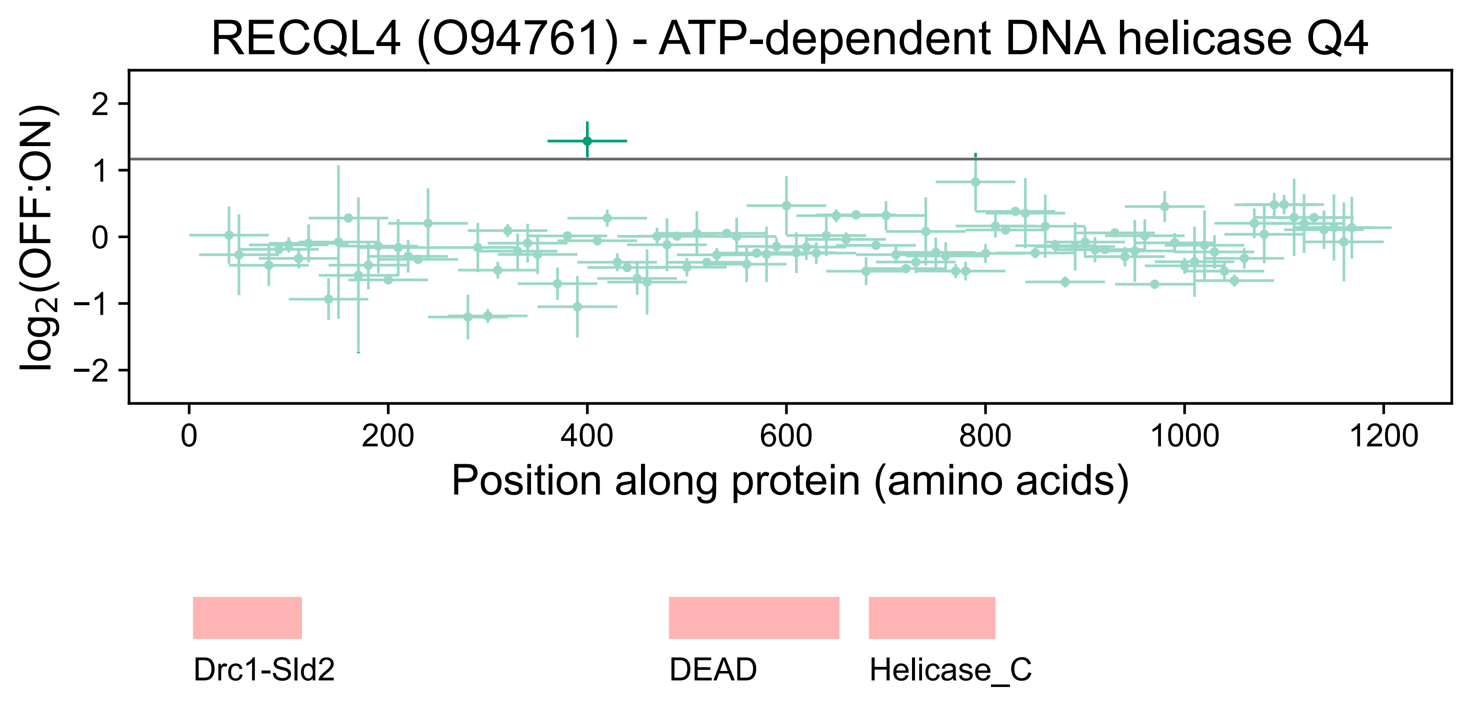
RECQL4 (O94761) - ATP-dependent DNA helicase Q4
log2(OFF:ON)
2
1
0
−1
−2
0
200
400
600
800
1000
1200
Position along protein (amino acids)
Drc1-Sld2
DEAD
Helicase_C

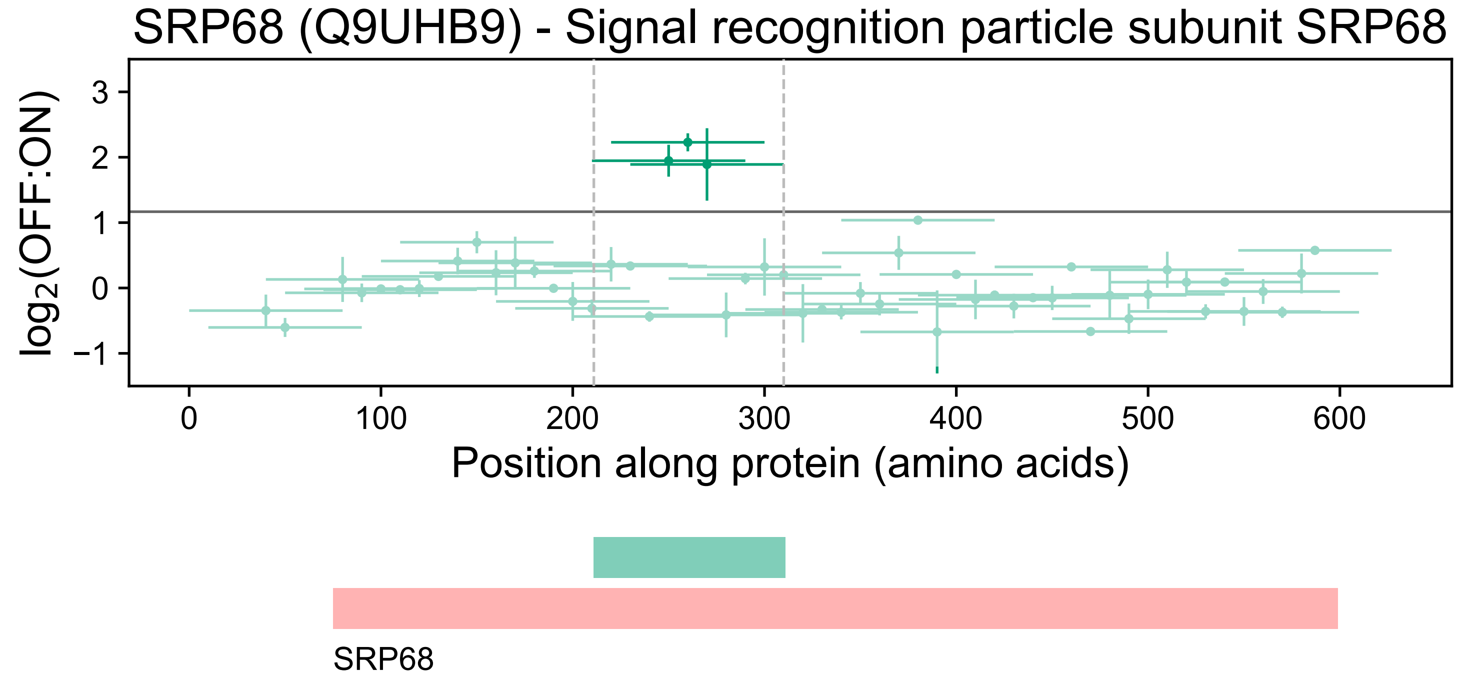

SRP68 (Q9UHB9) - Signal recognition particle subunit SRP68
log2(OFF:ON)
3
2
1
0
−1
0
100
200
300
400
500
600
Position along protein (amino acids)
SRP68

### TDRD6 (O60522) - Tudor domain-containing protein 6

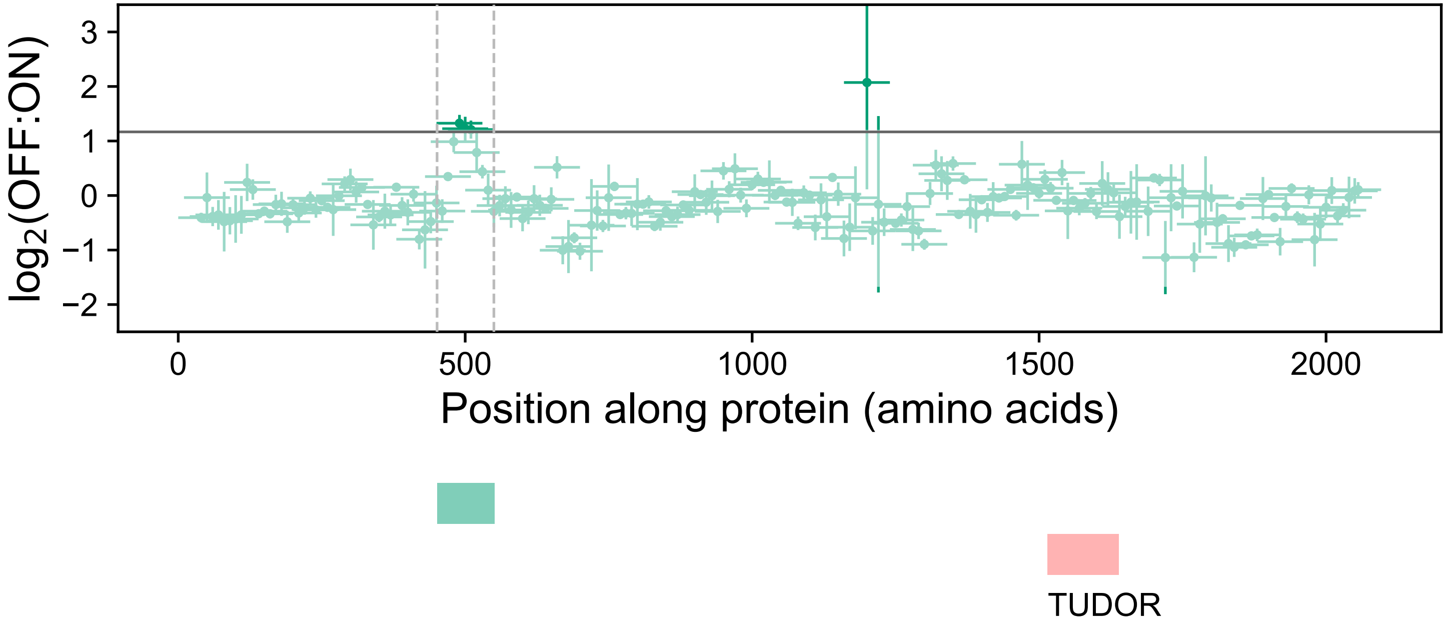

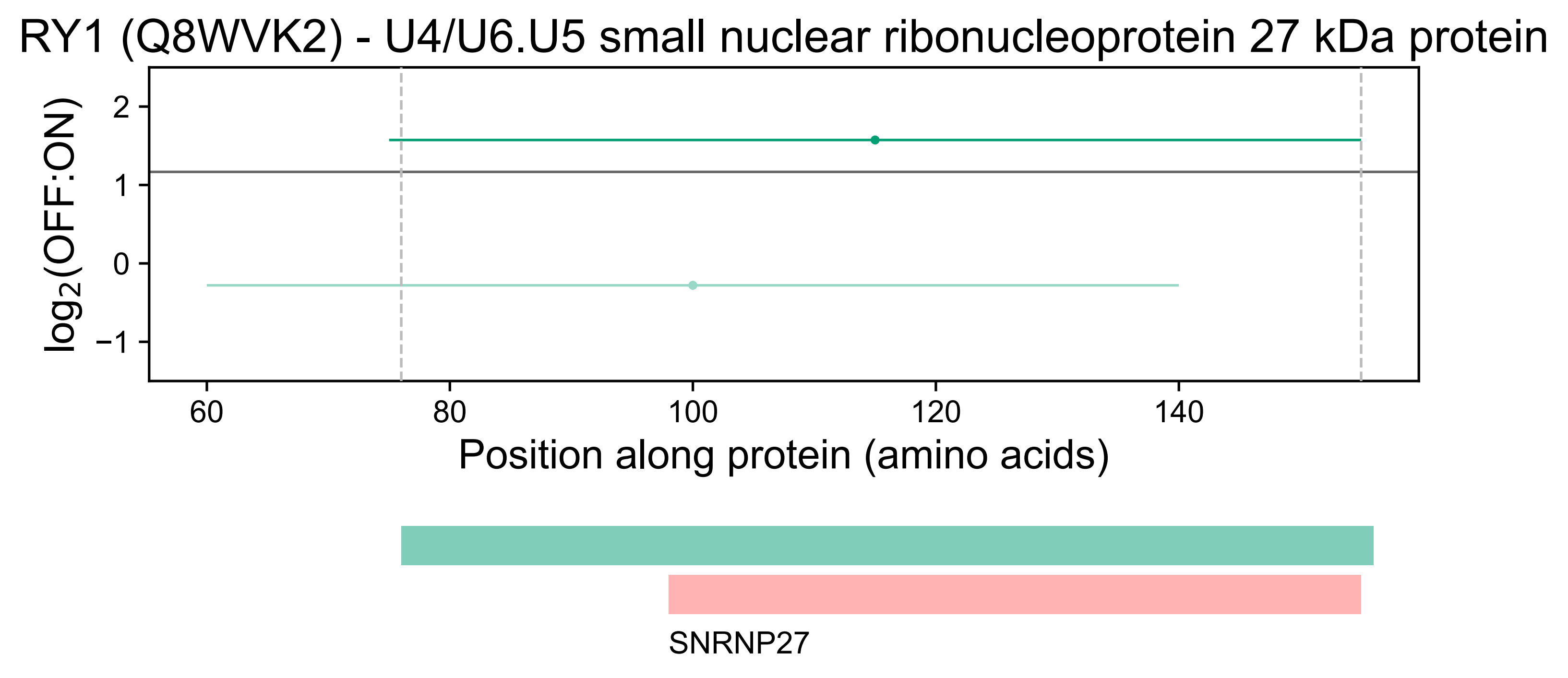
RY1 (Q8WVK2) - U4/U6.U5 small nuclear ribonucleoprotein 27 kDa protein
log2(OFF:ON)
2
1
0
−1
60
80
100
120
140
Position along protein (amino acids)
SNRNP27

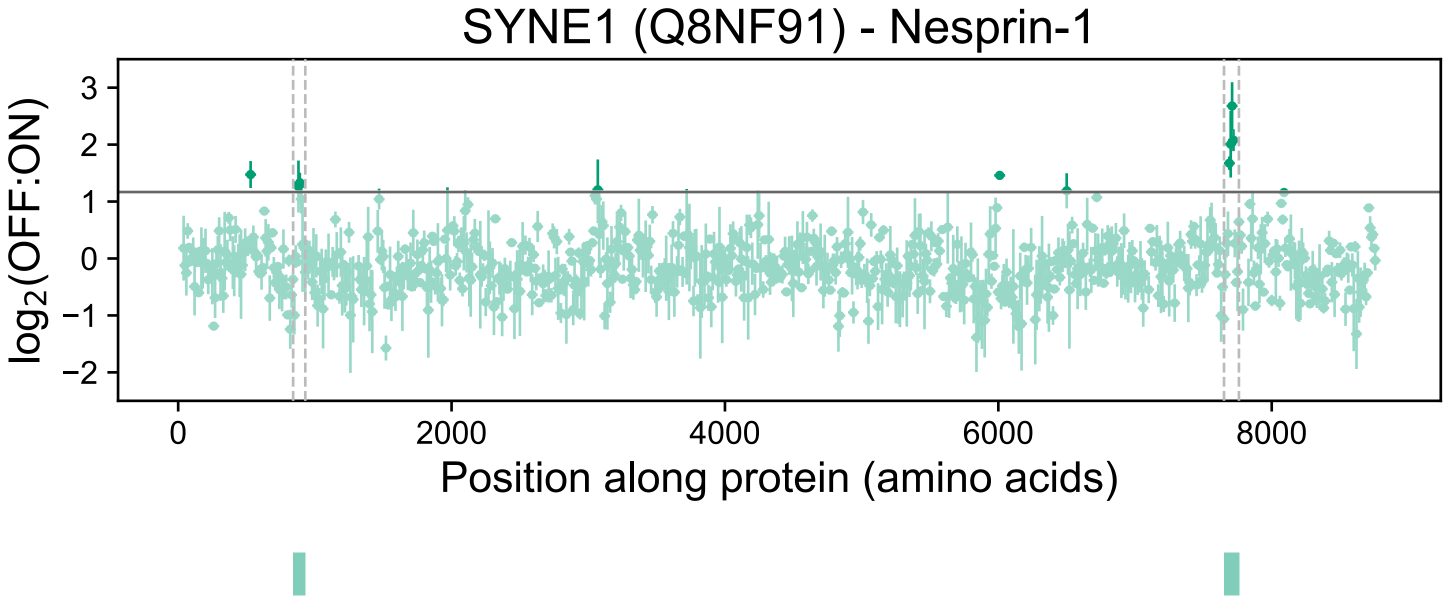

SYNE1 (Q8NF91) - Nesprin-1
log2(OFF:ON)
3
2
1
0
−1
−2
0
2000
4000
6000
8000
Position along protein (amino acids)

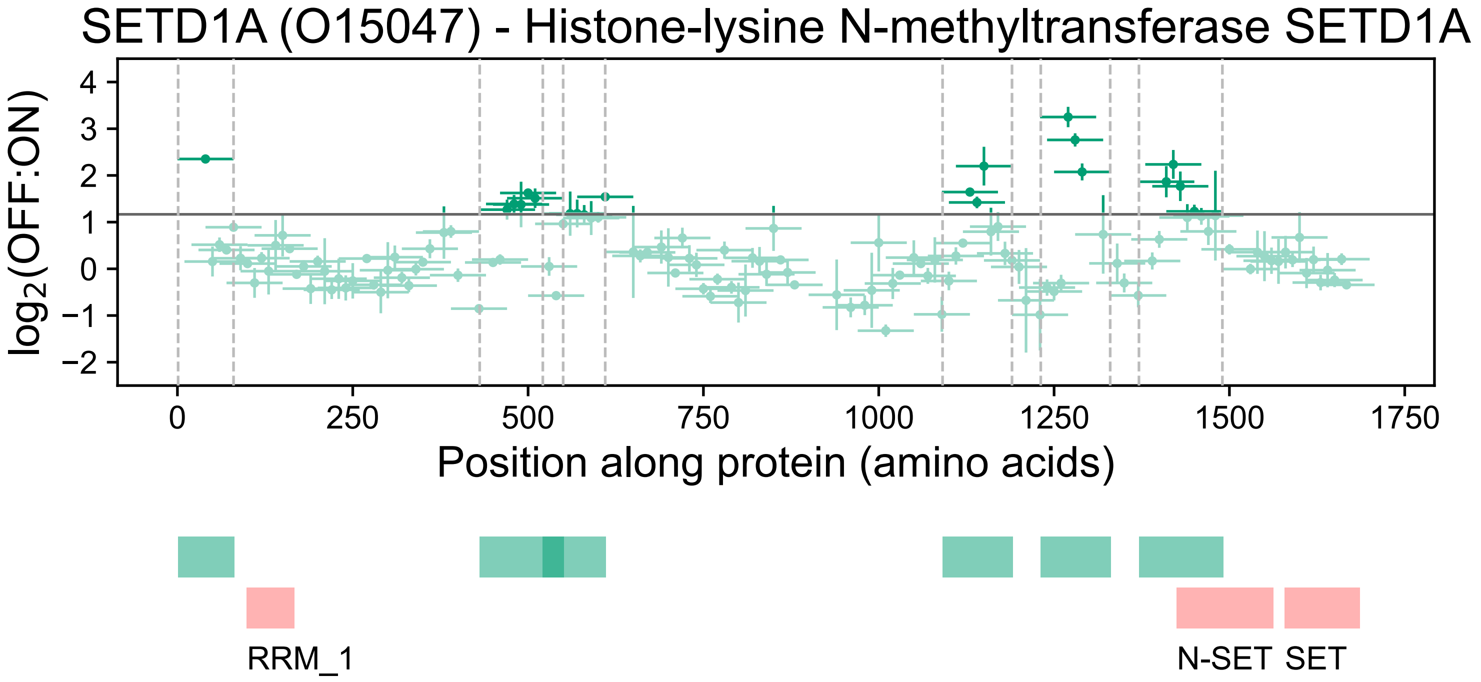

SETD1A (O15047) - Histone-lysine N-methyltransferase SETD1A
log2(OFF:ON)
4
3
2
1
0
−1
−2
0
250
500
750
1000
1250
1500
1750
Position along protein (amino acids)
RRM_1
N-SET
SET

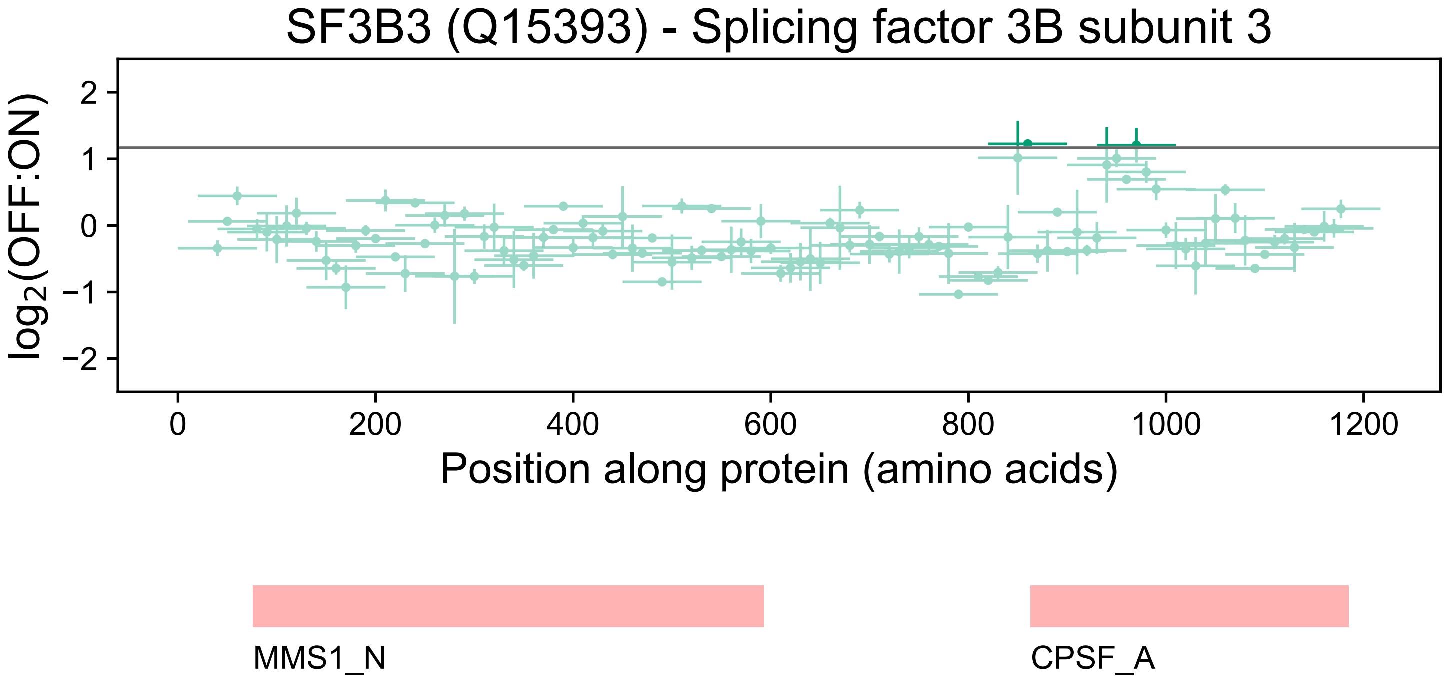
SF3B3 (Q15393) - Splicing factor 3B subunit 3
log2(OFF:ON)
2
1
0
−1
−2
0
200
400
600
800
1000
1200
Position along protein (amino acids)
MMS1_N
CPSF_A

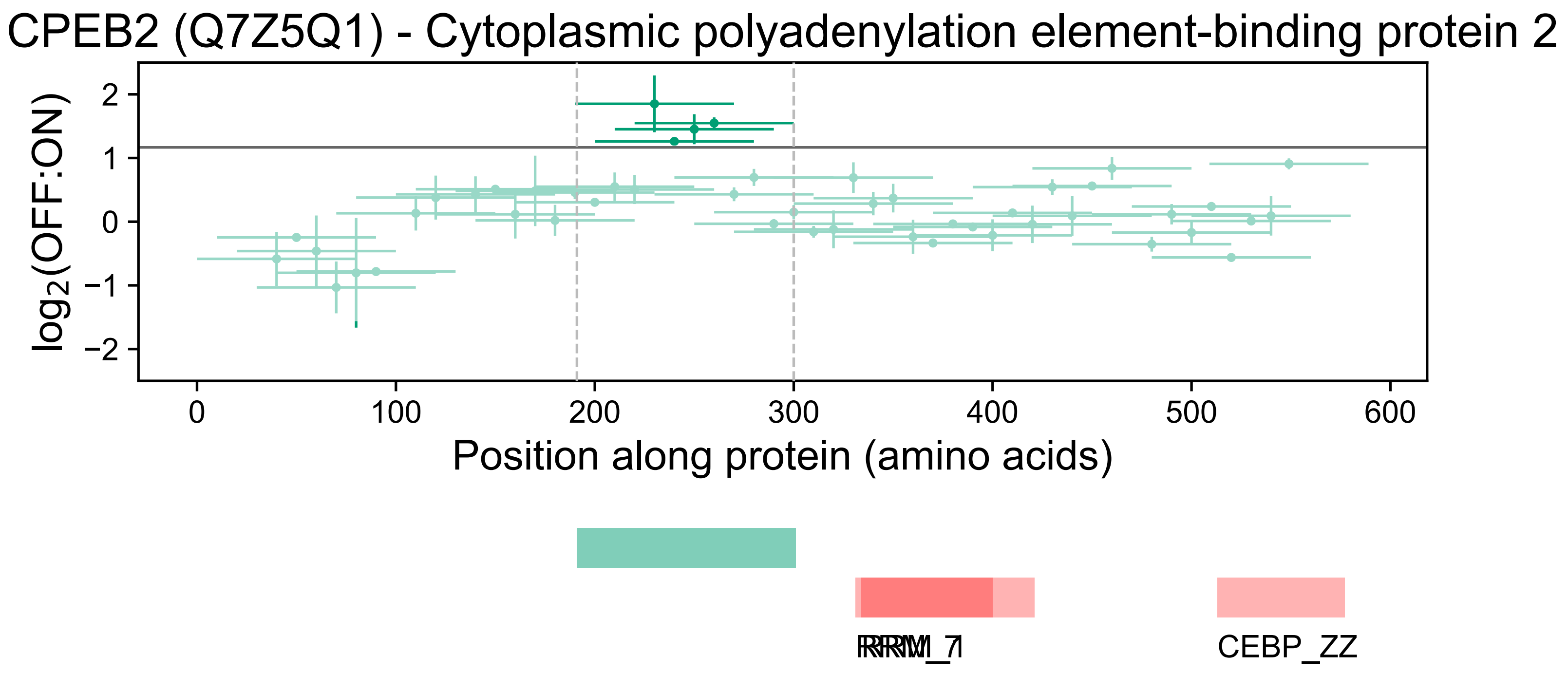
CPEB2 (Q7Z5Q1) - Cytoplasmic polyadenylation element-binding protein 2
log2(OFF:ON)
2
1
0
−1
−2
0
100
200
300
400
500
600
Position along protein (amino acids)
RRM_7
RRM_1
CEBP_ZZ

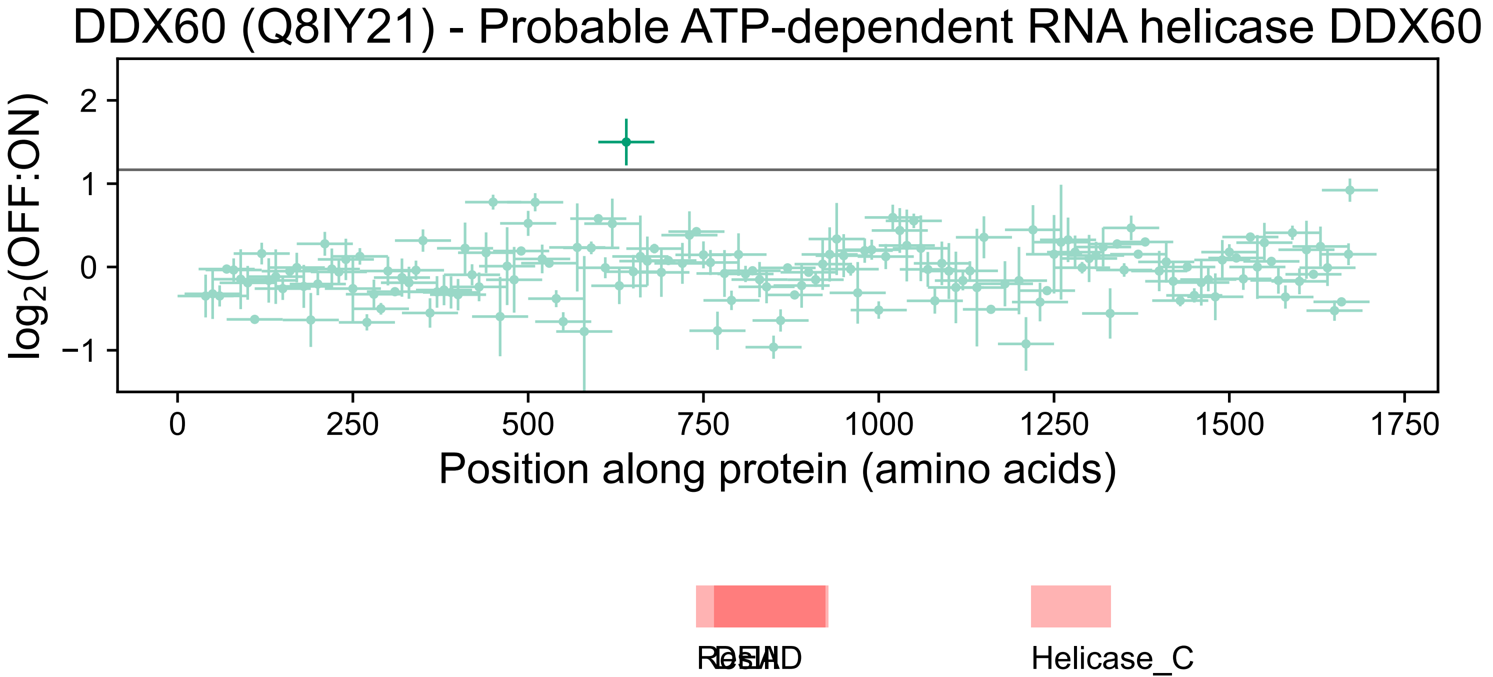

DDX60 (Q8IY21) - Probable ATP-dependent RNA helicase DDX60
log2(OFF:ON)
2
1
0
−1
0
250
500
750
1000
1250
1500
1750
Position along protein (amino acids)
ResIII
DEAD
Helicase_C

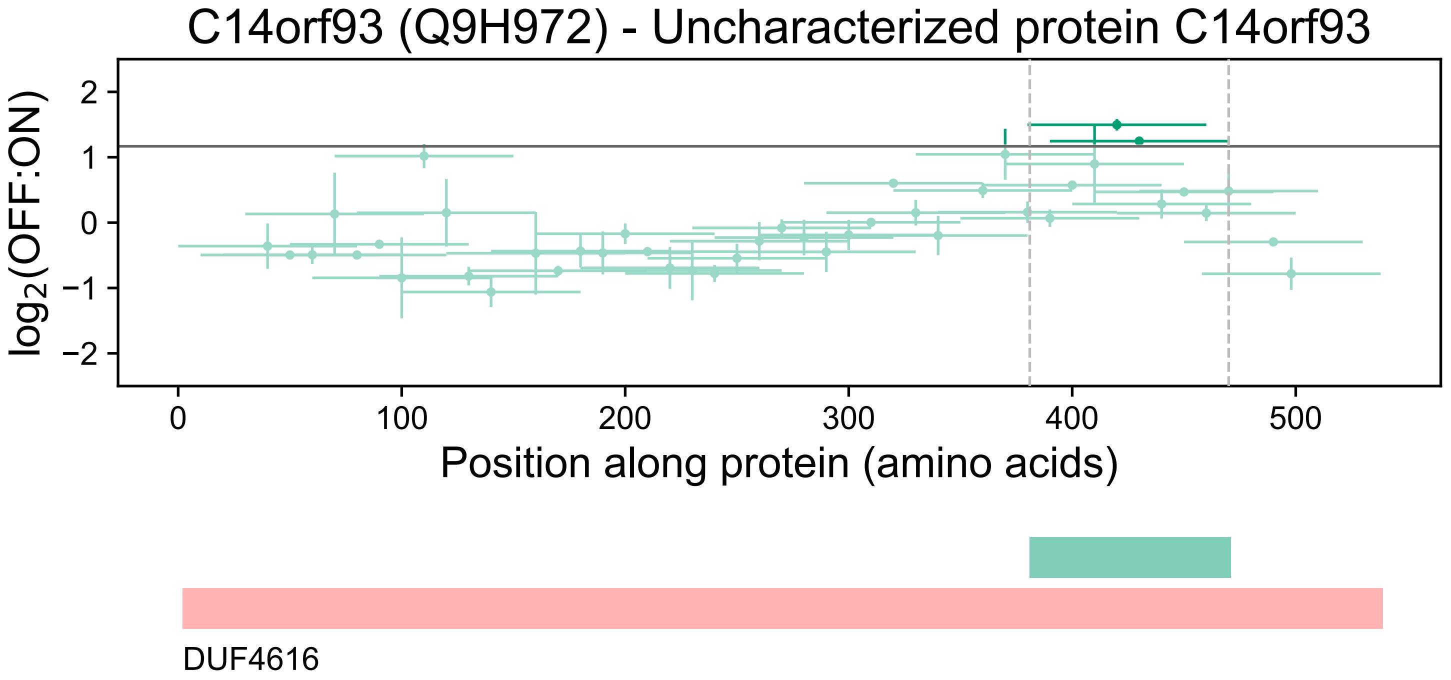
C14orf93 (Q9H972) - Uncharacterized protein C14orf93
log2(OFF:ON)
2
1
0
−1
−2
0
100
200
300
400
500
Position along protein (amino acids)
DUF4616

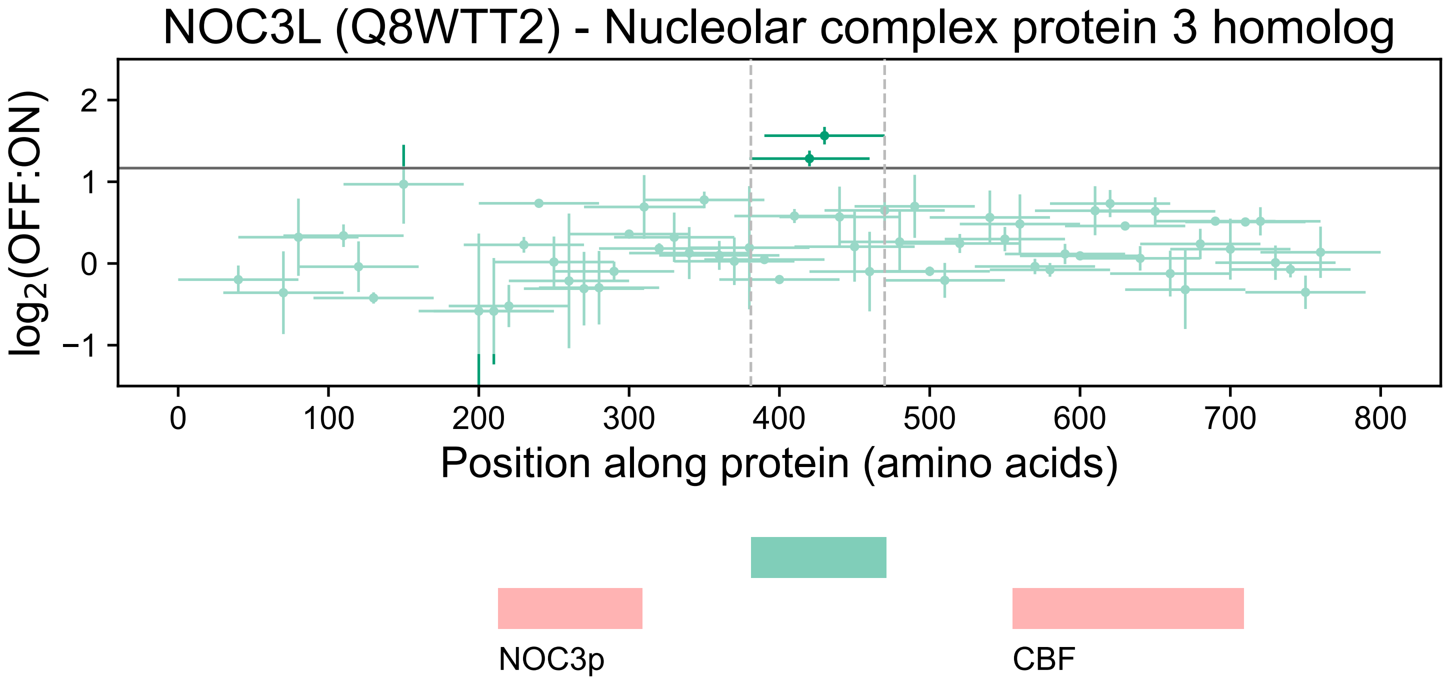

NOC3L (Q8WTT2) - Nucleolar complex protein 3 homolog
$\log_2$(OFF:ON)
2
1
0
−1
0
100
200
300
400
500
600
700
800
Position along protein (amino acids)
NOC3p
CBF

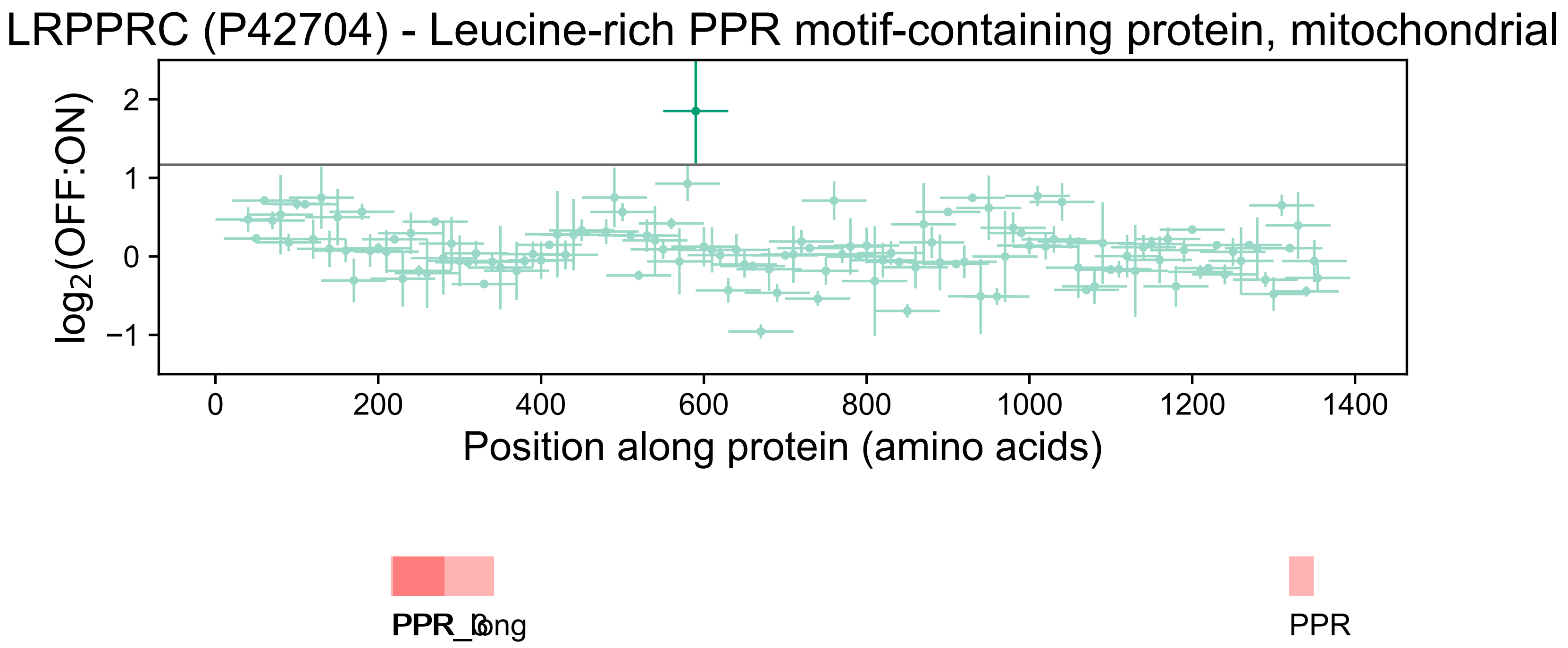

LRPPRC (P42704) - Leucine-rich PPR motif-containing protein, mitochondrial
$\log_2$(OFF:ON)
2
1
0
−1
0
200
400
600
800
1000
1200
1400
Position along protein (amino acids)
PPR_3
PPR_long
PPR

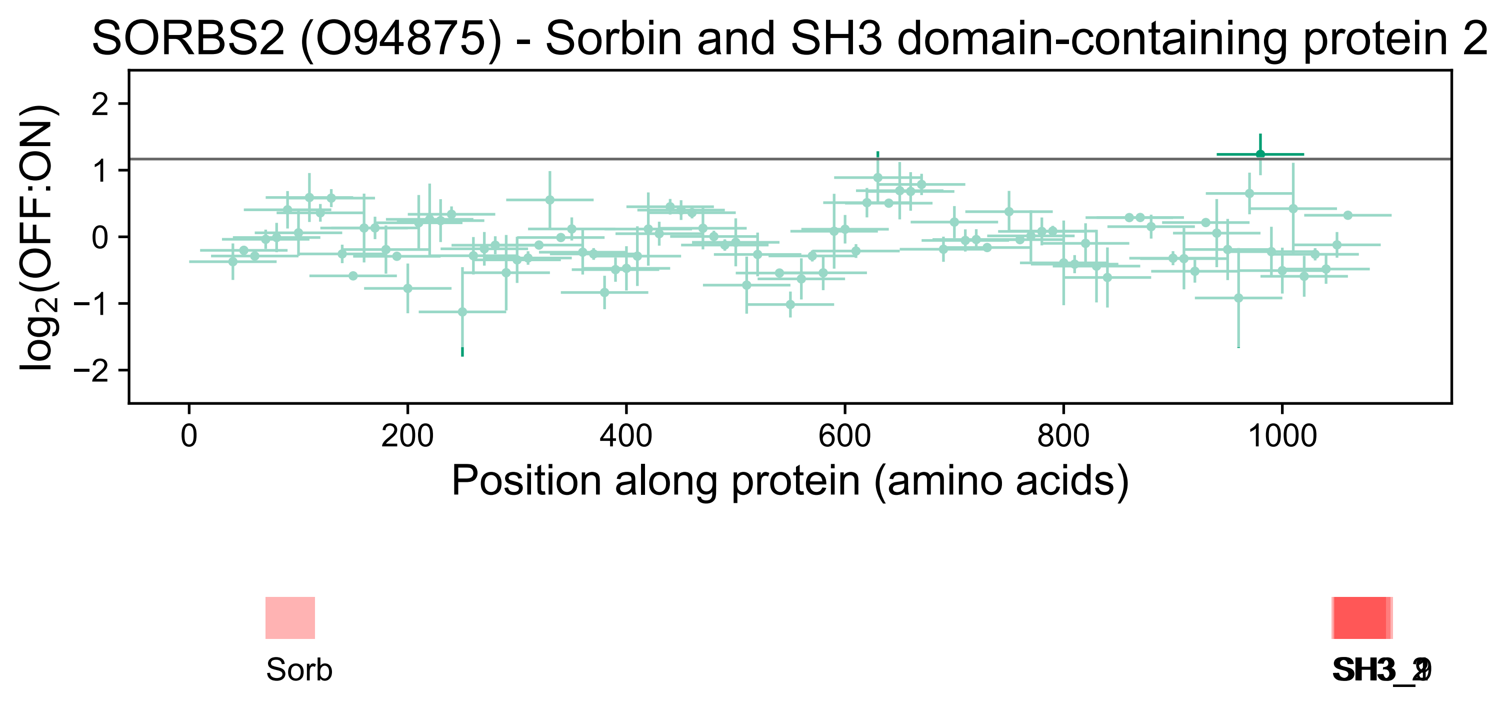
SORBS2 (O94875) - Sorbin and SH3 domain-containing protein 2
log2(OFF:ON)
2
1
0
−1
−2
0
200
400
600
800
1000
Position along protein (amino acids)
Sorb
SH3_2
SH3_9

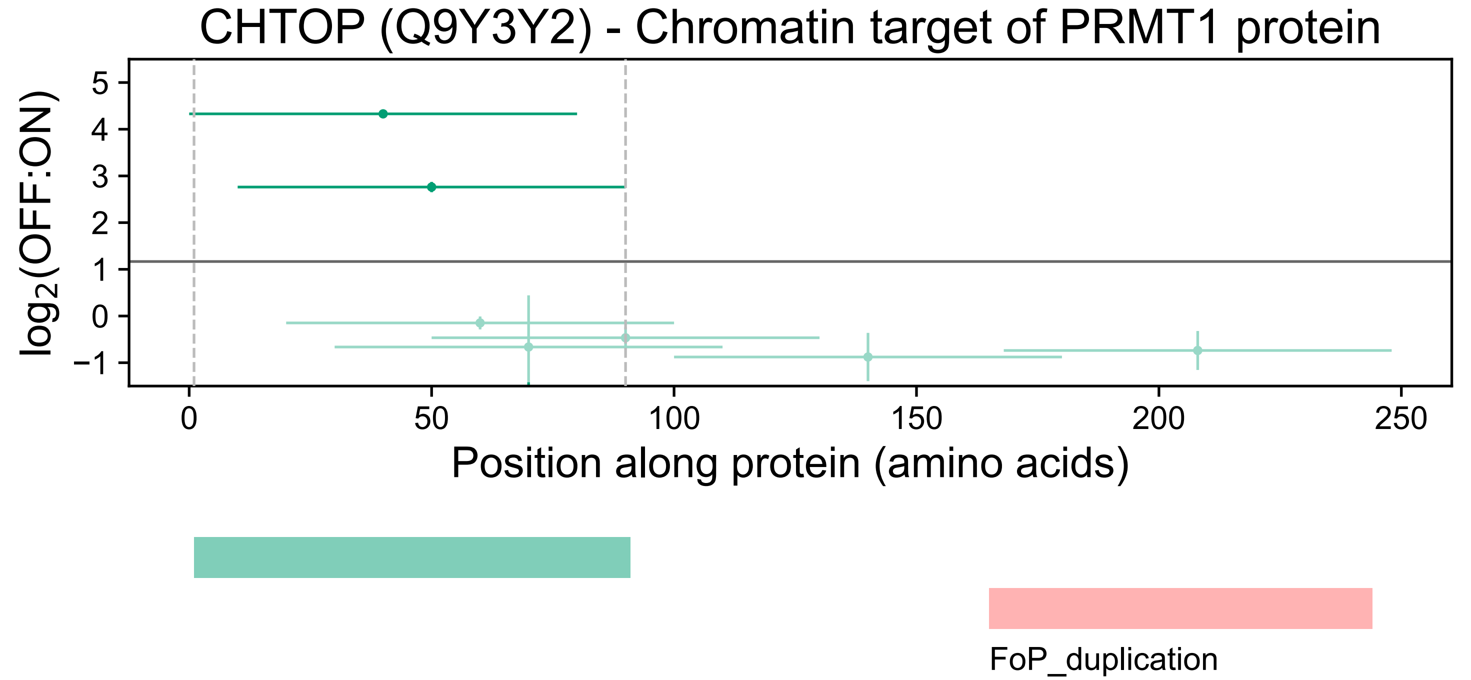
CHTOP (Q9Y3Y2) - Chromatin target of PRMT1 protein
log2(OFF:ON)
5
4
3
2
1
0
−1
0
50
100
150
200
250
Position along protein (amino acids)
FoP_duplication

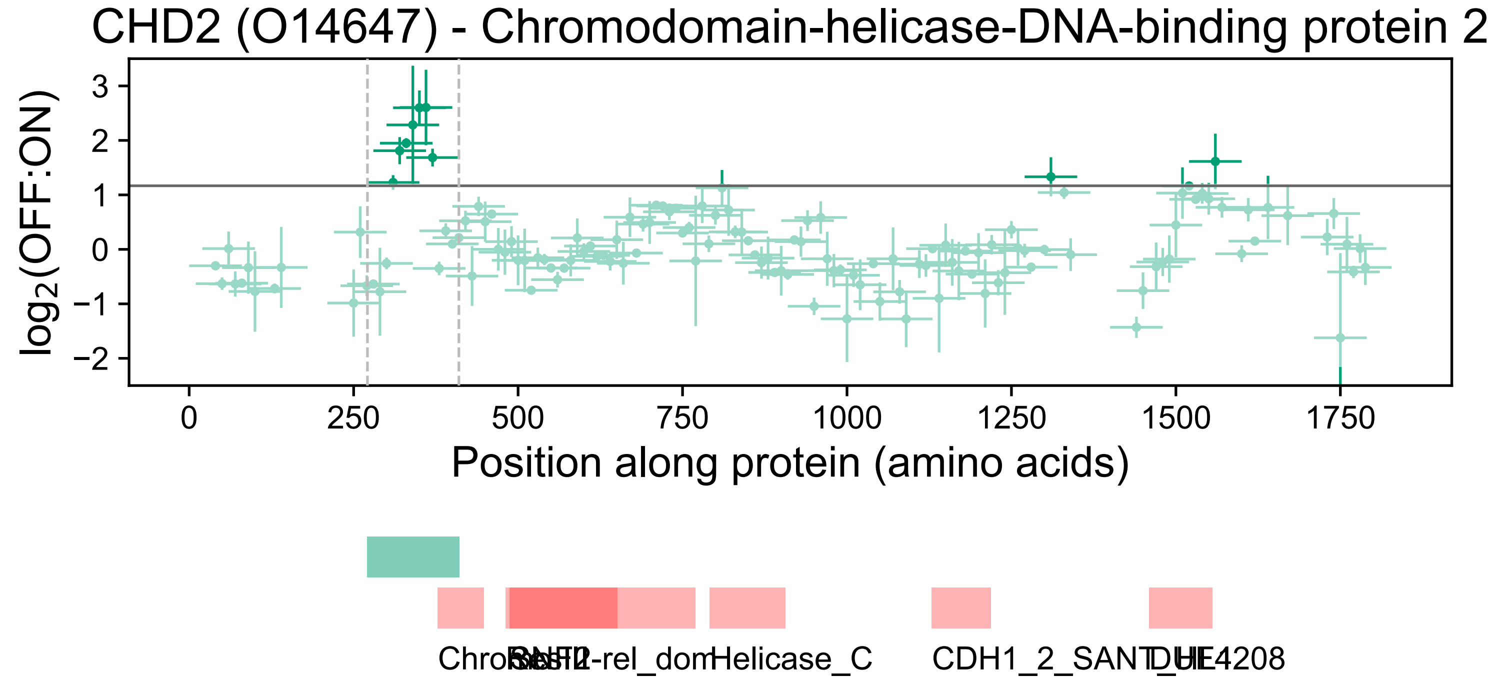

CHD2 (O14647) - Chromodomain-helicase-DNA-binding protein 2
log2(OFF:ON)
3
2
1
0
−1
−2
0
250
500
750
1000
1250
1500
1750
Position along protein (amino acids)
Helicase_C

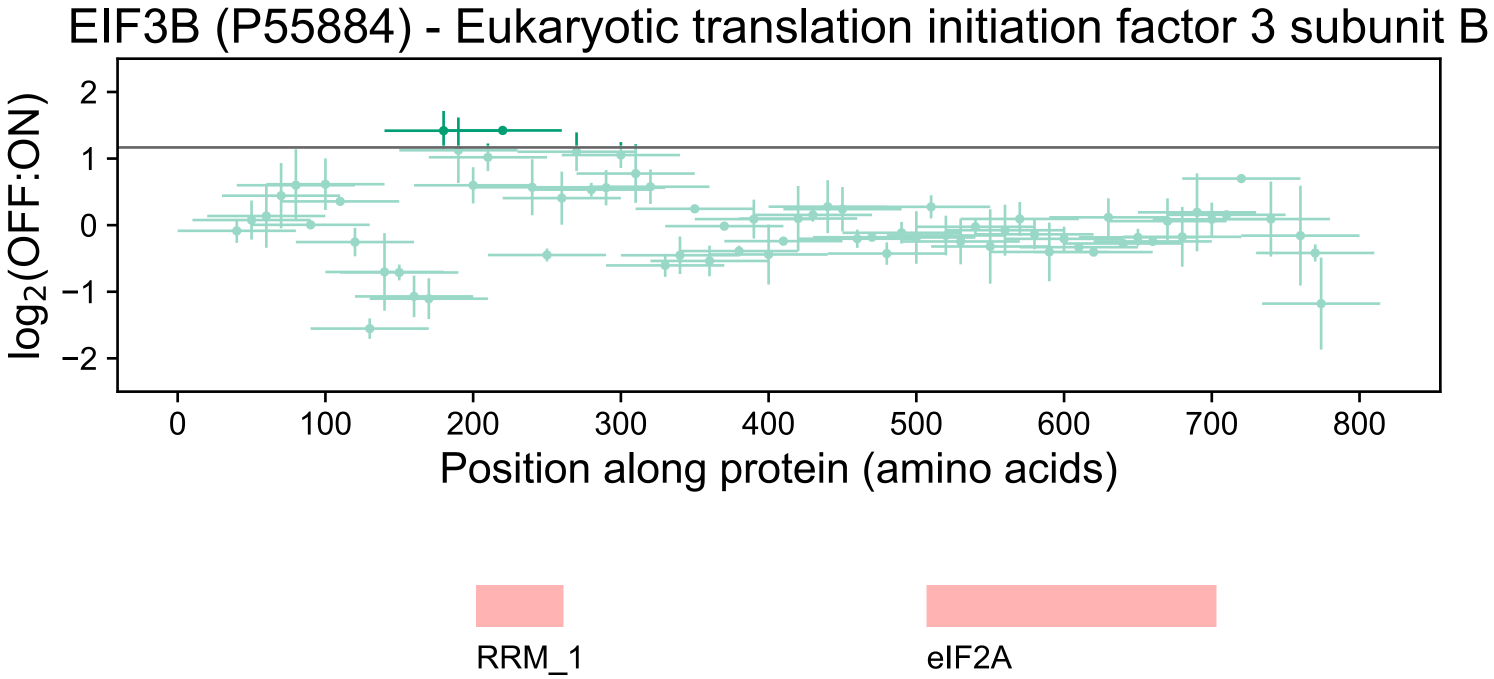

EIF3B (P55884) - Eukaryotic translation initiation factor 3 subunit B
log2(OFF:ON)
2
1
0
−1
−2
0
100
200
300
400
500
600
700
800
Position along protein (amino acids)
RRM_1
eIF2A

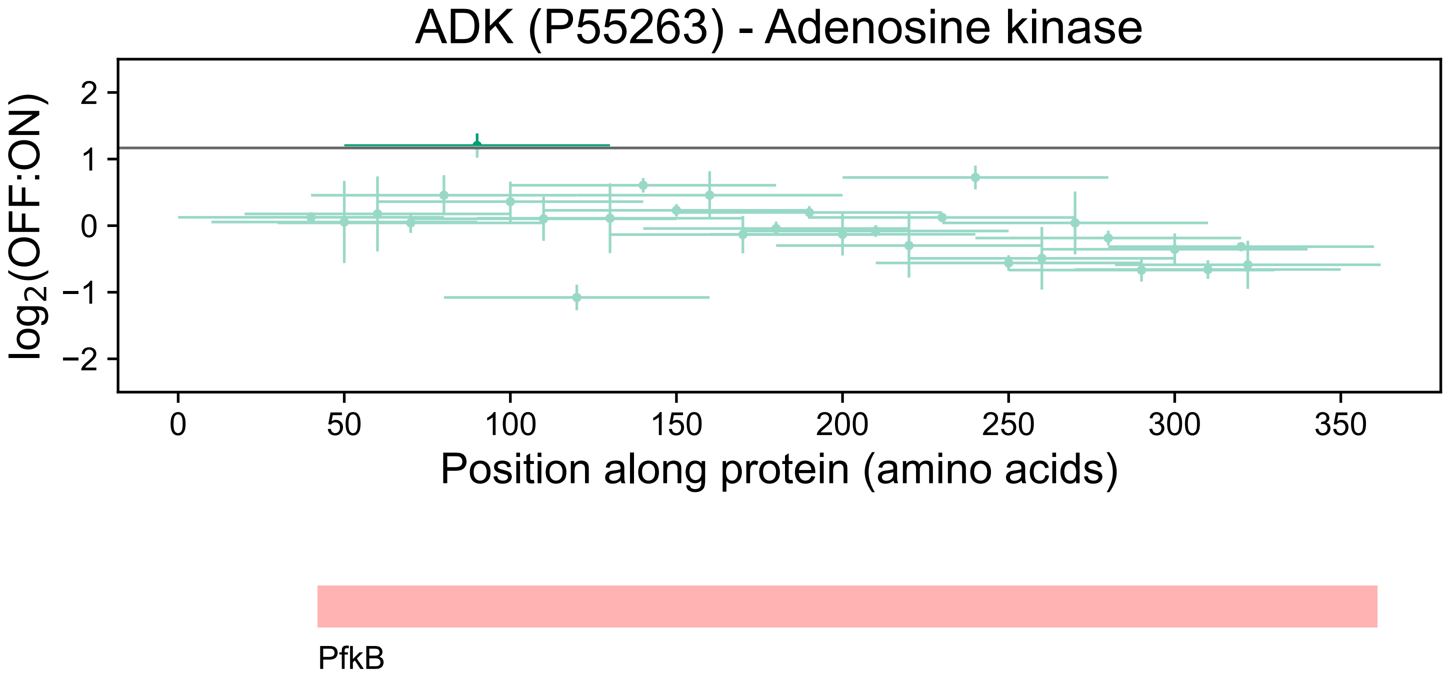

ADK (P55263) - Adenosine kinase
log2(OFF:ON)
2
1
0
−1
−2
0
50
100
150
200
250
300
350
Position along protein (amino acids)
PfkB

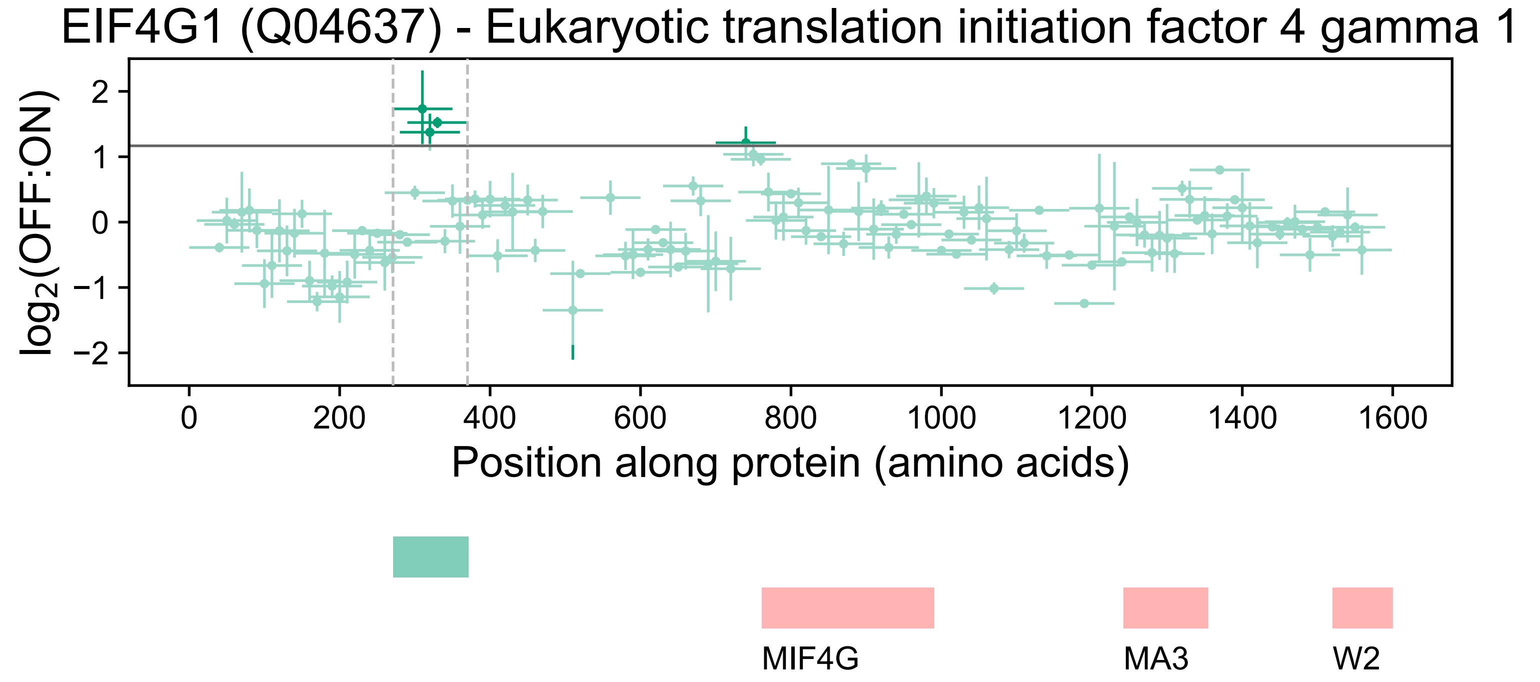

EIF4G1 (Q04637) - Eukaryotic translation initiation factor 4 gamma 1
log2(OFF:ON)
2
1
0
−1
−2
0
200
400
600
800
1000
1200
1400
1600
Position along protein (amino acids)
MIF4G
MA3
W2

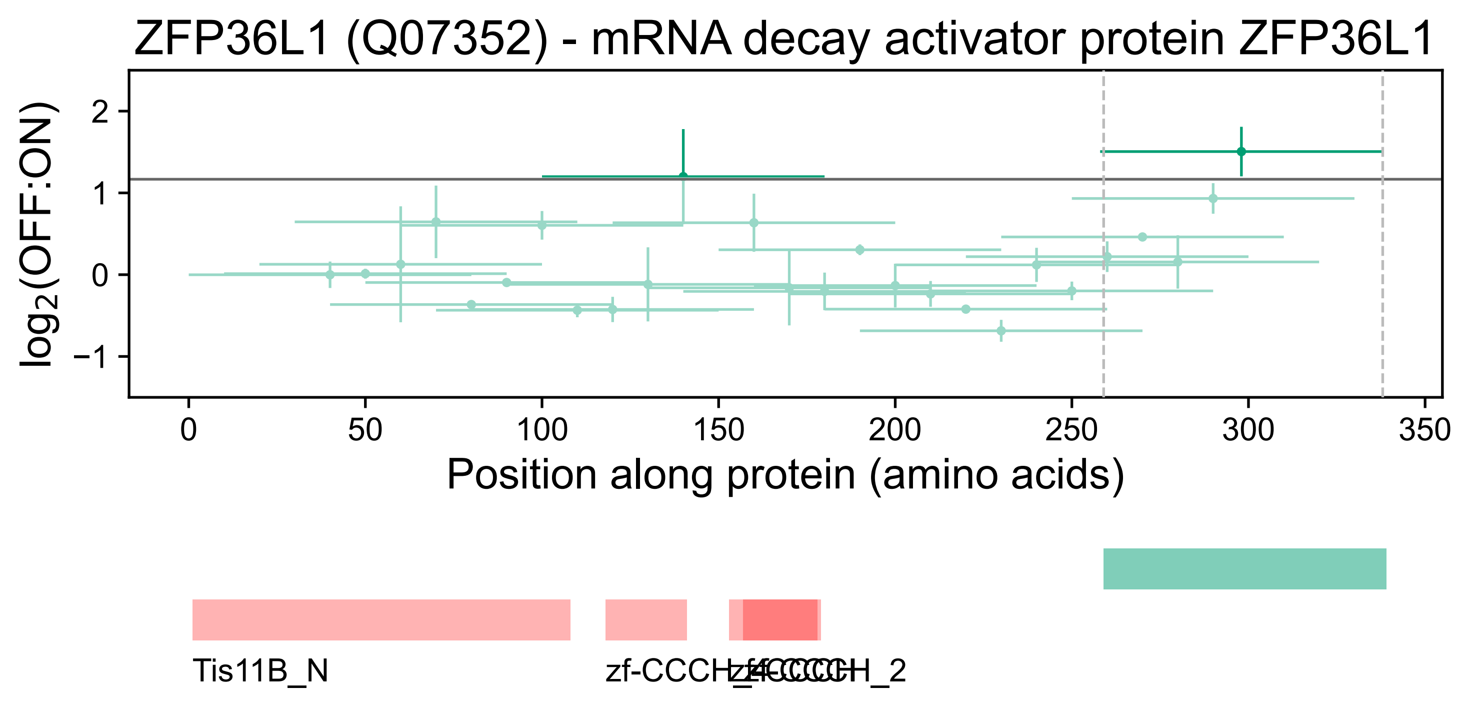

ZFP36L1 (Q07352) - mRNA decay activator protein ZFP36L1
log2(OFF:ON)
2
1
0
−1
0
50
100
150
200
250
300
350
Position along protein (amino acids)
Tis11B_N
zf-CCCH_4
zf-CCCH_2

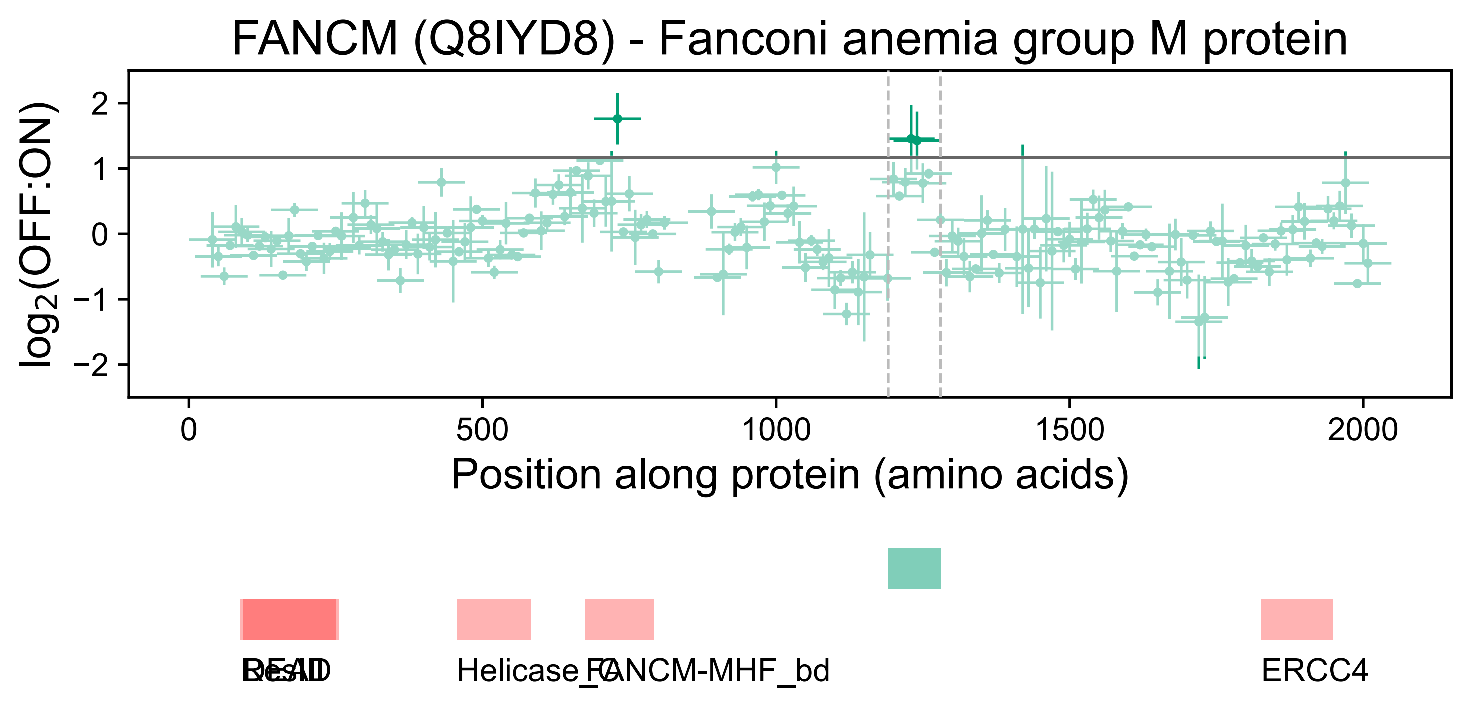
FANCM (Q8IYD8) - Fanconi anemia group M protein
log2(OFF:ON)
2
1
0
−1
−2
0
500
1000
1500
2000
Position along protein (amino acids)
DEAD
ResIII
Helicase_C
FANCM-MHF_bd
ERCC4

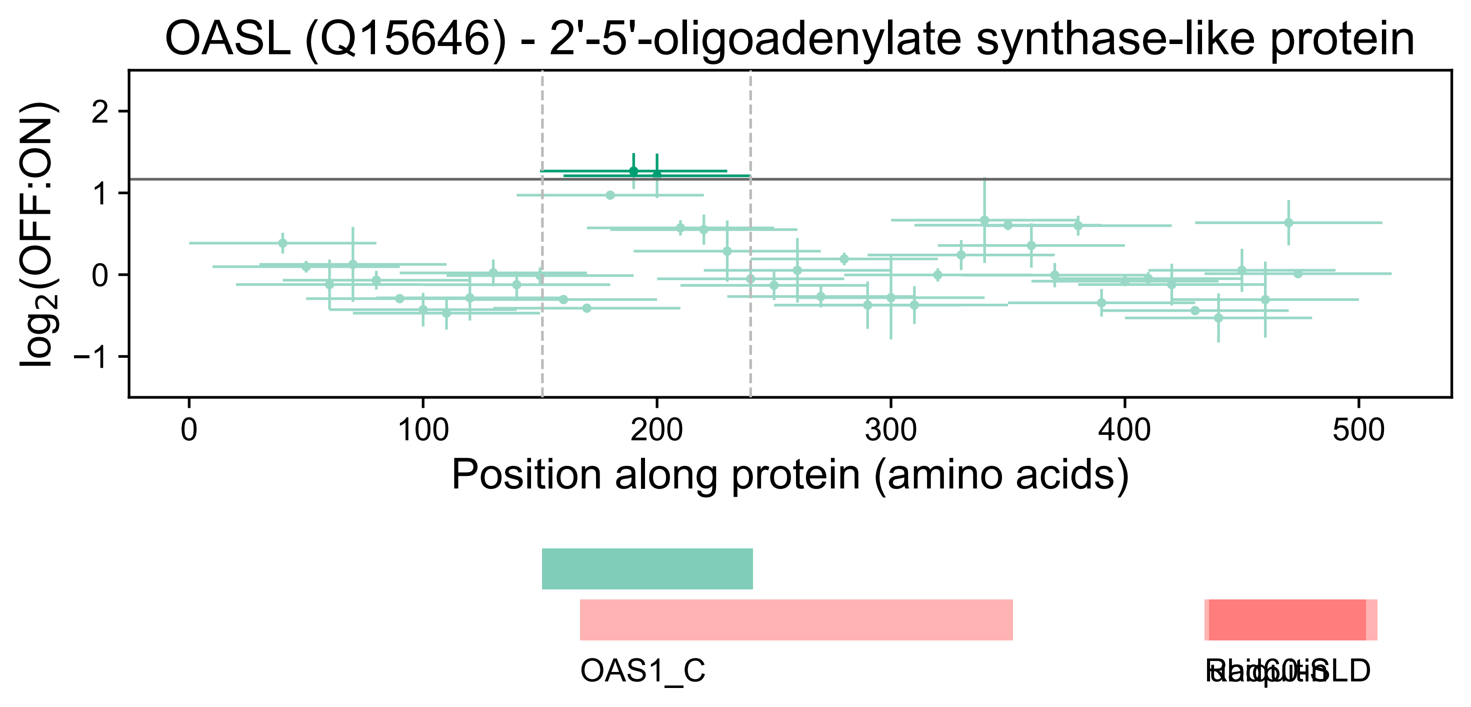
OASL (Q15646) - 2'-5'-oligoadenylate synthase-like protein
log2(OFF:ON)
2
1
0
−1
0
100
200
300
400
500
Position along protein (amino acids)
OAS1_C

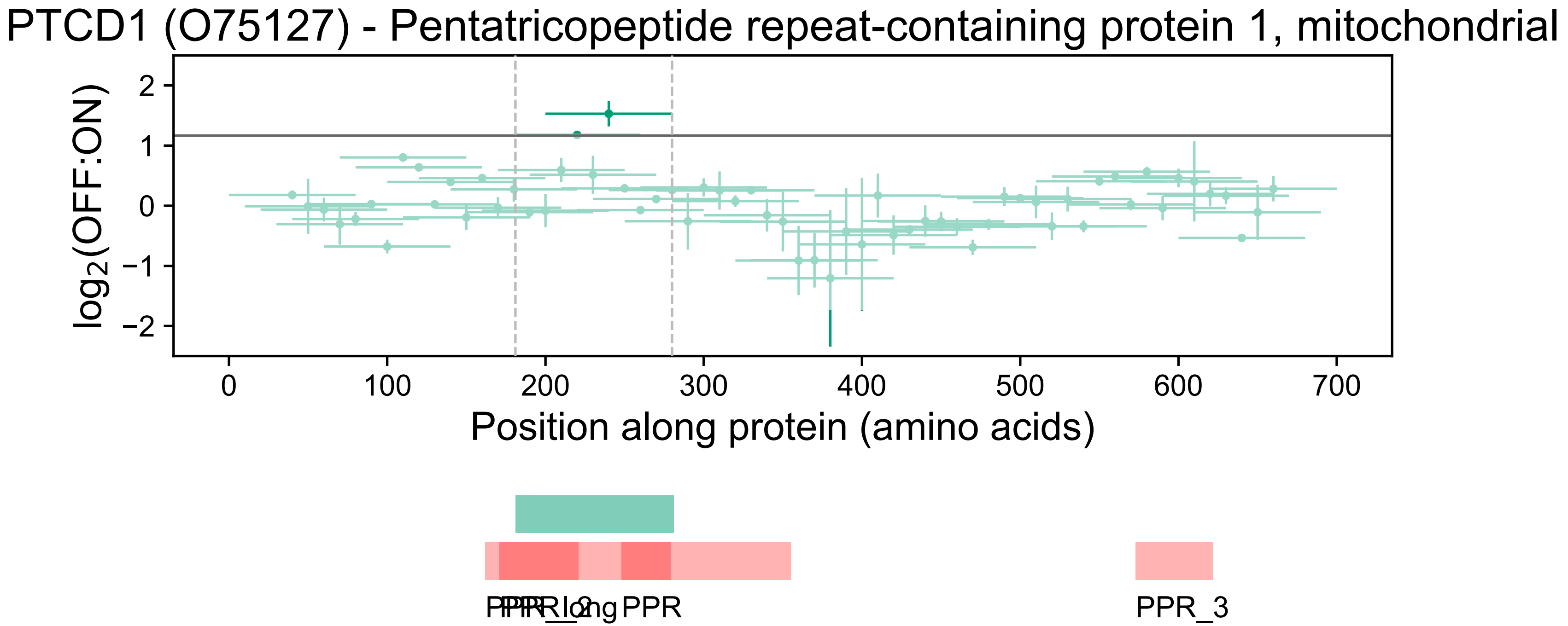

PTCD1 (O75127) - Pentatricopeptide repeat-containing protein 1, mitochondrial
log2(OFF:ON)
2
1
0
−1
−2
0
100
200
300
400
500
600
700
Position along protein (amino acids)
PPR_3

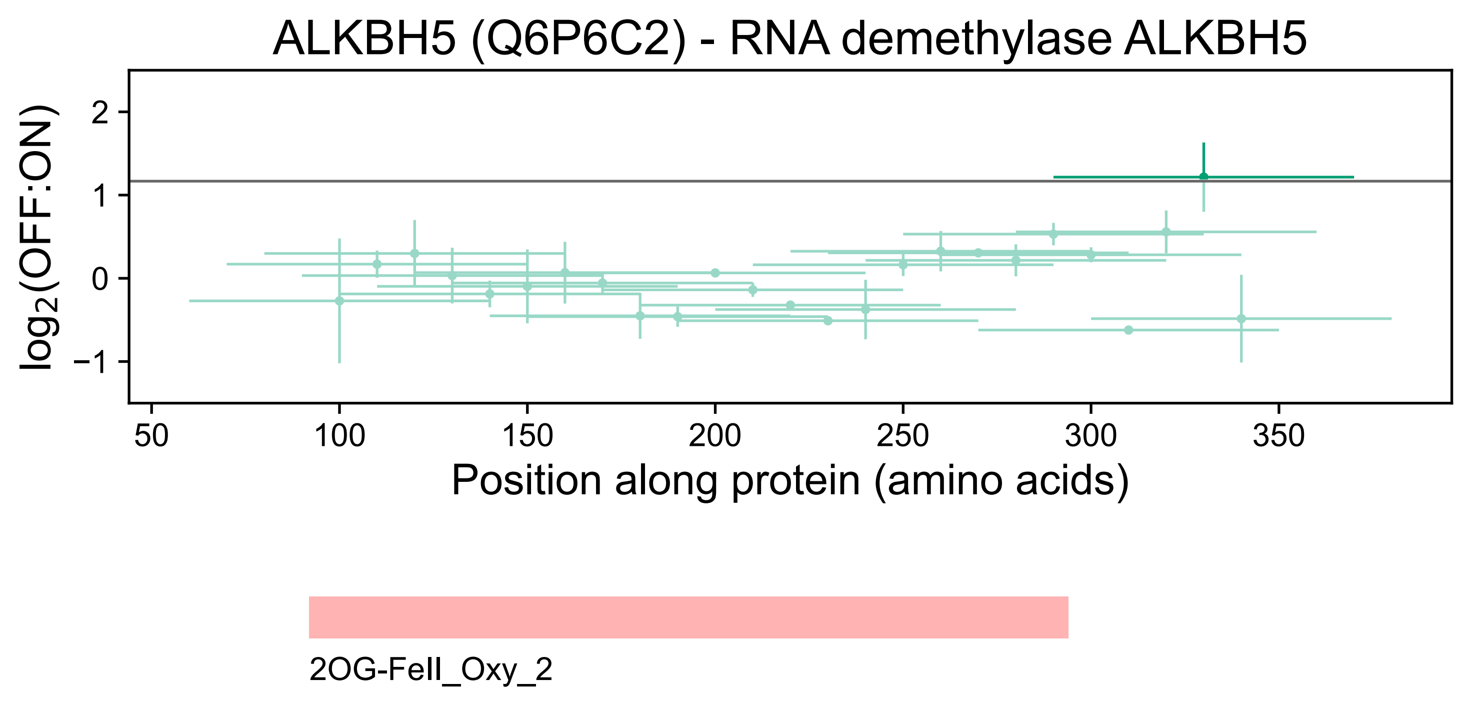

ALKBH5 (Q6P6C2) - RNA demethylase ALKBH5
log2(OFF:ON)
2
1
0
−1
50
100
150
200
250
300
350
Position along protein (amino acids)
2OG-FeII_Oxy_2

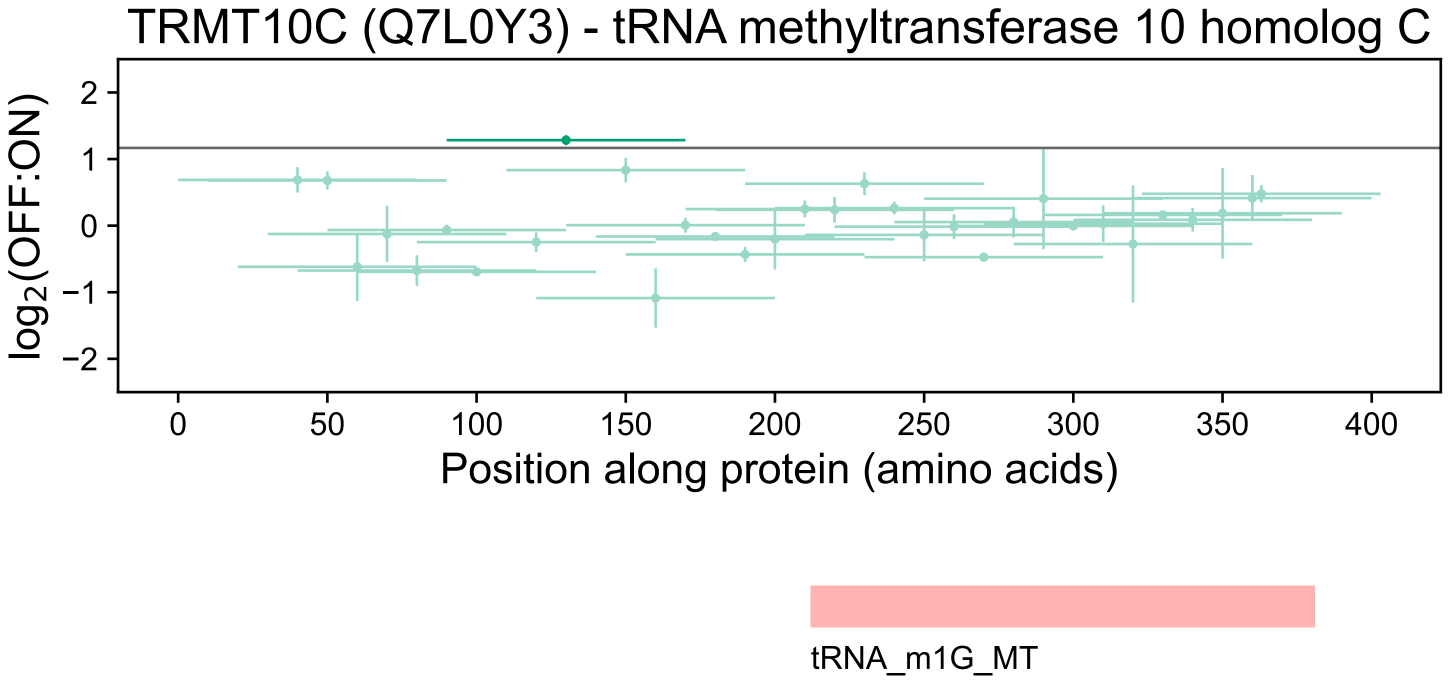
TRMT10C (Q7L0Y3) - tRNA methyltransferase 10 homolog C
log2(OFF:ON)
2
1
0
−1
−2
0
50
100
150
200
250
300
350
400
Position along protein (amino acids)
tRNA_m1G_MT

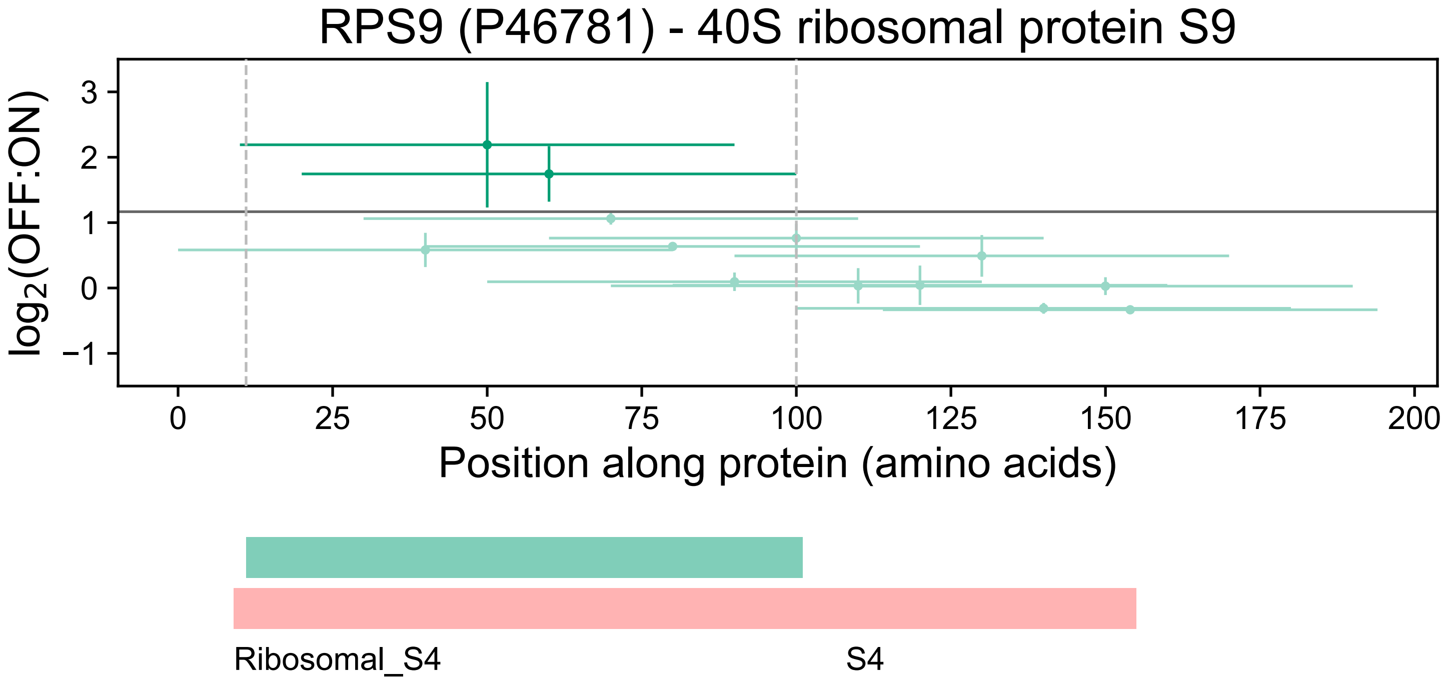
RPS9 (P46781) - 40S ribosomal protein S9
log2(OFF:ON)
3
2
1
0
−1
0
25
50
75
100
125
150
175
200
Position along protein (amino acids)
Ribosomal_S4
S4

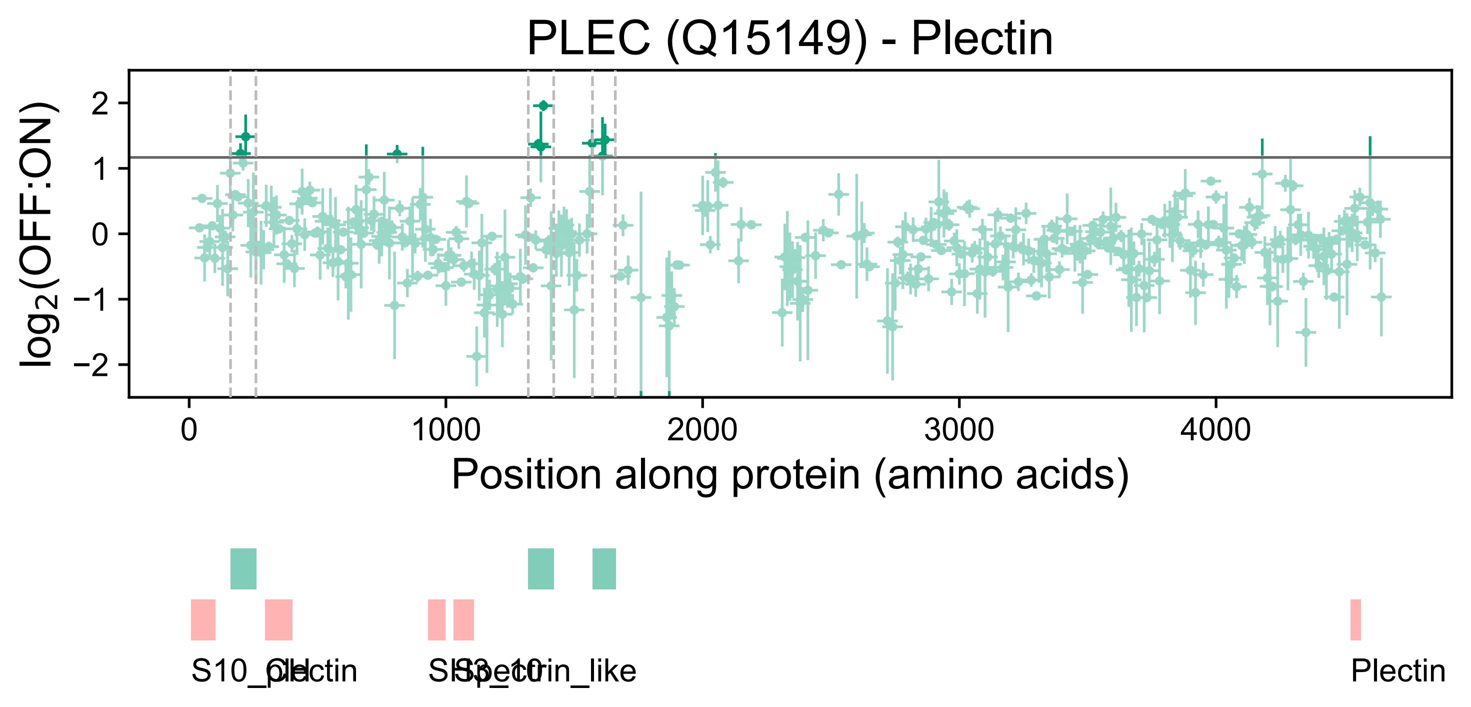
PLEC (Q15149) - Plectin
log2(OFF:ON)
2
1
0
−1
−2
0
1000
2000
3000
4000
Position along protein (amino acids)
S10_plectin
Ch
SH3_10
Spectrin_like
Plectin

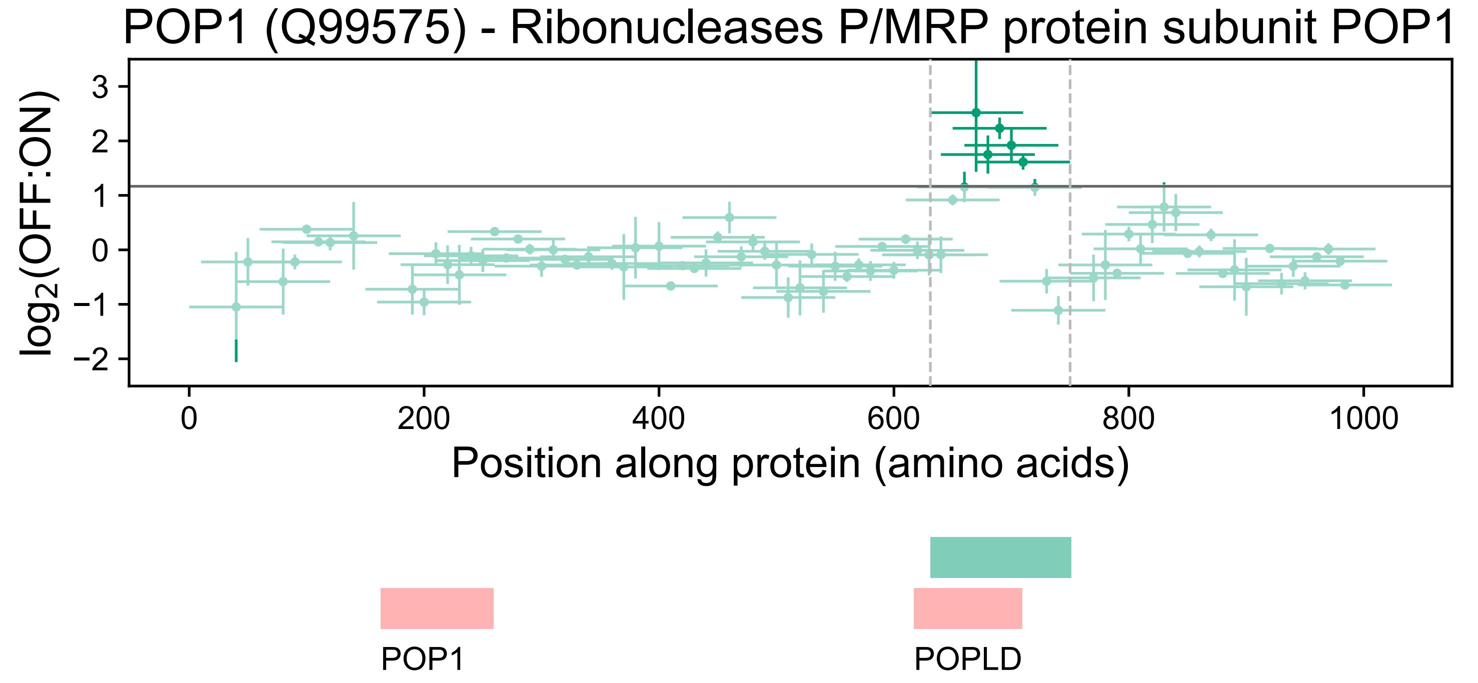

POP1 (Q99575) - Ribonucleases P/MRP protein subunit POP1
log2(OFF:ON)
3
2
1
0
−1
−2
0
200
400
600
800
1000
Position along protein (amino acids)
POP1
POPLD

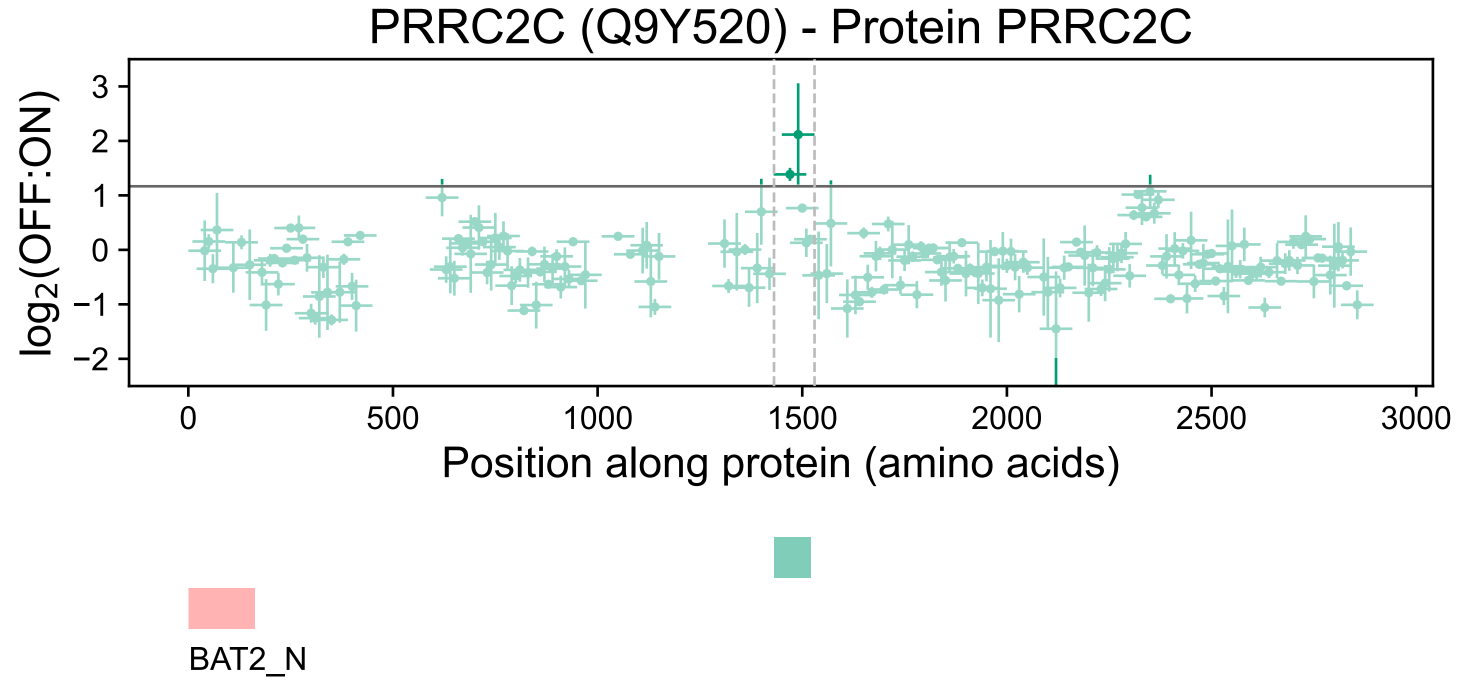

PRRC2C (Q9Y520) - Protein PRRC2C
log2(OFF:ON)
3
2
1
0
−1
−2
0
500
1000
1500
2000
2500
3000
Position along protein (amino acids)
BAT2_N

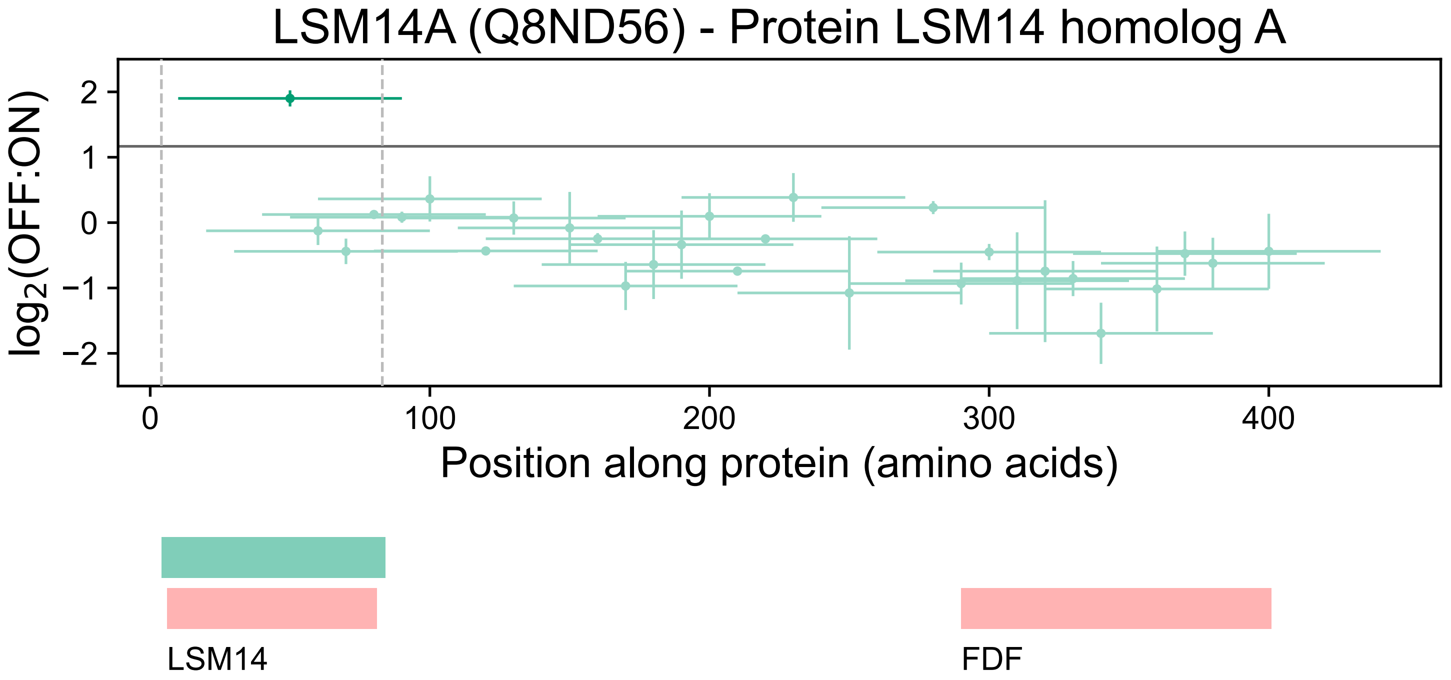

LSM14A (Q8ND56) - Protein LSM14 homolog A
log2(OFF:ON)
2
1
0
−1
−2
0
100
200
300
400
Position along protein (amino acids)
LSM14
FDF

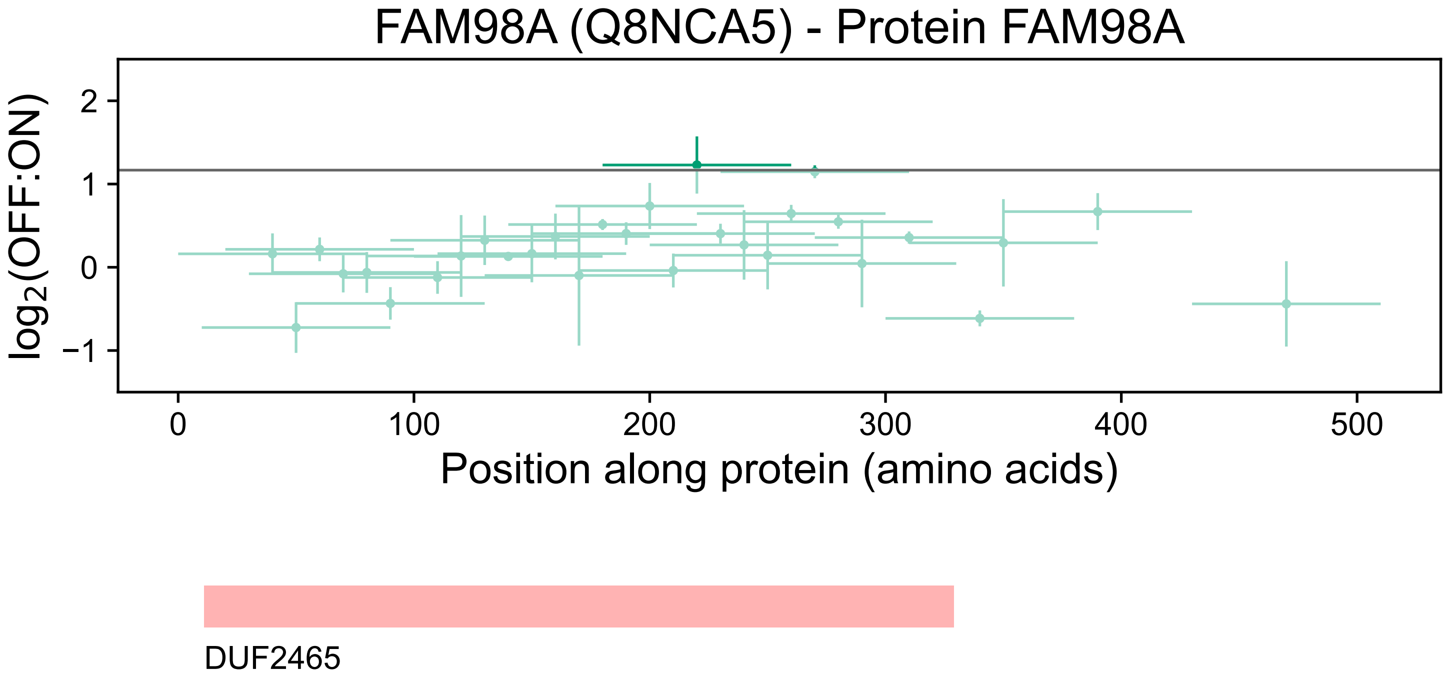
FAM98A (Q8NCA5) - Protein FAM98A
log2(OFF:ON)
2
1
0
−1
0
100
200
300
400
500
Position along protein (amino acids)
DUF2465

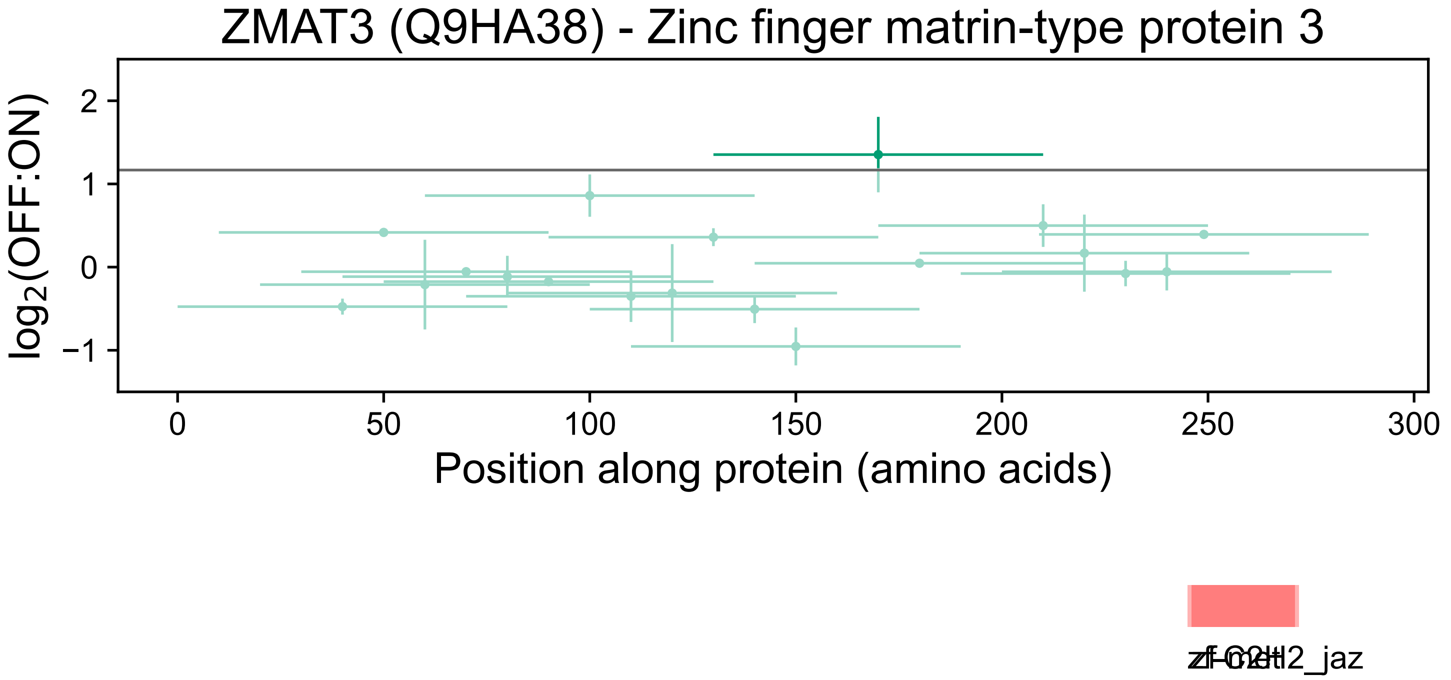
ZMAT3 (Q9HA38) - Zinc finger matrin-type protein 3
log2(OFF:ON)
2
1
0
−1
0
50
100
150
200
250
300
Position along protein (amino acids)

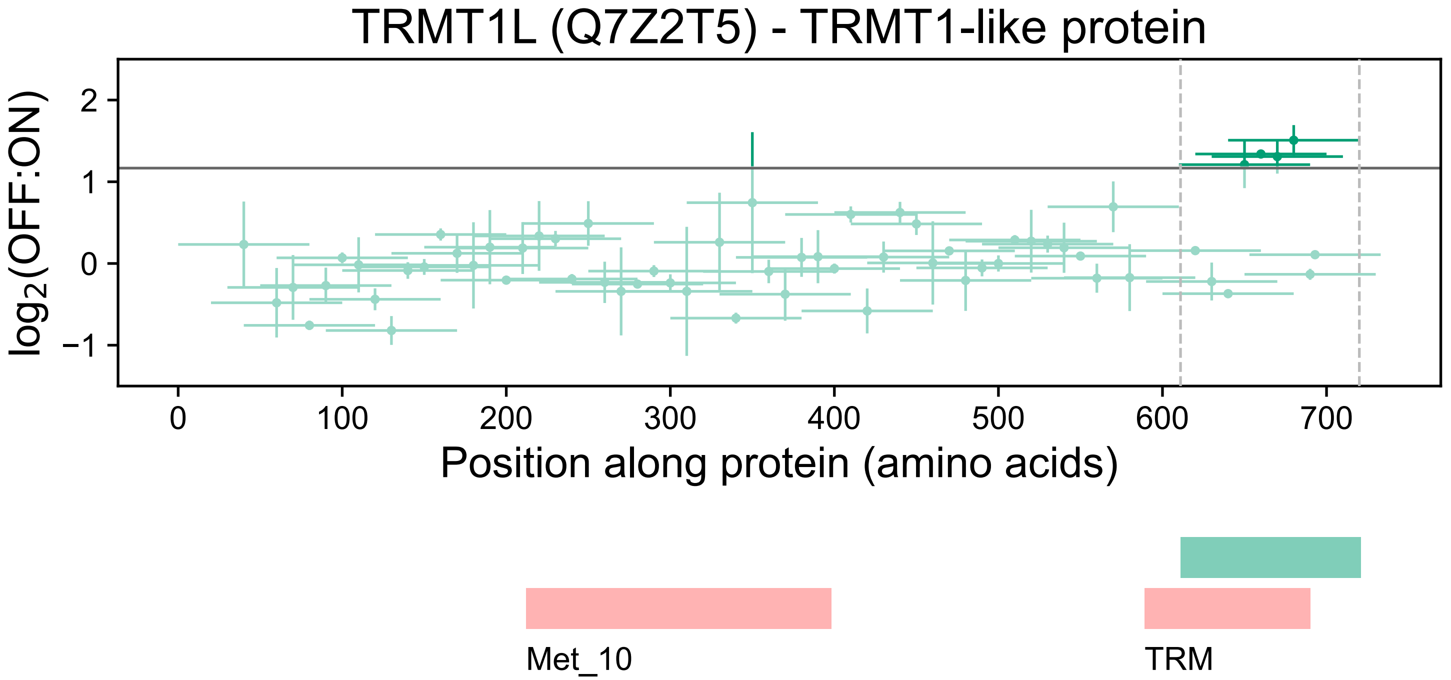
TRMT1L (Q7Z2T5) - TRMT1-like protein
log2(OFF:ON)
2
1
0
−1
0
100
200
300
400
500
600
700
Position along protein (amino acids)
Met_10
TRM

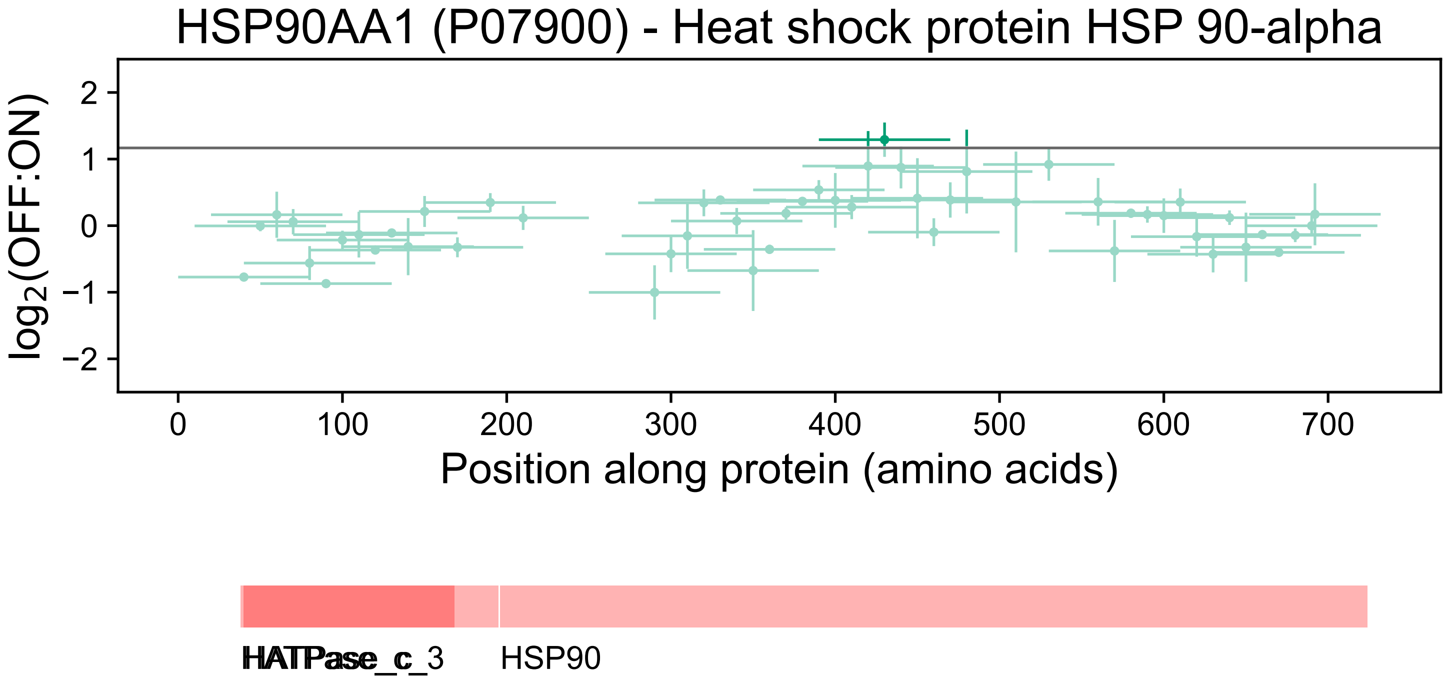
HSP90AA1 (P07900) - Heat shock protein HSP 90-alpha
log2(OFF:ON)
2
1
0
−1
−2
0
100
200
300
400
500
600
700
Position along protein (amino acids)
HATPase_c
HATPase_c_3
HSP90

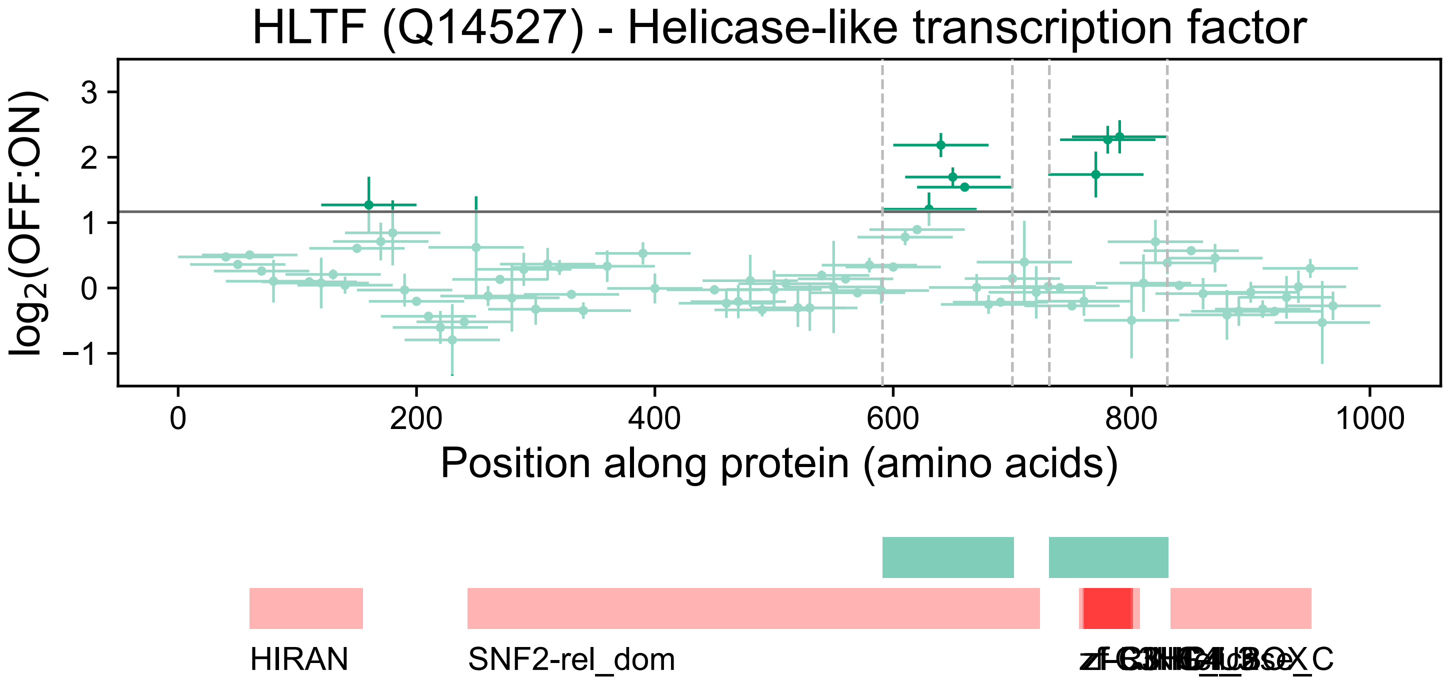
HLTF (Q14527) - Helicase-like transcription factor
log2(OFF:ON)
3
2
1
0
−1
0
200
400
600
800
1000
Position along protein (amino acids)
HIRAN
SNF2-rel_dom

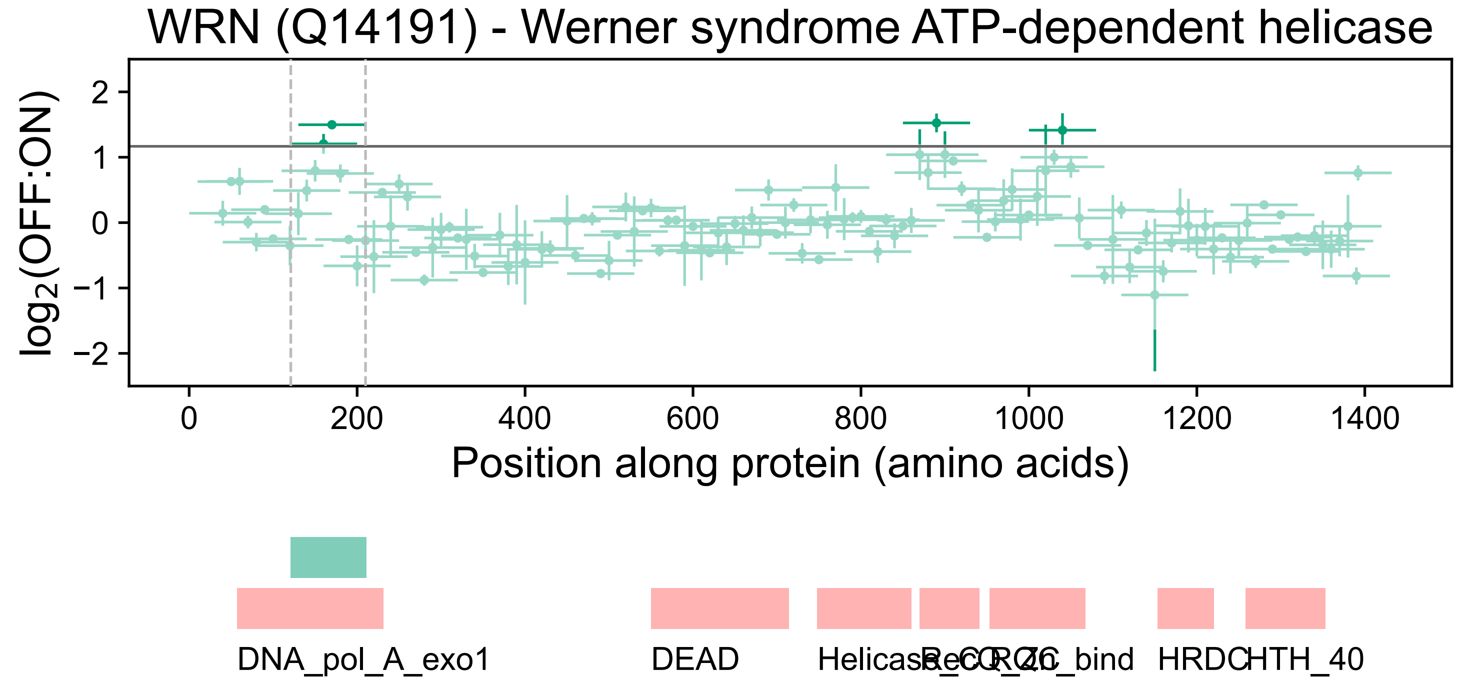
WRN (Q14191) - Werner syndrome ATP-dependent helicase
log2(OFF:ON)
2
1
0
−1
−2
0
200
400
600
800
1000
1200
1400
Position along protein (amino acids)
DNA_pol_A_exo1
DEAD
Helicase_C
RQC
Zn_bind
HRDC
HTH_40

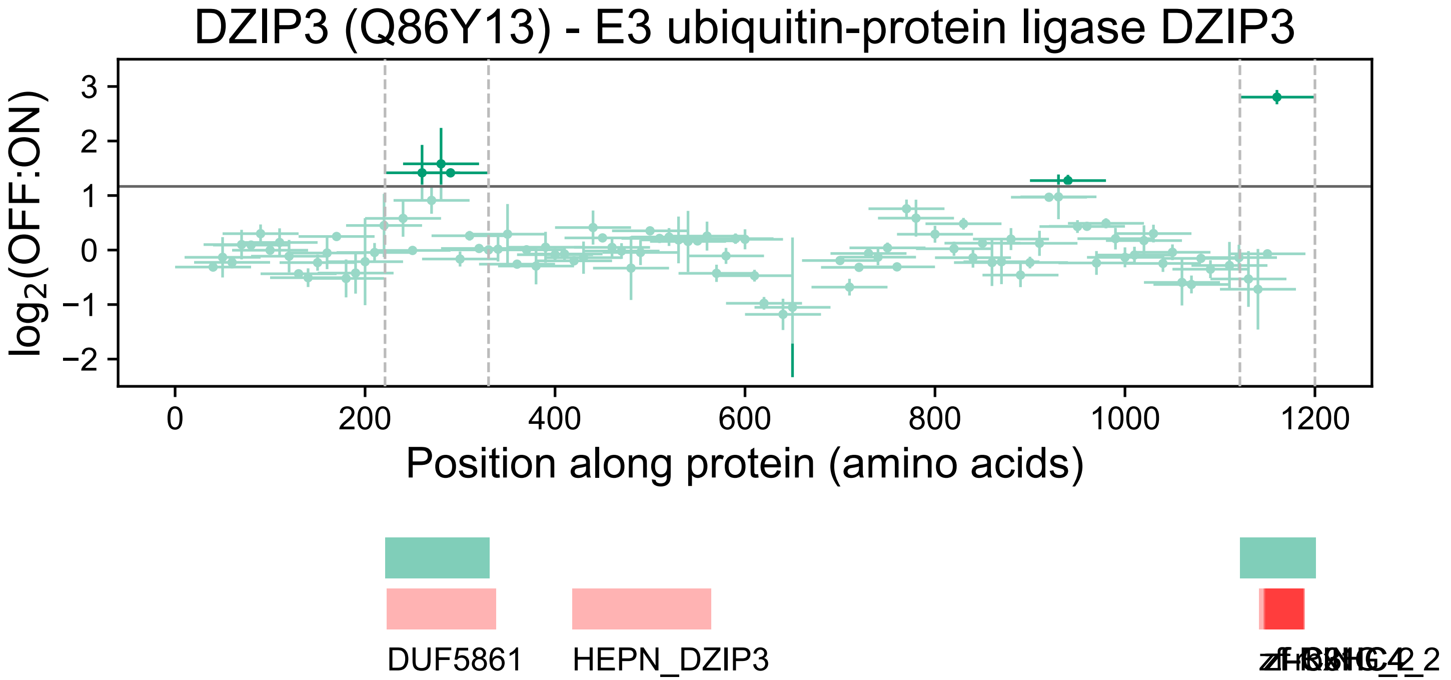

DZIP3 (Q86Y13) - E3 ubiquitin-protein ligase DZIP3
log2(OFF:ON)
3
2
1
0
−1
−2
0
200
400
600
800
1000
1200
Position along protein (amino acids)
DUF5861
HEPN_DZIP3

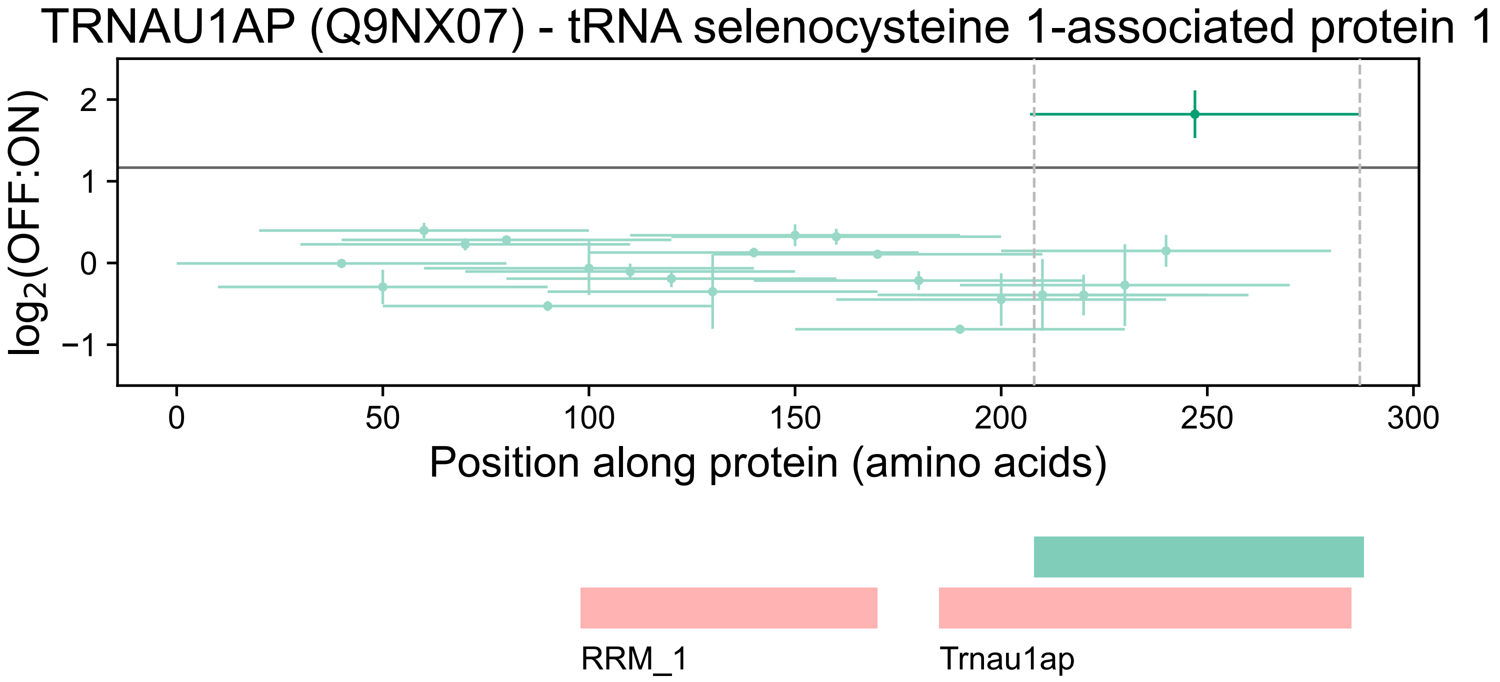
TRNAU1AP (Q9NX07) - tRNA selenocysteine 1-associated protein 1
log2(OFF:ON)
2
1
0
−1
0
50
100
150
200
250
300
Position along protein (amino acids)
RRM_1
Trnau1ap

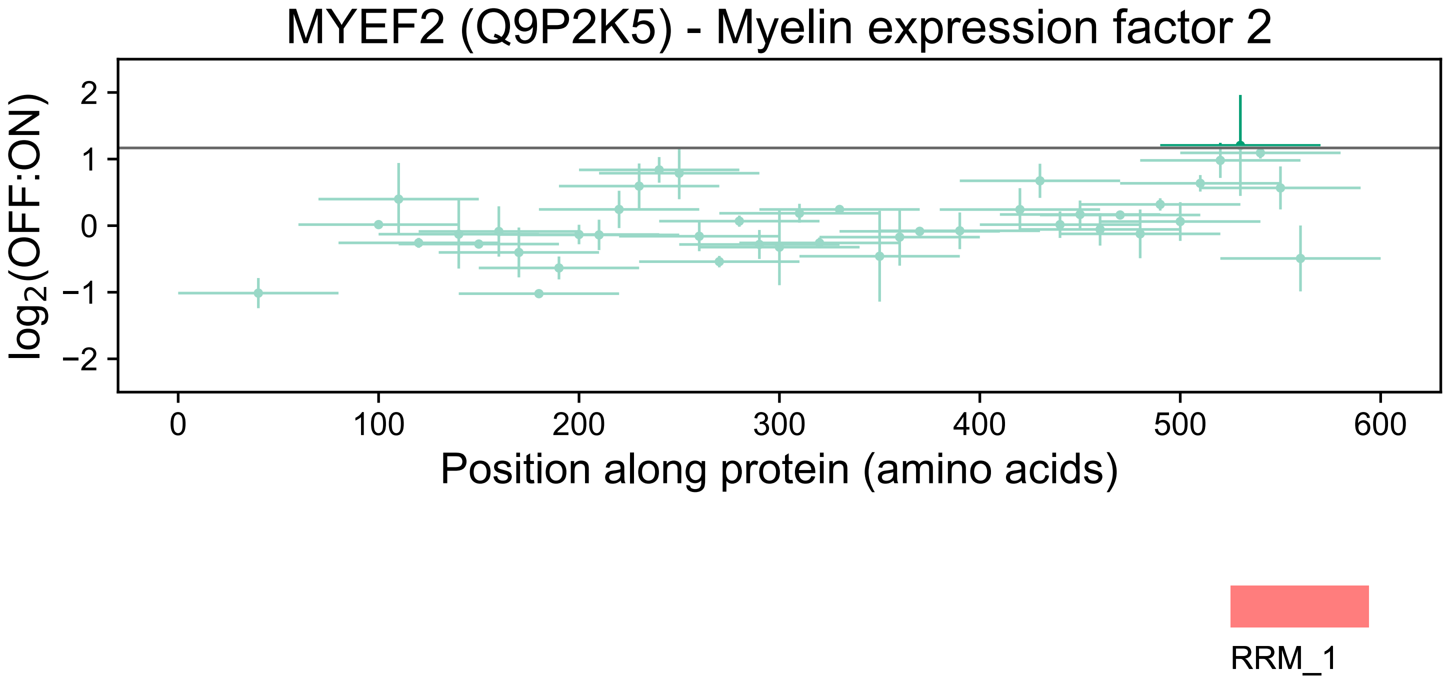
MYEF2 (Q9P2K5) - Myelin expression factor 2
log2(OFF:ON)
2
1
0
−1
−2
0
100
200
300
400
500
600
Position along protein (amino acids)
RRM_1

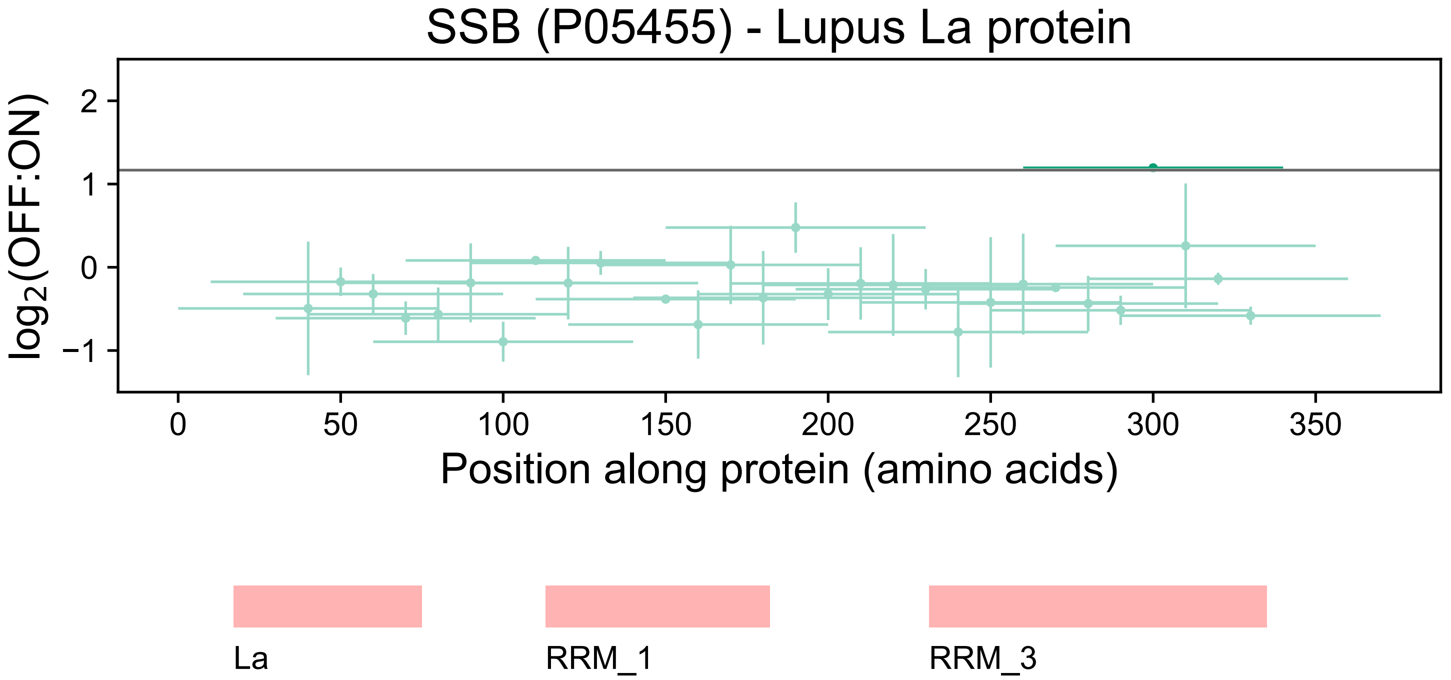
SSB (P05455) - Lupus La protein
log2(OFF:ON)
2
1
0
−1
0
50
100
150
200
250
300
350
Position along protein (amino acids)
La
RRM_1
RRM_3

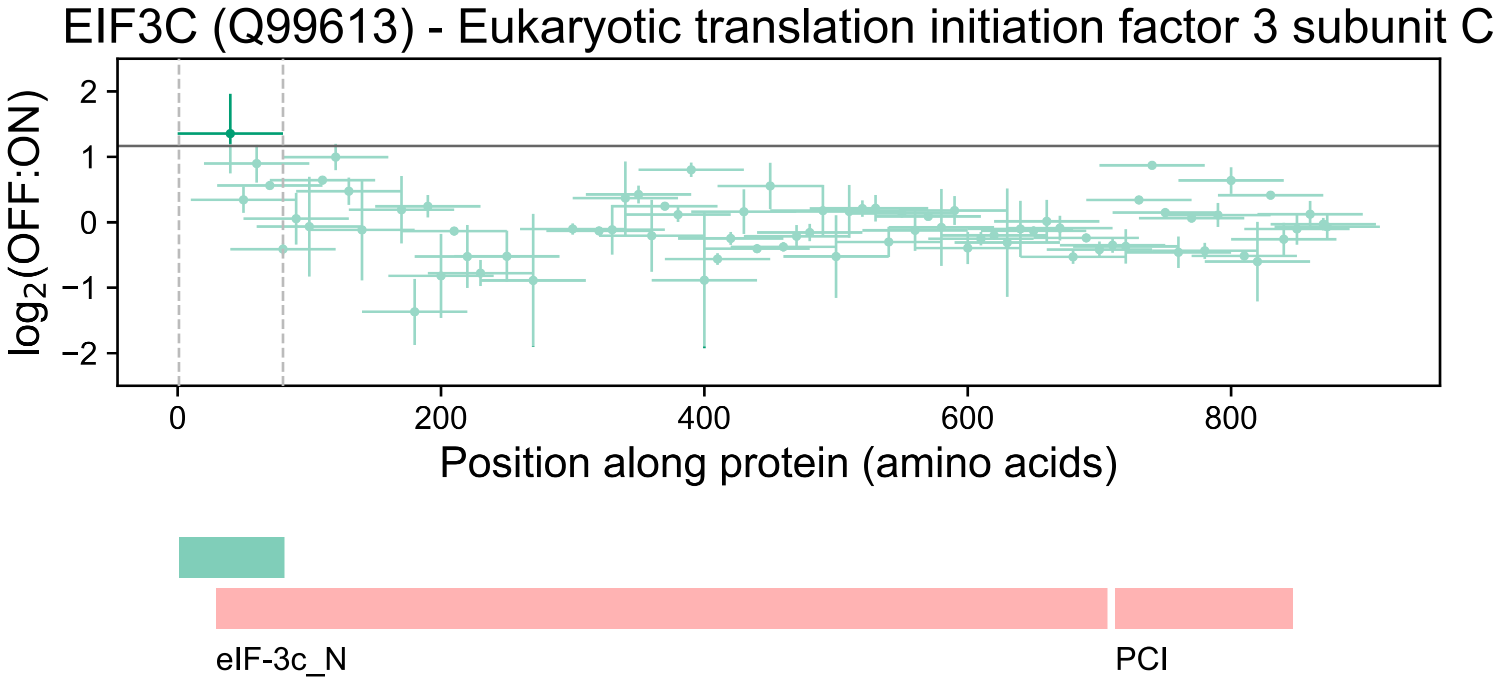
EIF3C (Q99613) - Eukaryotic translation initiation factor 3 subunit C
$\log_2$(OFF:ON)
2
1
0
−1
−2
0
200
400
600
800
Position along protein (amino acids)
eIF-3c_N
PCI

### TNRC6B (Q9UPQ9) - Trinucleotide repeat-containing gene 6B protein

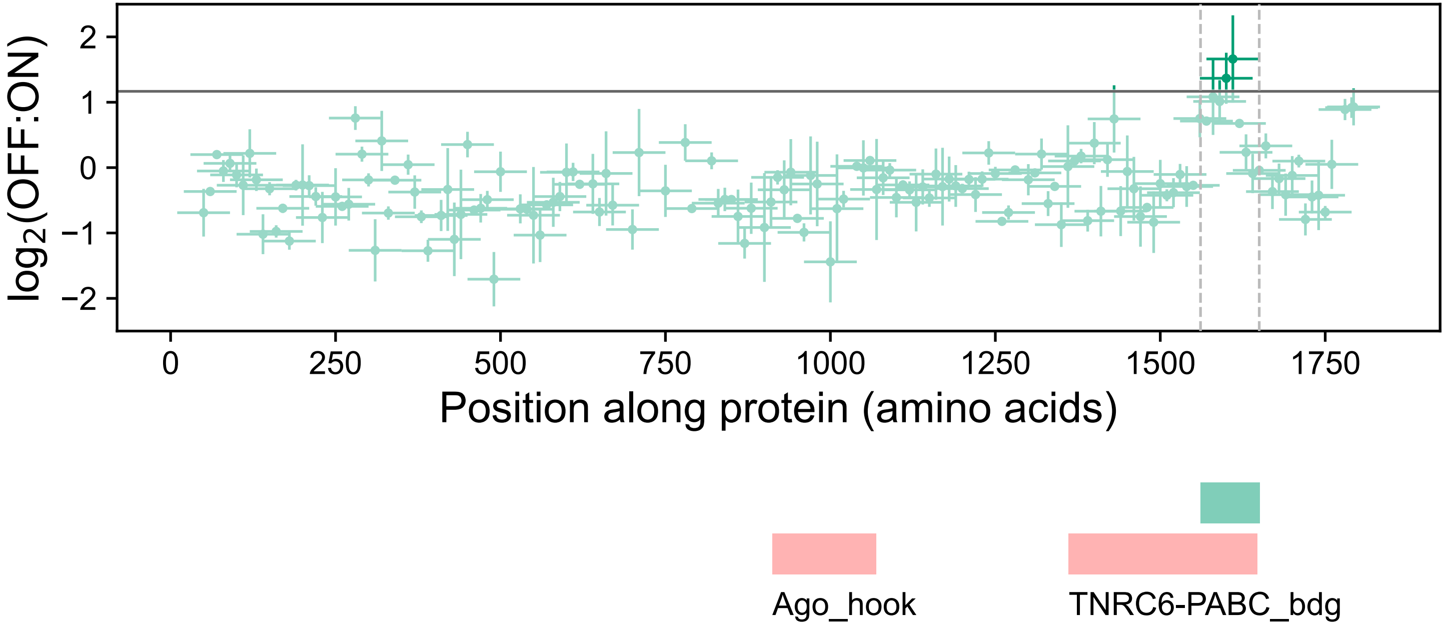

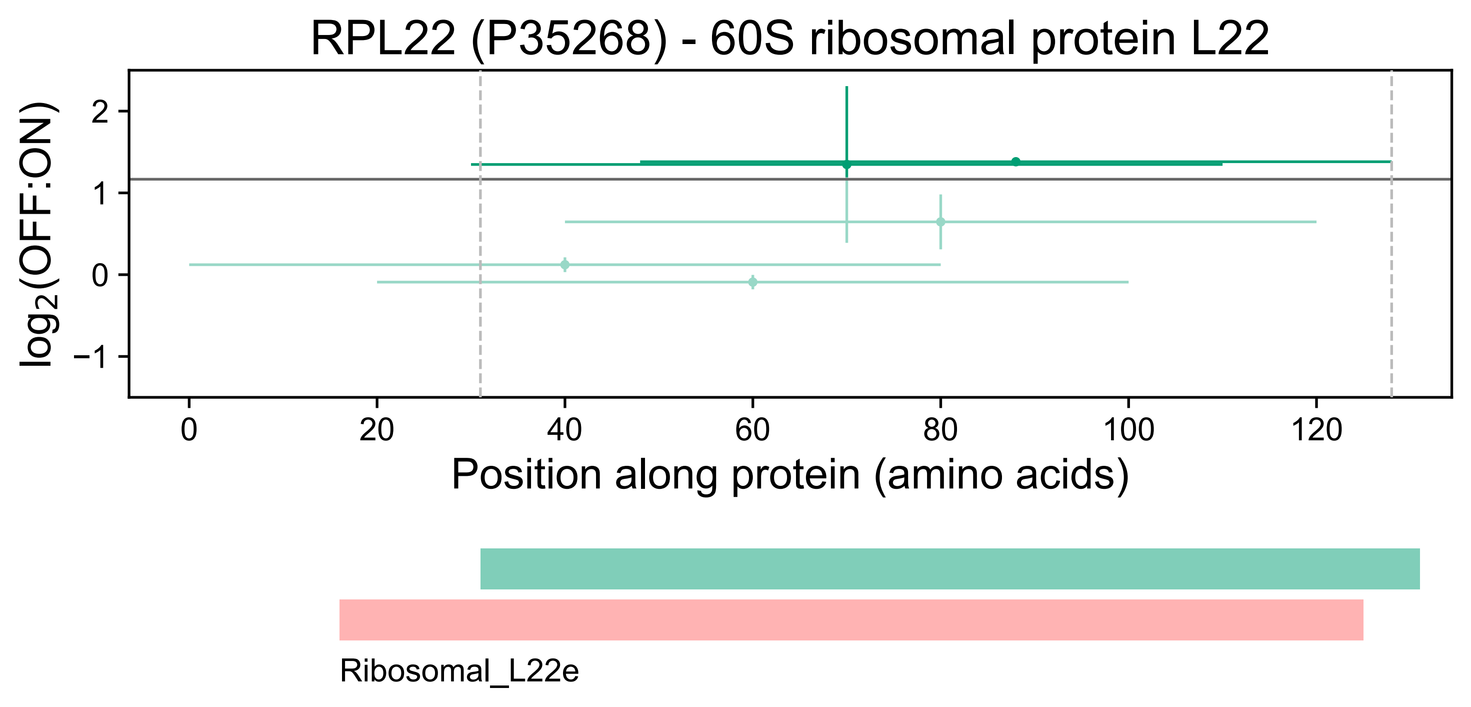
RPL22 (P35268) - 60S ribosomal protein L22
log2(OFF:ON)
2
1
0
−1
0
20
40
60
80
100
120
Position along protein (amino acids)
Ribosomal_L22e

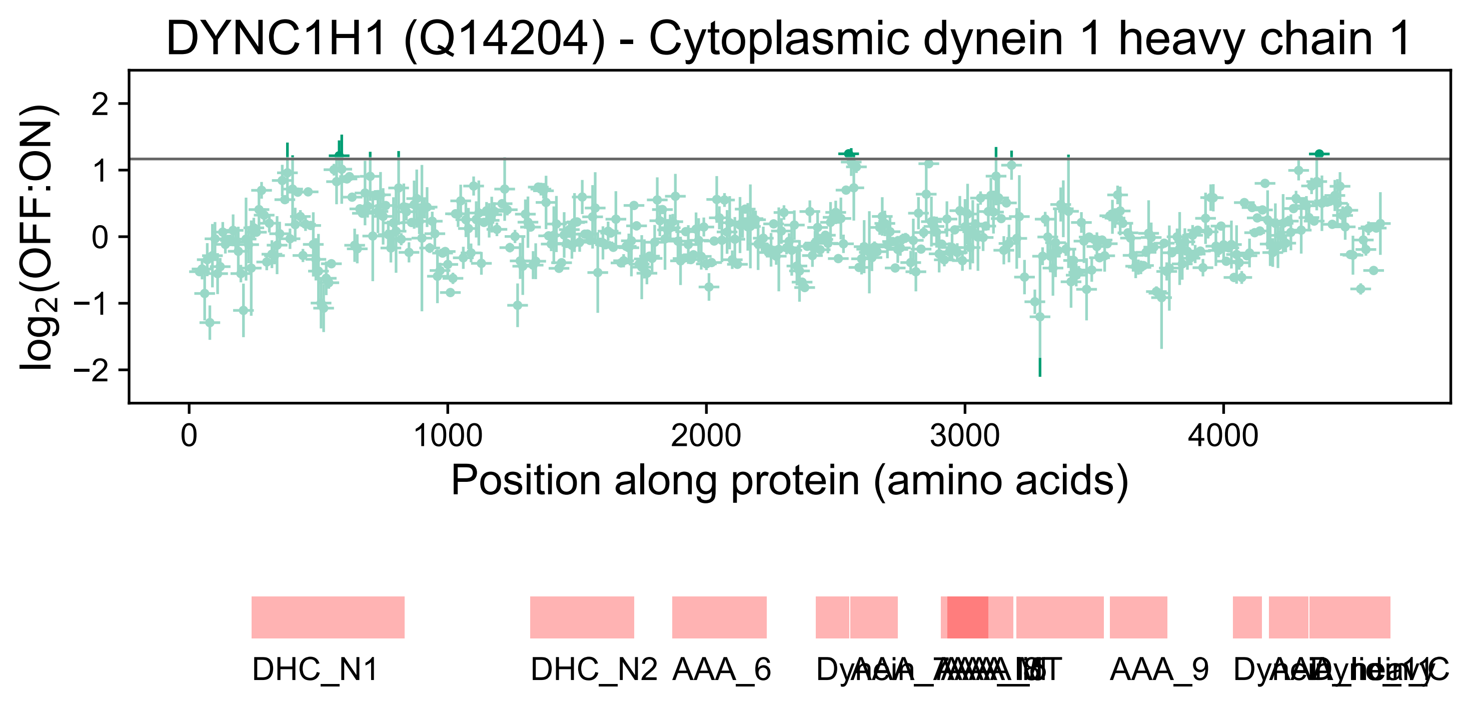
DYNC1H1 (Q14204) - Cytoplasmic dynein 1 heavy chain 1
log2(OFF:ON)
2
1
0
−1
−2
0
1000
2000
3000
4000
Position along protein (amino acids)
DHC_N1
DHC_N2
AAA_6
AAA_9

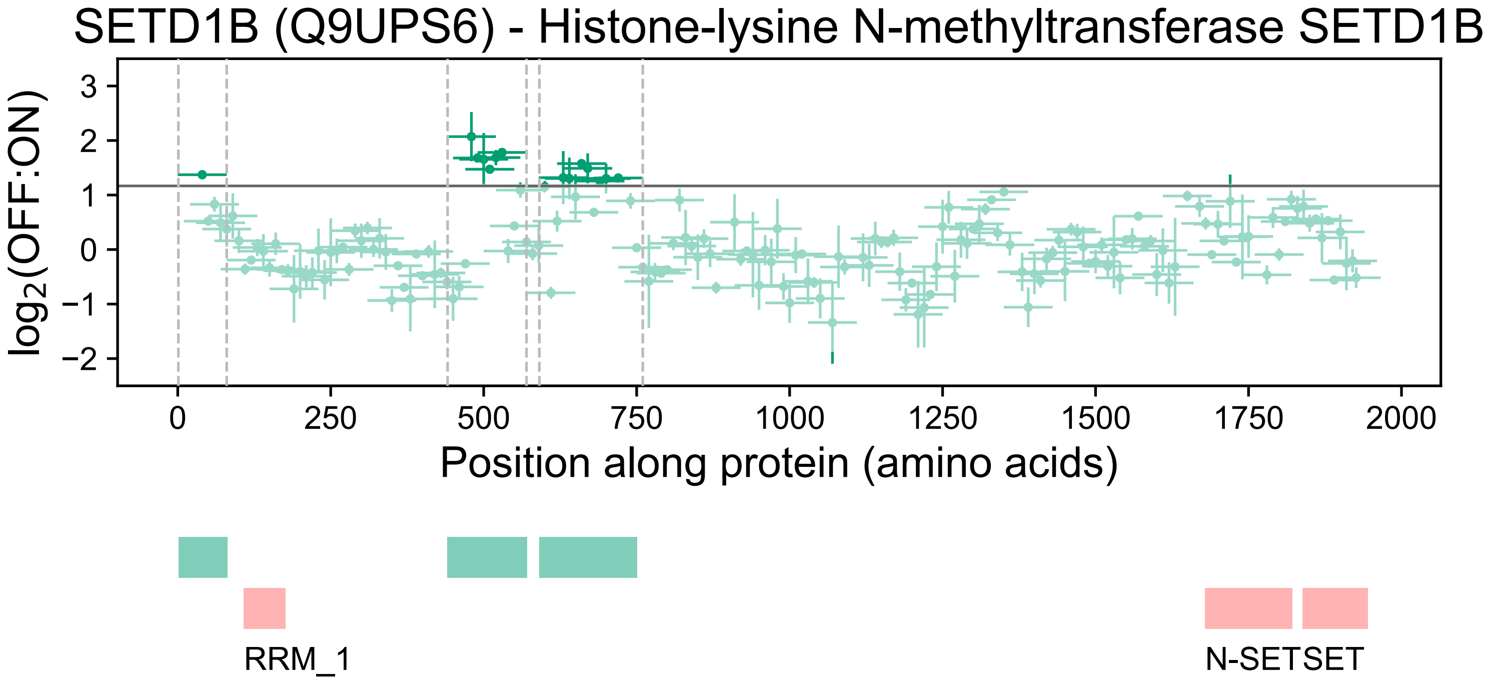

SETD1B (Q9UPS6) - Histone-lysine N-methyltransferase SETD1B
log2(OFF:ON)
3
2
1
0
−1
−2
0
250
500
750
1000
1250
1500
1750
2000
Position along protein (amino acids)
RRM_1
N-SETSET

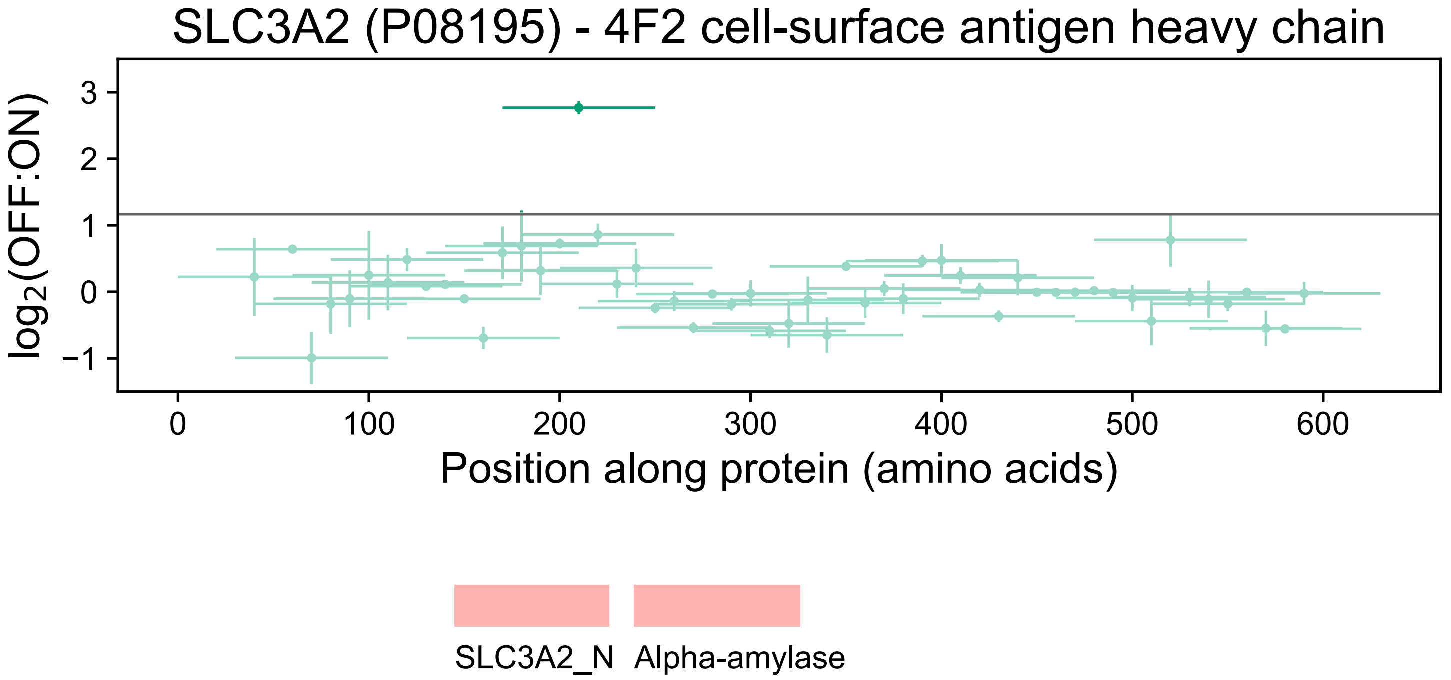
SLC3A2 (P08195) - 4F2 cell-surface antigen heavy chain
log2(OFF:ON)
3
2
1
0
−1
0
100
200
300
400
500
600
Position along protein (amino acids)
SLC3A2_N
Alpha-amylase

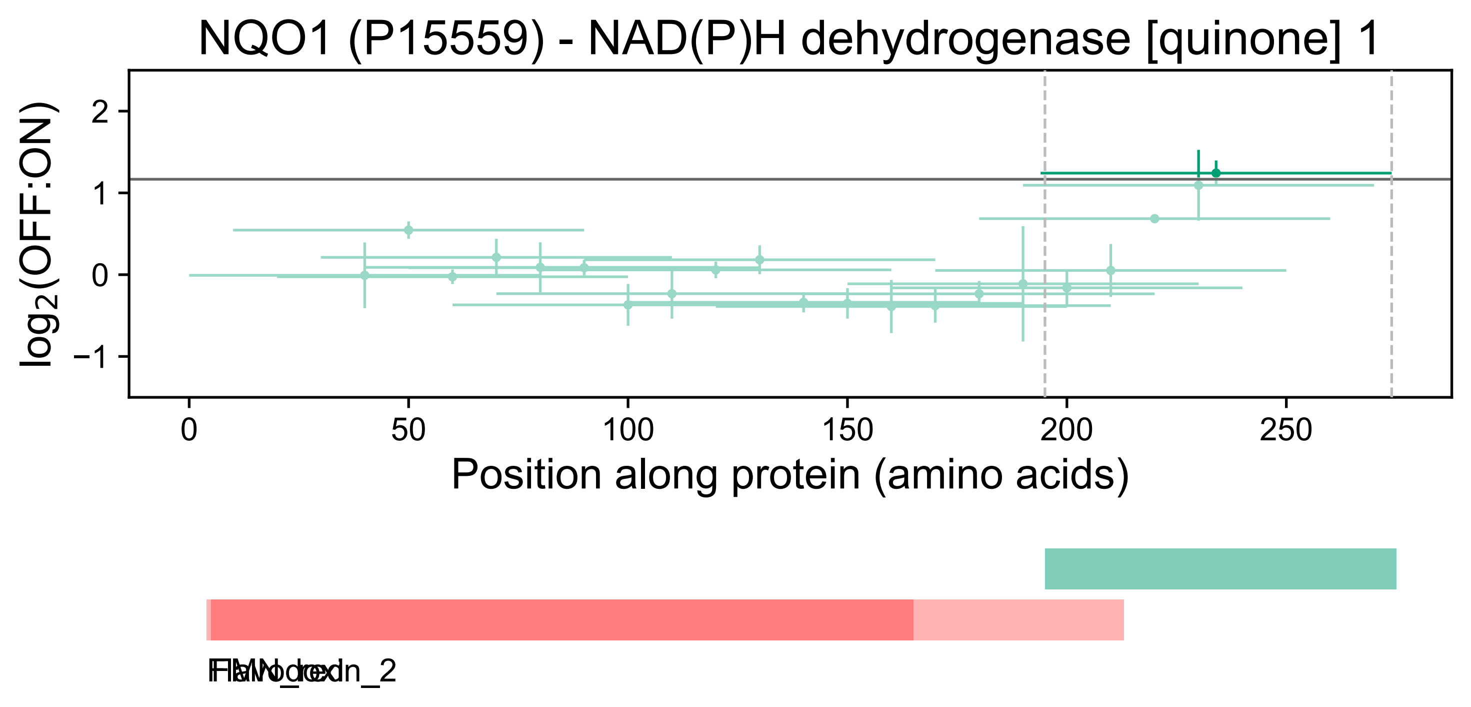
NQO1 (P15559) - NAD(P)H dehydrogenase [quinone] 1
log2(OFF:ON)
2
1
0
−1
0
50
100
150
200
250
Position along protein (amino acids)
FMN_red
Flavodoxin_2

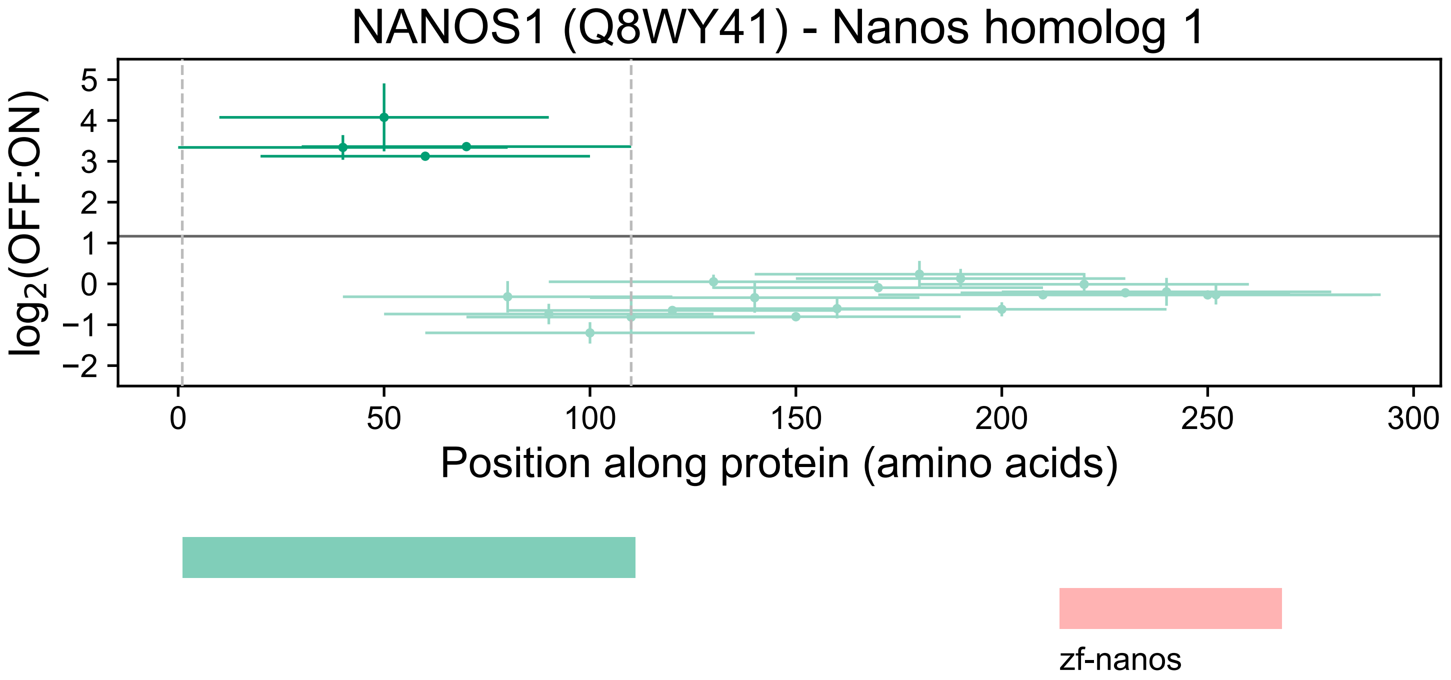
NANOS1 (Q8WY41) - Nanos homolog 1
log2(OFF:ON)
5
4
3
2
1
0
−1
−2
0
50
100
150
200
250
300
Position along protein (amino acids)
zf-nanos

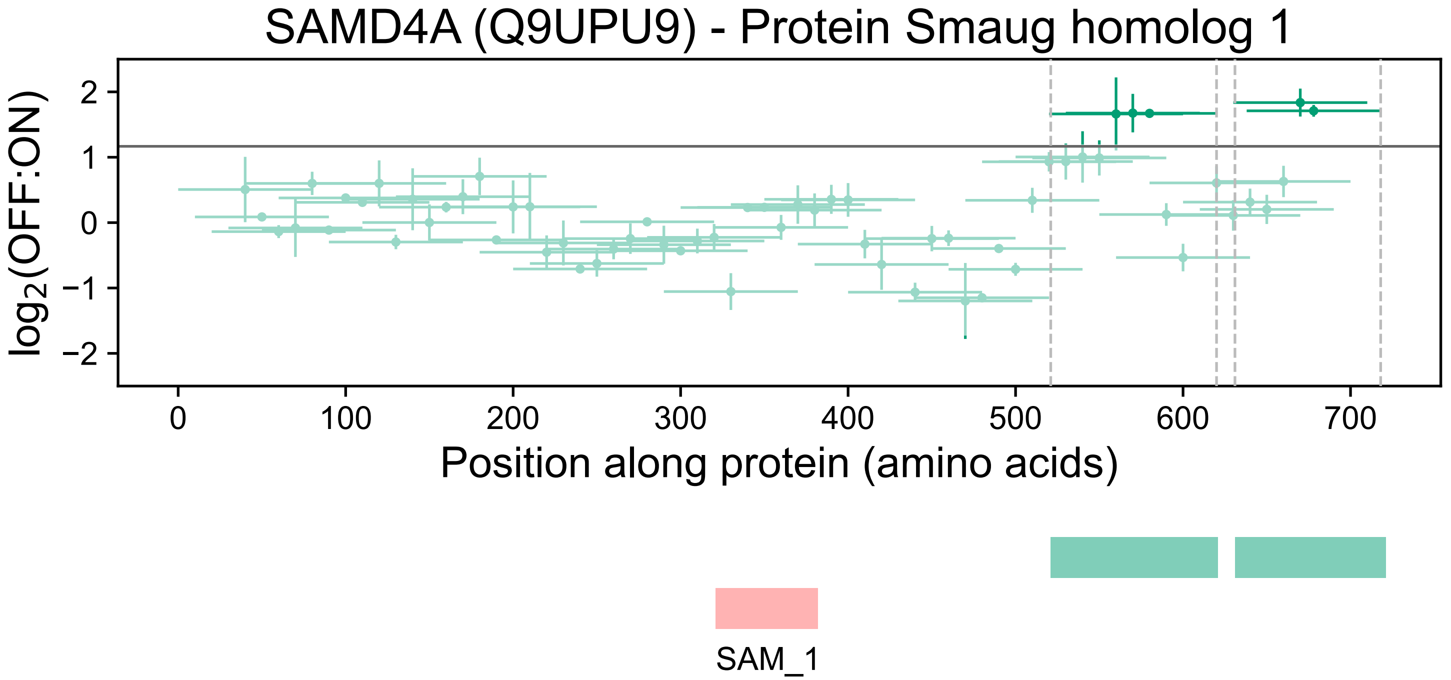
SAMD4A (Q9UPU9) - Protein Smaug homolog 1
log2(OFF:ON)
2
1
0
−1
−2
0
100
200
300
400
500
600
700
Position along protein (amino acids)
SAM_1

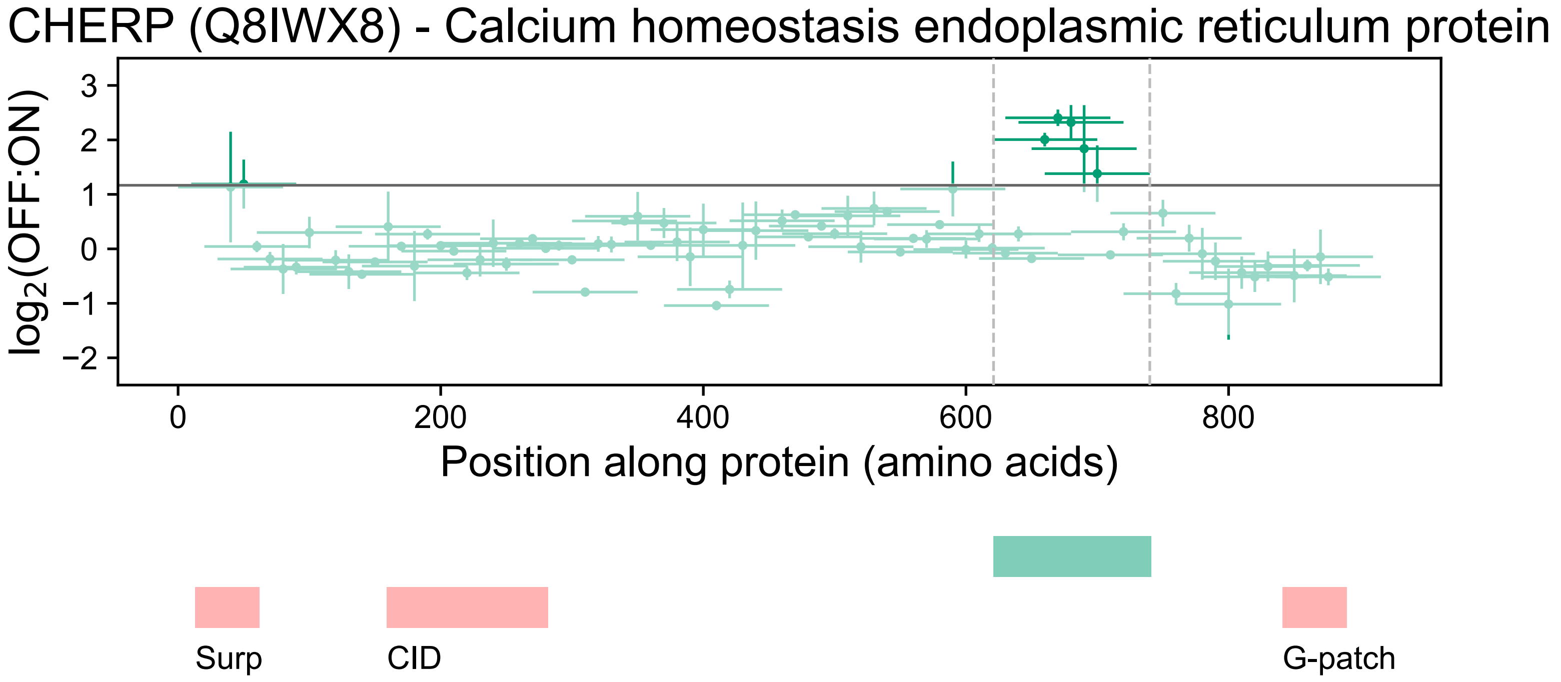

CHERP (Q8IWX8) - Calcium homeostasis endoplasmic reticulum protein
log2(OFF:ON)
3
2
1
0
−1
−2
0
200
400
600
800
Position along protein (amino acids)
Surp
CID
G-patch

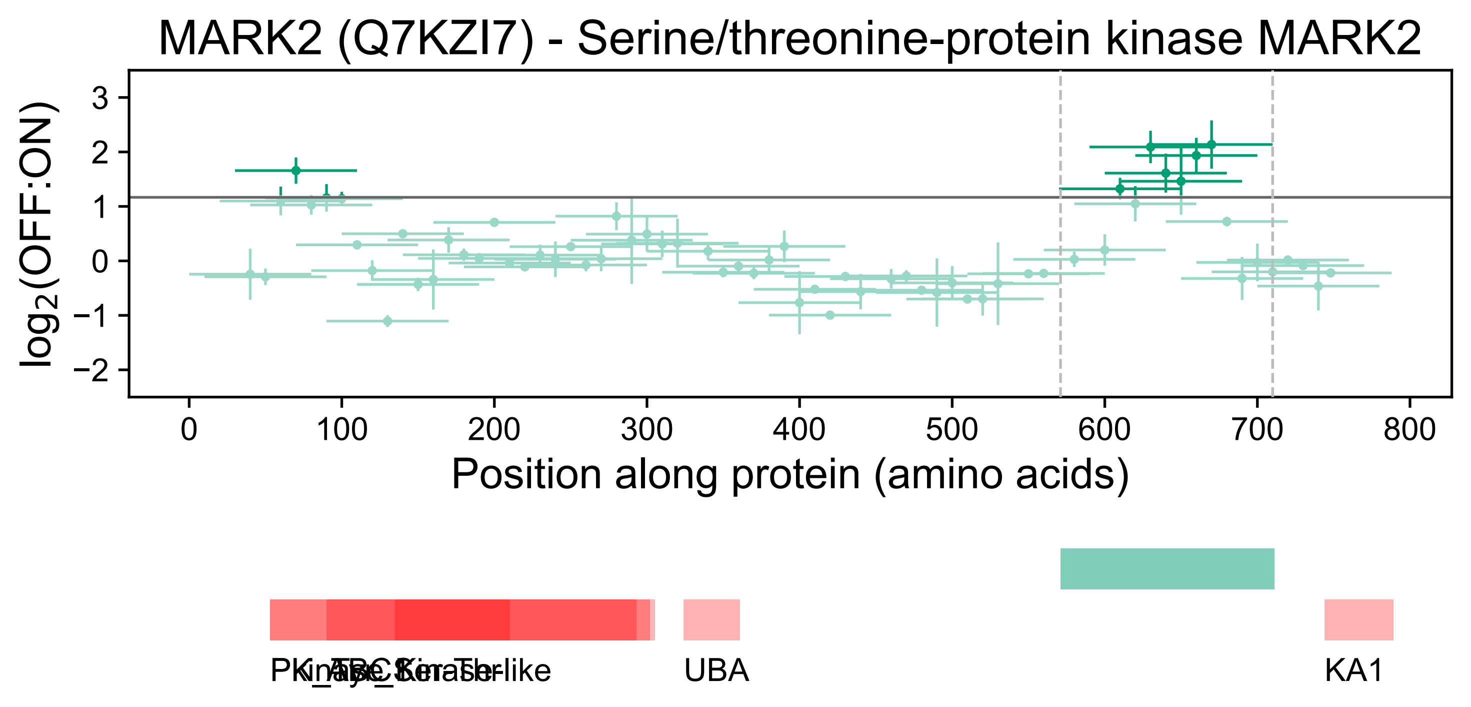
MARK2 (Q7KZI7) - Serine/threonine-protein kinase MARK2
log2(OFF:ON)
Position along protein (amino acids)
3
2
1
0
−1
−2
0
100
200
300
400
500
600
700
800
UBA
KA1

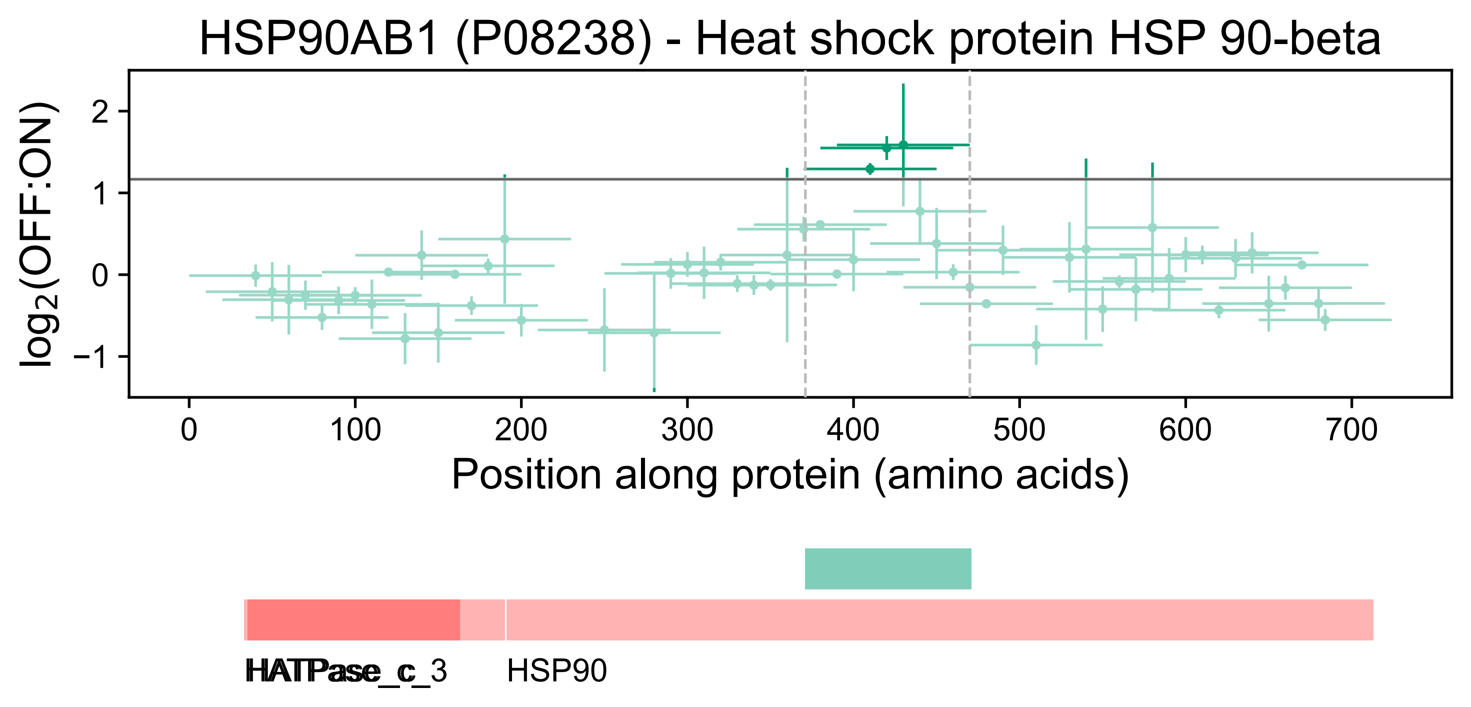
HSP90AB1 (P08238) - Heat shock protein HSP 90-beta
log2(OFF:ON)
2
1
0
−1
0
100
200
300
400
500
600
700
Position along protein (amino acids)
HATPase_c
HATPase_c_3
HSP90

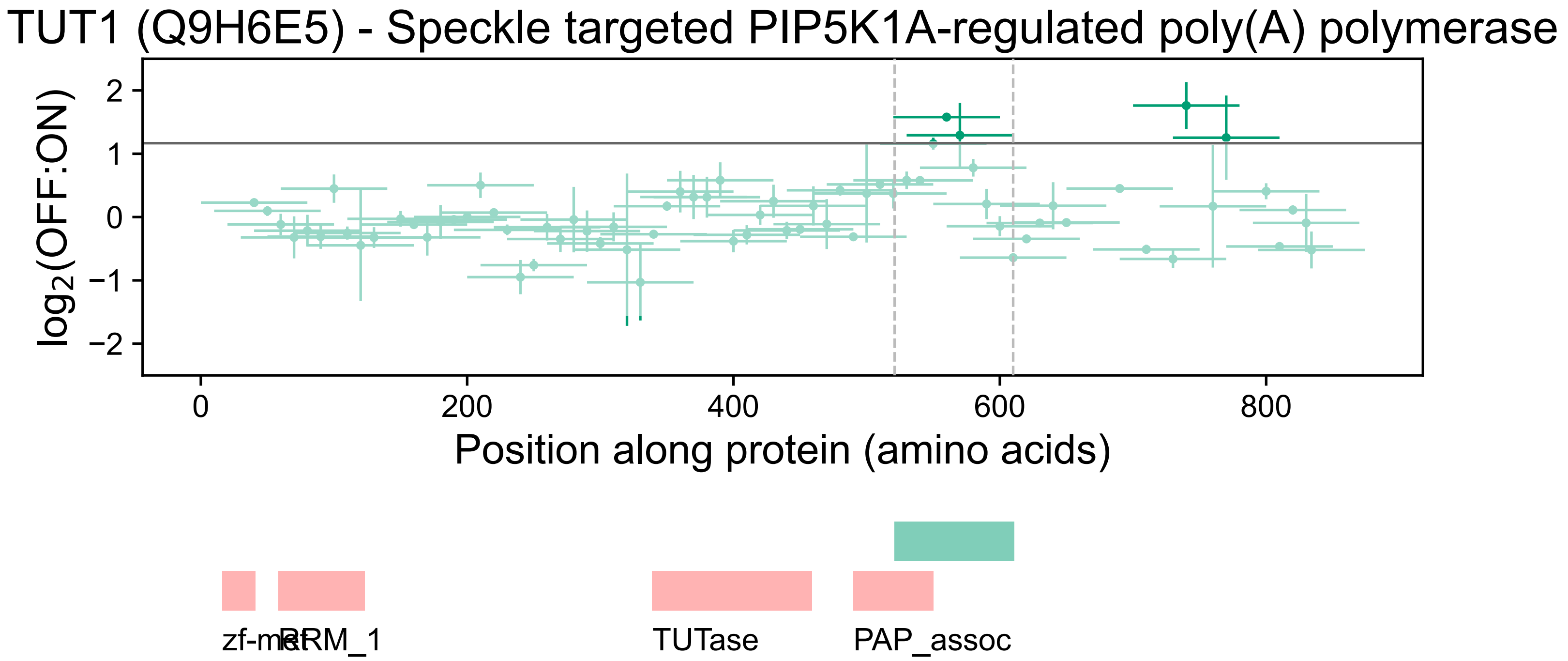
TUT1 (Q9H6E5) - Speckle targeted PIP5K1A-regulated poly(A) polymerase
log2(OFF:ON)
2
1
0
−1
−2
0
200
400
600
800
Position along protein (amino acids)
zf-met
RRM_1
TUTase
PAP_assoc

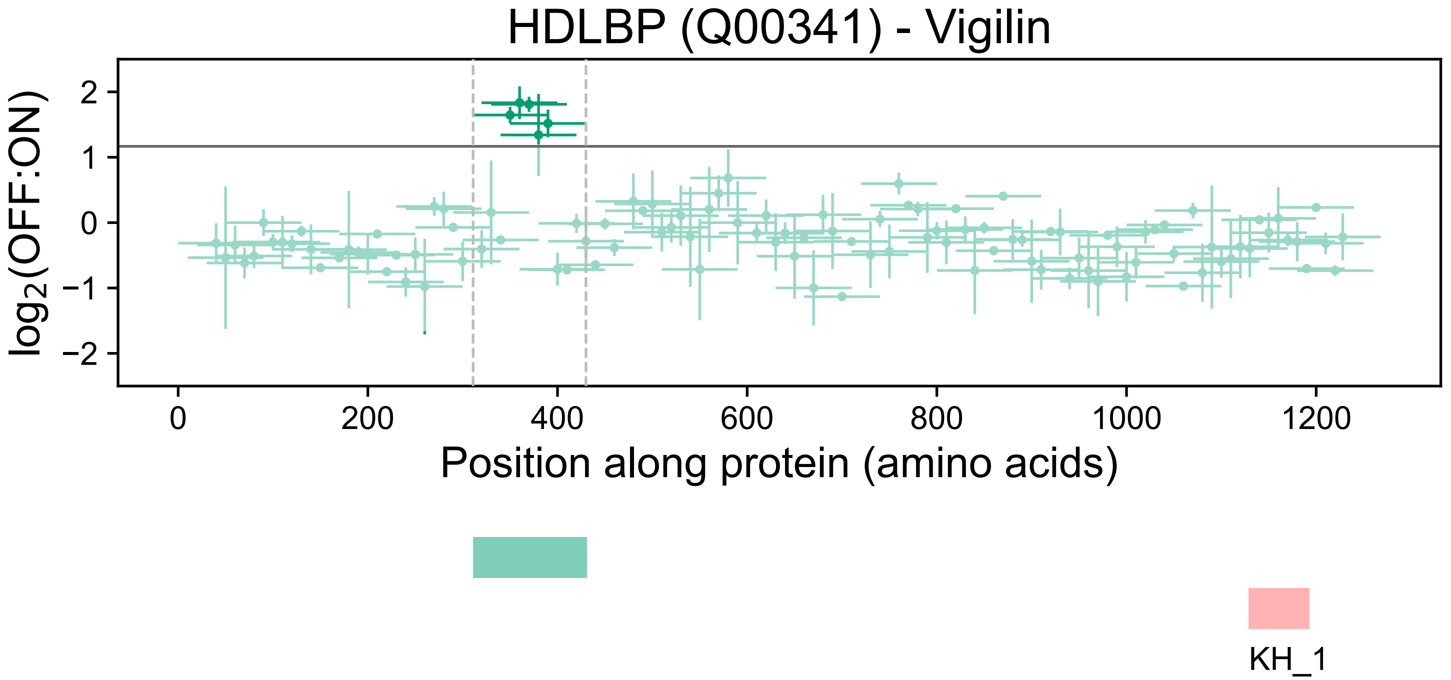
HDLBP (Q00341) - Vigilin
log2(OFF:ON)
2
1
0
−1
−2
0
200
400
600
800
1000
1200
Position along protein (amino acids)
KH_1

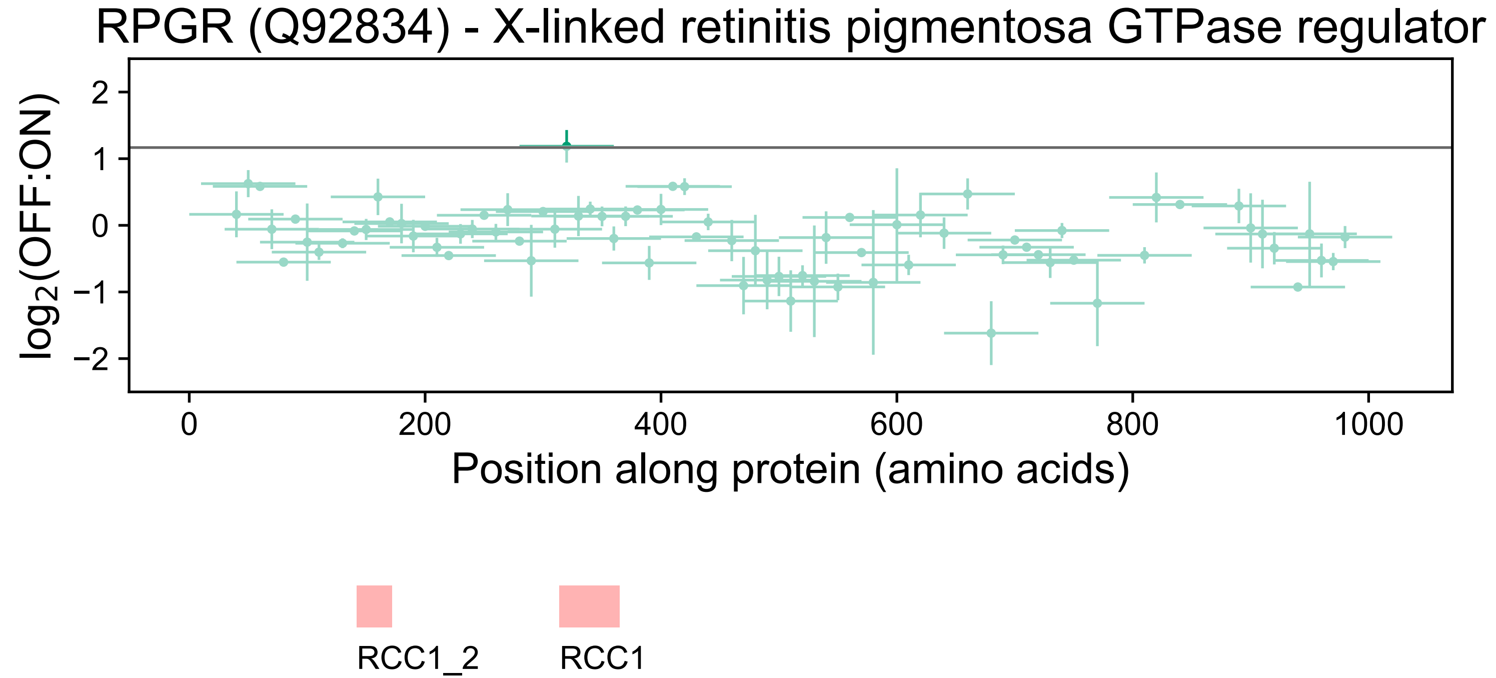
RPGR (Q92834) - X-linked retinitis pigmentosa GTPase regulator
log2(OFF:ON)
2
1
0
−1
−2
0
200
400
600
800
1000
Position along protein (amino acids)
RCC1_2
RCC1

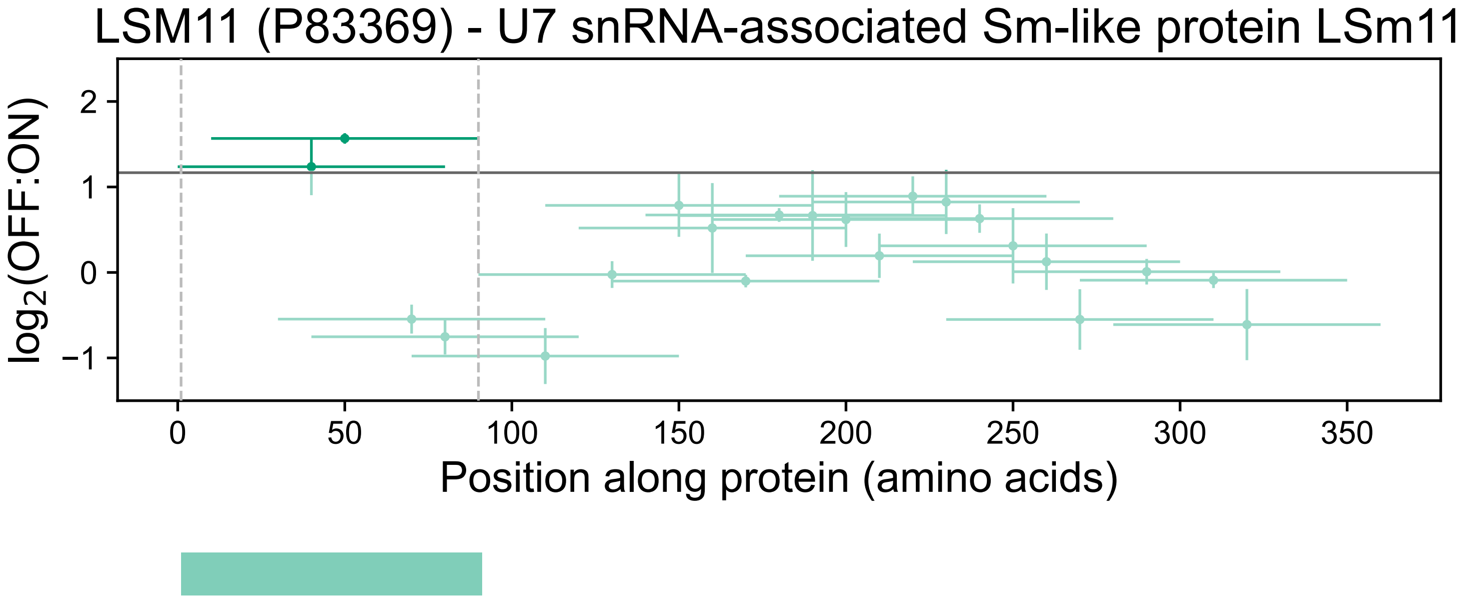

LSM11 (P83369) - U7 snRNA-associated Sm-like protein LSm11
log2(OFF:ON)
2
1
0
−1
0
50
100
150
200
250
300
350
Position along protein (amino acids)

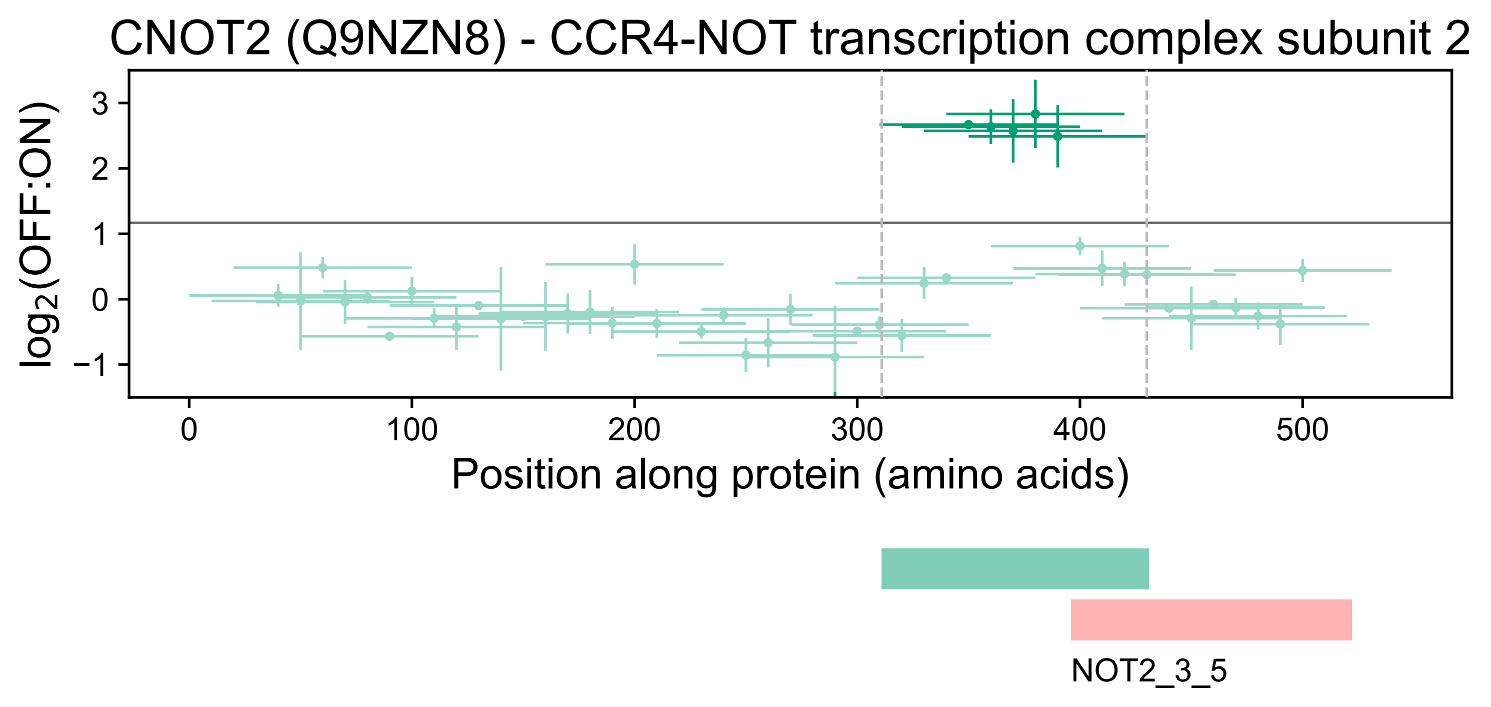

CNOT2 (Q9NZN8) - CCR4-NOT transcription complex subunit 2
log2(OFF:ON)
3
2
1
0
−1
0
100
200
300
400
500
Position along protein (amino acids)
NOT2_3_5

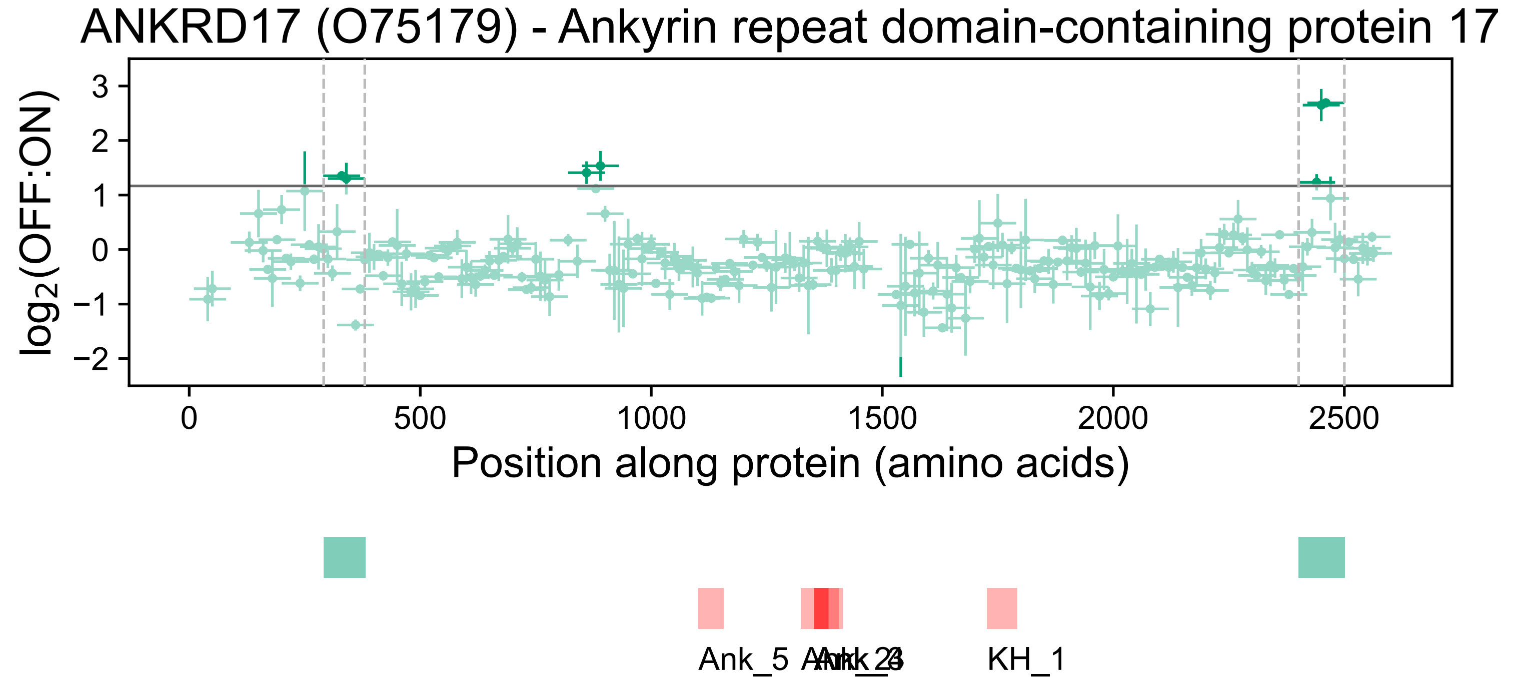
ANKRD17 (O75179) - Ankyrin repeat domain-containing protein 17
log2(OFF:ON)
Position along protein (amino acids)
Ank_5
Ank_2
Ank_4
KH_1

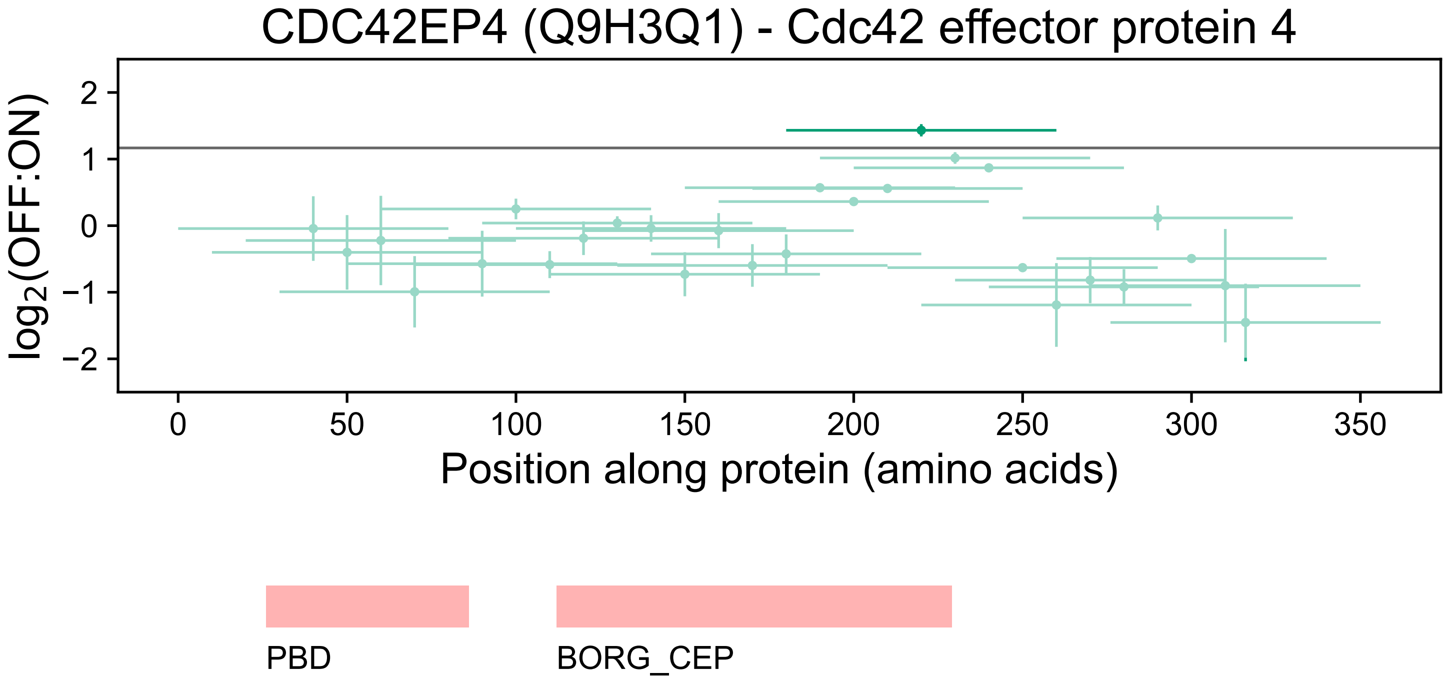

CDC42EP4 (Q9H3Q1) - Cdc42 effector protein 4
log2(OFF:ON)
2
1
0
−1
−2
0
50
100
150
200
250
300
350
Position along protein (amino acids)
PBD
BORG_CEP

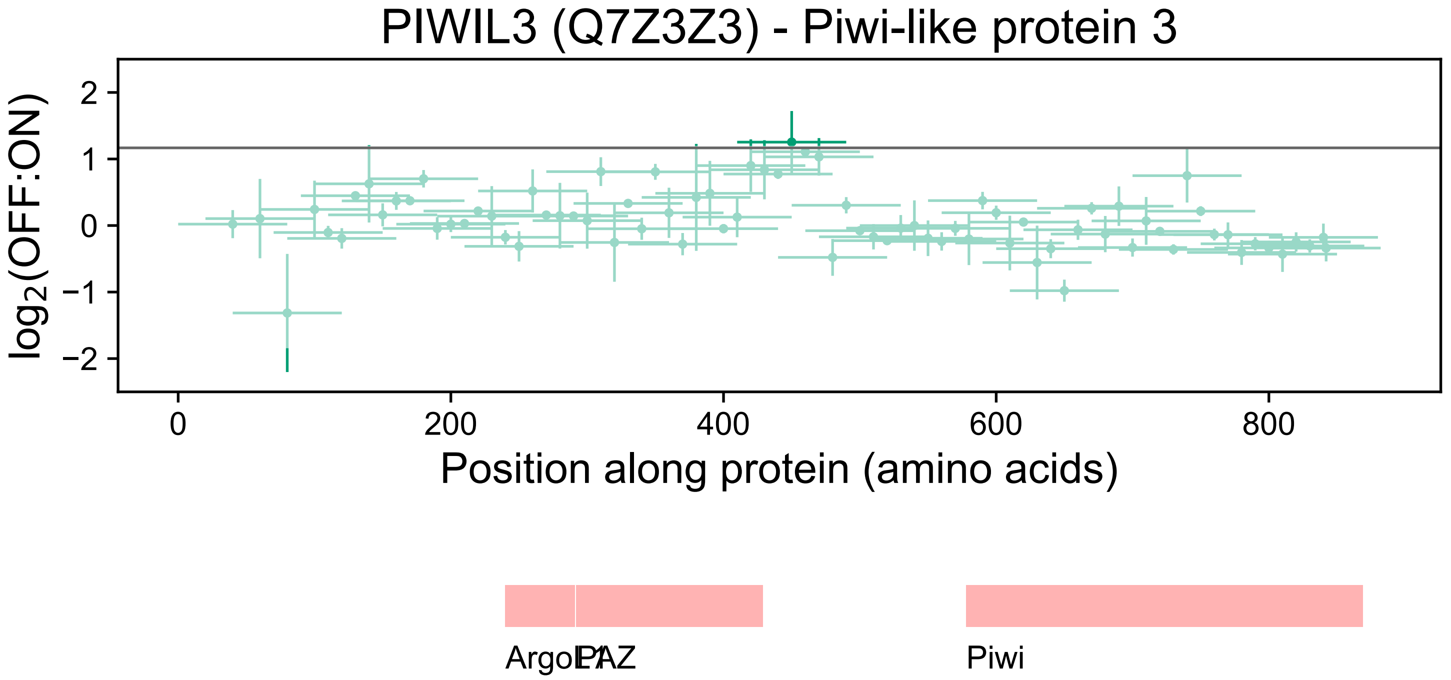
PIWIL3 (Q7Z3Z3) - Piwi-like protein 3
log2(OFF:ON)
2
1
0
−1
−2
0
200
400
600
800
Position along protein (amino acids)
ArgoL1
PAZ
Piwi

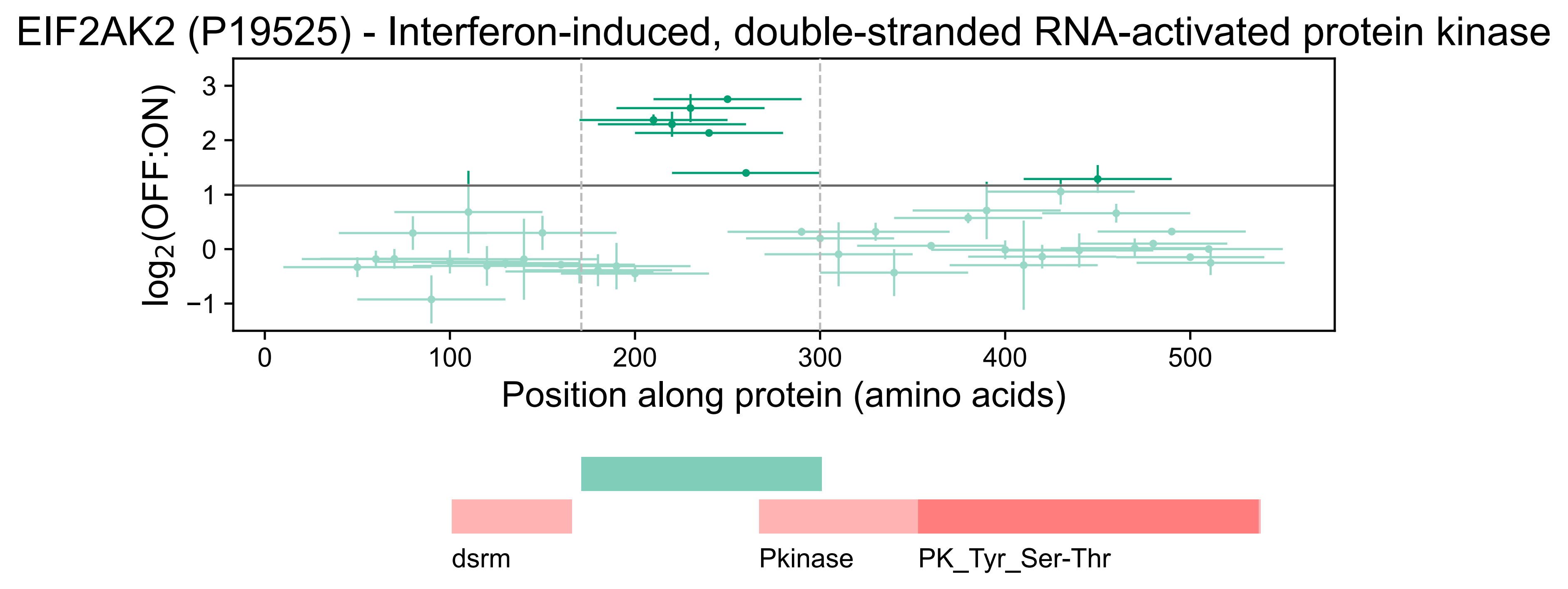

EIF2AK2 (P19525) - Interferon-induced, double-stranded RNA-activated protein kinase
log2(OFF:ON)
3
2
1
0
−1
0
100
200
300
400
500
Position along protein (amino acids)
dsrm
Pkinase
PK_Tyr_Ser-Thr

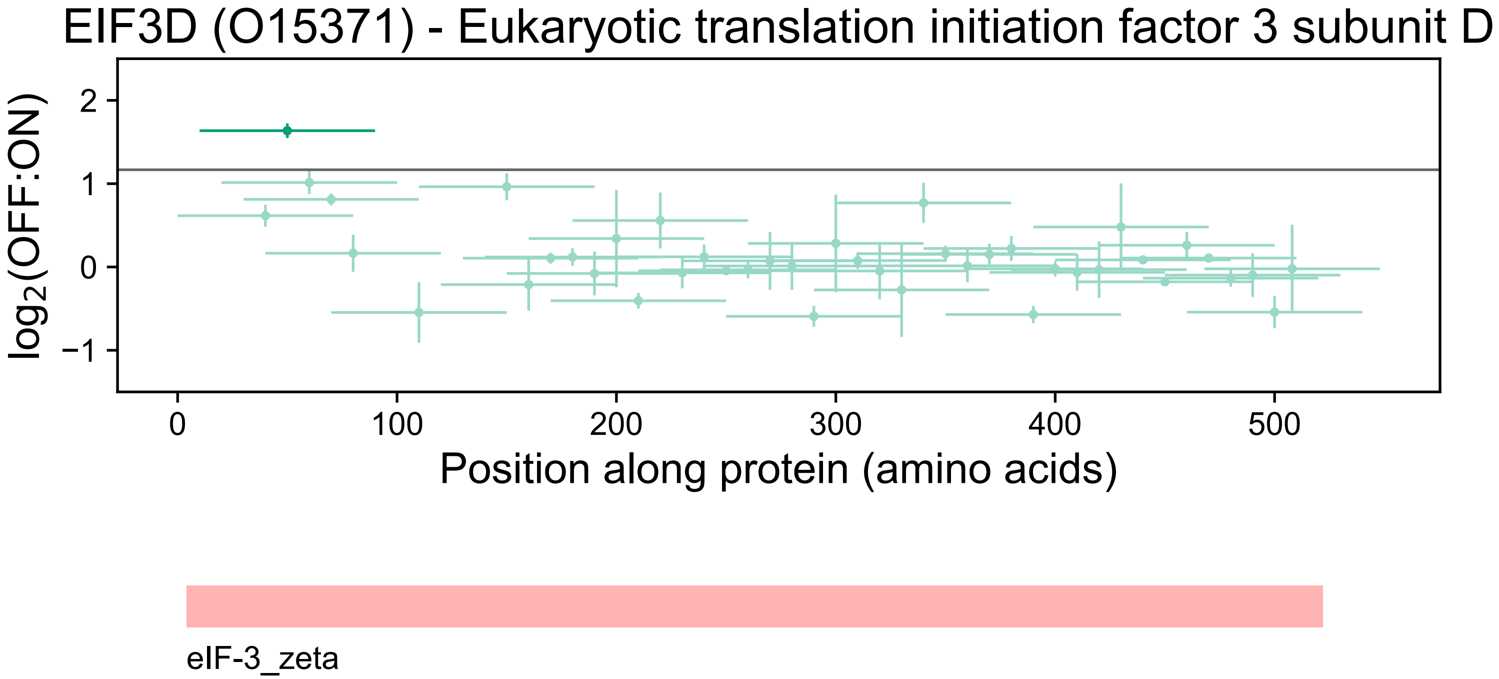
EIF3D (O15371) - Eukaryotic translation initiation factor 3 subunit D
log2(OFF:ON)
2
1
0
−1
0
100
200
300
400
500
Position along protein (amino acids)
eIF-3_zeta

### HNRNPUL1 (Q9BUJ2) - Heterogeneous nuclear ribonucleoprotein U-like protein 1

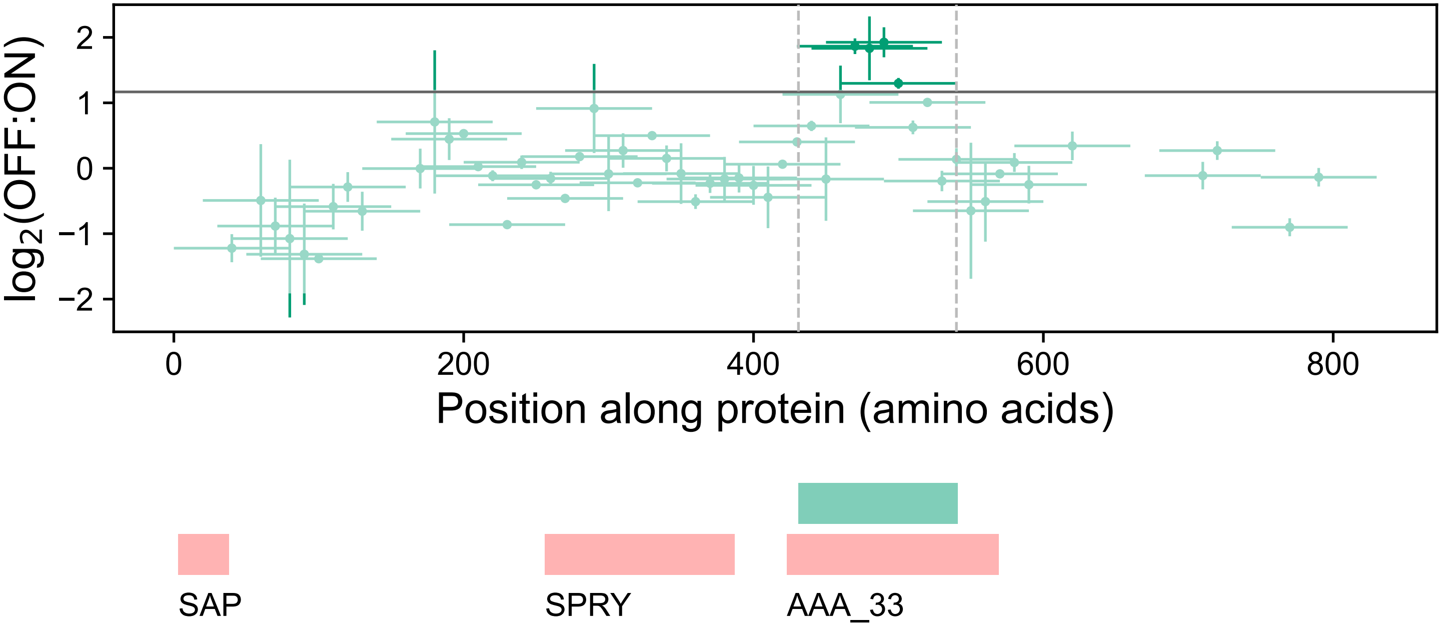

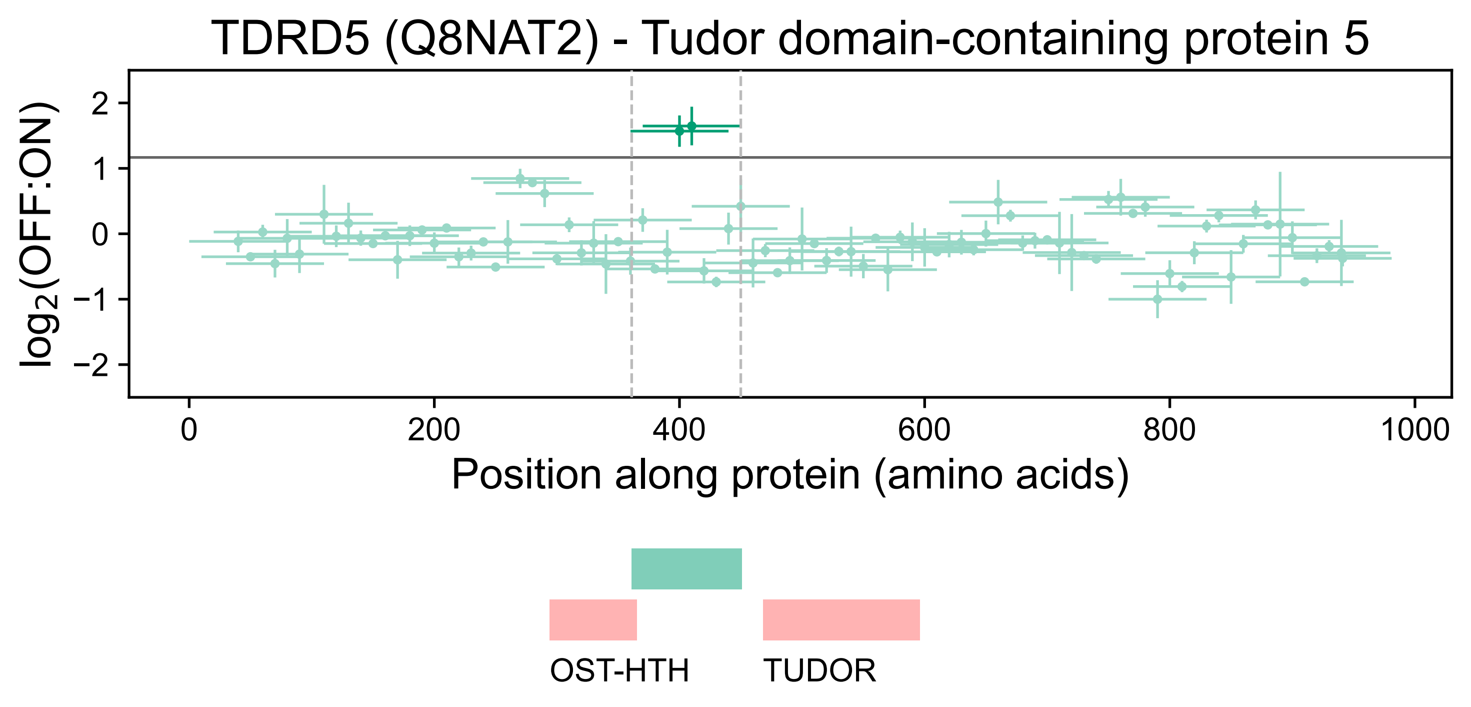
TDRD5 (Q8NAT2) - Tudor domain-containing protein 5
log2(OFF:ON)
2
1
0
−1
−2
0
200
400
600
800
1000
Position along protein (amino acids)
OST-HTH
TUDOR

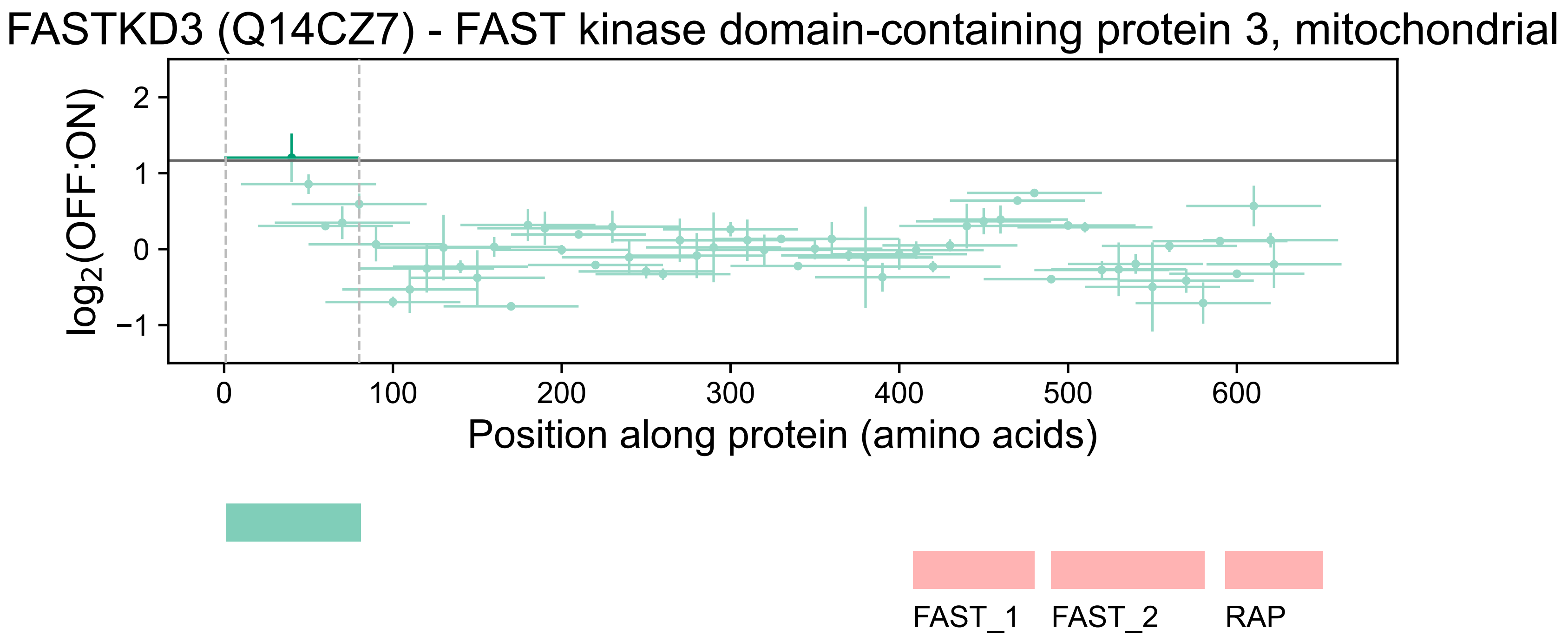

FASTKD3 (Q14CZ7) - FAST kinase domain-containing protein 3, mitochondrial
log2(OFF:ON)
2
1
0
−1
0
100
200
300
400
500
600
Position along protein (amino acids)
FAST_1
FAST_2
RAP

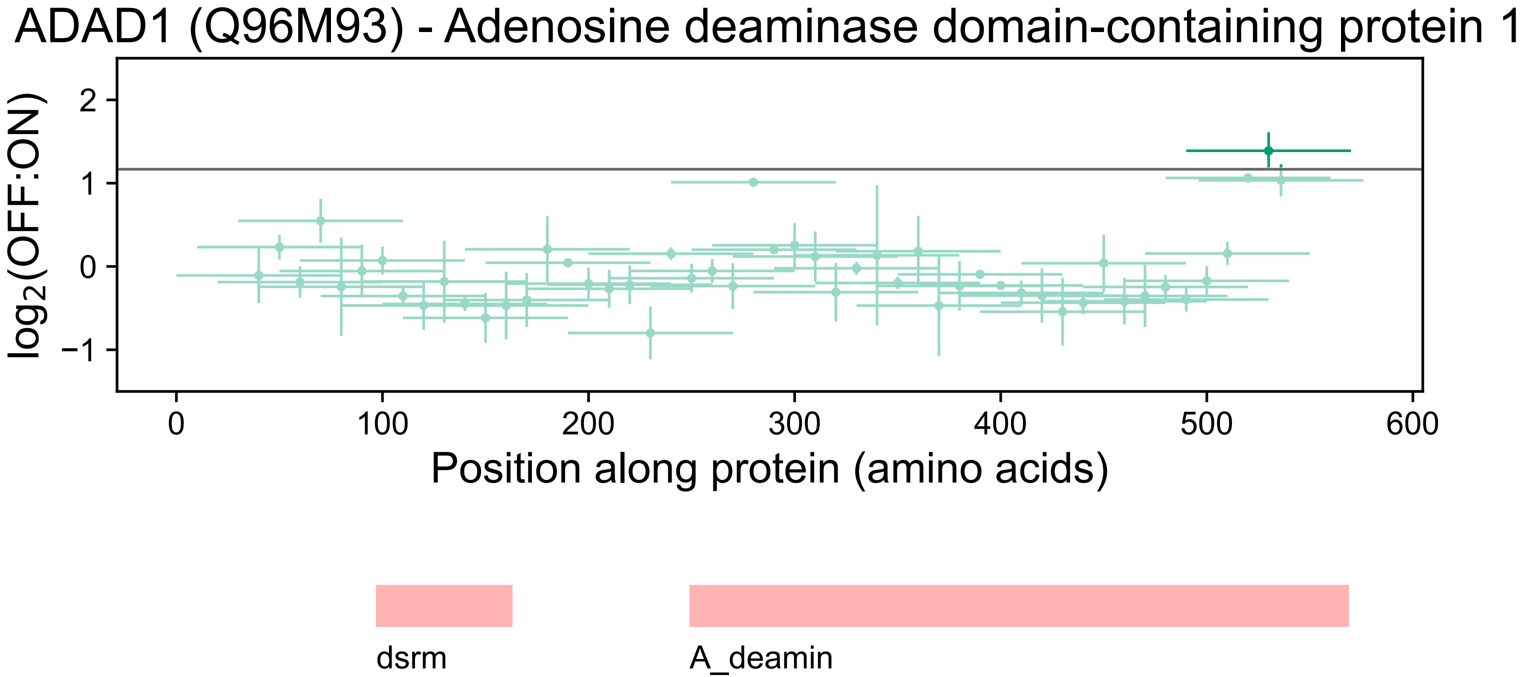

ADAD1 (Q96M93) - Adenosine deaminase domain-containing protein 1
log2(OFF:ON)
2
1
0
−1
0
100
200
300
400
500
600
Position along protein (amino acids)
dsrm
A_deamin

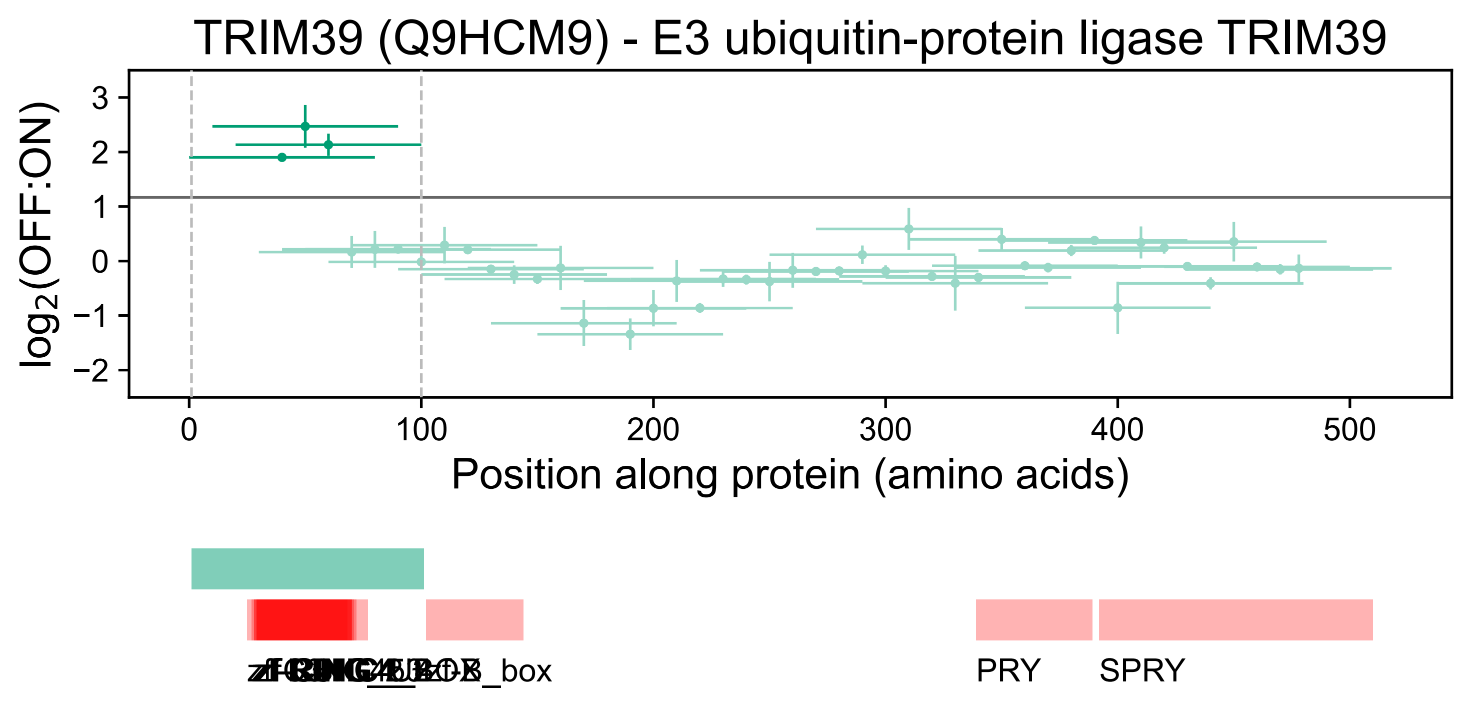
TRIM39 (Q9HCM9) - E3 ubiquitin-protein ligase TRIM39
log2(OFF:ON)
Position along protein (amino acids)
PRY
SPRY

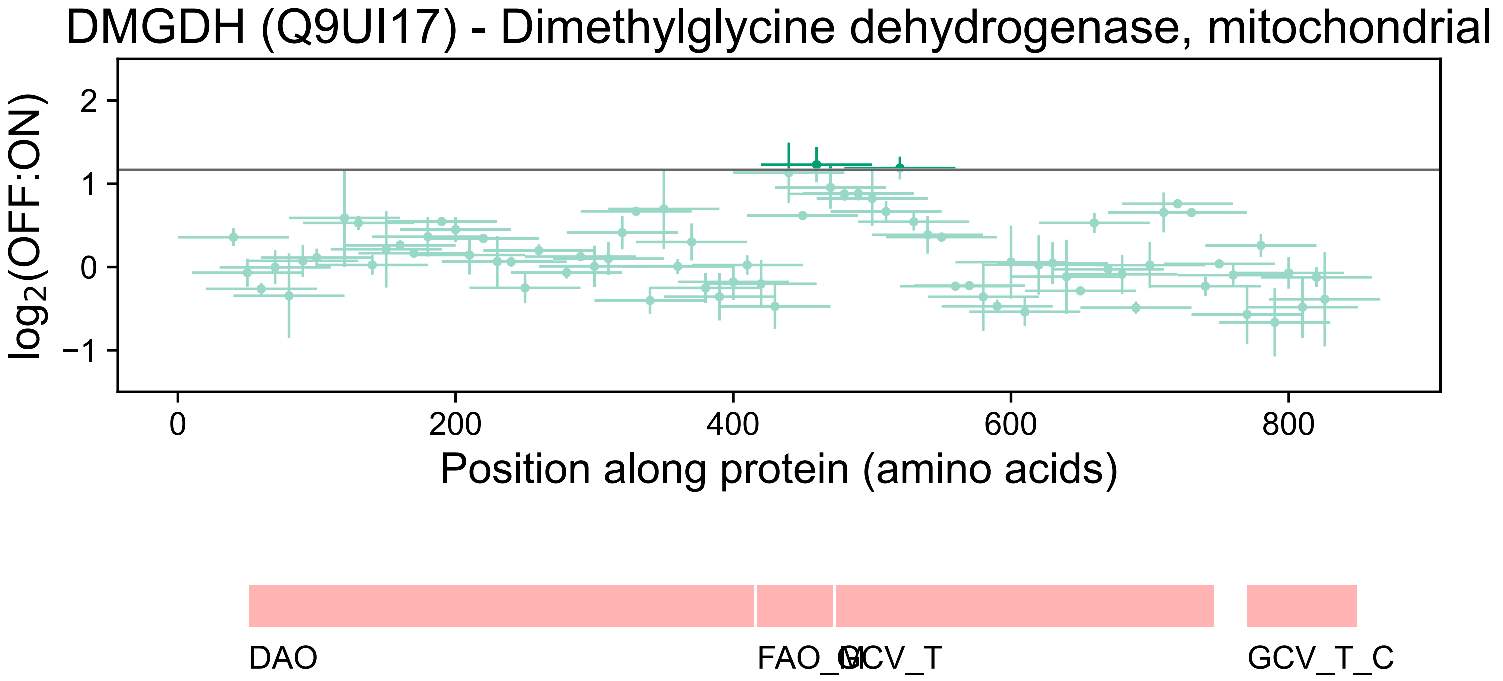
DMGDH (Q9UI17) - Dimethylglycine dehydrogenase, mitochondrial
log2(OFF:ON)
2
1
0
−1
0
200
400
600
800
Position along protein (amino acids)
DAO
FAO_M
GCV_T
GCV_T_C

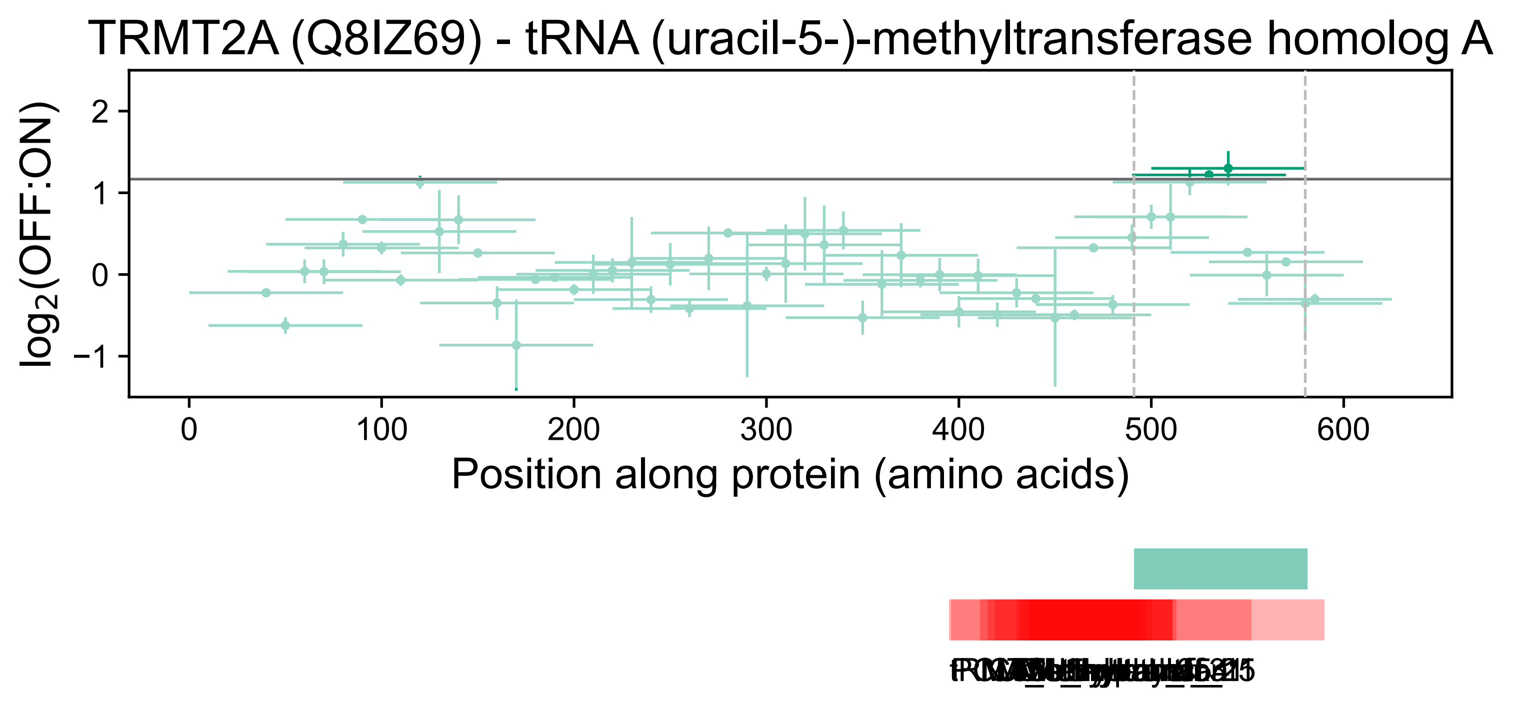
TRMT2A (Q8IZ69) - tRNA (uracil-5-)-methyltransferase homolog A
$\log_2$(OFF:ON)
2
1
0
−1
0
100
200
300
400
500
600
Position along protein (amino acids)

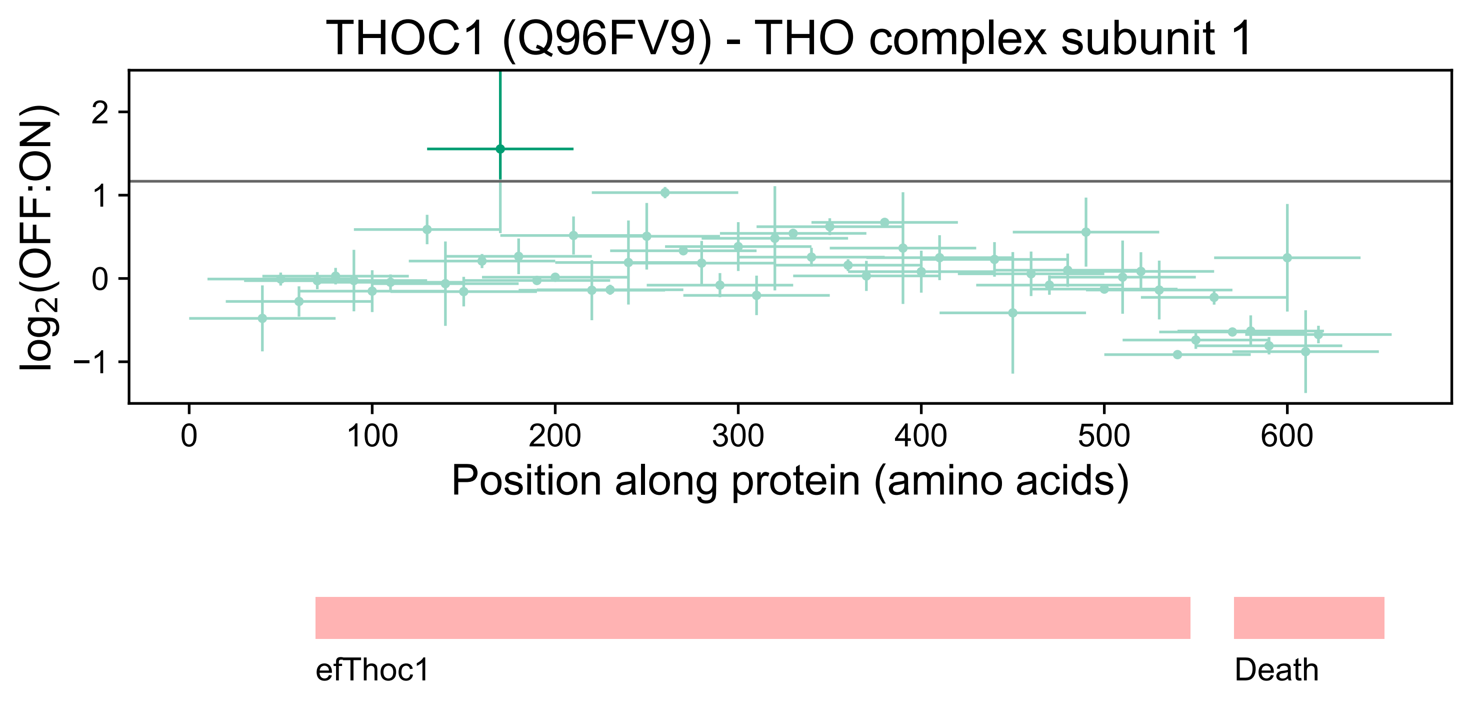
THOC1 (Q96FV9) - THO complex subunit 1
log2(OFF:ON)
2
1
0
−1
0
100
200
300
400
500
600
Position along protein (amino acids)
efThoc1
Death

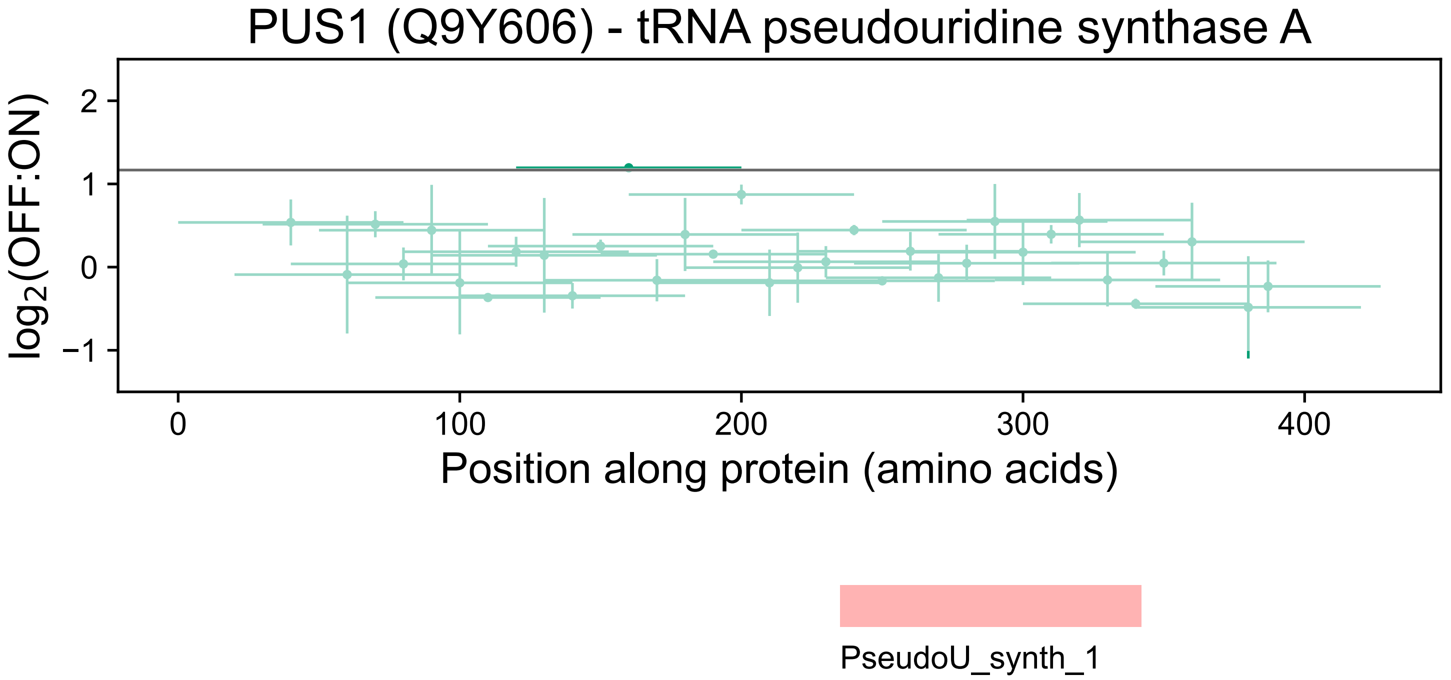
PUS1 (Q9Y606) - tRNA pseudouridine synthase A
log2(OFF:ON)
2
1
0
−1
0
100
200
300
400
Position along protein (amino acids)
PseudoU_synth_1

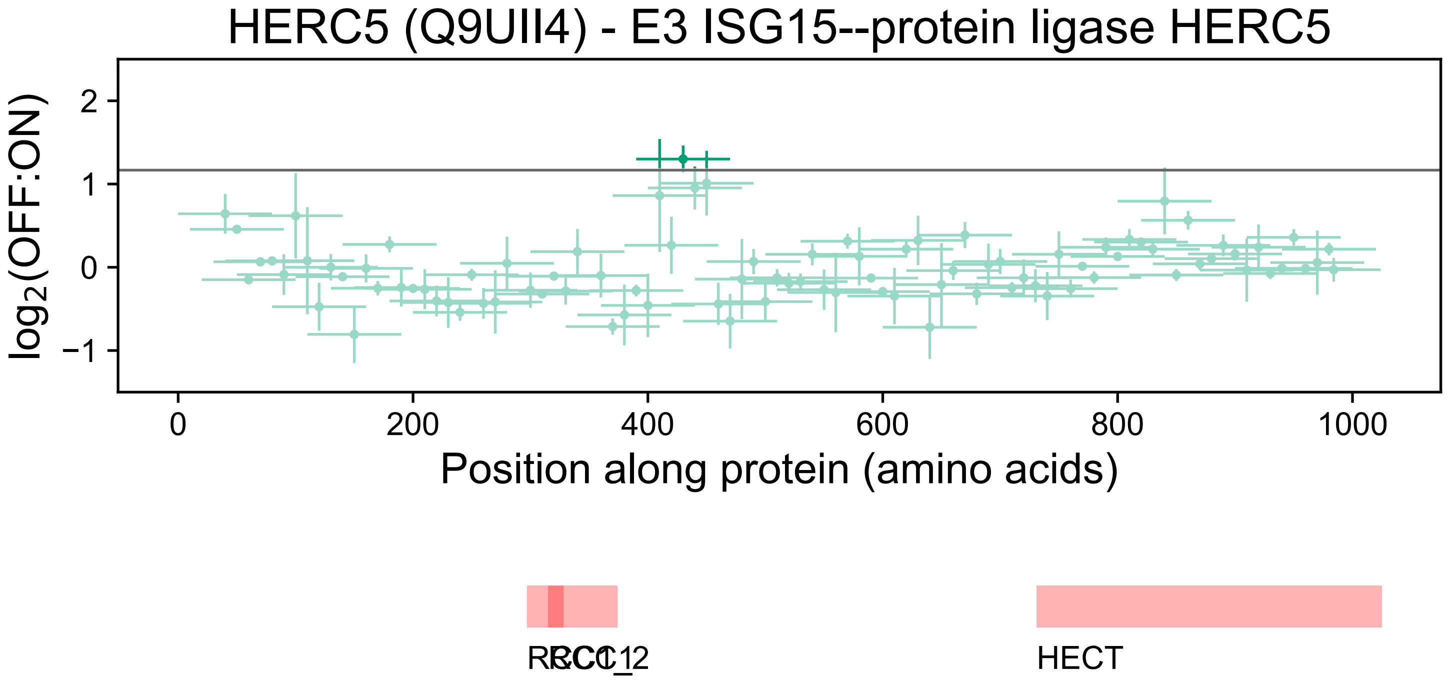

HERC5 (Q9UII4) - E3 ISG15--protein ligase HERC5
log2(OFF:ON)
2
1
0
−1
0
200
400
600
800
1000
Position along protein (amino acids)
RCC1_2
RCC1
HECT

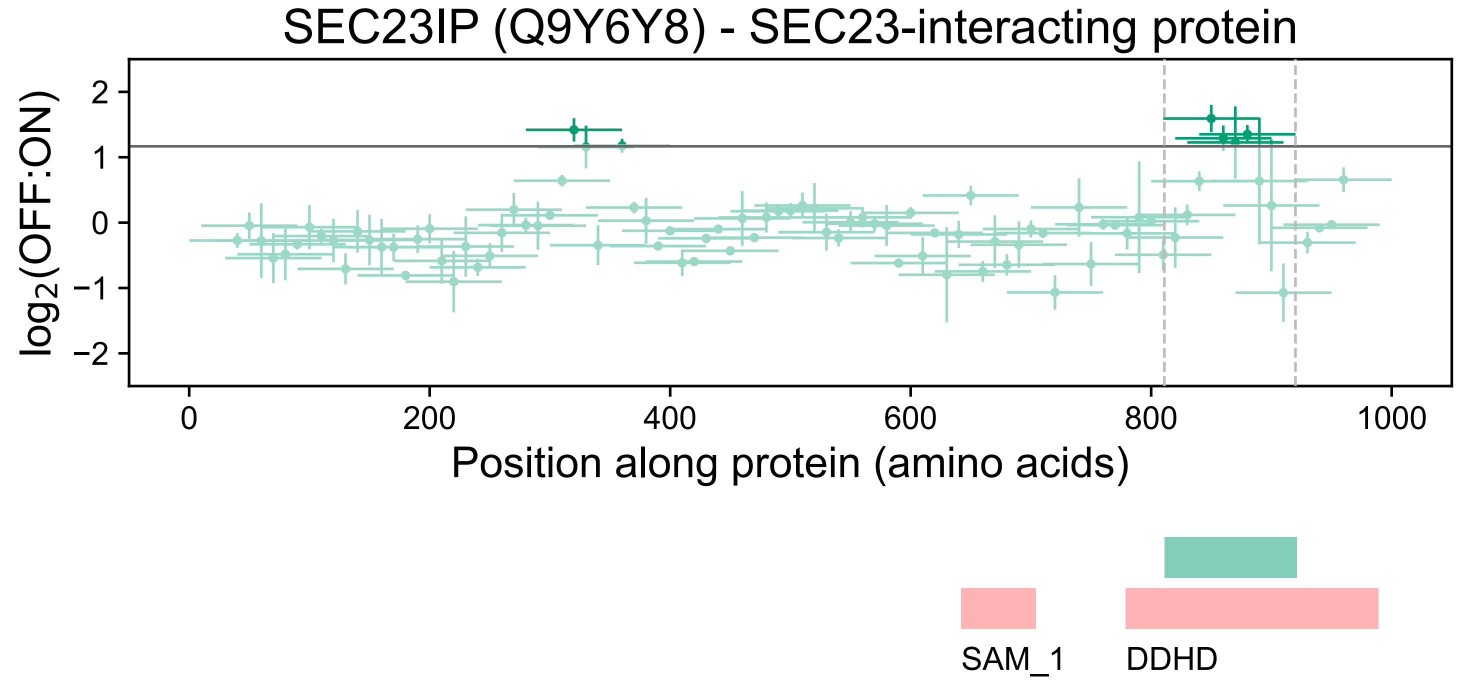

SEC23IP (Q9Y6Y8) - SEC23-interacting protein
log2(OFF:ON)
2
1
0
−1
−2
0
200
400
600
800
1000
Position along protein (amino acids)
SAM_1
DDHD

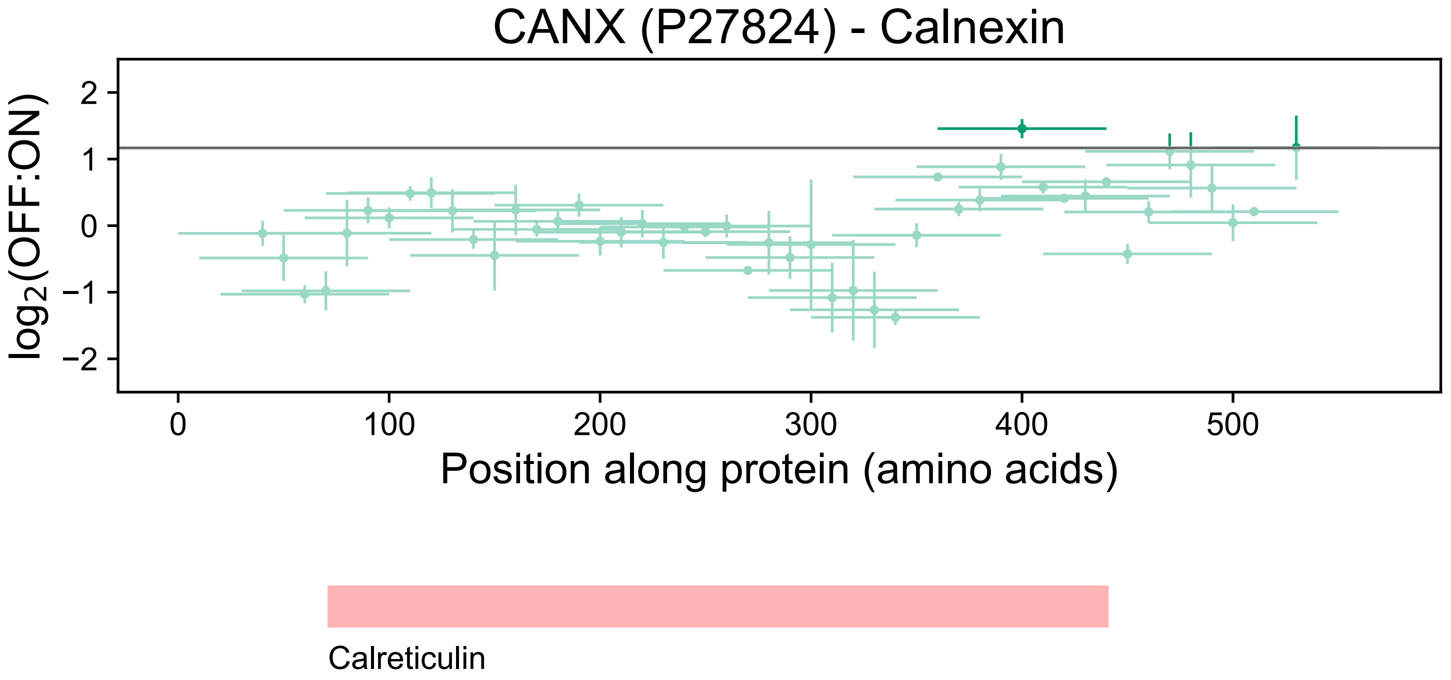

CANX (P27824) - Calnexin
log2(OFF:ON)
2
1
0
−1
−2
0
100
200
300
400
500
Position along protein (amino acids)
Calreticulin

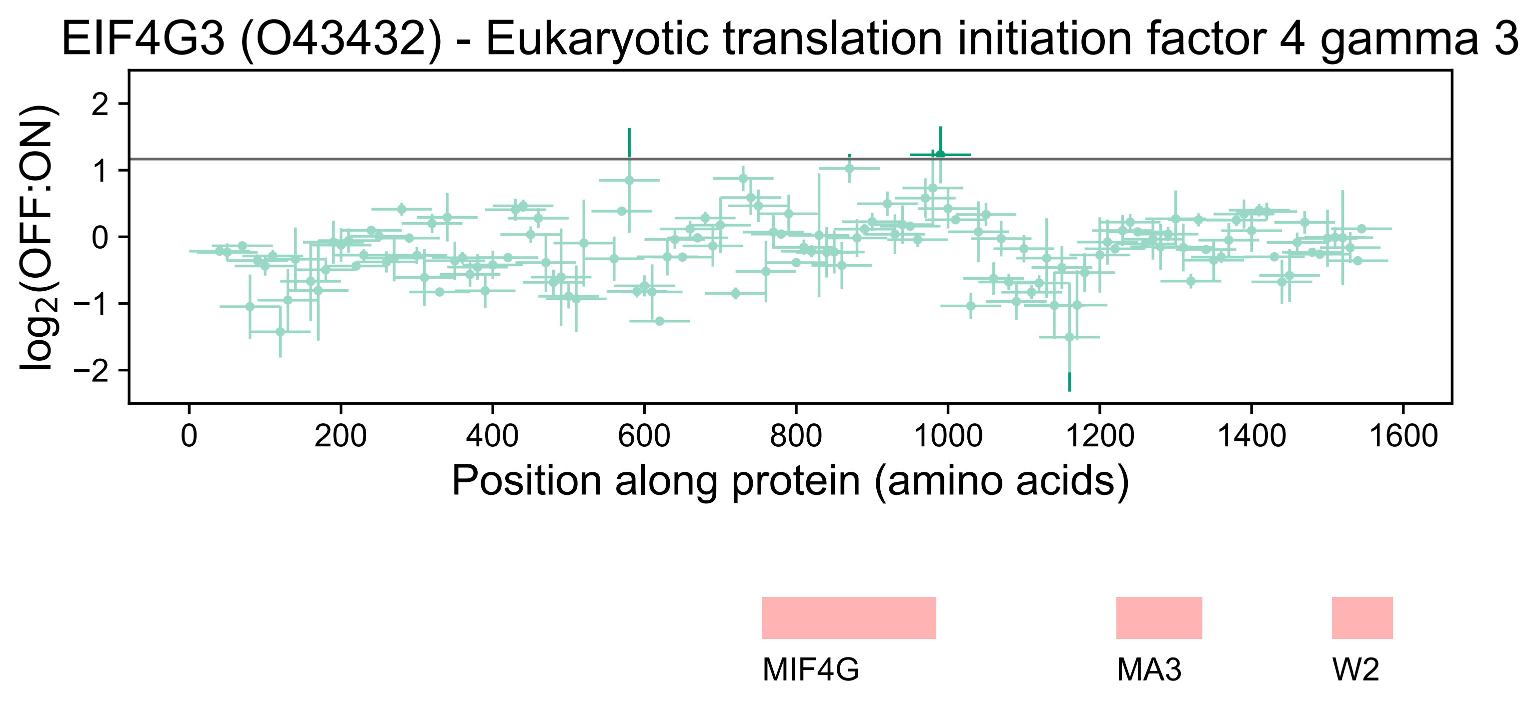
EIF4G3 (O43432) - Eukaryotic translation initiation factor 4 gamma 3
log2(OFF:ON)
2
1
0
−1
−2
0
200
400
600
800
1000
1200
1400
1600
Position along protein (amino acids)
MIF4G
MA3
W2

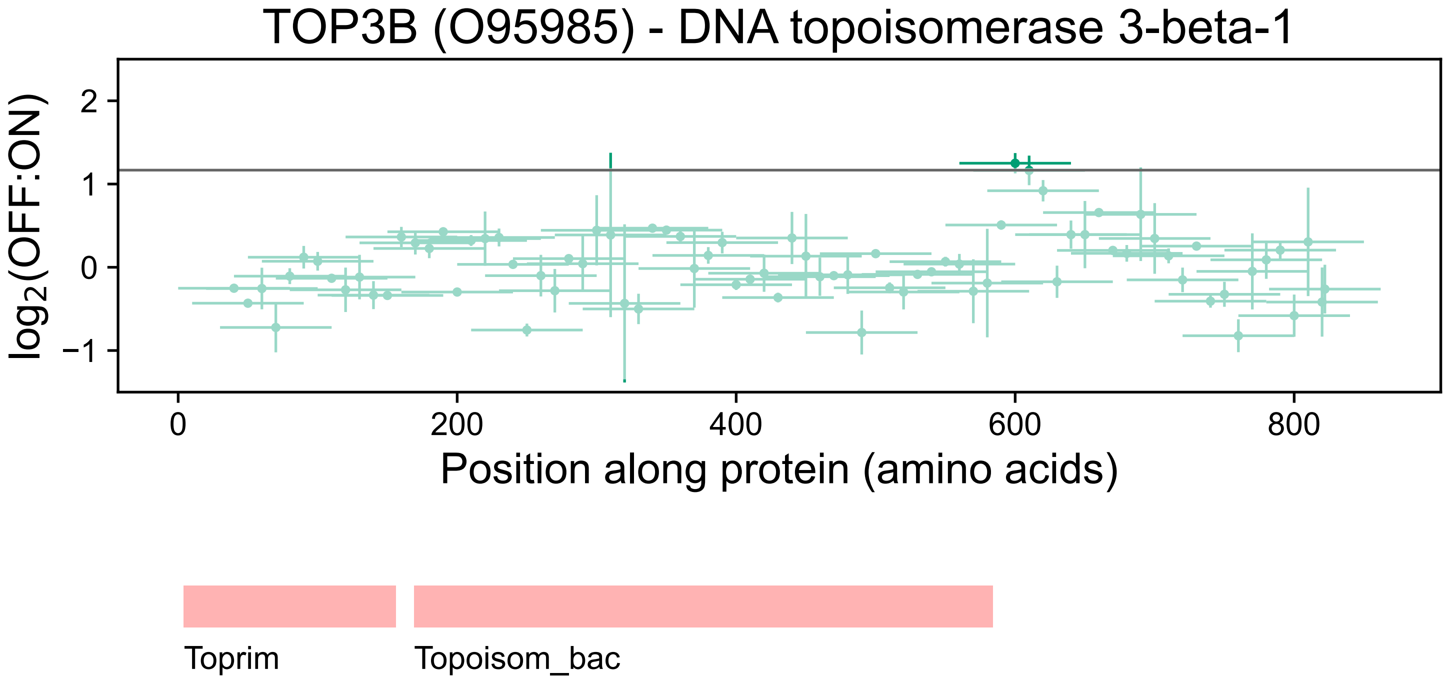
TOP3B (O95985) - DNA topoisomerase 3-beta-1
log2(OFF:ON)
2
1
0
−1
0
200
400
600
800
Position along protein (amino acids)
Toprim
Topoisom_bac

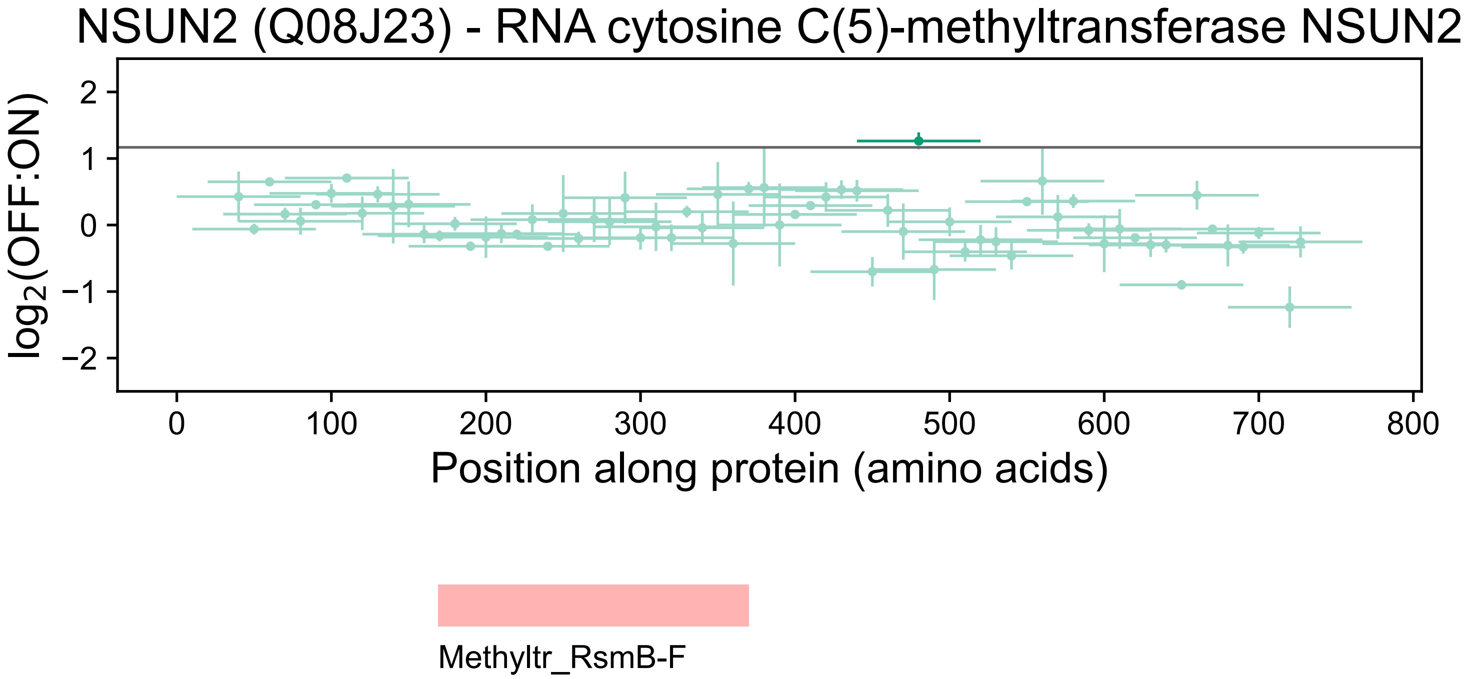
NSUN2 (Q08J23) - RNA cytosine C(5)-methyltransferase NSUN2
log2(OFF:ON)
2
1
0
−1
−2
0
100
200
300
400
500
600
700
800
Position along protein (amino acids)
Methyltr_RsmB-F

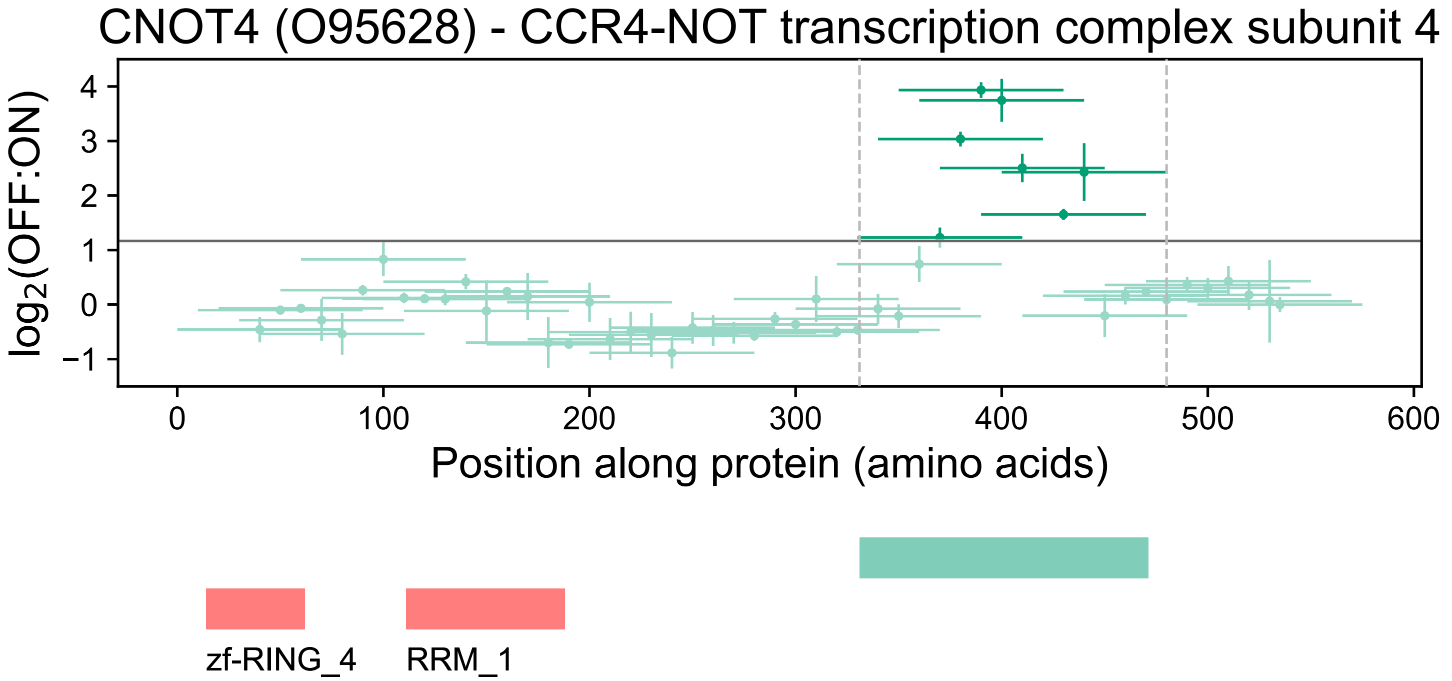
CNOT4 (O95628) - CCR4-NOT transcription complex subunit 4
log2(OFF:ON)
4
3
2
1
0
−1
0
100
200
300
400
500
600
Position along protein (amino acids)
zf-RING_4
RRM_1

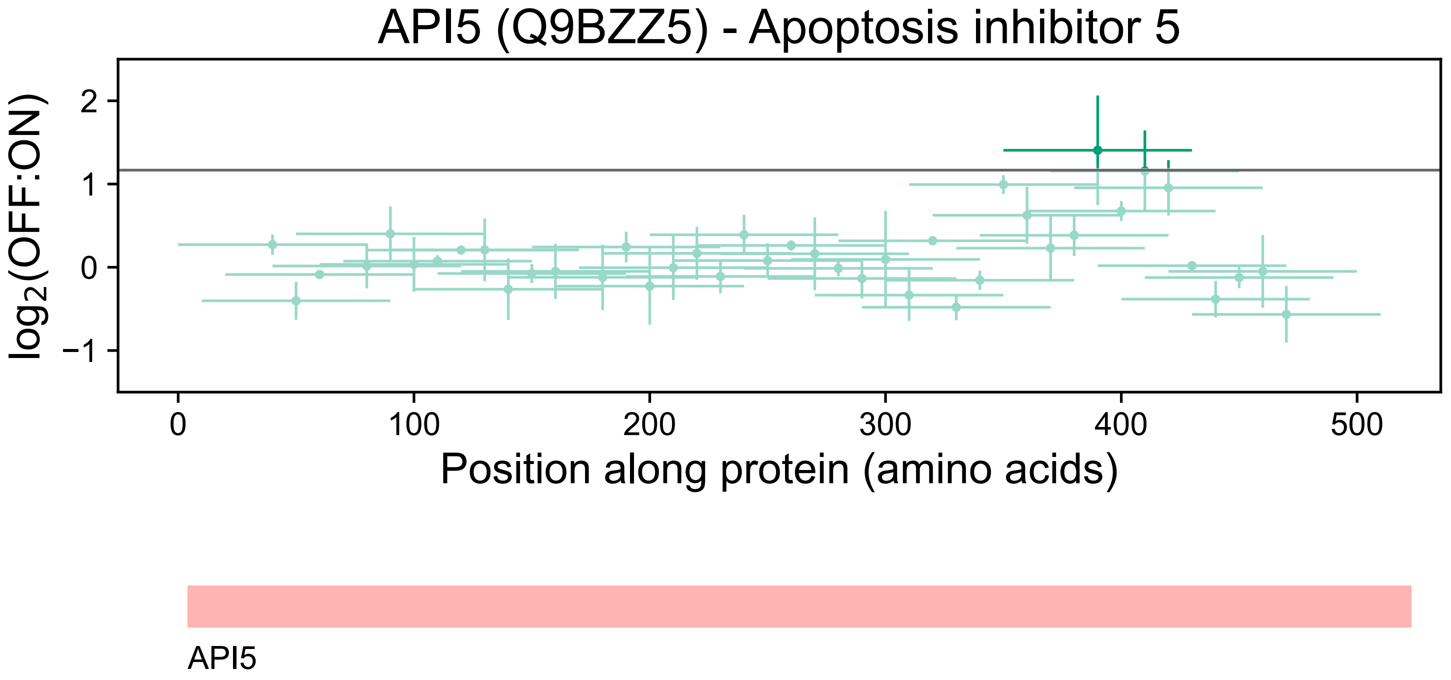

API5 (Q9BZZ5) - Apoptosis inhibitor 5
log2(OFF:ON)
2
1
0
−1
0
100
200
300
400
500
Position along protein (amino acids)
API5

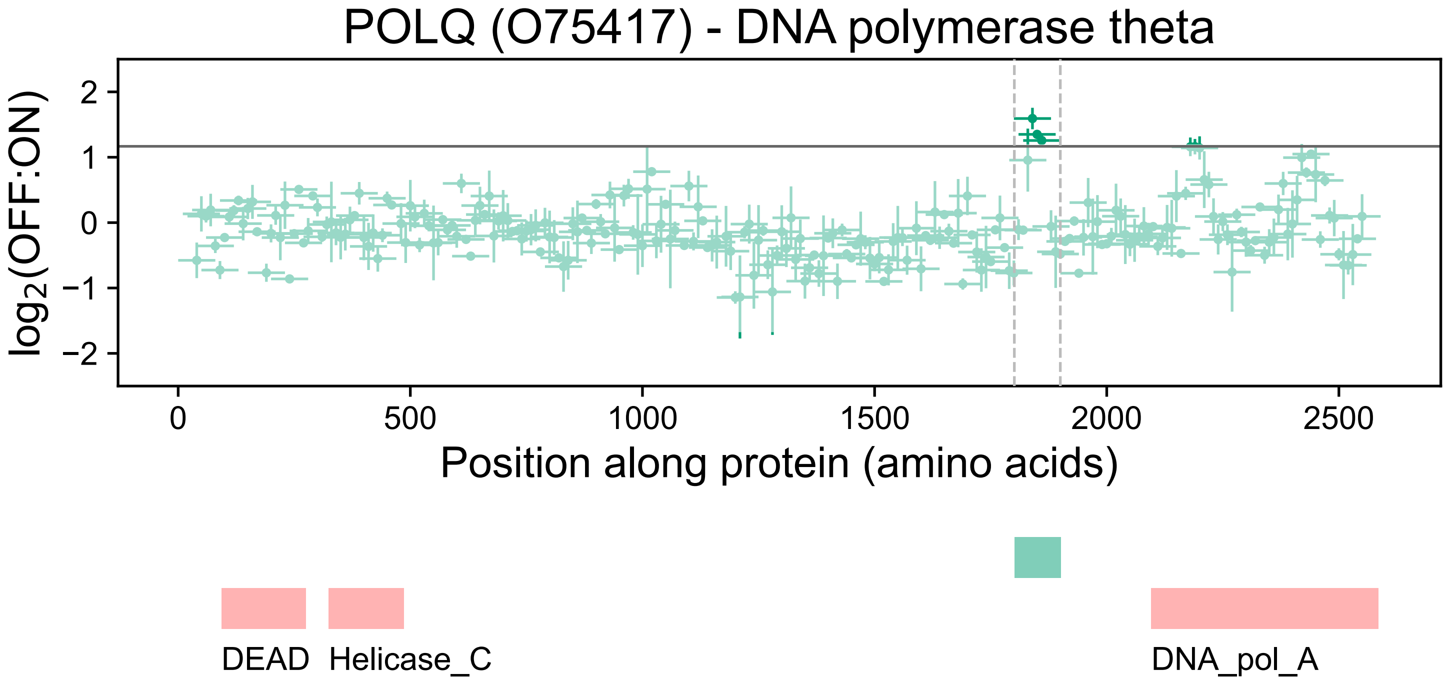
POLQ (O75417) - DNA polymerase theta
log2(OFF:ON)
2
1
0
−1
−2
0
500
1000
1500
2000
2500
Position along protein (amino acids)
DEAD
Helicase_C
DNA_pol_A

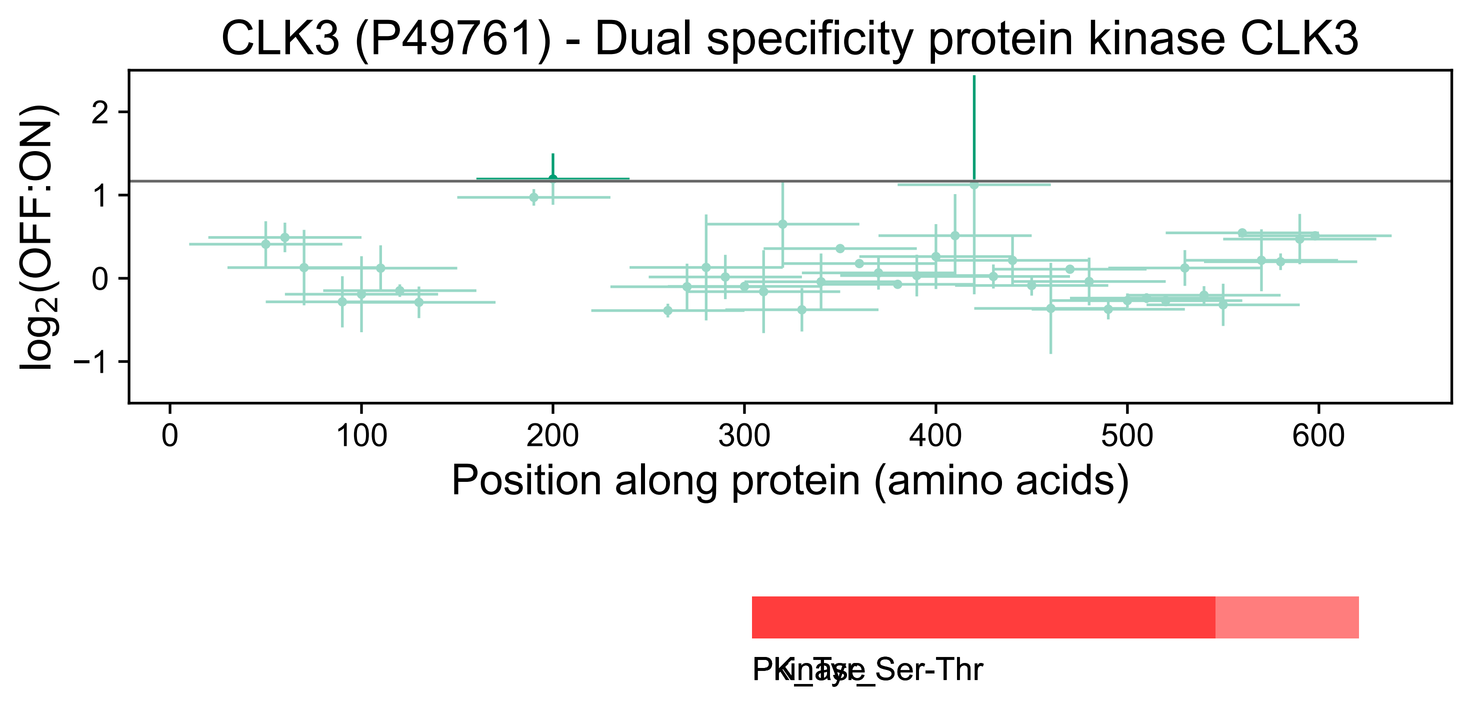
CLK3 (P49761) - Dual specificity protein kinase CLK3
log2(OFF:ON)
2
1
0
−1
0
100
200
300
400
500
600
Position along protein (amino acids)

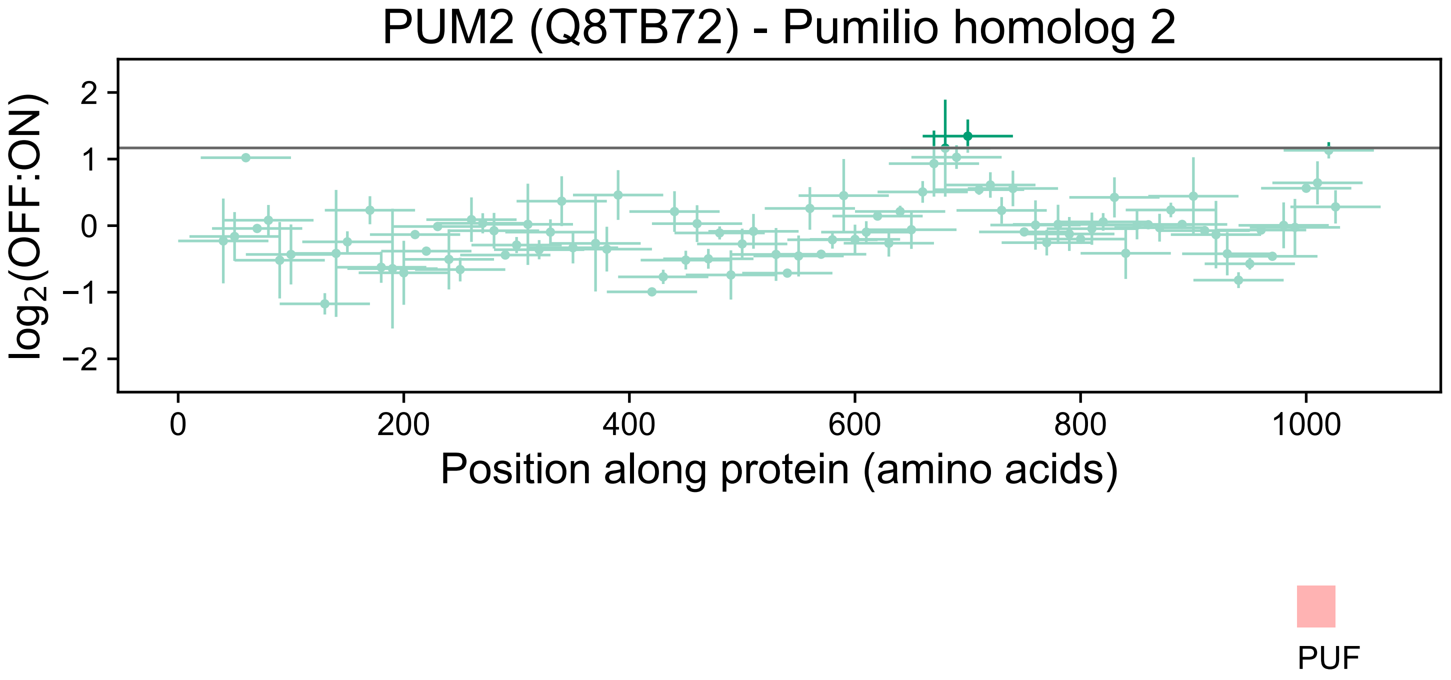

PUM2 (Q8TB72) - Pumilio homolog 2
log2(OFF:ON)
2
1
0
−1
−2
0
200
400
600
800
1000
Position along protein (amino acids)
PUF

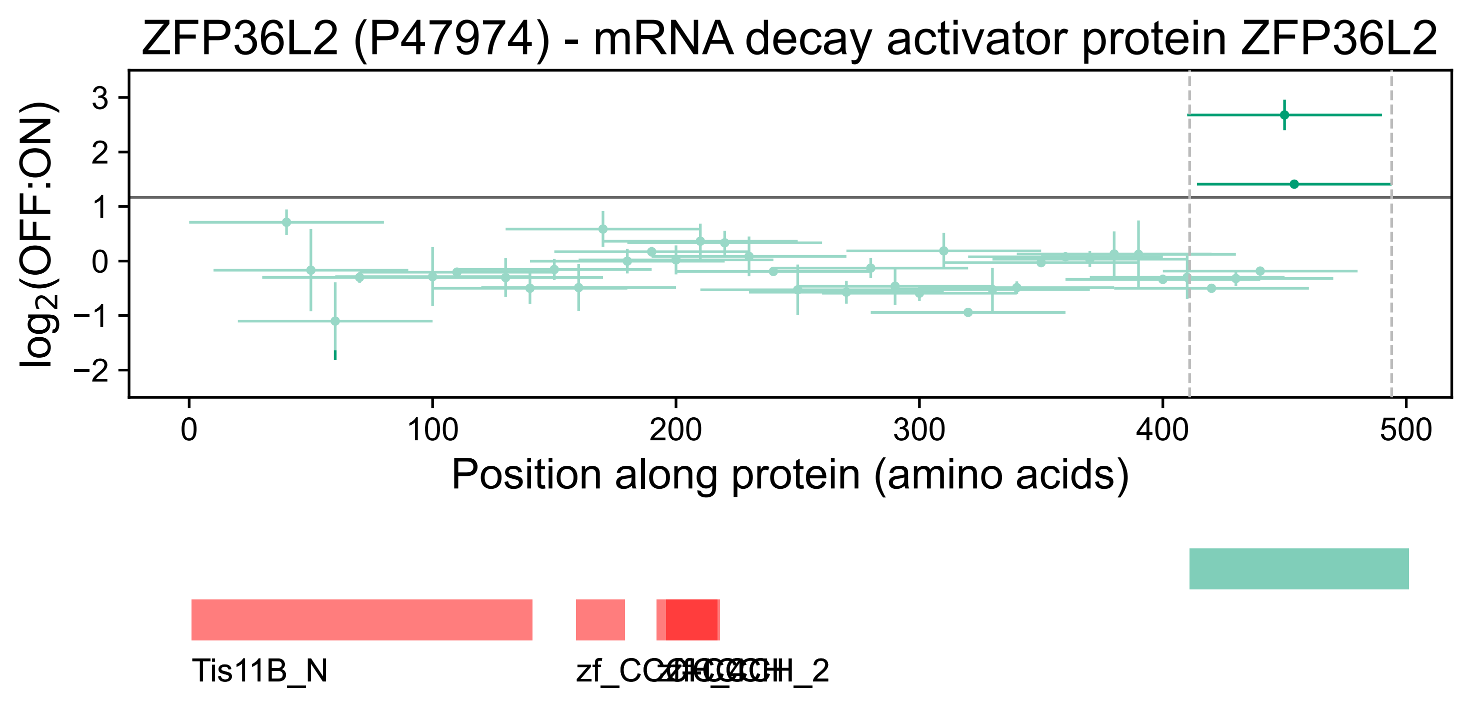

ZFP36L2 (P47974) - mRNA decay activator protein ZFP36L2
log2(OFF:ON)
3
2
1
0
−1
−2
0
100
200
300
400
500
Position along protein (amino acids)
Tis11B_N
zf_CCCH
zf-CCCH_2

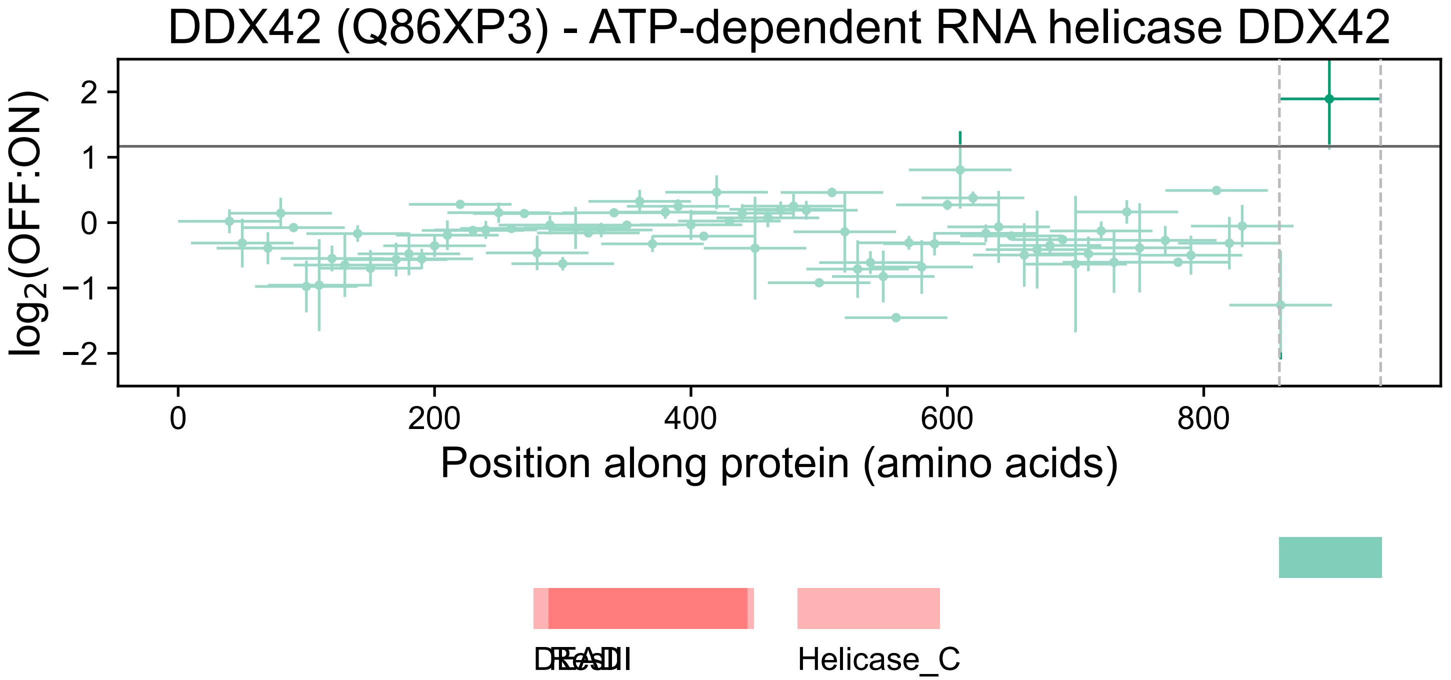

DDX42 (Q86XP3) - ATP-dependent RNA helicase DDX42
log2(OFF:ON)
2
1
0
−1
−2
0
200
400
600
800
Position along protein (amino acids)
DEAD
ResIII
Helicase_C

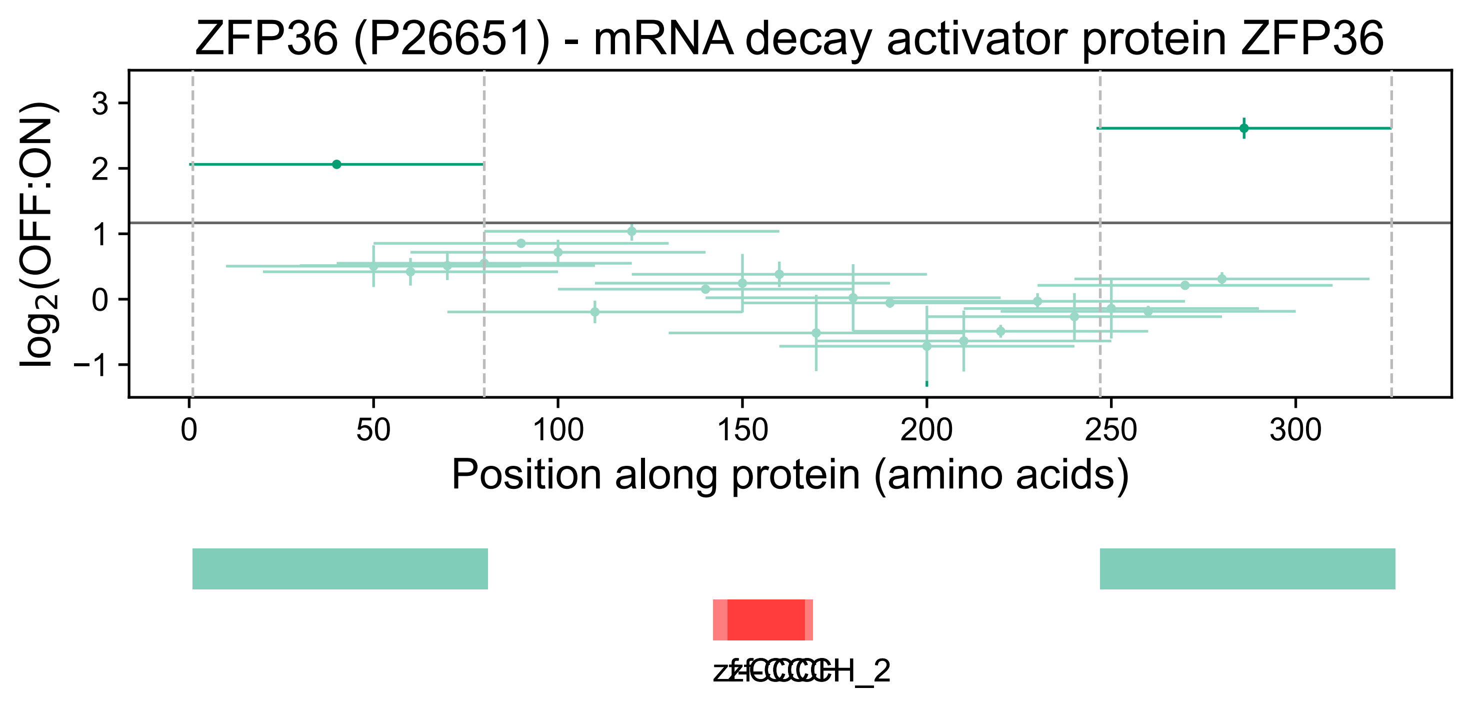

ZFP36 (P26651) - mRNA decay activator protein ZFP36
log2(OFF:ON)
3
2
1
0
−1
0
50
100
150
200
250
300
Position along protein (amino acids)
zf-CCCH
zf-CCCH_2

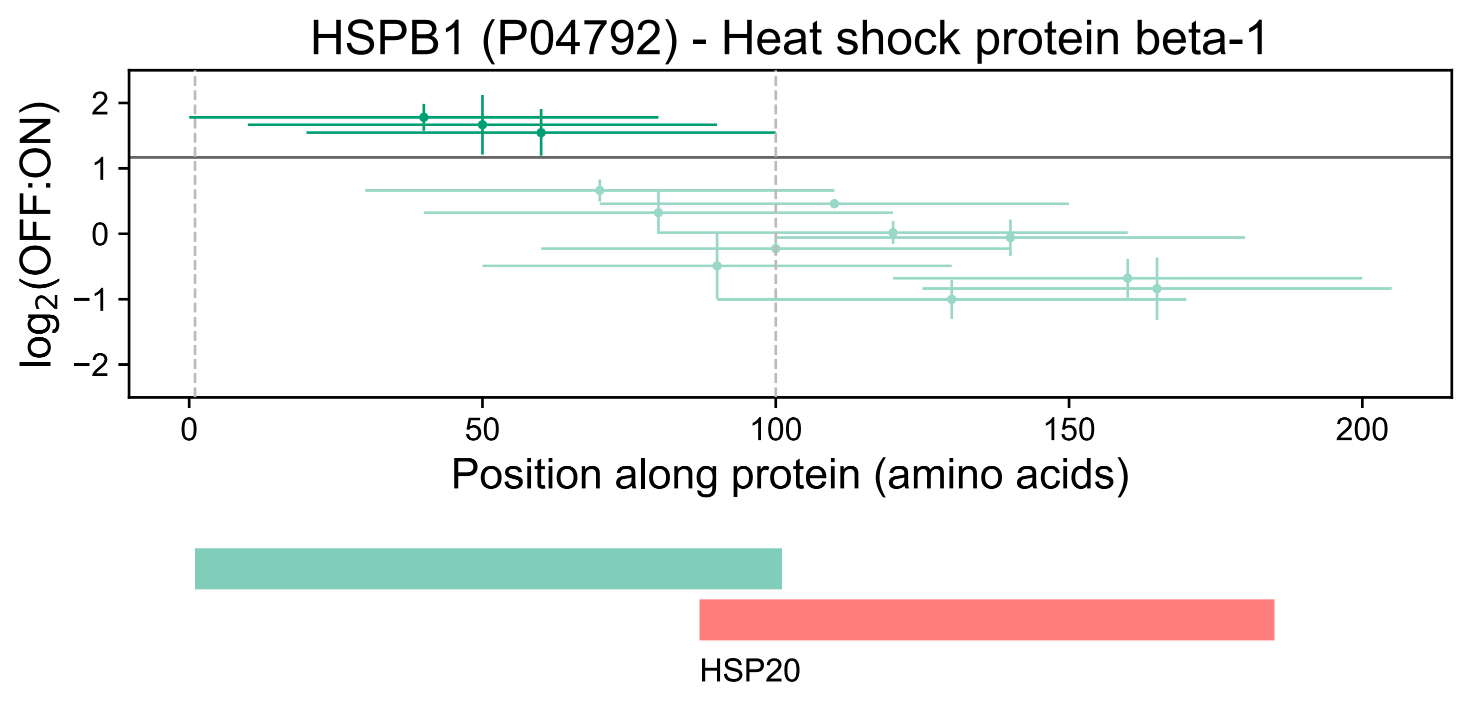
HSPB1 (P04792) - Heat shock protein beta-1
log2(OFF:ON)
2
1
0
−1
−2
0
50
100
150
200
Position along protein (amino acids)
HSP20

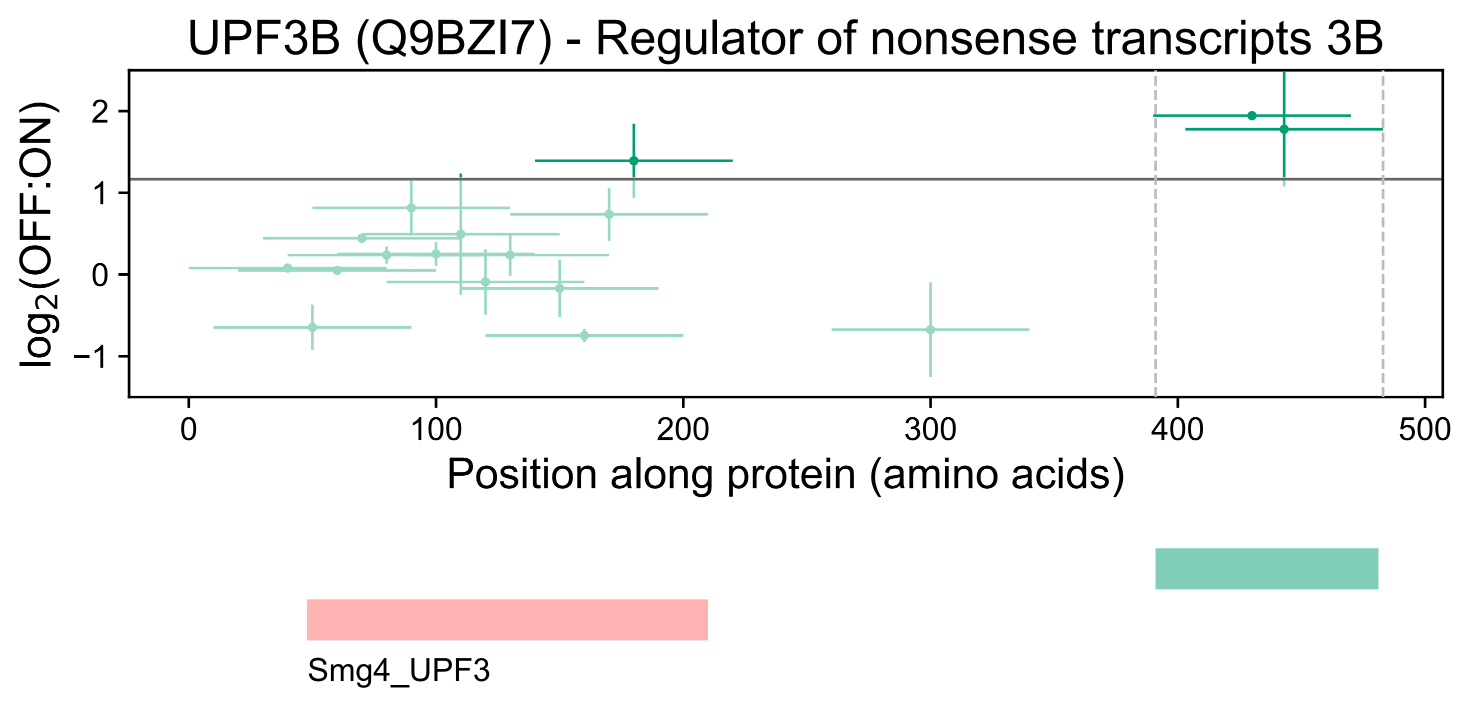
UPF3B (Q9BZI7) - Regulator of nonsense transcripts 3B
log2(OFF:ON)
2
1
0
−1
0
100
200
300
400
500
Position along protein (amino acids)
Smg4_UPF3

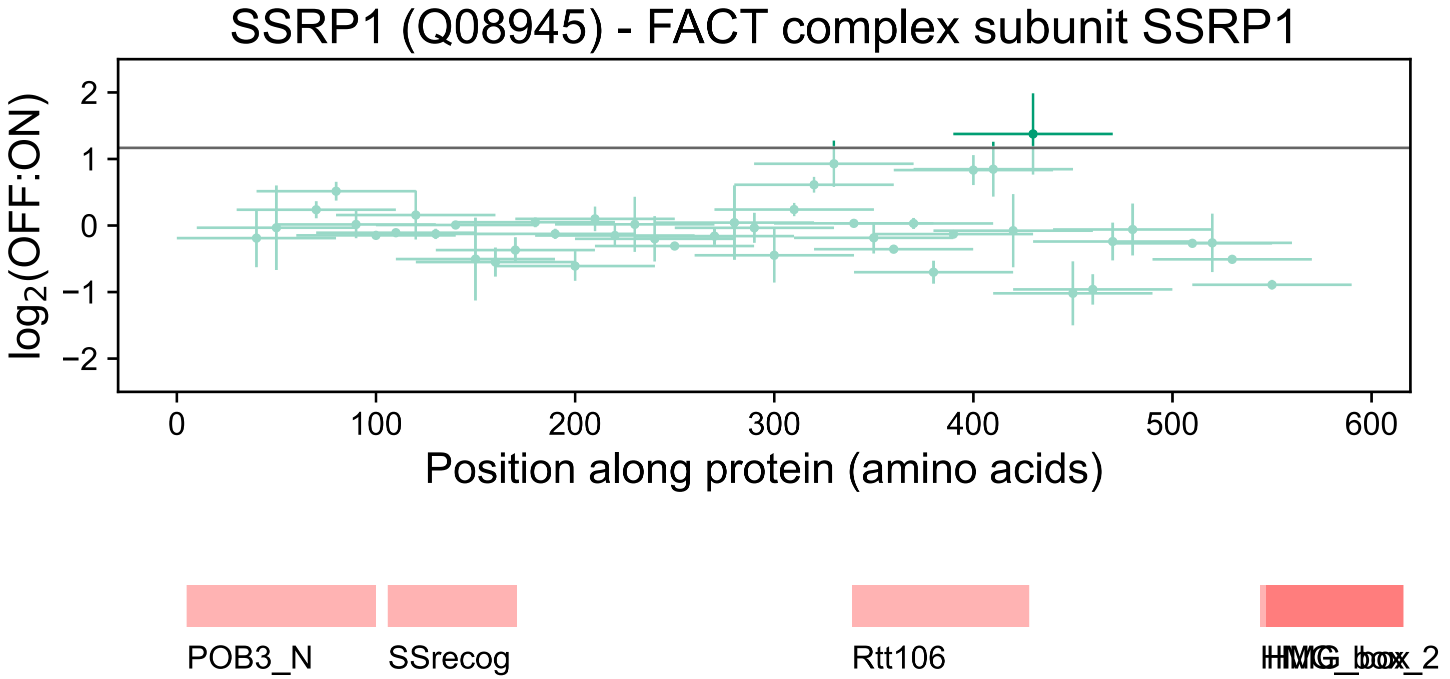

SSRP1 (Q08945) - FACT complex subunit SSRP1
log2(OFF:ON)
2
1
0
−1
−2
0
100
200
300
400
500
600
Position along protein (amino acids)
POB3_N
SSrecog
Rtt106
HMG_box
HMG_box_2

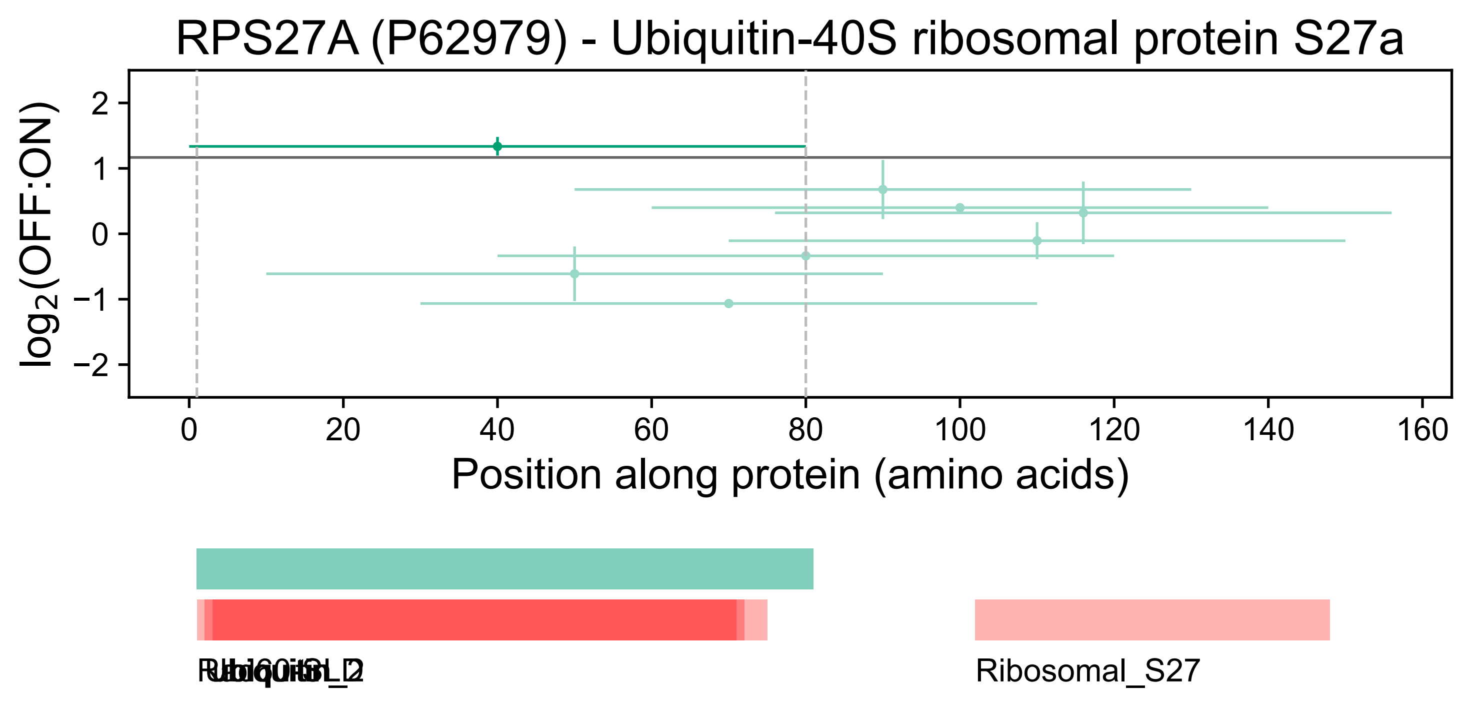

RPS27A (P62979) - Ubiquitin-40S ribosomal protein S27a
log2(OFF:ON)
2
1
0
−1
−2
0
20
40
60
80
100
120
140
160
Position along protein (amino acids)
Ribosomal_S27

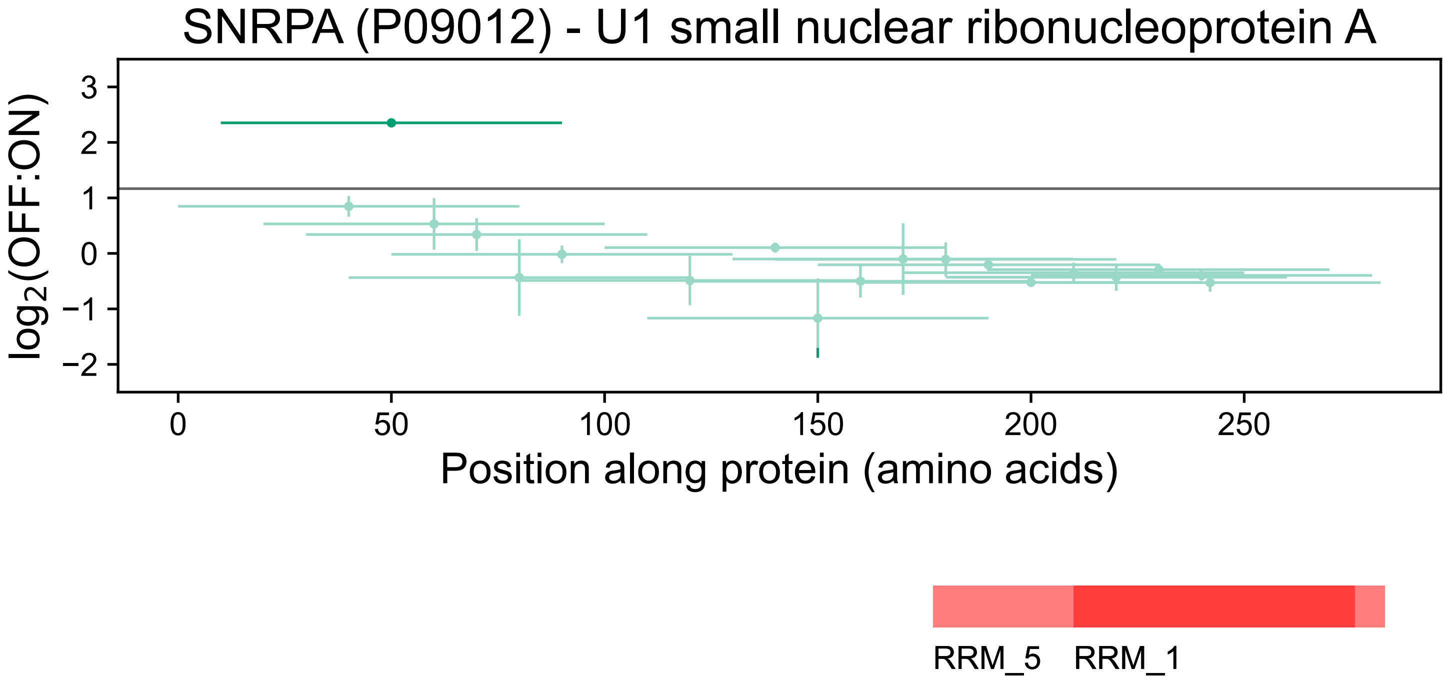
SNRPA (P09012) - U1 small nuclear ribonucleoprotein A
log2(OFF:ON)
3
2
1
0
−1
−2
0
50
100
150
200
250
Position along protein (amino acids)
RRM_5
RRM_1

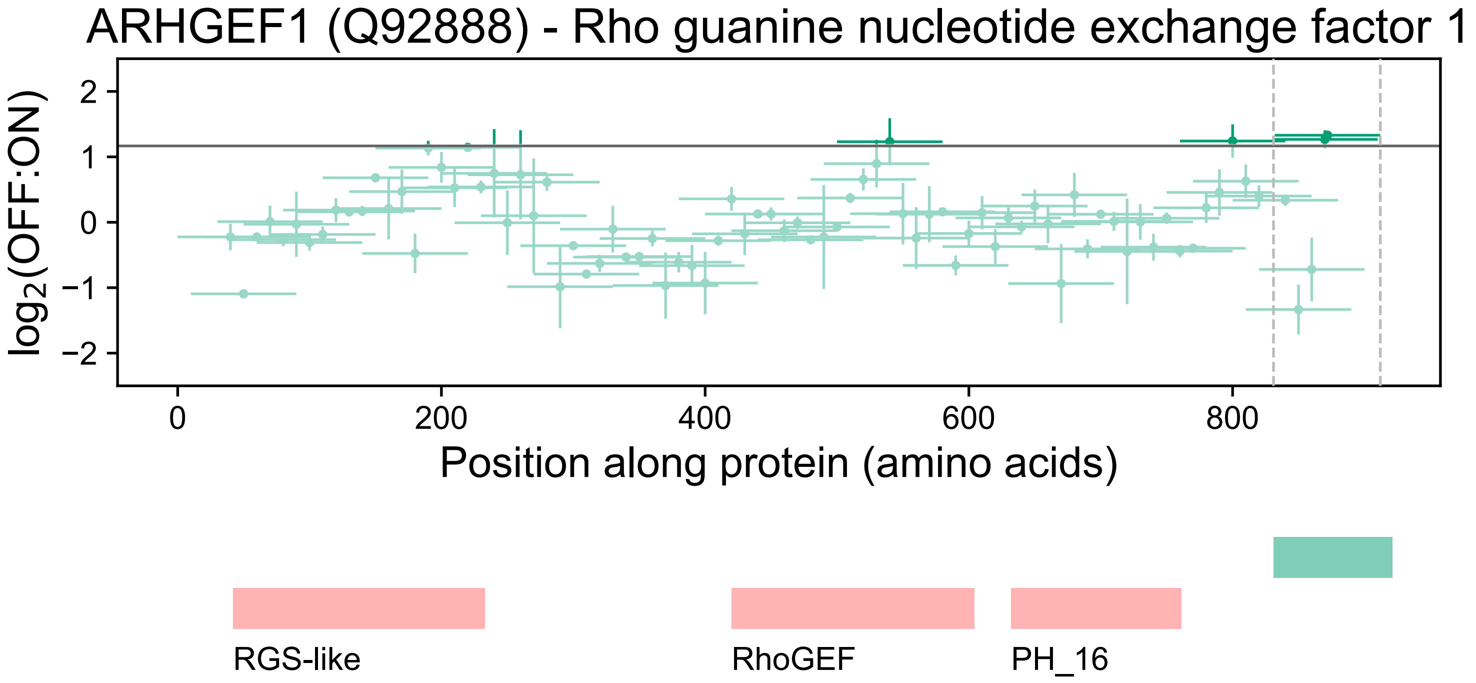

ARHGEF1 (Q92888) - Rho guanine nucleotide exchange factor 1
log2(OFF:ON)
2
1
0
−1
−2
0
200
400
600
800
Position along protein (amino acids)
RGS-like
RhoGEF
PH_16

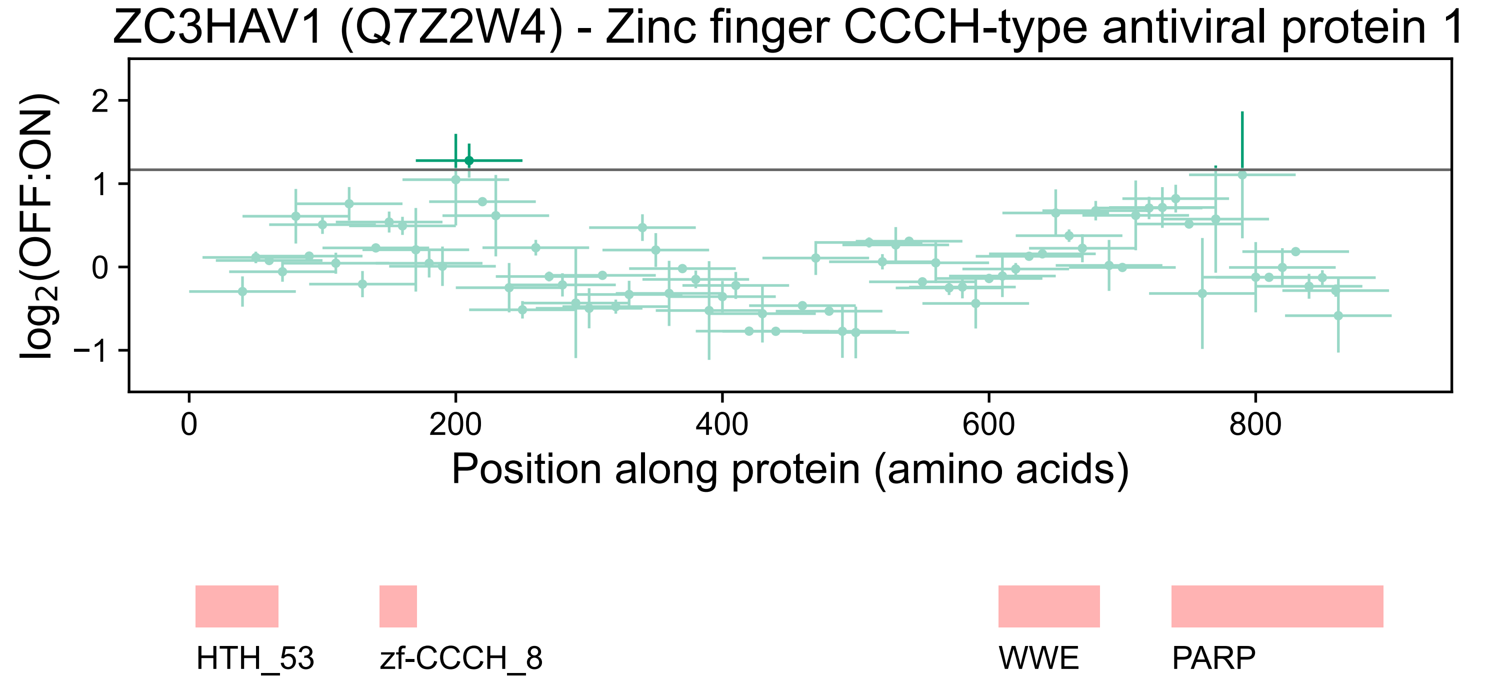

ZC3HAV1 (Q7Z2W4) - Zinc finger CCCH-type antiviral protein 1
log2(OFF:ON)
2
1
0
−1
0
200
400
600
800
Position along protein (amino acids)
HTH_53
zf-CCCH_8
WWE
PARP

CPEB4 (Q17RY0) - Cytoplasmic polyadenylation element-binding protein 4

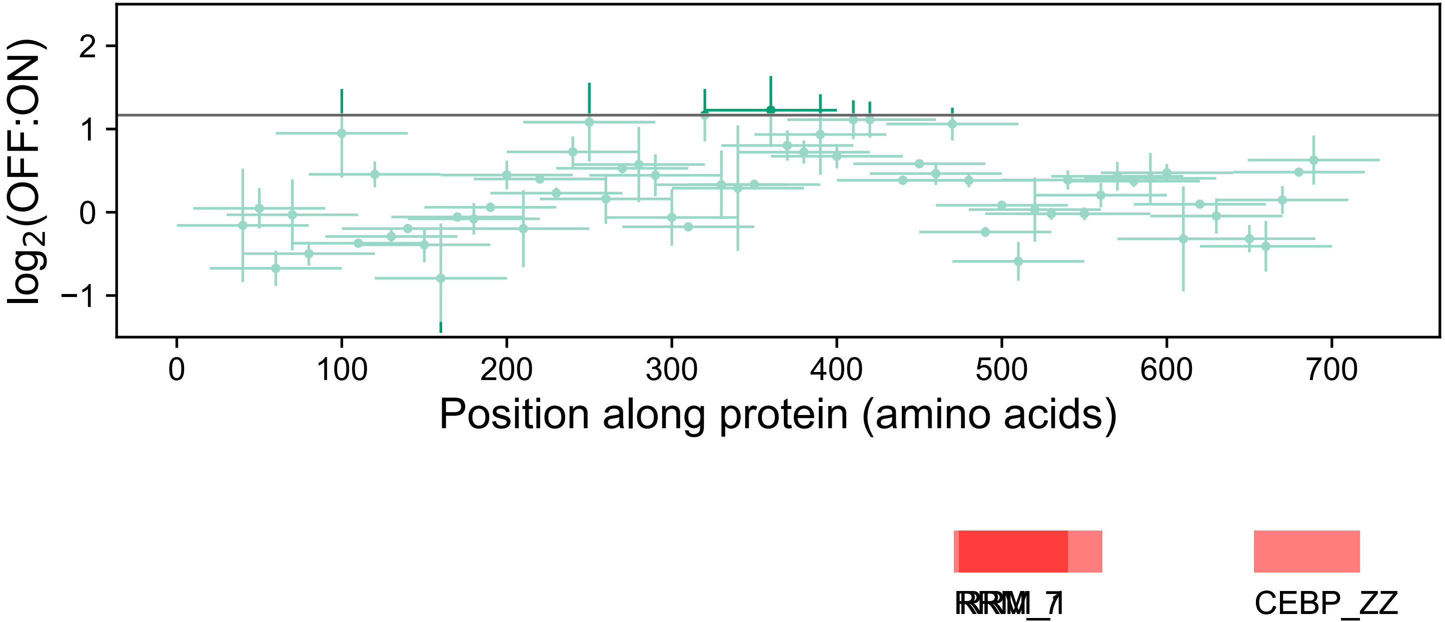

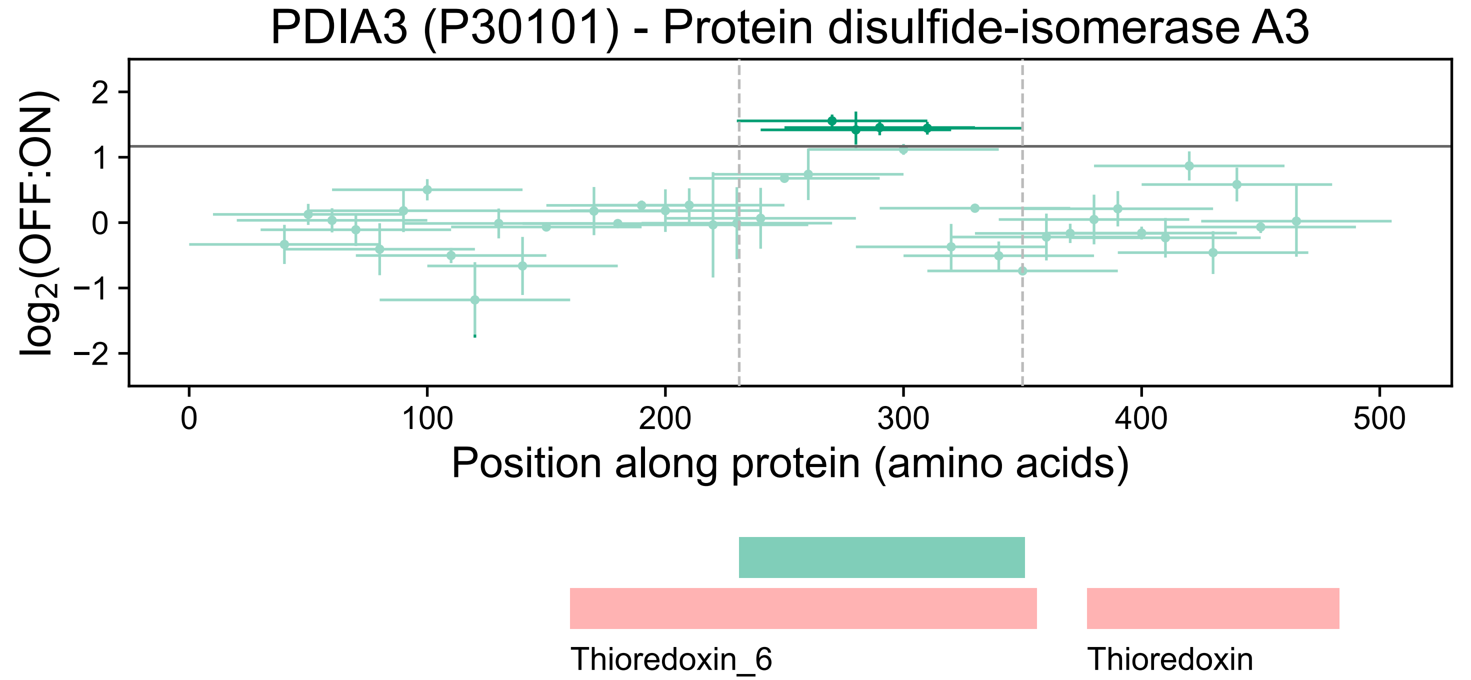
PDIA3 (P30101) - Protein disulfide-isomerase A3
log2(OFF:ON)
2
1
0
−1
−2
0
100
200
300
400
500
Position along protein (amino acids)
Thioredoxin_6
Thioredoxin

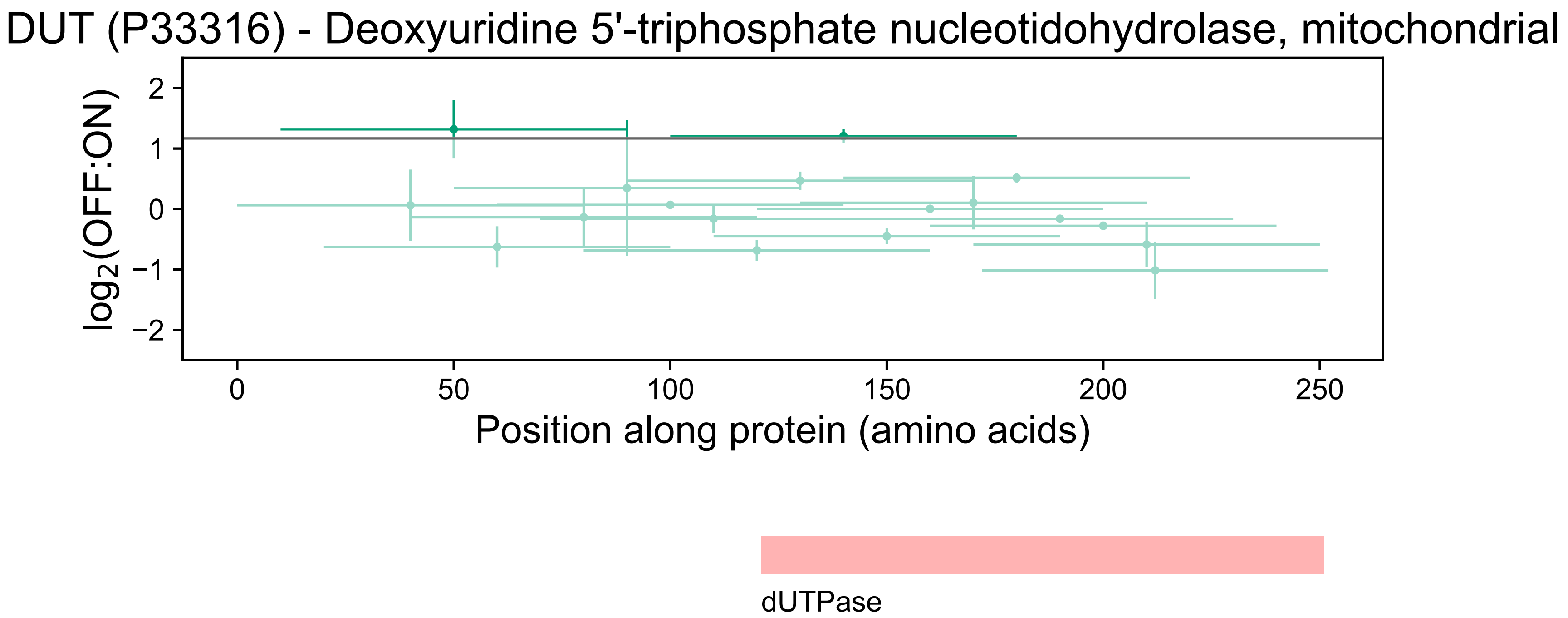
DUT (P33316) - Deoxyuridine 5'-triphosphate nucleotidohydrolase, mitochondrial
log2(OFF:ON)
2
1
0
−1
−2
0
50
100
150
200
250
Position along protein (amino acids)
dUTPase

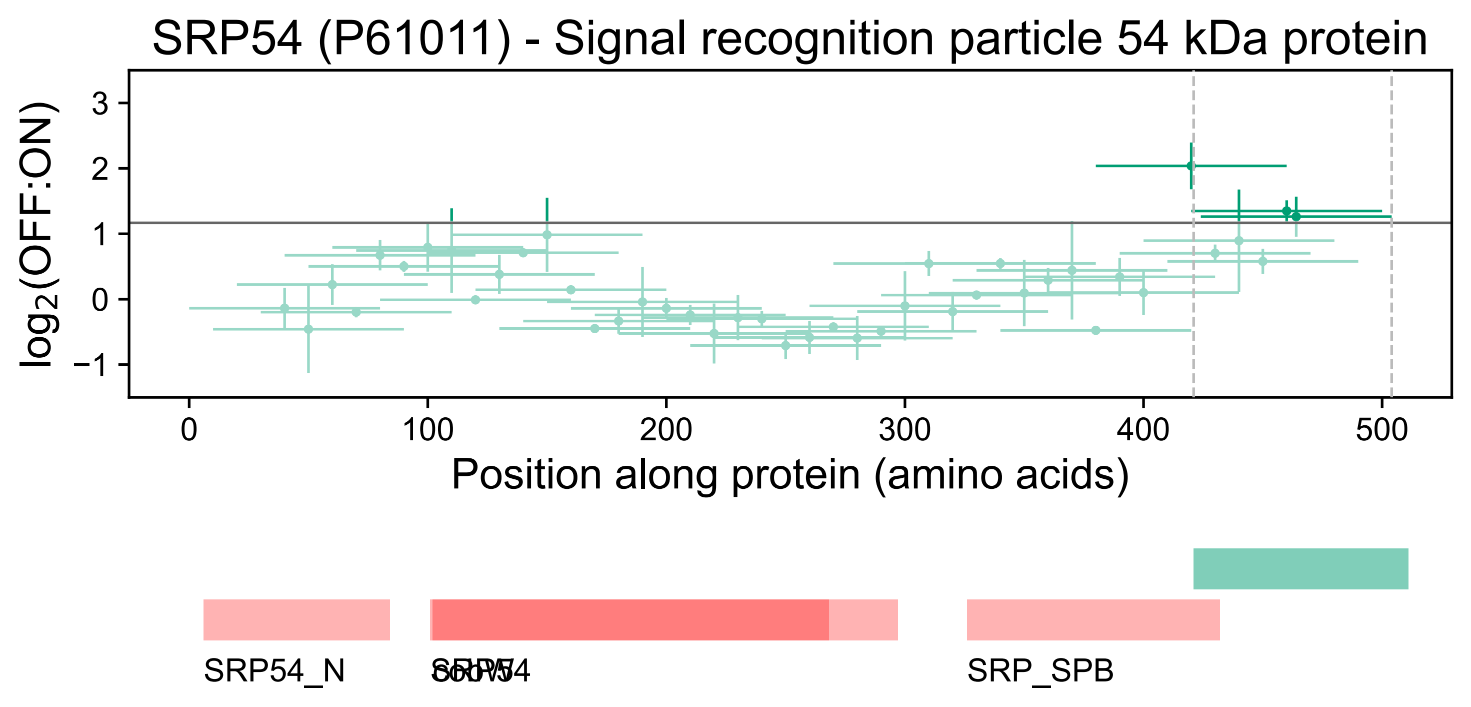
SRP54 (P61011) - Signal recognition particle 54 kDa protein
log2(OFF:ON)
3
2
1
0
−1
0
100
200
300
400
500
Position along protein (amino acids)
SRP54_N
SRP54
cobW
SRP_SPB

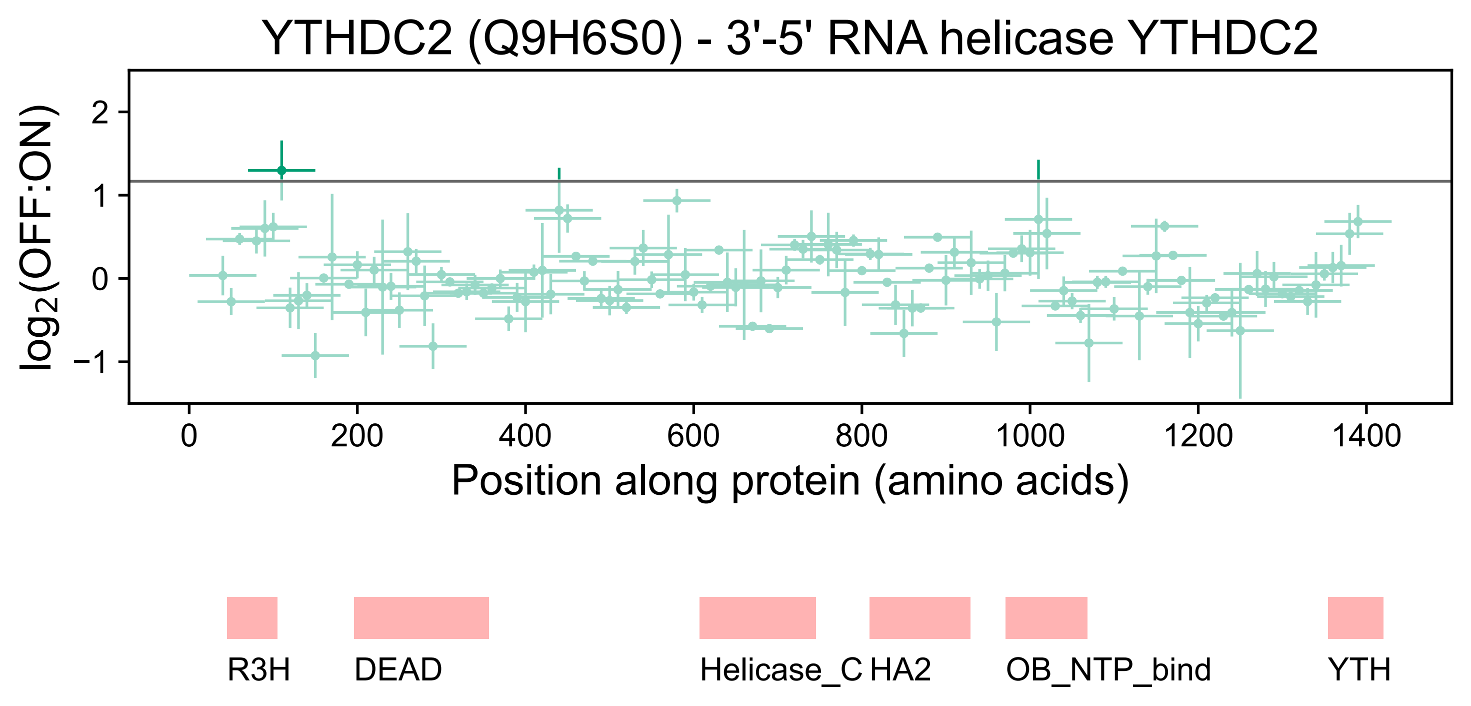
YTHDC2 (Q9H6S0) - 3'-5' RNA helicase YTHDC2
log2(OFF:ON)
2
1
0
−1
0
200
400
600
800
1000
1200
1400
Position along protein (amino acids)
R3H
DEAD
Helicase_C
HA2
OB_NTP_bind
YTH

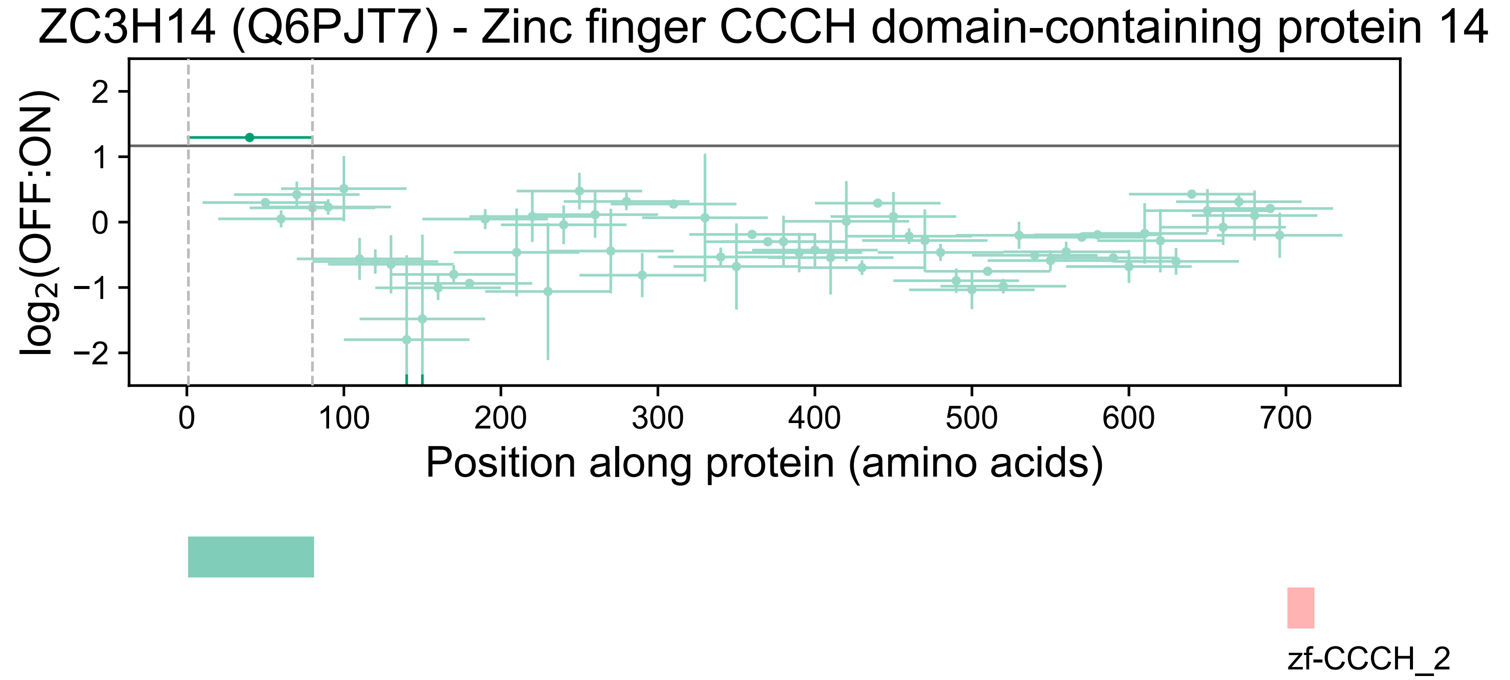
ZC3H14 (Q6PJT7) - Zinc finger CCCH domain-containing protein 14
log2(OFF:ON)
2
1
0
−1
−2
0
100
200
300
400
500
600
700
Position along protein (amino acids)
zf-CCCH_2

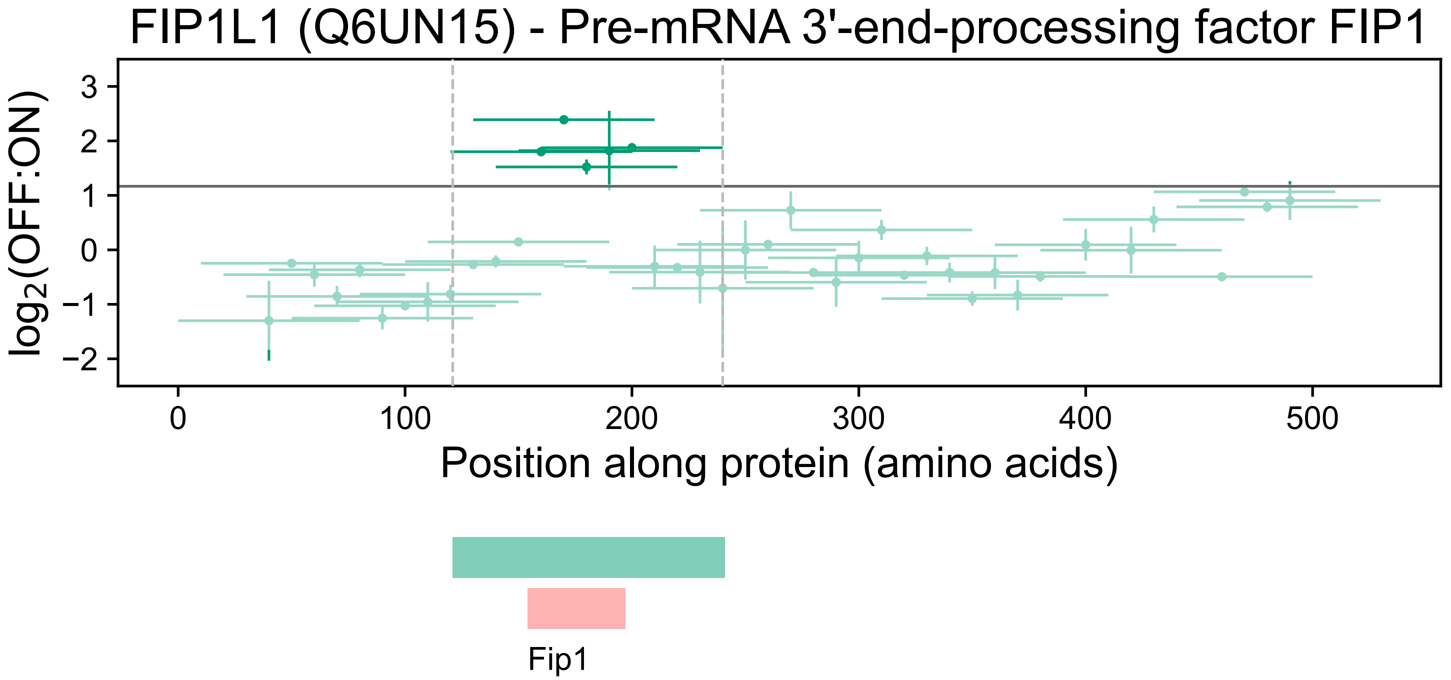

FIP1L1 (Q6UN15) - Pre-mRNA 3'-end-processing factor FIP1
log2(OFF:ON)
3
2
1
0
−1
−2
0
100
200
300
400
500
Position along protein (amino acids)
Fip1

### DGCR8 (Q8WYQ5) - Microprocessor complex subunit DGCR8

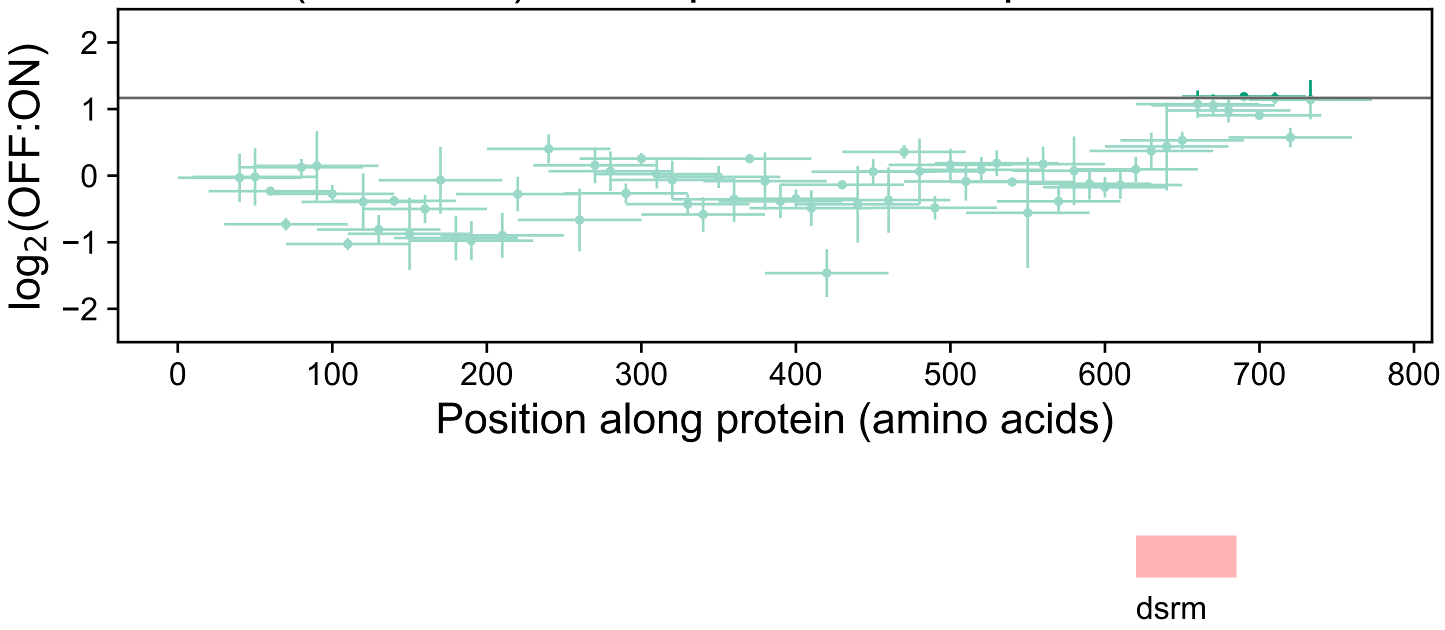

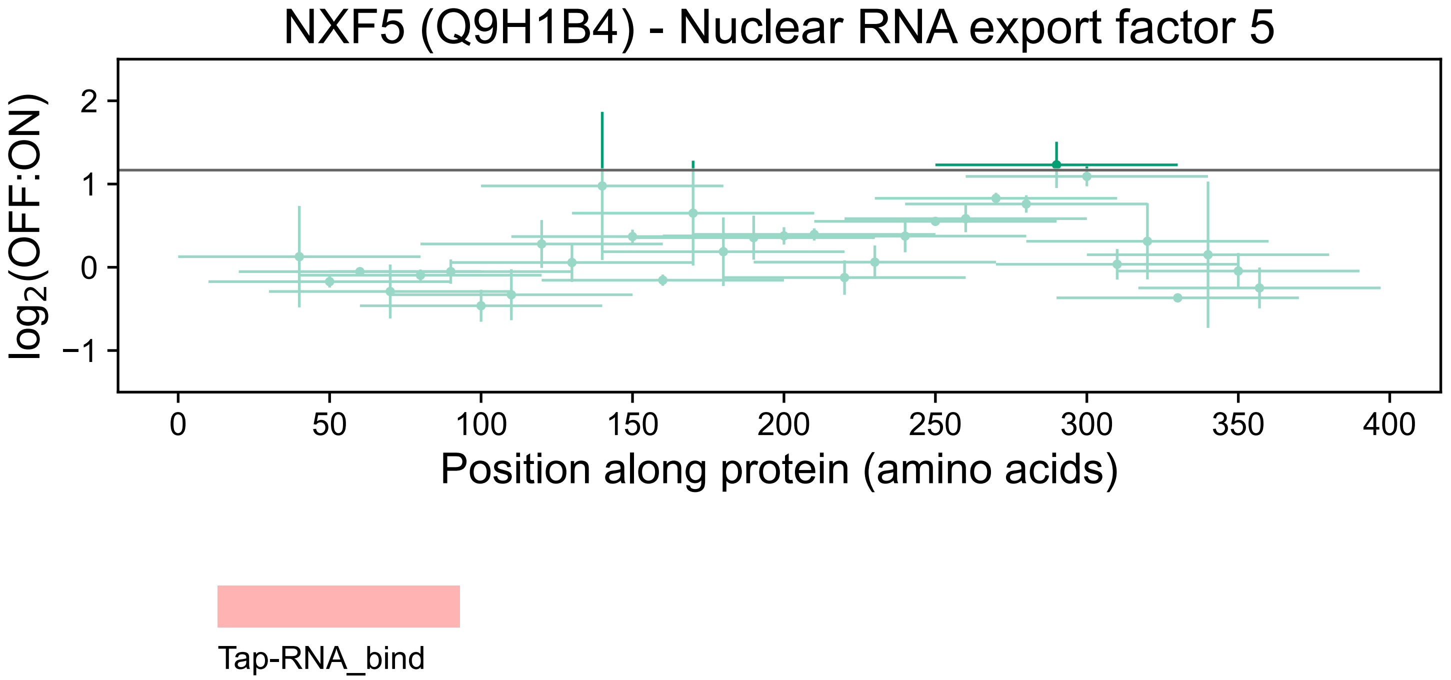
NXF5 (Q9H1B4) - Nuclear RNA export factor 5
log2(OFF:ON)
2
1
0
−1
0
50
100
150
200
250
300
350
400
Position along protein (amino acids)
Tap-RNA_bind

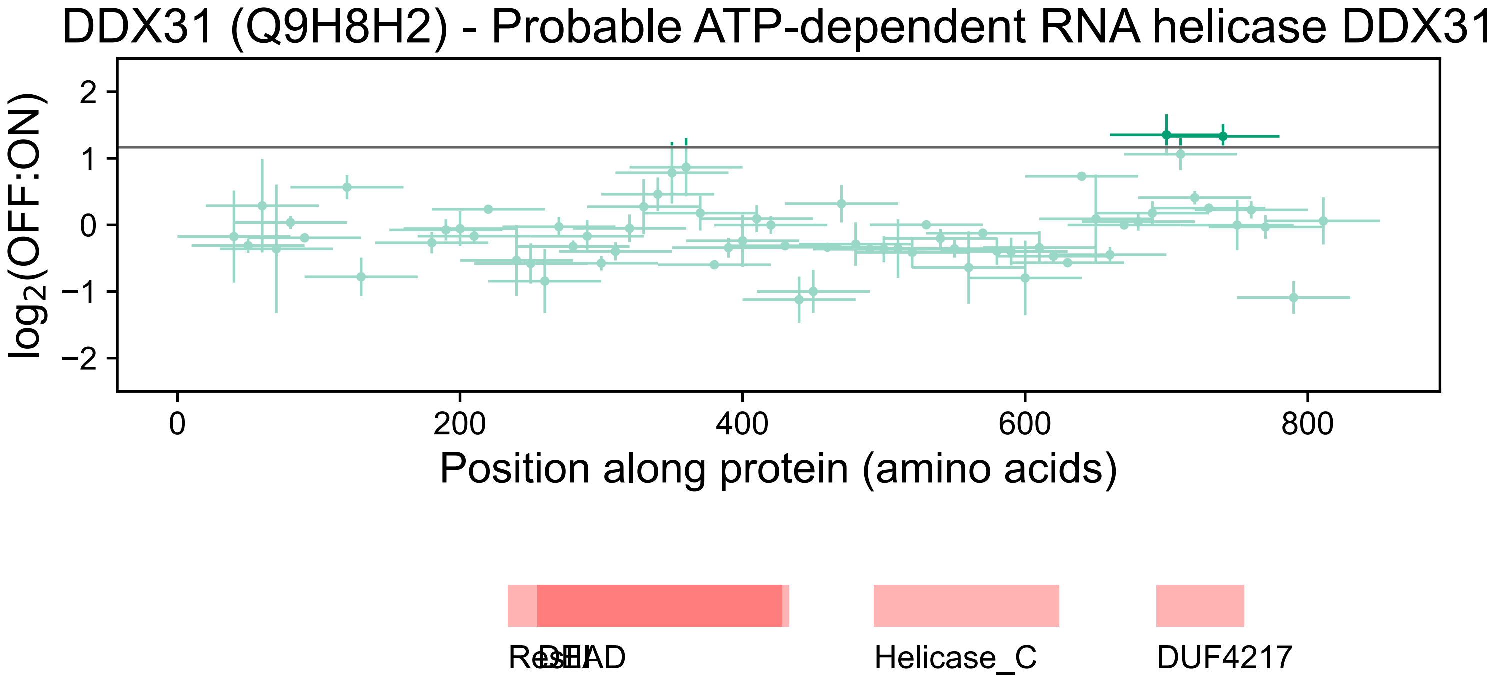
DDX31 (Q9H8H2) - Probable ATP-dependent RNA helicase DDX31
log2(OFF:ON)
2
1
0
−1
−2
0
200
400
600
800
Position along protein (amino acids)
ResIII
DEAD
Helicase_C
DUF4217

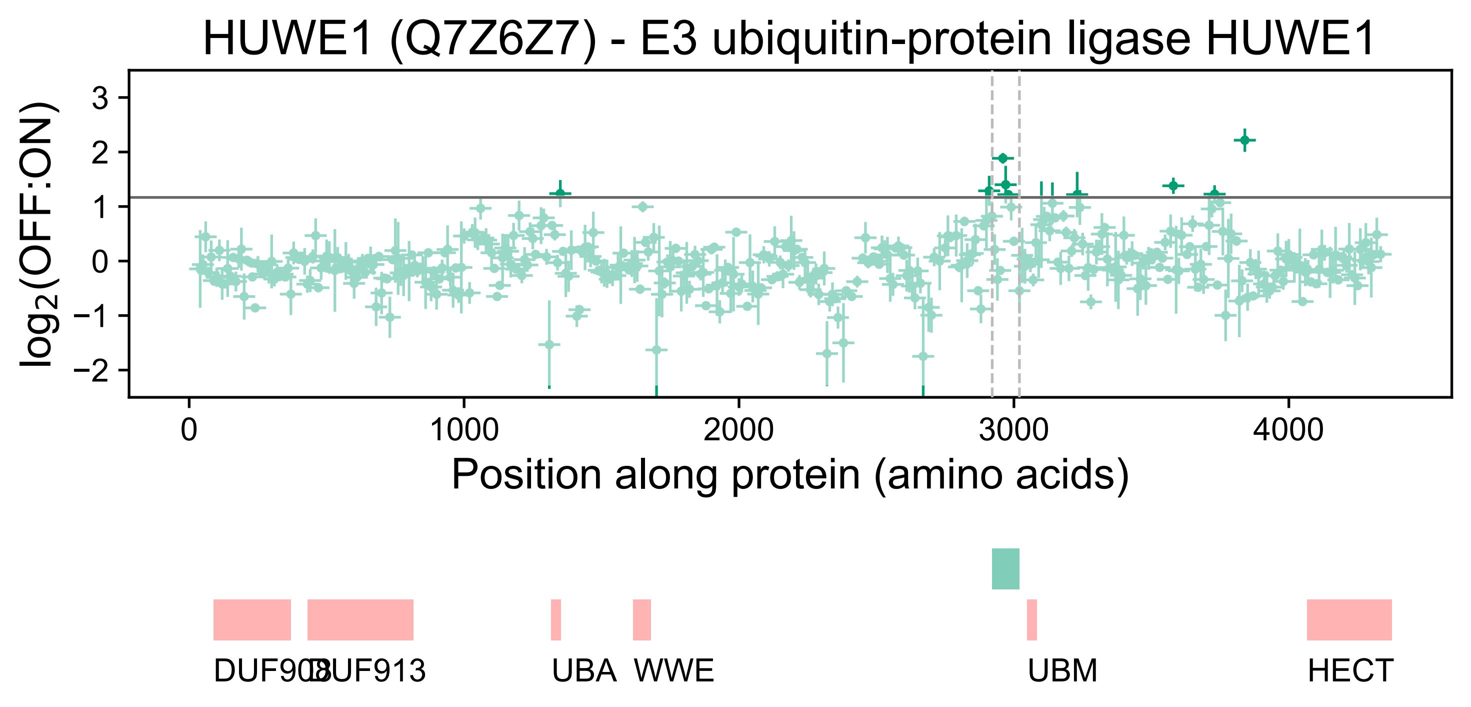

HUWE1 (Q7Z6Z7) - E3 ubiquitin-protein ligase HUWE1
log2(OFF:ON)
3
2
1
0
−1
−2
0
1000
2000
3000
4000
Position along protein (amino acids)
DUF908
DUF913
UBA
WWE
UBM
HECT

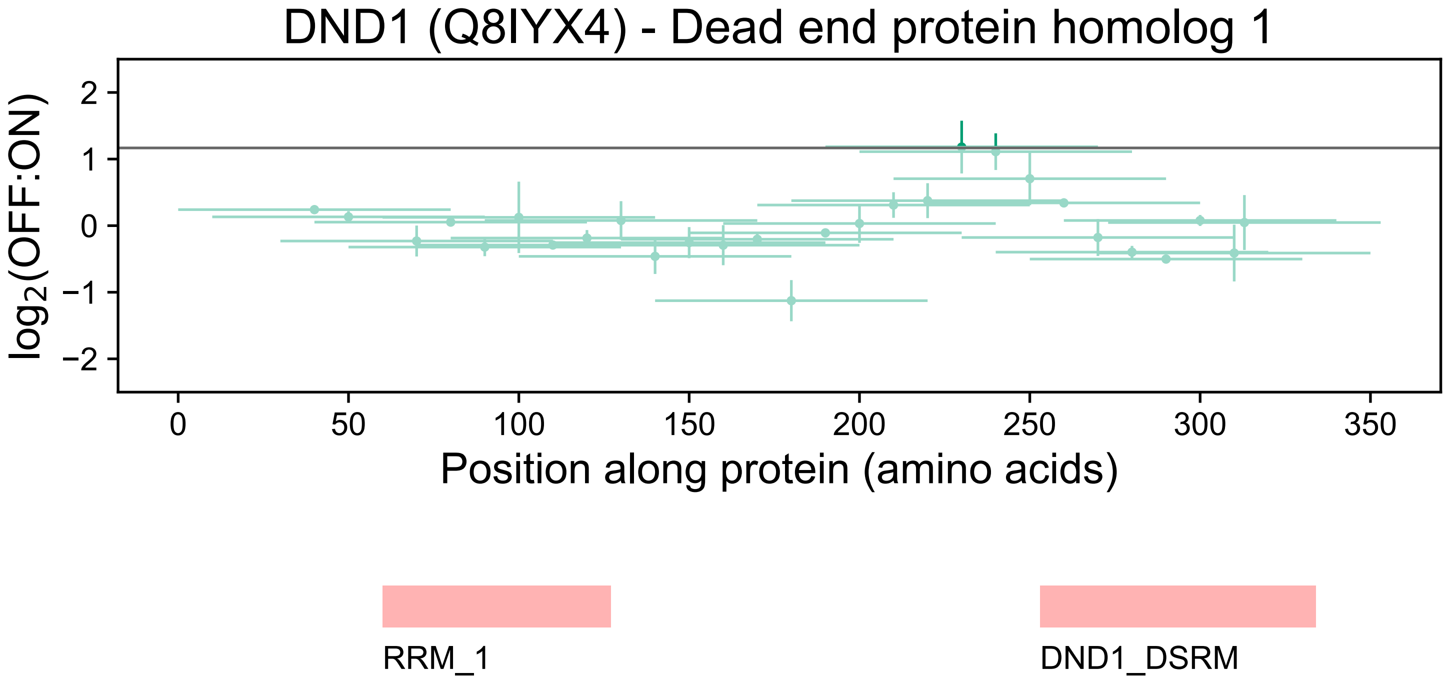
DND1 (Q8IYX4) - Dead end protein homolog 1
log2(OFF:ON)
2
1
0
−1
−2
0
50
100
150
200
250
300
350
Position along protein (amino acids)
RRM_1
DND1_DSRM

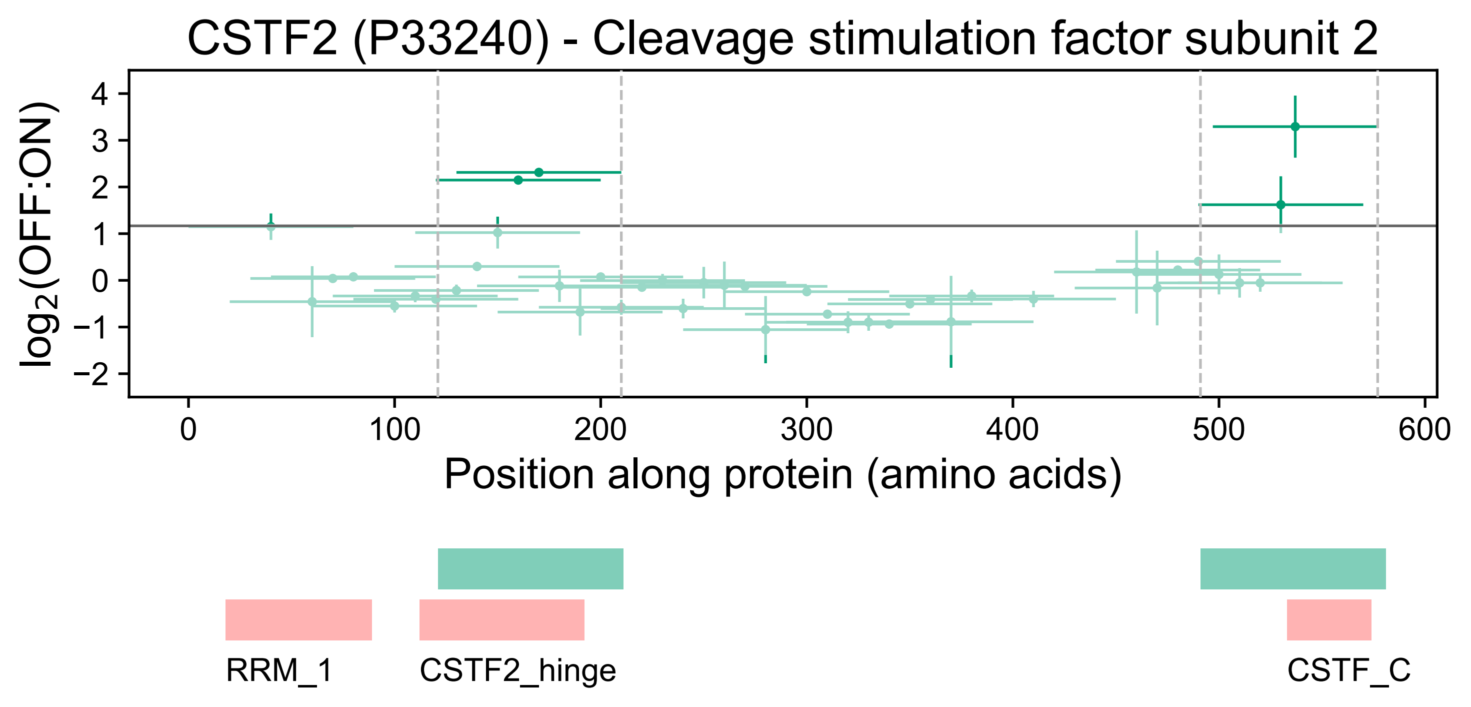

CSTF2 (P33240) - Cleavage stimulation factor subunit 2
log2(OFF:ON)
4
3
2
1
0
−1
−2
0
100
200
300
400
500
600
Position along protein (amino acids)
RRM_1
CSTF2_hinge
CSTF_C

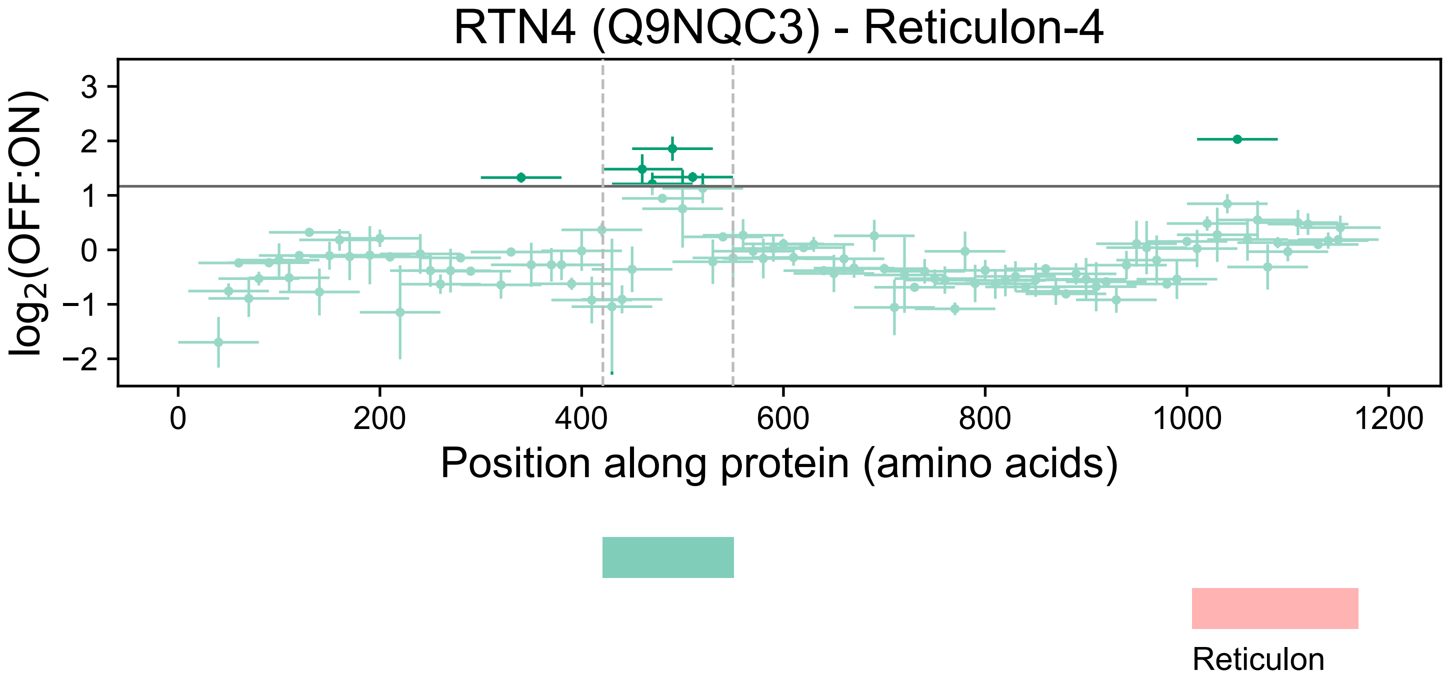
RTN4 (Q9NQC3) - Reticulon-4
log2(OFF:ON)
3
2
1
0
−1
−2
0
200
400
600
800
1000
1200
Position along protein (amino acids)
Reticulon

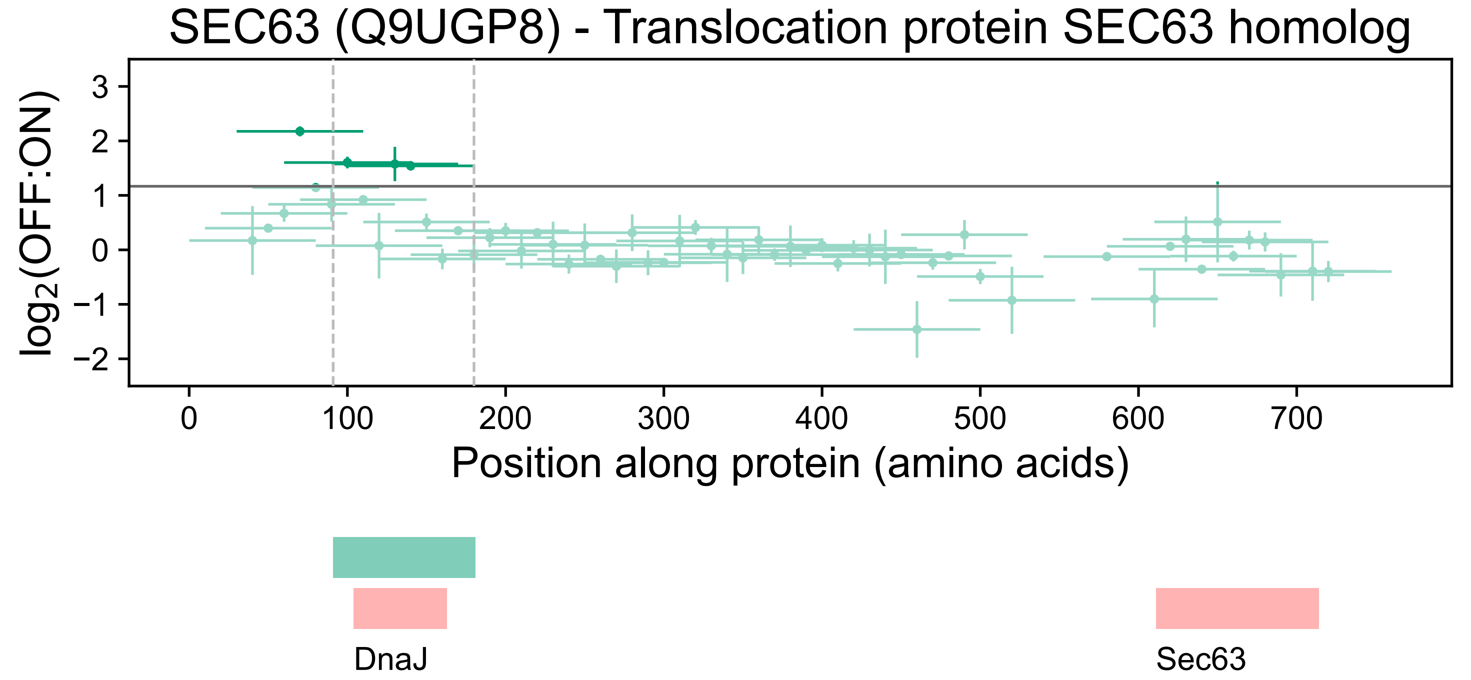
SEC63 (Q9UGP8) - Translocation protein SEC63 homolog
log2(OFF:ON)
3
2
1
0
−1
−2
0
100
200
300
400
500
600
700
Position along protein (amino acids)
DnaJ
Sec63

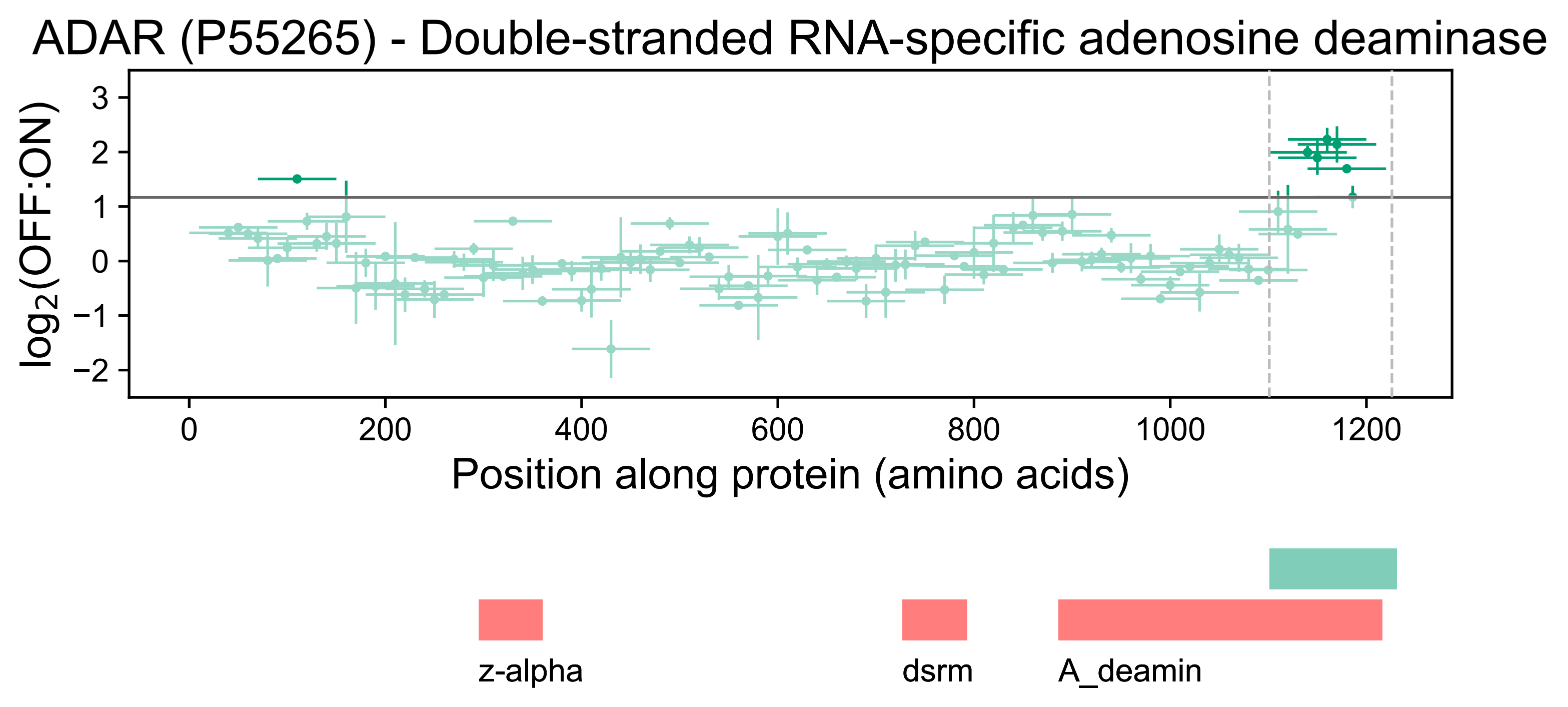
ADAR (P55265) - Double-stranded RNA-specific adenosine deaminase
log2(OFF:ON)
3
2
1
0
−1
−2
0
200
400
600
800
1000
1200
Position along protein (amino acids)
z-alpha
dsrm
A_deamin

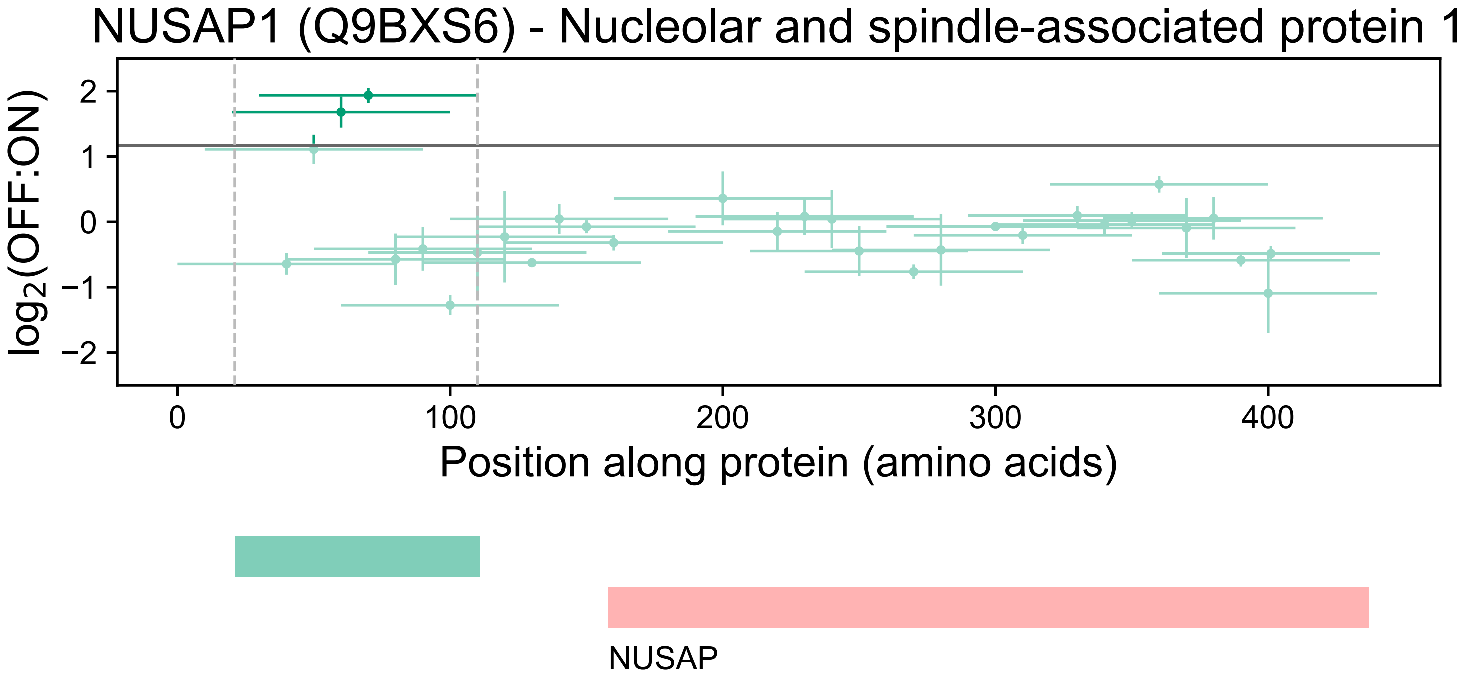
NUSAP1 (Q9BXS6) - Nucleolar and spindle-associated protein 1
log2(OFF:ON)
2
1
0
−1
−2
0
100
200
300
400
Position along protein (amino acids)
NUSAP

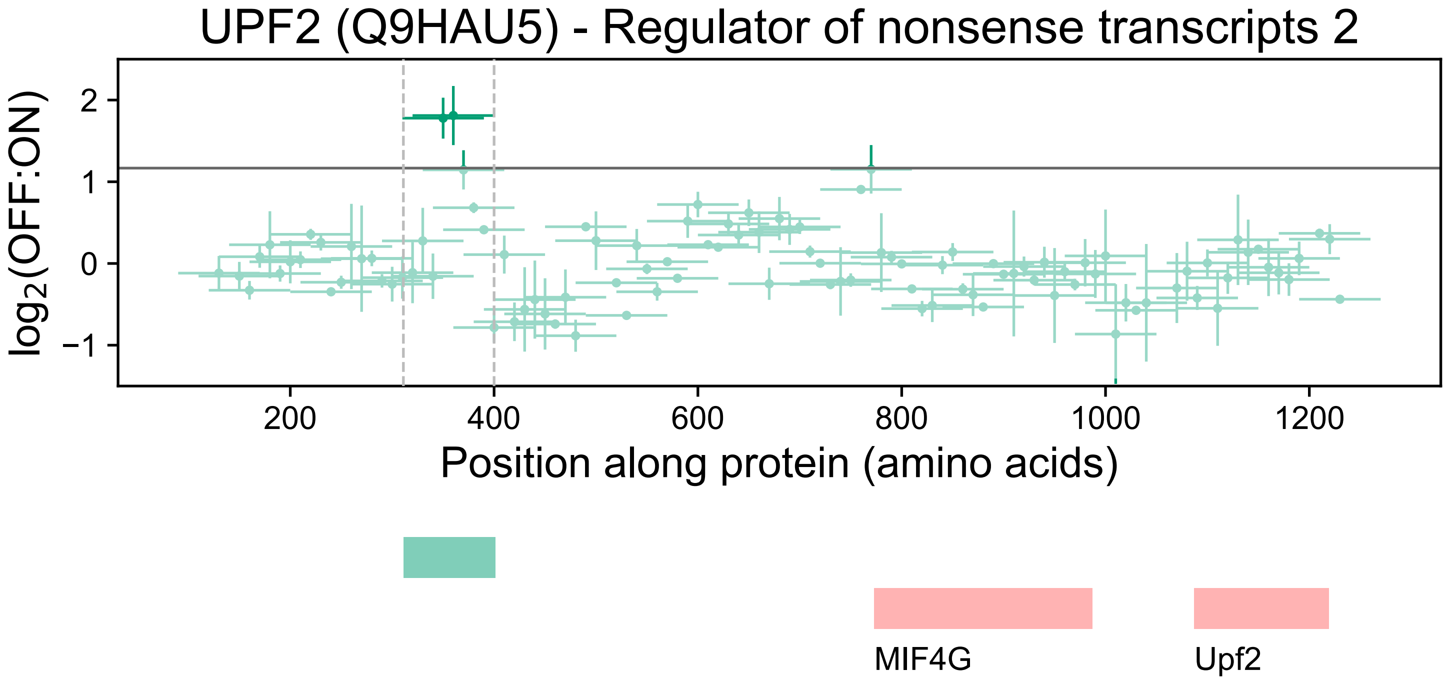

UPF2 (Q9HAU5) - Regulator of nonsense transcripts 2
log2(OFF:ON)
2
1
0
−1
200
400
600
800
1000
1200
Position along protein (amino acids)
MIF4G
Upf2

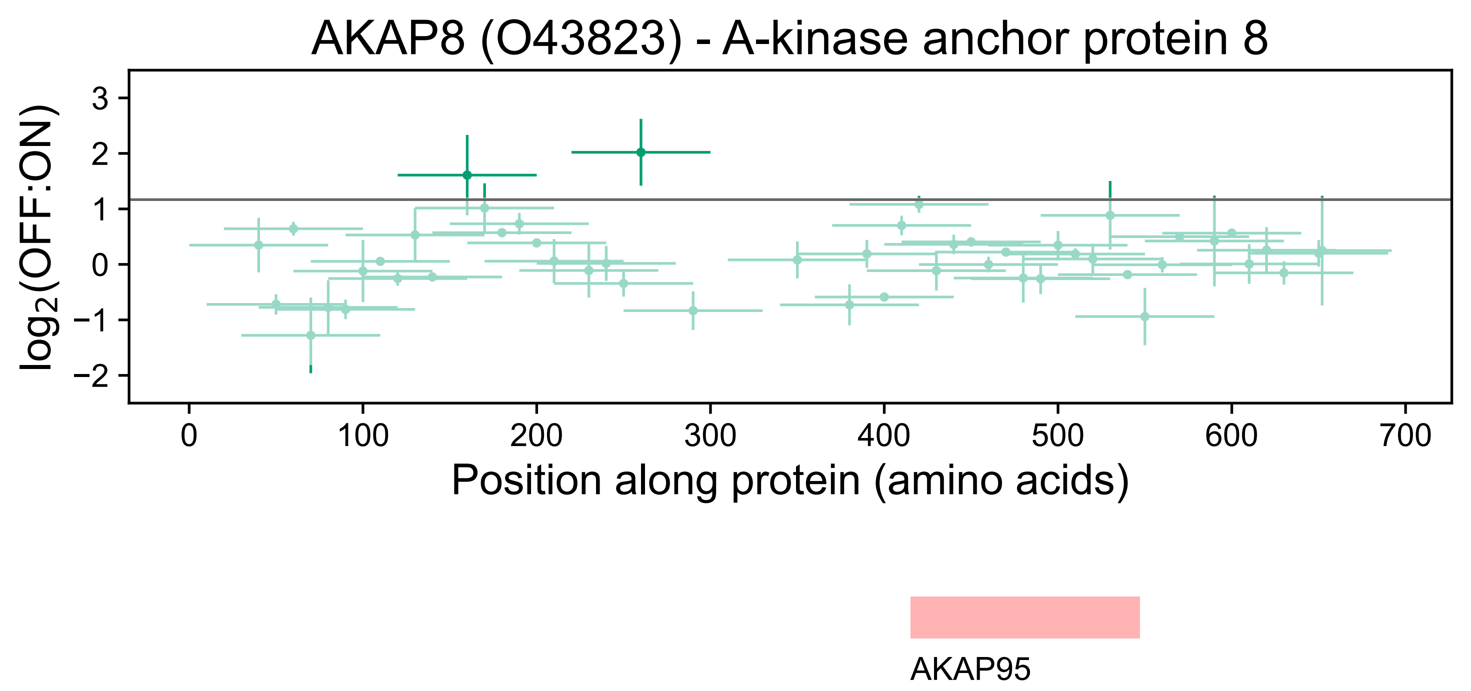

AKAP8 (O43823) - A-kinase anchor protein 8
log2(OFF:ON)
3
2
1
0
−1
−2
0
100
200
300
400
500
600
700
Position along protein (amino acids)
AKAP95

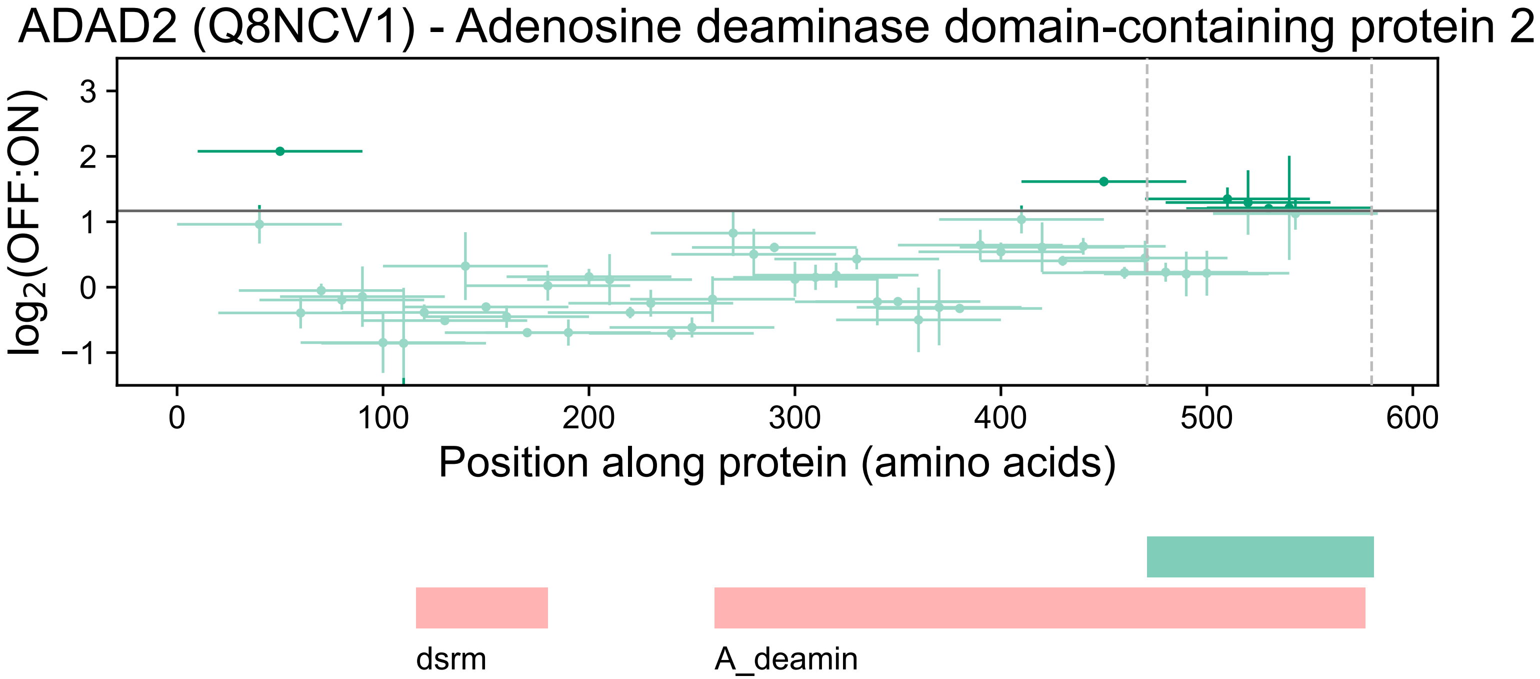
ADAD2 (Q8NCV1) - Adenosine deaminase domain-containing protein 2
log2(OFF:ON)
3
2
1
0
−1
0
100
200
300
400
500
600
Position along protein (amino acids)
dsrm
A_deamin

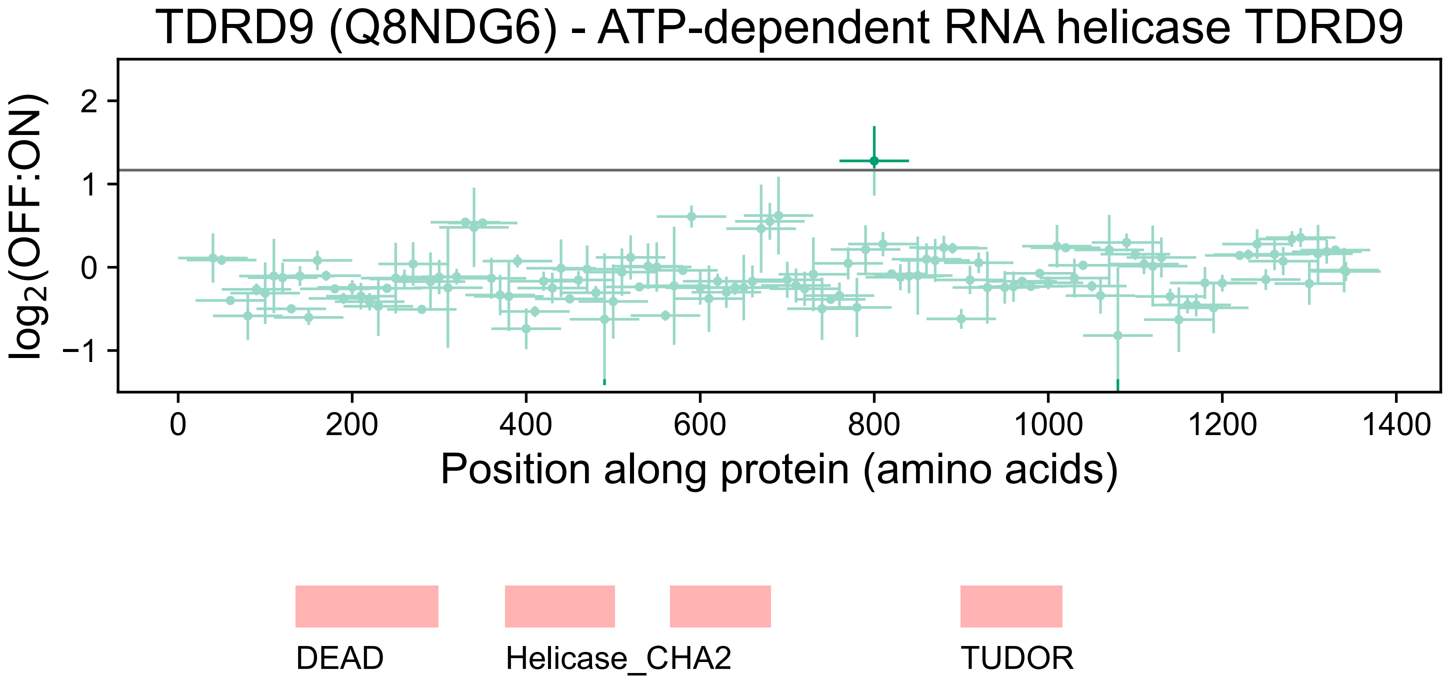

TDRD9 (Q8NDG6) - ATP-dependent RNA helicase TDRD9
$log_2$(OFF:ON)
2
1
0
−1
0
200
400
600
800
1000
1200
1400
Position along protein (amino acids)
DEAD
Helicase_CHA2
TUDOR

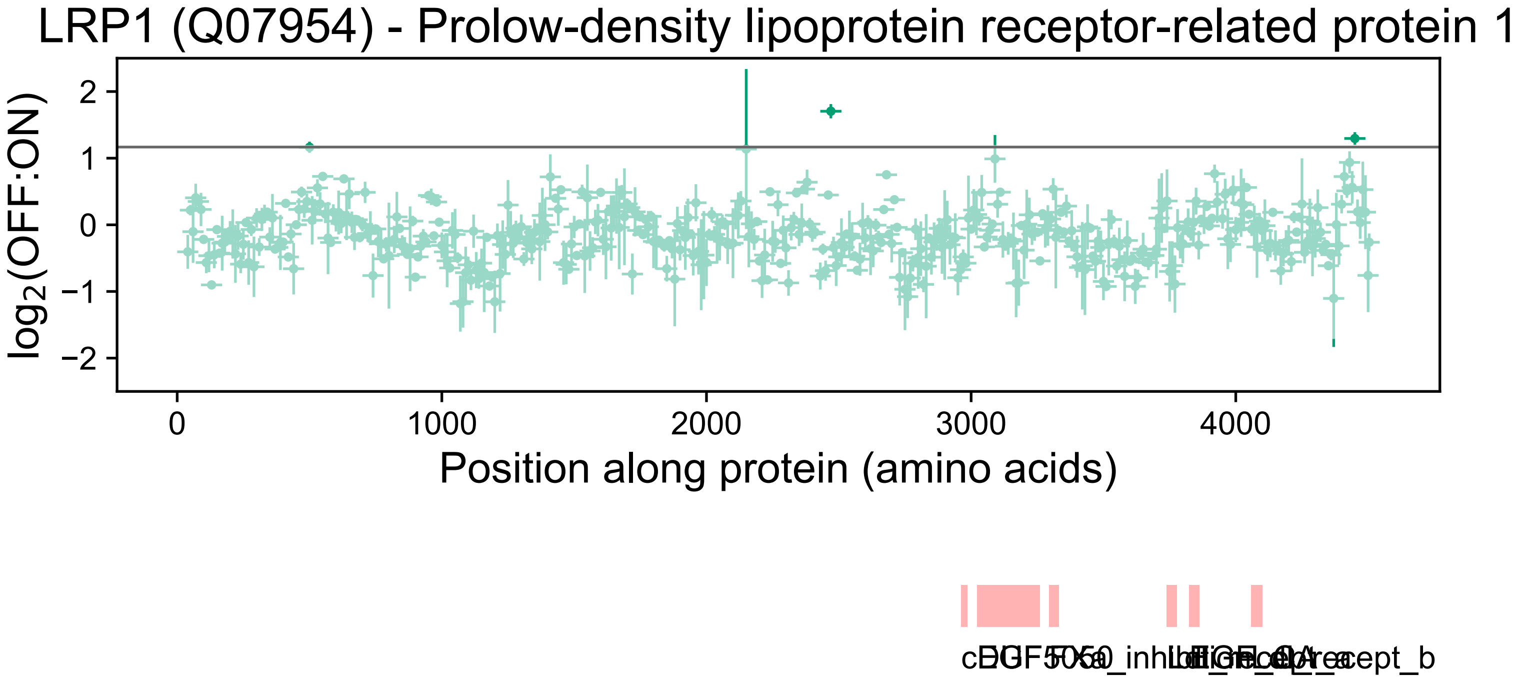
LRP1 (Q07954) - Prolow-density lipoprotein receptor-related protein 1
log2(OFF:ON)
2
1
0
−1
−2
0
1000
2000
3000
4000
Position along protein (amino acids)

### ZC3H11A (O75152) - Zinc finger CCCH domain-containing protein 11A

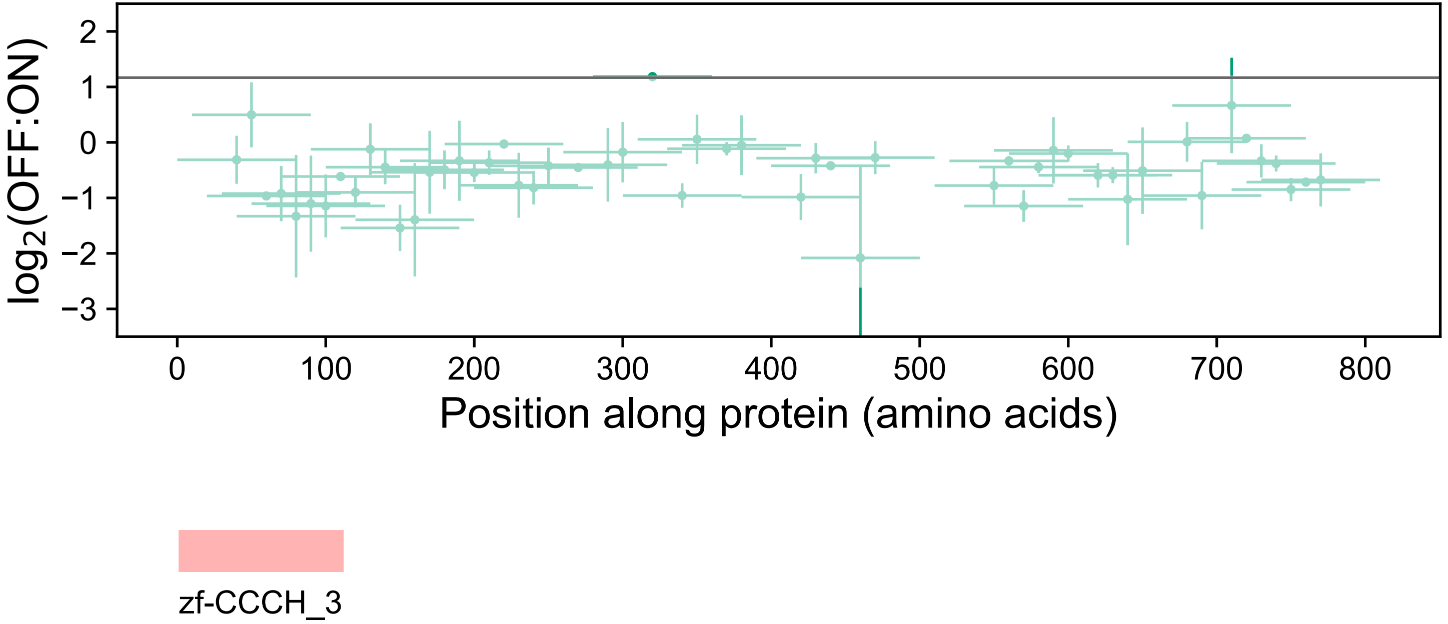

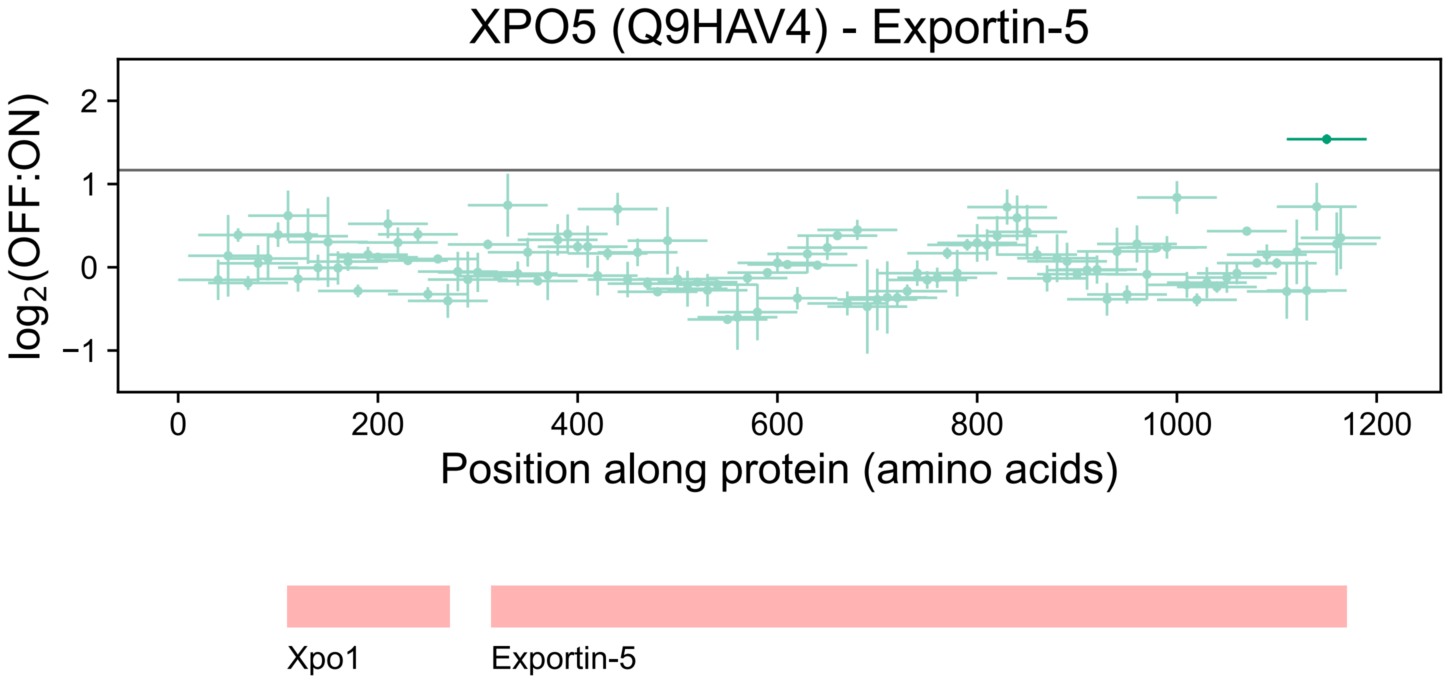
XPO5 (Q9HAV4) - Exportin-5
log2(OFF:ON)
2
1
0
−1
0
200
400
600
800
1000
1200
Position along protein (amino acids)
Xpo1
Exportin-5

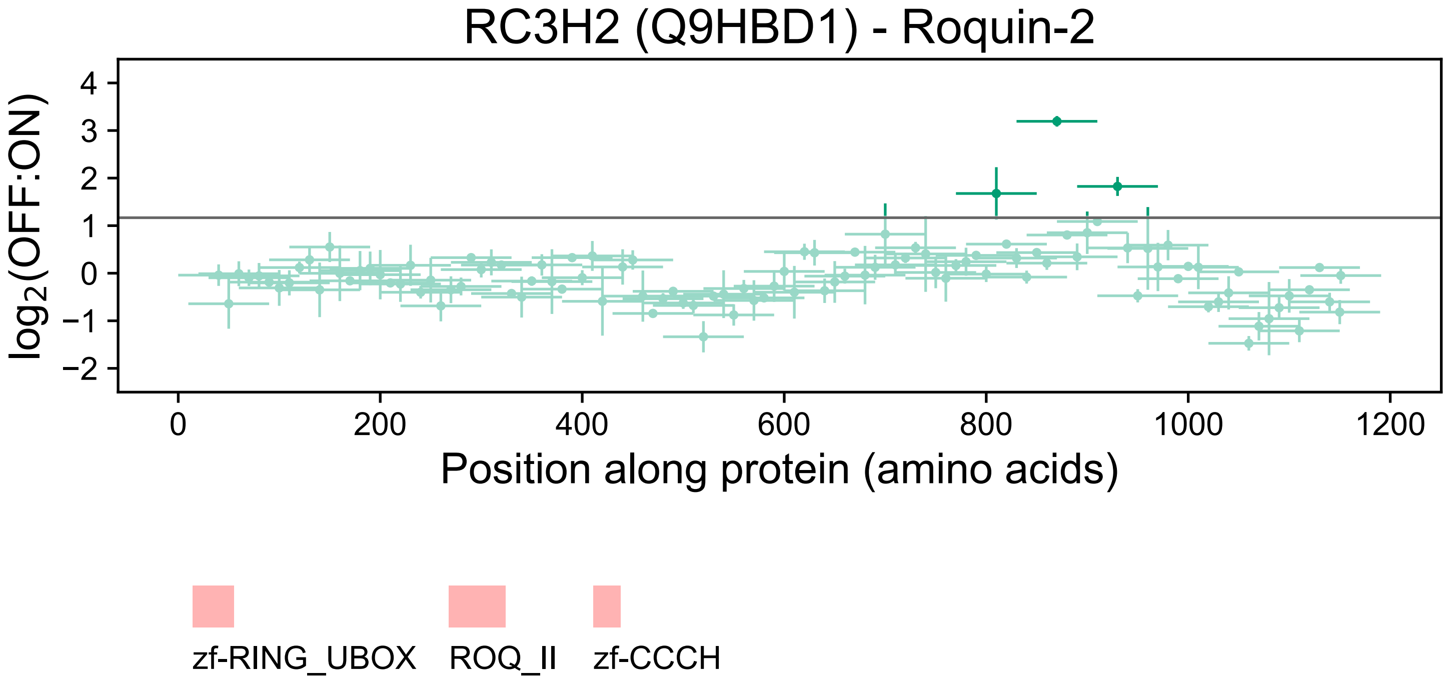
RC3H2 (Q9HBD1) - Roquin-2
log2(OFF:ON)
4
3
2
1
0
−1
−2
0
200
400
600
800
1000
1200
Position along protein (amino acids)
zf-RING_UBOX
ROQ_II
zf-CCCH

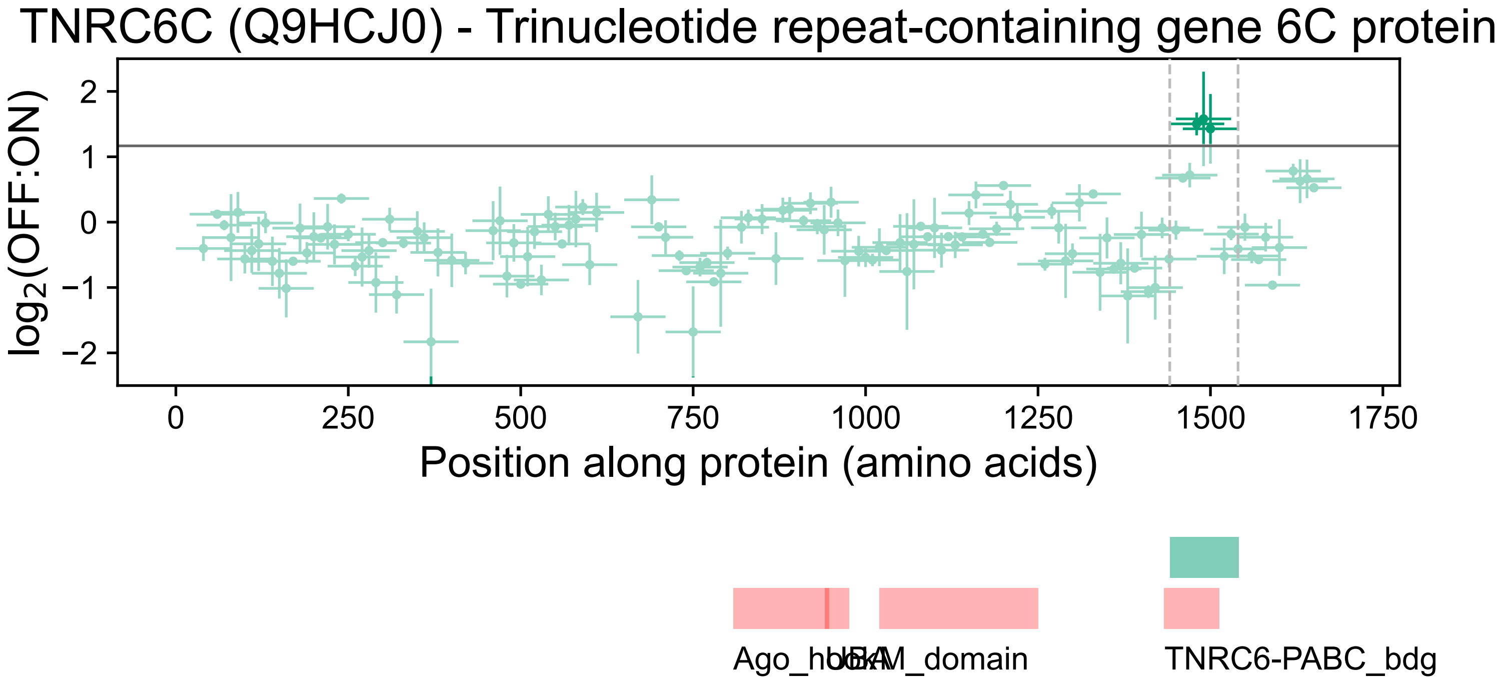

TNRC6C (Q9HCJ0) - Trinucleotide repeat-containing gene 6C protein
log2(OFF:ON)
2
1
0
−1
−2
0
250
500
750
1000
1250
1500
1750
Position along protein (amino acids)
Ago_hook
UBA
M_domain
TNRC6-PABC_bdg

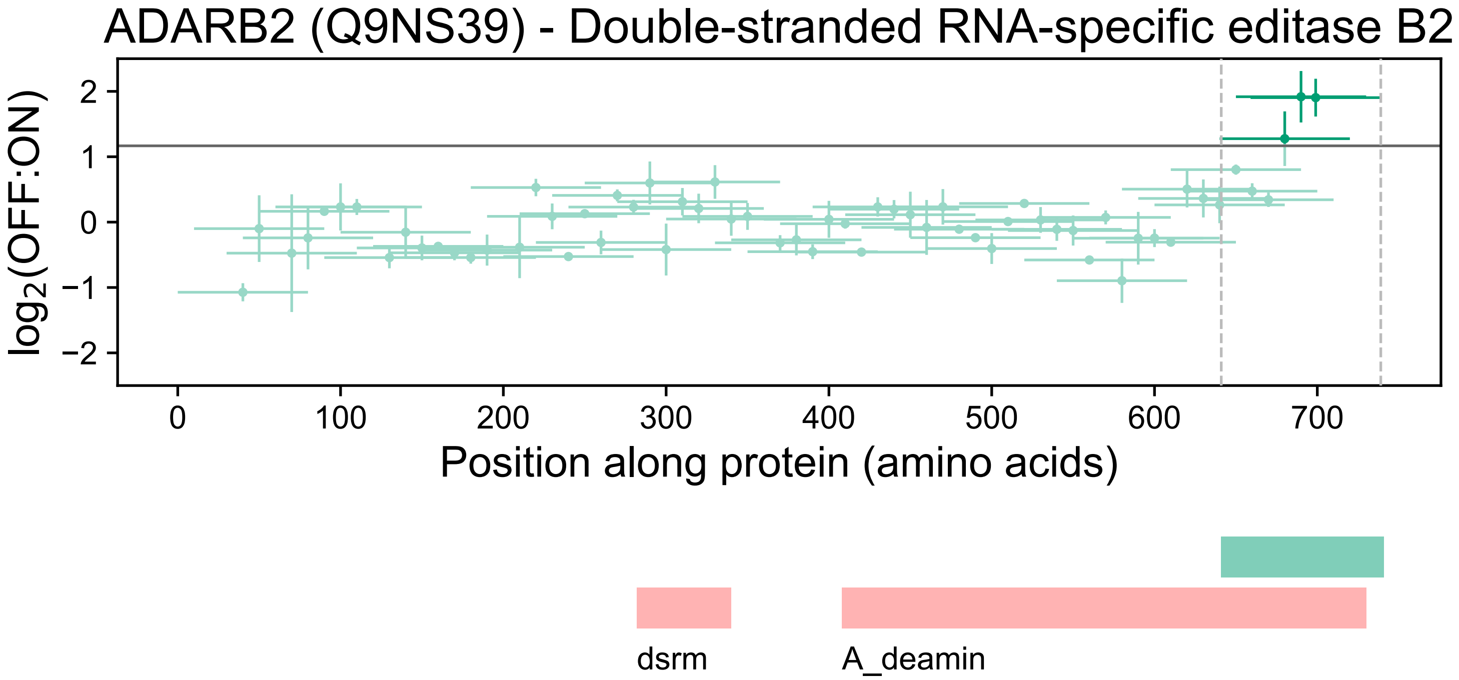

ADARB2 (Q9NS39) - Double-stranded RNA-specific editase B2
log2(OFF:ON)
2
1
0
−1
−2
0
100
200
300
400
500
600
700
Position along protein (amino acids)
dsrm
A_deamin

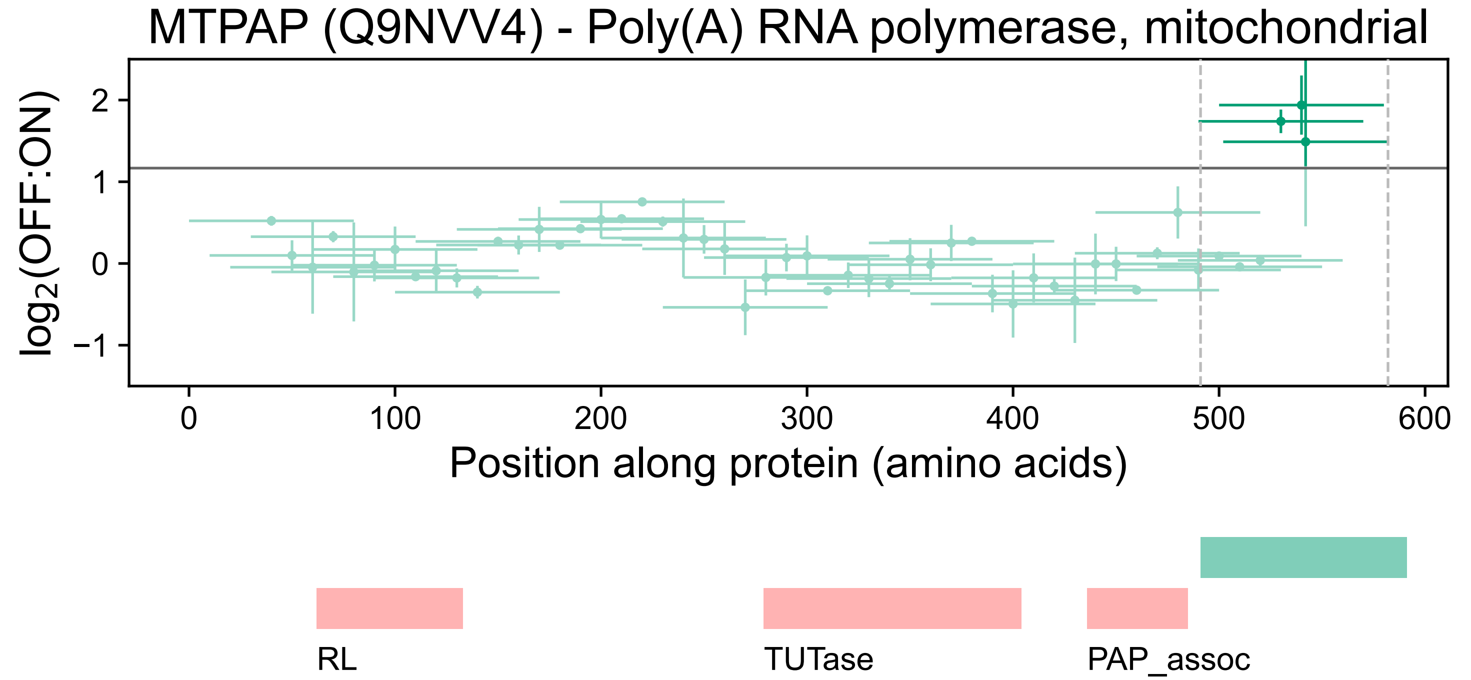
MTPAP (Q9NVV4) - Poly(A) RNA polymerase, mitochondrial
log2(OFF:ON)
2
1
0
−1
0
100
200
300
400
500
600
Position along protein (amino acids)
RL
TUTase
PAP_assoc

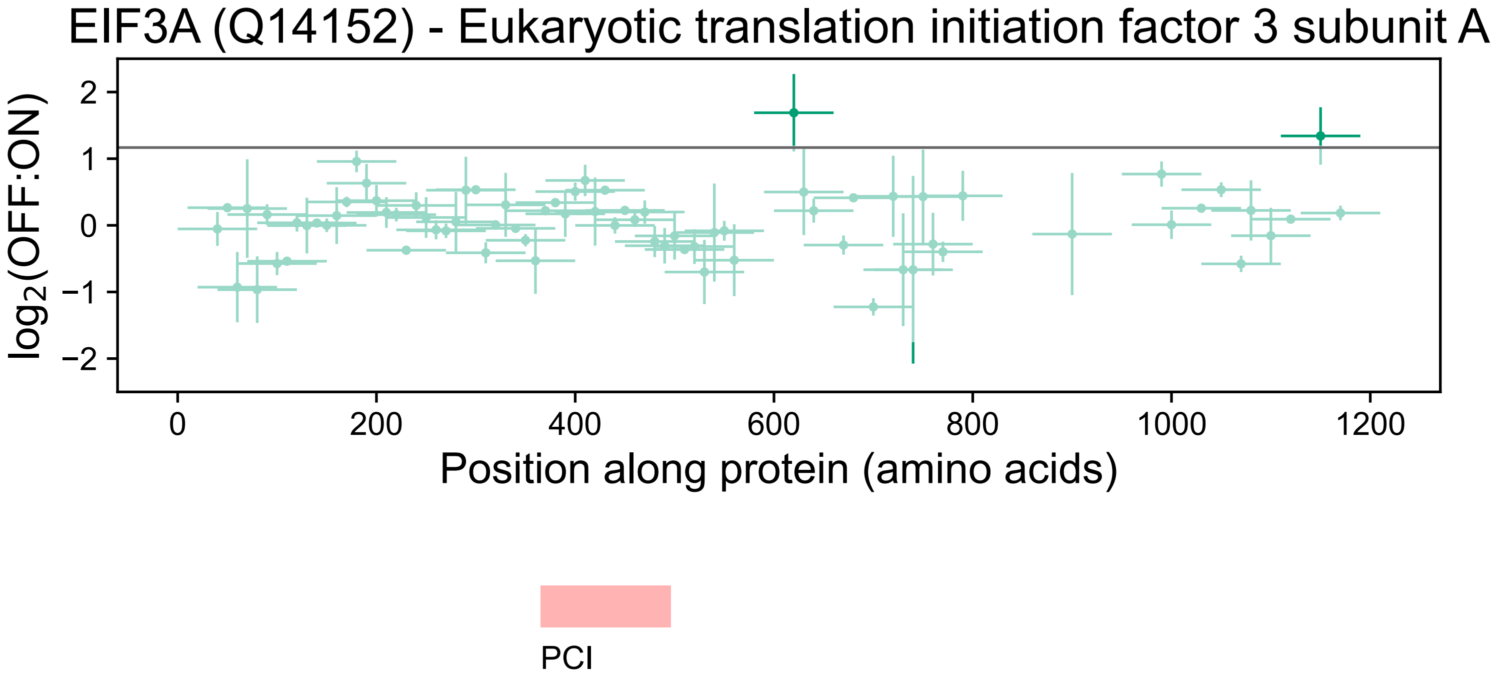

EIF3A (Q14152) - Eukaryotic translation initiation factor 3 subunit A
log2(OFF:ON)
2
1
0
−1
−2
0
200
400
600
800
1000
1200
Position along protein (amino acids)
PCI

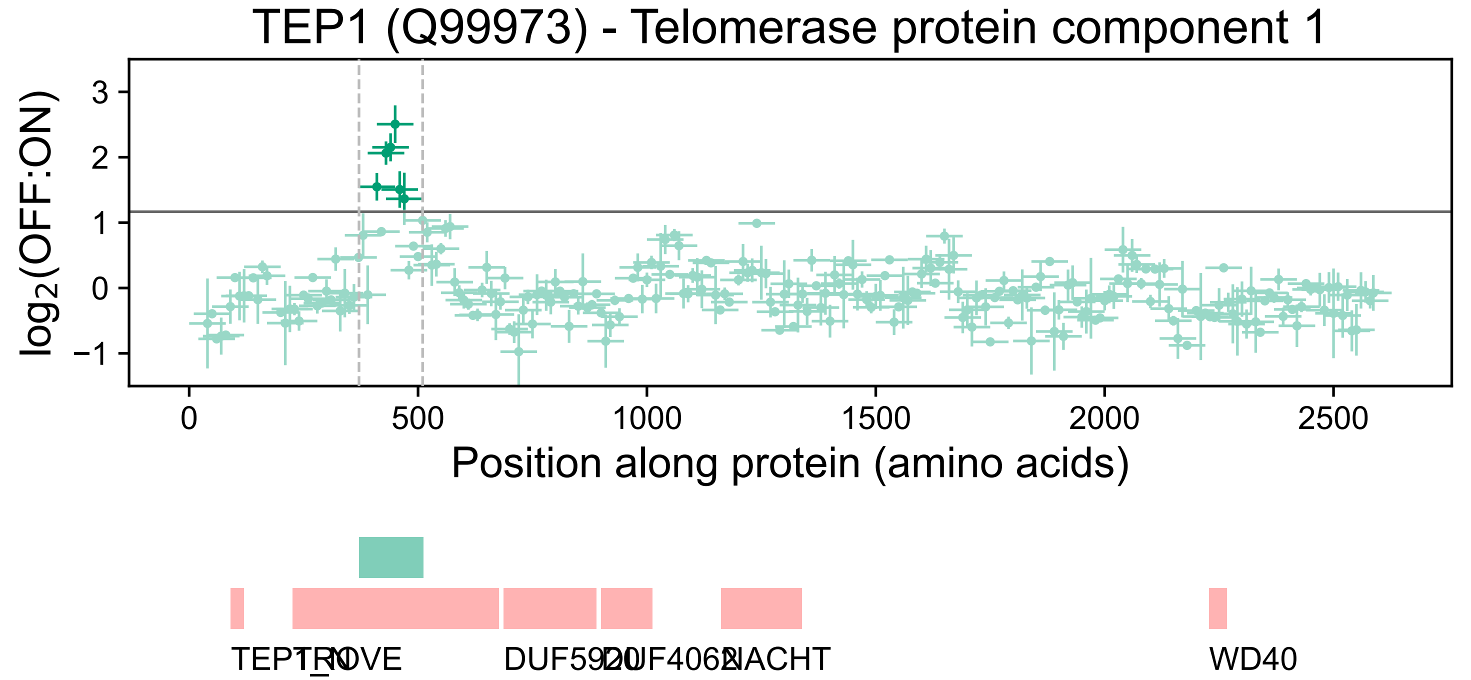

TEP1 (Q99973) - Telomerase protein component 1
log2(OFF:ON)
3
2
1
0
−1
0
500
1000
1500
2000
2500
Position along protein (amino acids)
TEP1_N
TROVE
DUF5920
DUF4062
NACHT
WD40

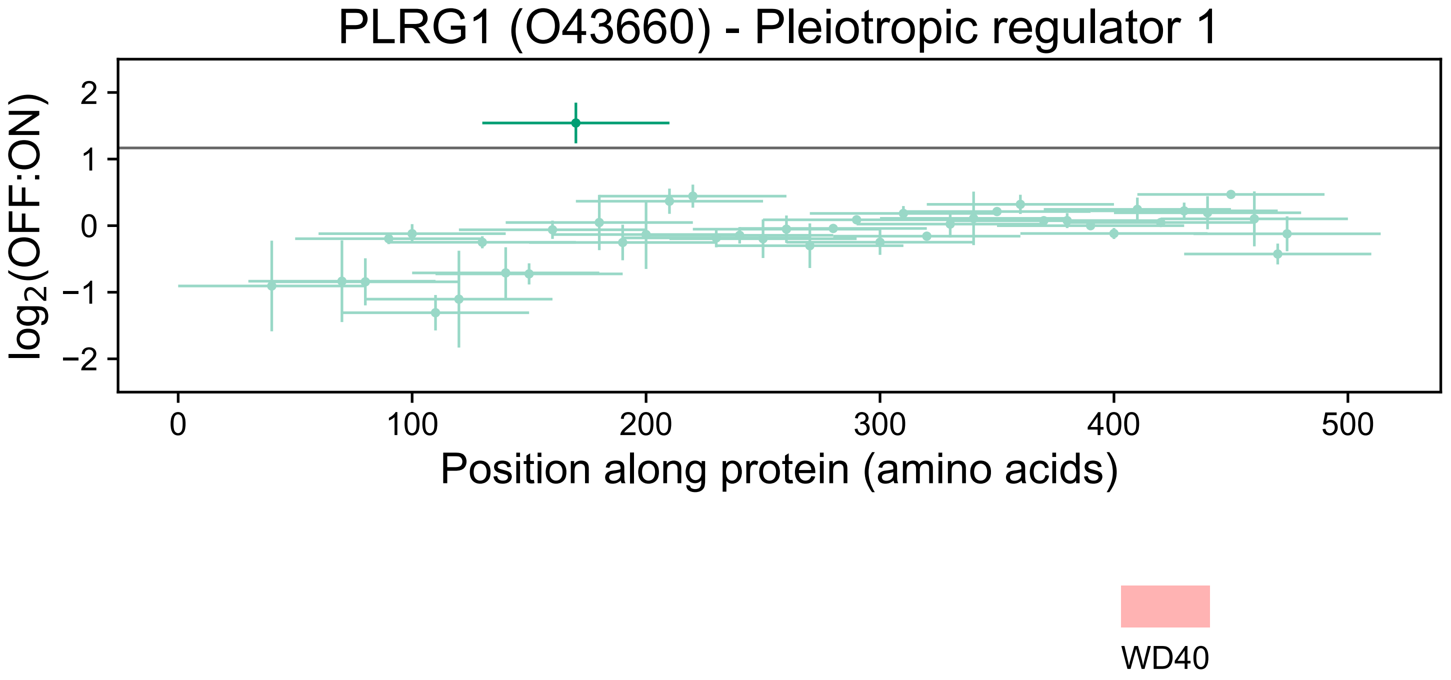

PLRG1 (O43660) - Pleiotropic regulator 1
log2(OFF:ON)
2
1
0
−1
−2
0
100
200
300
400
500
Position along protein (amino acids)
WD40

### RCC2 (Q9P258) - Protein RCC2

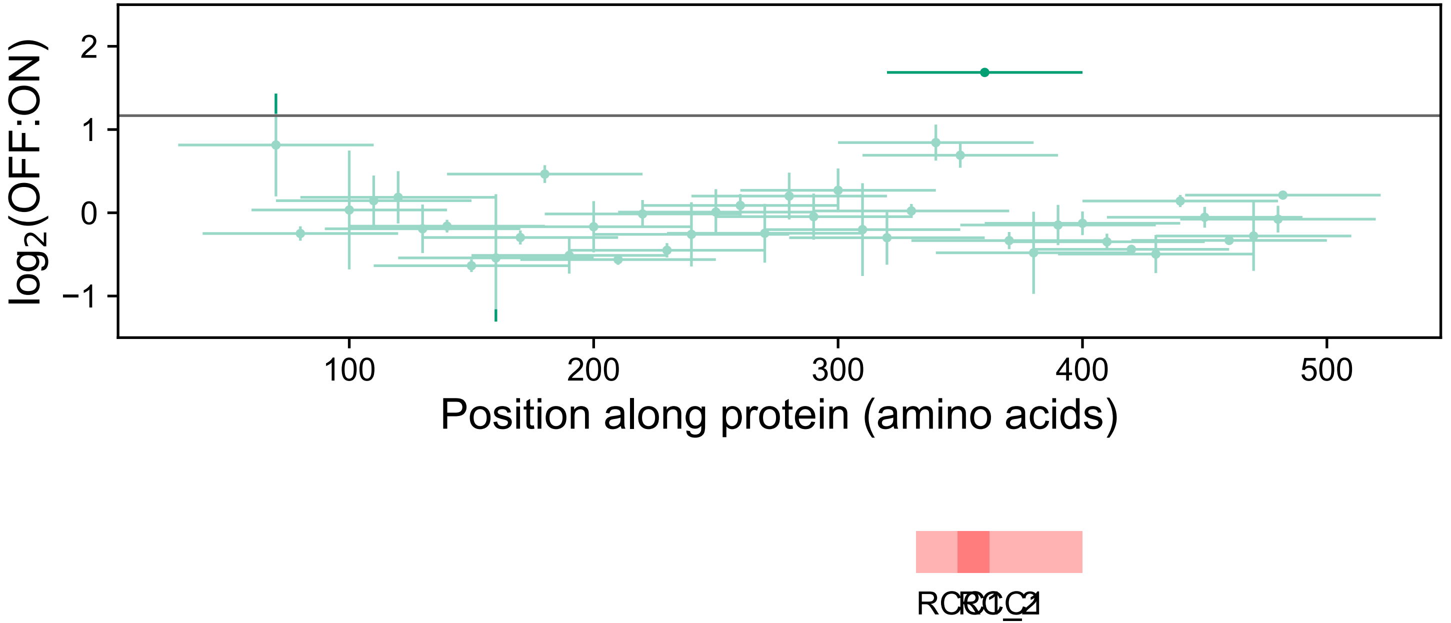

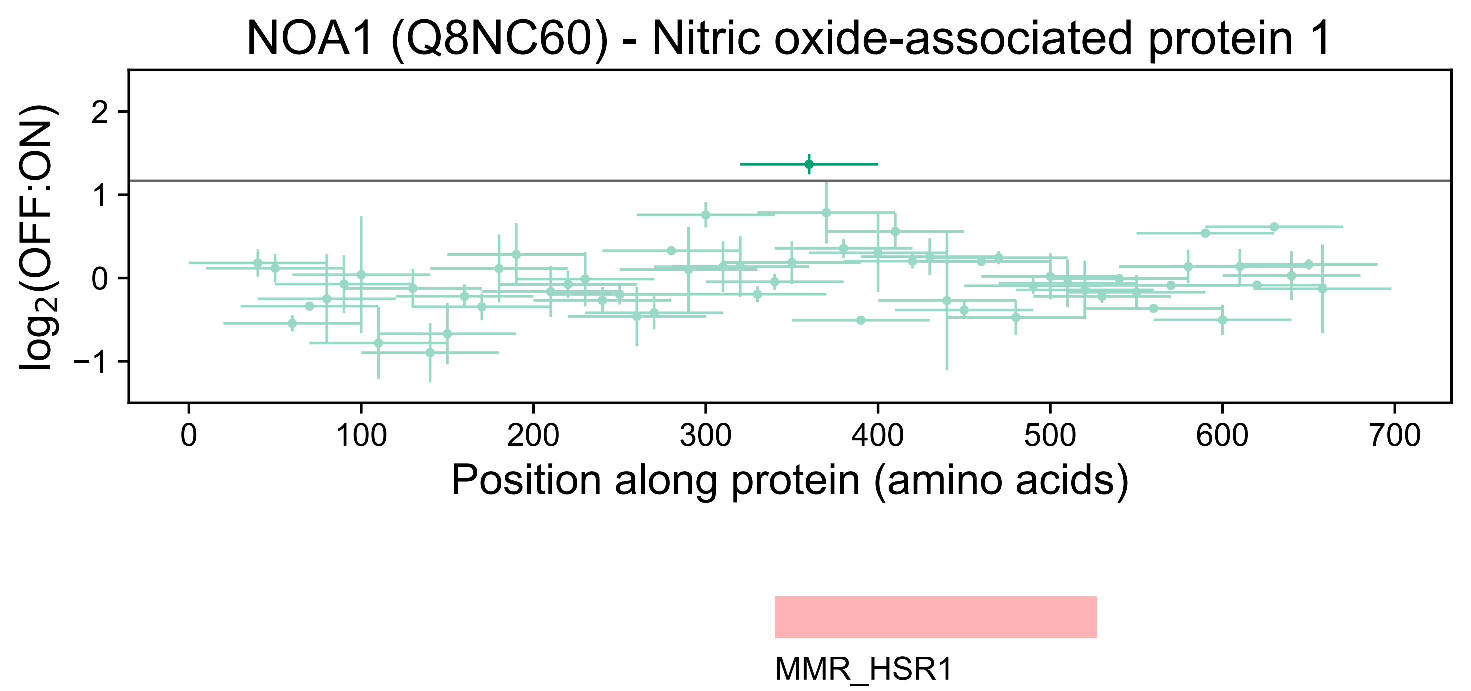
NOA1 (Q8NC60) - Nitric oxide-associated protein 1
log2(OFF:ON)
2
1
0
−1
0
100
200
300
400
500
600
700
Position along protein (amino acids)
MMR_HSR1

### DDX3Y (O15523) - ATP-dependent RNA helicase DDX3Y

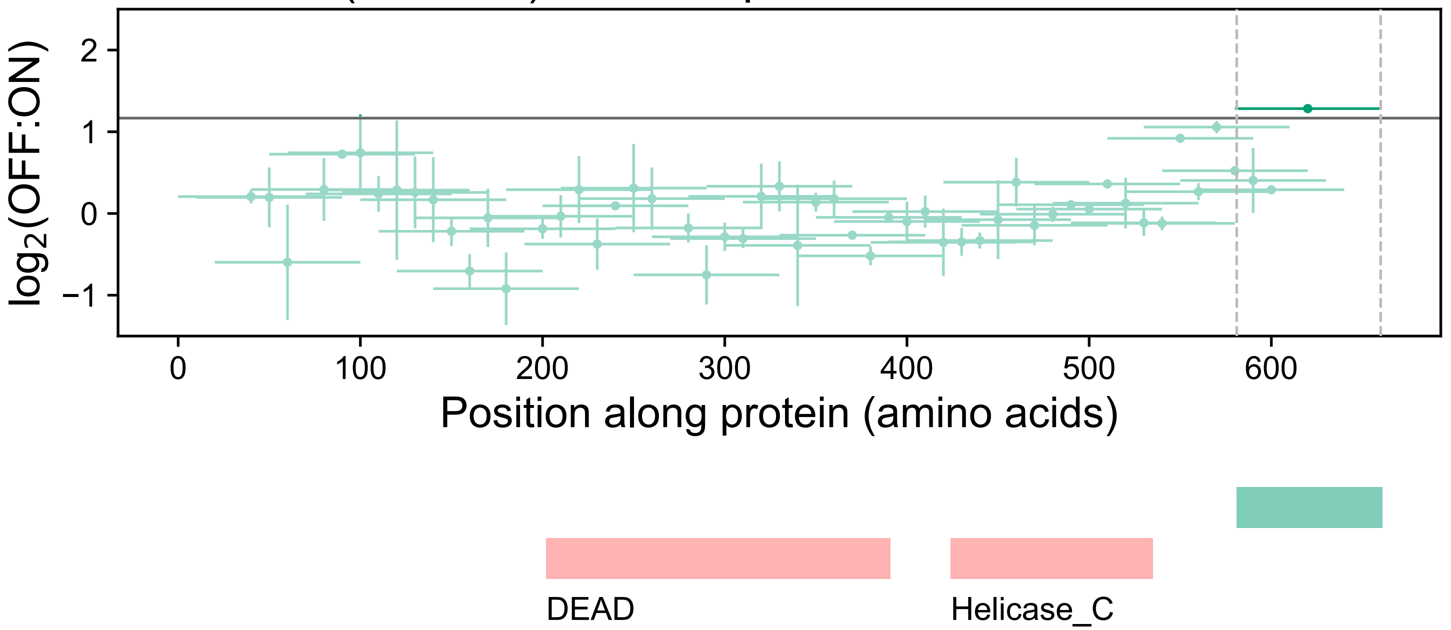
